# Trait divergence and trade-offs among Brassicaceae species differing in elevational distribution

**DOI:** 10.1101/2022.02.02.478839

**Authors:** Alessio Maccagni, Yvonne Willi

## Abstract

Species have restricted geographic distributions and the causes are still largely unknown. Temperature has long been associated with distribution limits, suggesting that there are ubiquitous constraints to the evolution of the climate niche. Here we investigated the traits involved in such constraints by macroevolutionary comparisons involving around 100 Brassicaceae species differing in elevational distribution. Plants were grown under three temperature treatments (regular frost, mild, regular heat) and phenotyped for phenological, morphological and thermal resistance traits. Trait values were analysed by assessing the effect of temperature and elevational distribution, by comparing models of evolutionary trajectories, and by correlative approaches to identify trade-offs. Analyses pointed to size, leaf morphology and growth under heat as among the most discriminating traits between low- and high-elevation species, with high-elevation species growing faster under the occurrence of regular heat bouts, at the cost of much reduced size. Mixed models and evolutionary models supported adaptive divergence for these traits, and correlation analysis indicated their involvement in moderate trade-offs. Finally, we found asymmetry in trait evolution, with evolvability across traits being 50% less constrained under regular frost. Overall, results suggest that trade-offs between traits under adaptive divergence contribute to the disparate distribution of species along the elevational gradient.

## INTRODUCTION

Species have restricted geographic distributions, but the causes behind this phenomenon are still unsolved (MacArthur 1972; Gaston 2003; Connallon and Sgrò 2018; Willi and Van Buskirk 2019). From an ecological point of view, range limits reflect dispersal limitation or limits of the ecological niche, with the niche being defined as the abiotic and biotic conditions that allow a species to persist (i.e., the realized niche *sensu* Hutchinson 1957; Leibold 1995). From an evolutionary point of view, range limits reflect limits to the evolution of the ecological niche. But why is it that species fail to adapt to environmental conditions beyond their current range? MacArthur (1972) suggested that a possible reason is exclusive divergent adaptation across habitats. He envisioned that specialization to one environment imposes high demographic costs under colonization of a new environment, or in other words, a trade-off. Trade-offs are a key concept in evolution, likely affecting all aspects of ecological specialization (Rosenzweig 1995) and including species distribution limits, but they have been rarely studied in this context.

Among the many ecological factors that may affect the persistence of organisms, climate is known to be critical in controlling large-scale distribution (MacArthur 1972). Many past studies noticed coincidences between geographic or elevational range limits and temperature isotherms (Salisbury 1926; Iversen 1944; Dahl 1951; Root 1988). More recently, the field of species distribution modelling confirmed the good agreement between range limits and climate variables (e.g., Normand et al. 2009; Lee◻Yaw et al. 2016). Further studies looked into phenotypic patterns associated with the most limiting aspects of climate at range limits, particularly at the cold end of distribution. Loehle (1998) suggested that the northern range limit of North American tree species was determined by cold tolerance. Phenotypic data supported that species from higher latitudes were usually more tolerant to the cold than those from lower latitudes (Addo-Bediako et al. 2000; Hawkins et al. 2014; Wen et al. 2018; Sunday et al. 2019). Similarly, abiotic stress appeared to be linked with the upper elevational range limit for some mountainous plant species, suggesting a predominant role of negative temperatures (Vetaas 2002; Macek et al. 2009; Körner et al. 2016). Also, the warm end of distribution may be strongly affected by climate even though the prevailing hypothesis has emphasized the importance of negative species interactions (MacArthur, 1972; Gaston 2003; Louthan et al. 2015). So far, no clear evidence exists that e.g., competition explains the southern range limit of species on a global scale. Some studies supported the hypothesis (Loehle 1998; Pither 2003), while others did not (Cahill et al. 2014). Probably because of the general dismissal of climate as a factor determining warm-end limits, few studies focused on how organisms cope with heat in the context of species distribution limits (e.g., Sunday et al. 2012; Kellermann et al. 2012;), particularly in plants (e.g., Kappen 1981; Wos and Willi 2015).

What are the sources of constraints in the evolution of the climate niche? According to simple evolutionary principles, genetic variation and selection are needed for a response to selection and adaptation (Falconer and Mackay 1996). Genetic constraints may involve low genetic variation of traits under selection. However, microevolutionary studies have shown that there is commonly ample genetic variation in single traits, and natural selection acting on populations is often strong (Mousseau & Roff 1987, Houle 1992, Kingsolver & Diamond 2011). These findings suggest generally rapid and ubiquitous adaptation through highly evolvable traits. Another type of genetic constraint is trade-offs in fitness-relevant traits, often seen as an obstacle to adaptive evolution by limiting the rate of adaptation (Futuyama and Moreno 1988; Bennett and Lenski 2007; Walker 2007). Negative genetic correlations among traits in regard to their fitness consequences appear mainly due to two non-exclusive causes. The first is that both the environment and the genetics of traits exert a limitation on trait values through differential allocation of limited amounts of resources (Bell 1984; van Noordwijk and de Jong 1986). The second cause is purely genetic; pleiotropic antagonism occurs when an allele increases the fitness via a first trait but reduces it via a second (Rose 1983). If we translate this into a thermobiology context, it is reasonable to assert that thermal extremes impose selection on some traits, resulting in a better thermal performance under one type of extreme, paid at the price of a reduction in performance in a contrasting environment or a contrasting aspect of the biology of the species. In ectothermic animals, relatively common trade-offs involve thermal resistance on the one hand, and growth, starvation resistance, longevity or reproduction on the other hand (Luckinbill 1998; Norry and Loeschcke 2002; Hoffmann et al. 2005; Stoks and De Block 2011; Casanueva et al. 2012), or cold and heat tolerance (Norry et al. 2007). Temperature can also mediate trade-offs between traits, e.g., between lifespan and reproduction (Mockett and Sohal 2006), or longevity and body size (Norry and Loeschcke 2002), or it can reverse the sign of a correlation (reviewed in Sgrò and Hoffmann 2004). In plants, trade-offs were discovered between cold tolerance and frost resistance (e.g., *Raphanus raphanistrum*; Agrawal *et al*. 2004), and between speed of development and frost tolerance (Koehler et al. 2012; Molina-Montenegro et al. 2012; Bucher et al. 2019).

While micro-evolutionary studies can shed-light on trade-offs, those involving traits related to the climate niche have not revealed any cohesive patterns (e.g., Williams et al. 2012; Kelly et al. 2013). However, in the last decades, the field of comparative phylogenetics has developed macro-evolutionary models that allow the study of adaptive evolution of more than one trait while accounting for the shared history among species (summarized in Garamszegi 2014). Based on comparative models, the phylogenetic signal of traits can be estimated and interpreted in the context of niche conservatism (Cooper et al. 2010). Furthermore, the contribution of different evolutionary processes and constraints to respond to selection can be inferred (Butler and King 2004). Three evolutionary processes are typically modelled. A first is *genetic drift*, by which inherited characters slowly change in random direction and accumulate differences over time. The process is typically modelled by Brownian motion (BM). A second process is *stabilizing selection*, a likely result of dependencies among characters under opposing selection (Wagner and Schwenk 2000). It is modelled by Ornstein-Uhlenbeck (OU1) diffusion, which constrains BM toward an optimal trait value. Recent improvements allow variation in the direction of OU diffusion across lineages, depicting the third process of *divergent selection* (OUM, Beaulieu et al. 2012). This approach has been used in evolutionary studies linking traits with the climate niche, particularly on plants, and they highlighted a link between life-form or growth strategy, and adaptation (or exposure) to a cold environment (Boucher et al. 2012; Kostikova et al. 2013; Tonnabel et al. 2018). Examples emphasize the great potential the approach has in detecting traits of adaptation to climate and revealing potential trade-offs in such adaptation or indicating general evolutionary constraints.

Here we studied trait divergence associated with the predominant elevational distribution of plant species and analysed trait data for patterns of trade-offs in a macroevolutionary context. The study of elevational gradients is promising in the context for at least two reasons. On the one hand, elevation provides a steep climatic gradient in most mountainous regions, where over a short geographic distance a reduction of the mean temperature of 0.5 K every 100 m of elevation is found rather consistently (Körner 2003). On the other hand, species often occupy narrow elevational ranges (Körner 2003), making elevational gradients unique systems for studying adaptation to thermal stress and constraints in such evolution. Our study involved 100 Brassicaceae species occurring in the central Alps of Europe, with median elevational occurrence varying from 400 to 2800 m a.s.l. Seeds of the species were raised in climate chambers under three different temperature regimes (regular frost, mild, regular heat), and over a dozen traits depicting growth, leaf morphology and coping with thermal extremes were measured. Four main hypotheses were tested. (i) Species differ in trait expression depending on their elevational distribution. (ii) Traits differ in the signature of past evolutionary processes having acted on them. (iii) Phylogenetic conservatism in traits depends on the growth (thermal) environment. And (iv) there are trade-offs among traits associated with adaptation to elevation.

## MATERIAL AND METHODS

### Plant species

One hundred taxa (i.e., species and subspecies) belonging to the Brassicaceae family and naturally occurring in the Swiss Alps (and Jura) from the colline to the alpine life zone were selected. Apart from a good representation of the elevational gradient, other criteria were level of ploidy (diploid taxa preferred) and good representation of the phylogeny (list in Supplementary material A1). In the general area, around 180 species of Brassicaceae occur, of which 28 are strictly high-elevation species. On a global scale, Brassicaceae is an angiosperm family composed of 3’700 species (including important agricultural cultivars) subdivided into three main lineages (Al-Shehbaz et al. 2006).

For this study, seeds were collected from March to September during the years 2015-2017 at two different sites for each species within Switzerland. The sites were around the most common elevation for each species, at least 50 km apart from each other and preferentially from different biogeographic regions (Jura, Plateau, northern Prealps, western and eastern Central Alps and southern Prealps). For plants with very restricted distributions, only one population was sampled, but the number of individuals was doubled. At each site, seeds were collected from 10 to 30 different mother plants over an area of usually 50 m^2^ and spaced out from each other by 5 m. For endangered species on the Red List 2002 for Switzerland (Moser et al. 2002), authorization for sampling was obtained from the respective Cantonal authority. Sampled seeds of each mother plant were stored in separate paper bags (80 g m^−2^, 60 × 90 / 12 mm, ELCO AG, Brugg, Switzerland) under cold (4 °C), dark and dry (added silica gel) conditions until sowing.

### Raising of plants under three growth treatments and trait assessment

#### Design

The experimental design involved the raising of 100 taxa, each represented by 2 populations and 3 maternal lines per population, i.e. 6 maternal lines per species. The experiment was split into 6 blocks, with a different maternal line per species in a block. Within block, plants of a maternal line were exposed to 3 temperature treatments (regular frost, mild, regular heat). The final design resulted in 1’800 individuals (100 taxa × 6 maternal lines each in a different block × 3 treatments = 1’800 individuals). Maternal lines of a population were selected randomly, and seeds of a maternal line were selected haphazardly. A first round of sowing (S1) was done without the use of giberellic acid (GA_3_), resulting in some species (20) not germinating and some heterogeneity in the timing of germination. In a second round of sowing (S2), seeds were treated with giberellic acid (GA_3_), resulting in the germination of 14 additional species (but 5 were now lacking that germinated in S1) and a more similar timing of germination.

#### Plant rearing

Seeds were germinated in climate chambers under controlled conditions, with similar procedures in S1 and in S2 (S2 described in detail below). Two seeds were placed in a 1.5 ml eppendorf tube filled with 500 μl of GA_3_ solution (500 ppm, Merck KGeA, Dornstadt, Germany), with 3 tubes per maternal line. Seeds were incubated for 1 week in dark and cold (4 °C constant in Climecabs; Kälte 3000, Landquart, Switzerland) and then sown in multipot-trays (0.06 L, 54 pots per tray with Ø 4.4 cm each, BK Qualipot; gvz-rossat.ch, Otelfingen, Switzerland). Each pot had been filled with a mixture of soil (bark compost, peat and perlite, Aussaat-und Pikiererde; Oekohum, Herrenhof, Switzerland) and sand (0-4mm) in a ratio of 2:1. Multi-pot trays were covered with a garden fleece (Windhager, Hünenberg, Switzerland) and set up in blocks within growth chambers (MobyLux GroBanks; CLF Plant Climatics, Wertingen, Germany). Growth chambers were located inside a PlantMaster (CLF Plant Climatics) with managed humidity and temperature. Trays were kept at 18 °C during daytime (8 h) and 15 °C during nighttime (16 h), at 75% relative humidity (RH), and a light intensity of 150 μmol m^−2^ s^−1^ (fluorescent white lamps and red-LED). Twice a week, blocks were moved to a different chamber, with re-randomized positioning of trays. After 3 weeks, excess seedlings were used to fill pots with no germination with the following priority: use of the same maternal line within block, or the same population, or the same species. In week 4, germinated plants were moved back to climate chambers and entire trays were subjected to one of three temperature treatments.

#### Treatment

The three temperature treatments were: “frost” (F), “mild/control” (M) and “heat” (H). Conditions of the treatments were the following: <u>frost:</u> 20 °C (daytime), then −2 °C for 1 h (−4.8 K h^−1^; nighttime) and back to 20 °C (+7.3 K h^−1^; night); <u>mild/control:</u> 20 °C constant; and <u>heat:</u> 20 °C (beginning of day), then 40 °C for 1 h (+5 K h^−1^; day), back to 20 °C (−8.3 K h^−1^; day), 20 °C (night). All treatments were conducted at cycles of 12:12 h light:dark and a light intensity of about 300 μmol m^−2^ s^−1^ (LED white lamp) and 75% RH. Plants were acclimated two days before the beginning of treatment by exposing them to milder extremes, 2 °C for the frost treatment, and 35 °C for the heat treatment. We selected extreme temperatures based on records in the field during the vegetative period (Larcher and Wagner 1976; Sutinen et al. 2001; Körner 2003), while for the mild treatment we used a common standard temperature. Trays were randomized daily within each block, while blocks where moved to different climate chambers twice a week. Plants were kept under these conditions until the 9^th^ week after sowing, when trait assessments were performed. Mean species numbers across blocks that were assessed for a particular trait within the treatments ranged from 82.1 ± 3.6 (heat) to 85.5 ± 3.5 (mild) in S2 (N = 1406 plants), and from 52.1 ± 24.0 (heat) to 74.6 ± 1.1 (mild) in S1 (N = 862 plants).

#### Traits

Two traits were assessed before treatment started: seed size (SSIZ, in mm^2^) and days to germination (TGER). Five traits depicted the trajectory of plant growth based on leaf lenght: the initial growth rate (IGR, in mm day^−1^), parameters of a 3-parameter logistic model including the maximal growth rate (MGR, scale^−1^), the mid-point until final size was reached (XMID, in days) and asymptotic size (ASYM, in mm), and finally the number of plants on day 35 of treatment (NLEA). Since smaller values of XMID meant that a plant achieved mid-size faster, values were multiplied by −1 ([−]XMID) to represent speed of growth. Five leaf functional traits were assessed: leaf area (LA, in mm^2^), specific leaf area (SLA, area over dry weight in mm^2^ mg^−1^), leaf dry matter content (LDMC, ratio of dry weight over fresh weight in mg g^−1^), leaf dissection index (LDI, no unit), and leaf thickness (LTh, in mm). Resistance of leaves to thermal extremes was assessed under −10 °C (*minus*T2) and −5 °C (*minus*T1), and +45 °C (*plus*T1) and +50 °C (*plus*T2). Resistance to T1 was tested only on non-acclimated plants (i.e., plants of the mild growth treatment), while T2 was tested on non-acclimated and acclimated plants (i.e., plants pre-exposed to frost for assessing frost resistance, and plants pre-exposed to heat for assessing heat resistance). Tolerance to repeated frost or heat during the growth phase was calculated as MGR, -XMID or ASYM under frost or heat treatment minus the respective estimate in the mild treatment, divided by the estimate in the mild treatment. We used the term frost/heat tolerance *sensu lato* (*s.l*.) to refer to tolerance and resistance together. Full details are given in Supplementary material A2. For analyses, means of replicate trait measures per plant were calculated, on which species means per treatment and sowing round and finally species means per treatment across sowing rounds were calculated.

### Statistical analysis

#### Trait expression differing with temperature treatment during growth and elevational distribution

The effect of temperature treatment, median elevation of species distribution, and their interaction on traits was tested using generalised linear mixed models based on Markov Chain Monte Carlo techniques with the ‘brm’ function of the R package {brms} (Bürkner 2017). The fixed effect of treatment was coded as a categorical variable, and contrasts were performed against the “mild” treatment or, for tolerance, against “frost”. The fixed effect of median elevation of species distribution was calculated based on reported species occurrences of a nation-wide species inventory (infoflora.ch). Median elevation was mean-centred prior to analyses. Random effects were the round of sowing (i.e., S1 and S2) and the relatedness among species. A phylogeny produced based on several dozen chloroplast genes (Patsiou et al. 2021) was pruned to species included in this study with the function ‘treedata’ of package {geiger} (Harmon et al. 2008). The final matrix was obtained with the function ‘vcv’ {ape} (Paradis and Schliep 2018) and called with the ‘cov_ranef’ argument in brm. For each model, the contribution of the phylogenetic effect was tested by comparing the model that included it as a random effect to one that did not. Model comparisons were performed using the leave-one-out cross validation (i.e., LOO), which was calculated with the ‘add_criterion’ {brms} function combined with the expected log pointwise predictive density (i.e., ELPD) with the ‘loo_compare’ {brms} function. Resistance traits were modelled by a beta distribution because of their constrained nature between 0 and 1 (i.e., 100%), -XMID and tolerances by a gaussian distribution, and the remaining traits by a log-normal distribution because values could only be positive. Sampling behaviour of MCMC was inspected visually, and number of iterations, warmup and sampling interval adapted to each model to retain an effective sampling size of 1’000. Significance was tested by probability of direction calculated with the ‘p_direction’ function in {bayestestR} (Makowski et al. 2019). All analyses and figures were done with the statistics software R v. 4.0.3 (R Core Team 2014), and calculations were performed at sciCORE (http://scicore.unibas.ch/) scientific computing center of the University of Basel.

#### Past evolutionary forces

Further analyses were performed on species trait means averaged across rounds of sowing. Phylogenetic analyses on the evolutionary processes that had shaped trait divergence among species were run separately for the three temperature treatments, and by considering variance in trait means of species of the two rounds of sowing. We tested five evolutionary models using the R package {geiger} and {mvMORPH} (Clavel et al. 2015): White noise (WN) with trait evolution independent of phylogeny, BM, BMM with different speeds of the different regimes, OU1 and OUM. For BMM and OUM, the contrasting environmental regimes were low-*vs*. high-elevation distribution of species. Assignment to one of the two classes was made using the InfoFlora (infoflora.ch) distribution information, with a threshold at 1500 m a.s.l. (splitting species of the foothills/hills from those of sub-/alpine areas). For less frequent species on Swiss territory, the assignment was verified by data on the entire Alps and neighbouring mountain massifs (based on Aeschimann et al 2004). Ancestral state reconstruction and model comparison are described in Supplementary material A2. Validation of the results was performed by simulations on synthetic data and analyses after the random removal of species (A2).

Phylogenetic half-life, i.e. the time required for a trait to evolve halfway toward its adaptive optimum, was calculated for all traits assessed in the three growth environments and in each simulation described above. Values were extracted from an OU1 model, except when elevation had a significant effect – either in mixed models or evolutionary analysis; in those cases, values were derived from an OUM process. Small values of half-life indicate fast adaptation toward the optima and a lack of phylogenetic inertia, while high values indicate that traits retain the influence of the ancestral states for a longer time. We tested for an effect of growth environment (a factor with 3 levels, with ‘mild’ as baseline) on the evolutionary lability of traits with a generalised linear mixed model with ‘brm’ (as specified above). Phylogenetic half-life was modelled assuming a lognormal distribution (only positive values), and trait was a random effect.

#### Multi-trait relationships and trade-offs

To identify putative trade-offs between pairs of traits, Pearson correlation coefficients were calculated using the ‘rcorr’ function of the package {Hmisc} (Harrell 2019). Before performing correlations, some traits were log_10_-transformed (i.e., SSIZ, MGR, NLEA, LA, LDI and RESpT2), and all traits were centred to a mean of zero and scaled to the variance. Then, highly collinear traits were removed from the dataset using the ‘vifstep’ function {usdm} with threshold of 10, which resulted in the drop of 10 traits (i.e., ASYM_Frost_, ASYM_Heat_, NLEA_Mild_, NLEA_Heat_, LA_Frost_, LA_Heat_, SLA_Frost_, LDI_Mild_, LDI_Heat_ and TOL_IGR_Frost_; correlation matrix in Supplementary material A5). To further reduce the number of traits while maintaining the most discriminating ones depending on the elevation of origin of species, discriminant analysis of principal components (DAPC) was performed with ‘dapc’ of the package {adegenet} (Jombart 2008). The optimal number of PCs to retain was selected based on stratified cross-validation with ‘xvalDapc’ function of the package {adegenet} and 10’000 simulations for each level of PC retention. Traits contributing with a loading higher than 0.024 (i.e., the third quartile of the variables contribution) were selected and used for correlation analysis.

## RESULTS

### Trait expression differing with temperature treatment during growth and with elevational distribution

Results on trait expression differing between growth treatments and species depending on their elevational distribution are summarized in Table 1, Fig. 1 and Supplementary material A3. A high fraction of traits (~70%) responded to temperature. Under regular frost compared to mild conditions, plants reached the midpoint of growth earlier (A3 Fig. 1E), but they had smaller asymptotic size (Fig. 1B) and fewer and smaller leaves (A3 Fig. 1G, H). Their leaves had less surface area per dry mass and were thicker (smaller SLA and LTh; Fig. 1C and A3 Fig. 1I, K). However, frost resistance of leaves was not significantly different after pre-exposure to frost during growth (A3 Fig. 1R). Under regular heat during growth compared to mild conditions, the maximal growth rate of plants was significantly higher (Fig. 1A), the time to maximal growth shorter (A3 Fig. 1E) and plants had smaller asymptotic size (Fig. 1B, A3 Fig. 1F) and smaller leaves (Fig. A3 Fig. 1H). Furthermore, leaves had more surface area per dry mass and less dry mass per wet weight (larger SLA, smaller LDMC; A3 Fig. 1I, J and Fig. 1C). Finally, tolerance to heat was generally higher compared to tolerance to frost for maximal growth rate and asymptotic size (Fig. 1E, G).

**Table 1.**
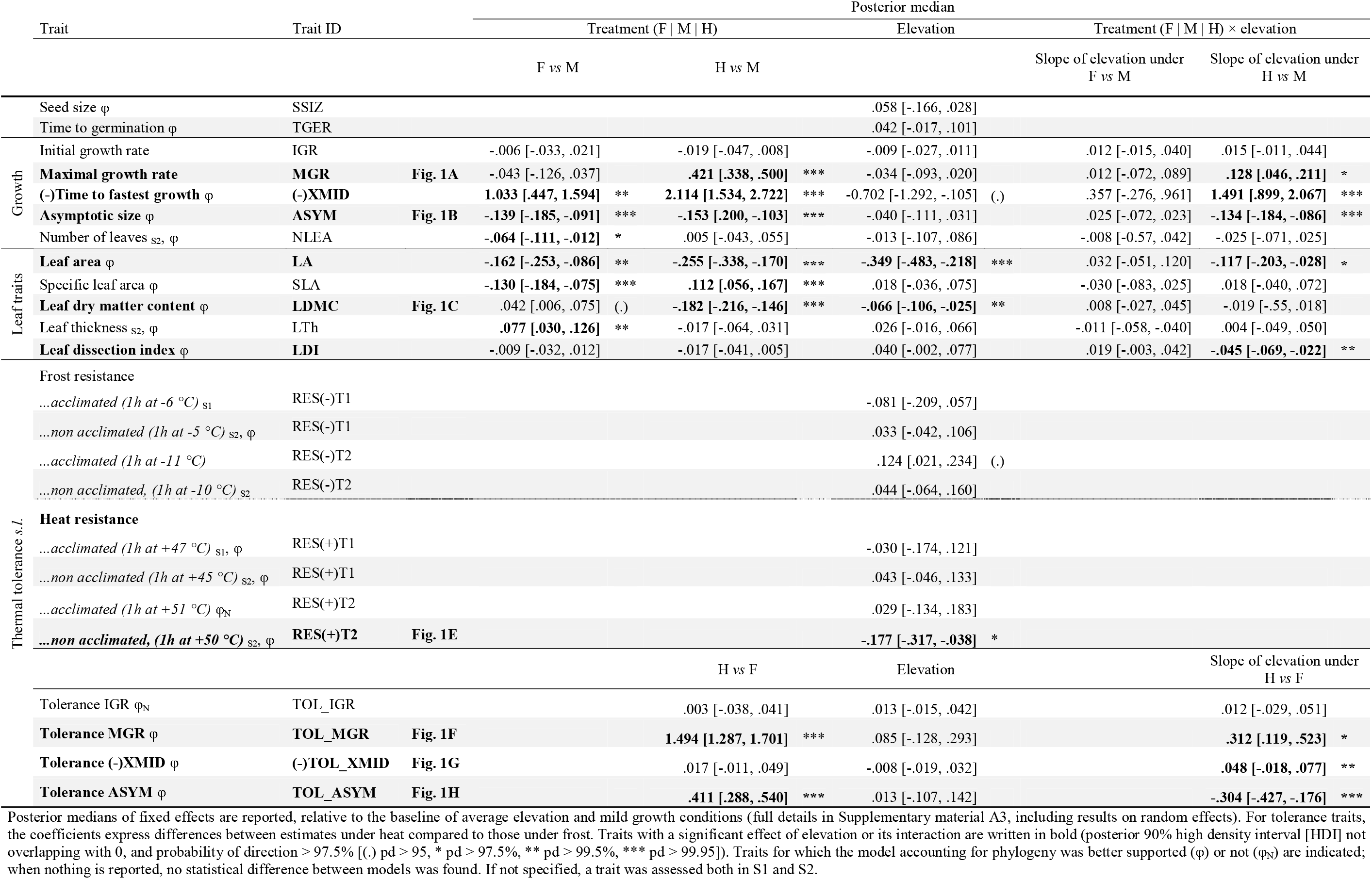
Results of mixed-effects models on the relationship between median elevation of species distribution, treatment during plant growth (regular frost [F], mild conditions [M], and regular heat [H]) and their interaction on plant traits

**Figure 1.**
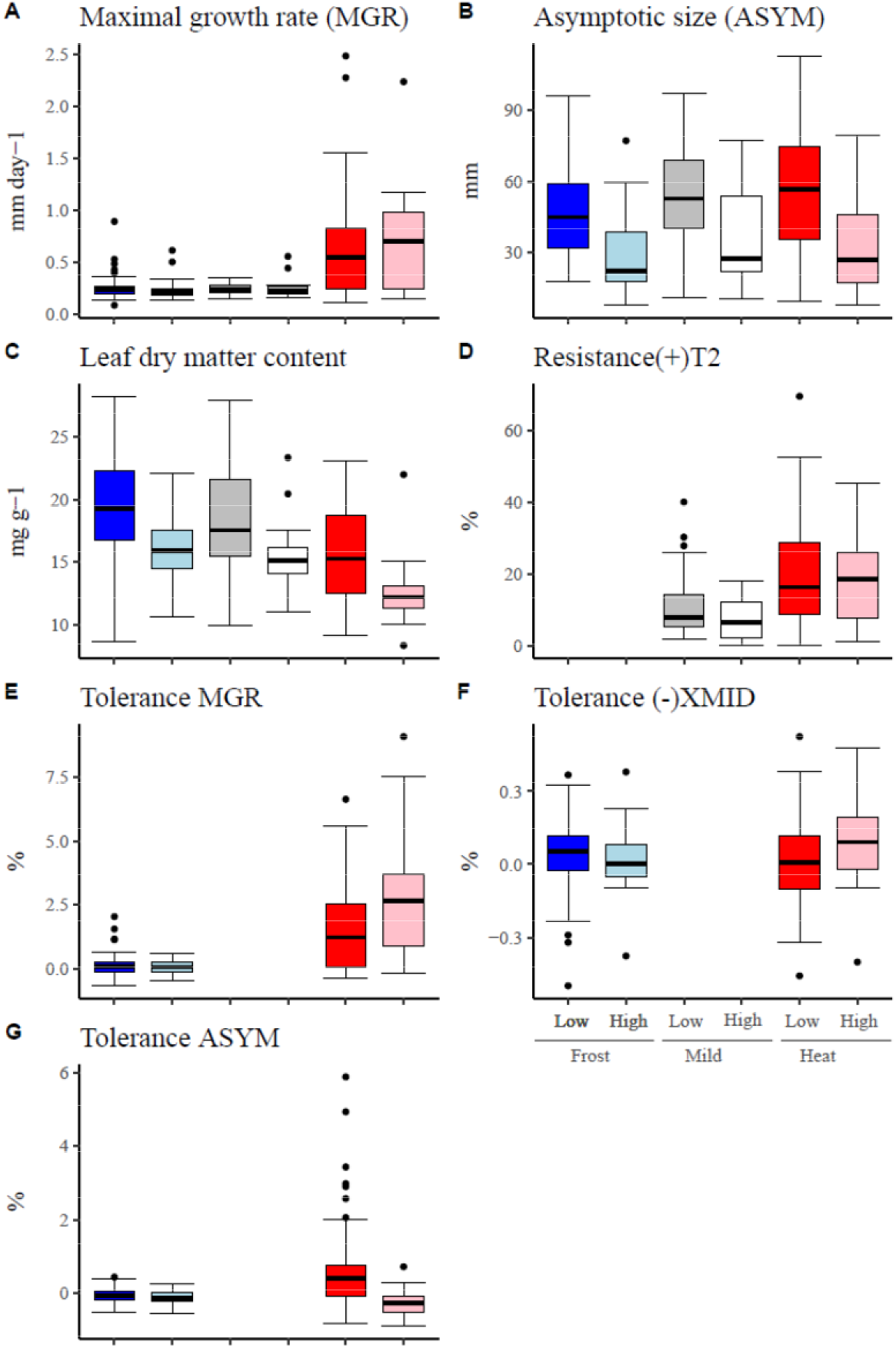
Boxplot showing the distribution of species-mean trait values for which species differed depending on their median elevation (low-*vs* high-elevation), either across growth treatments or in a particular growth treatment (frost, mild, or heat). For simplicity, only data of the second round of sowing are included and traits for which mixed-effects models and evolutionary models produced concordant results (panels for both rounds of sowing and all traits are in Supplementary material A3). Colours inside boxes represent the treatments (blue for frost, greyscale for control and red for heat), while the intensity represents median elevation of species occurrence (darker colours for low elevation and lighter colour for high elevation). The thick horizontal line is the median, the lower and upper hinges are the 25th and 75th percentiles; whiskers extends from the hinges to the smallest (largest) value at most (no further than) 1.5 * IQR of the hinges, and dots are values beyond that range.

Median elevation of species distribution alone explained only significant variation in the general expression of three traits (Tab. 1). Species occurring at higher elevation had smaller leaves (A3 Fig. 1H), lower dry-matter content (Fig. 1C) and lower heat resistance under no acclimation (RES(+)T2; Fig. 1D). A considerable fraction of traits was significantly affected by an interaction between median elevation of distribution and treatment, but only in the comparison between mild conditions and the heat treatment. The only notable exception was that higher-elevation species had increased frost resistance (after acclimation), but only for the first round of sowing (A3 Fig. 1R). When exposed to heat, higher-compared to lower-elevation species had faster growth (Fig. 1A), reached maximal growth earlier (higher -XMID, Fig. A3 Fig. 1E), but ended up being smaller (Fig. 1B), with smaller and less dissected leaves (A3 Fig. 1H, L). In line, higher-elevation species showed heightened tolerance to heat – compared to frost – by having a faster maximum growth (Fig. 1E), which was reached earlier (Fig. 1F), but they also showed lower tolerance to heat by ending up being smaller (Fig. 1G). Comparisons between models with and without considering the phylogeny revealed that its inclusion improved the model for about 70% of traits (Tab. 1, A3).

### Past evolutionary forces

Table 2 summarizes results on analyses of evolutionary processes having acted on traits, for each growth environment (for a full account see Supplementary material A4). The comparison between the two evolutionary switch models (i.e., ‘ER’, under which low and high elevation are predicted to change at equal rate; or ‘ARD’, under which for- and backward rates between states can take different values) indicated a slightly better performance of the more parameterised model (AIC_ER_ 106.898; AIC_ARD_ 103.111), with a fitted value of Q from low → high of .030 and low ← high of.910.

**Table 2.**
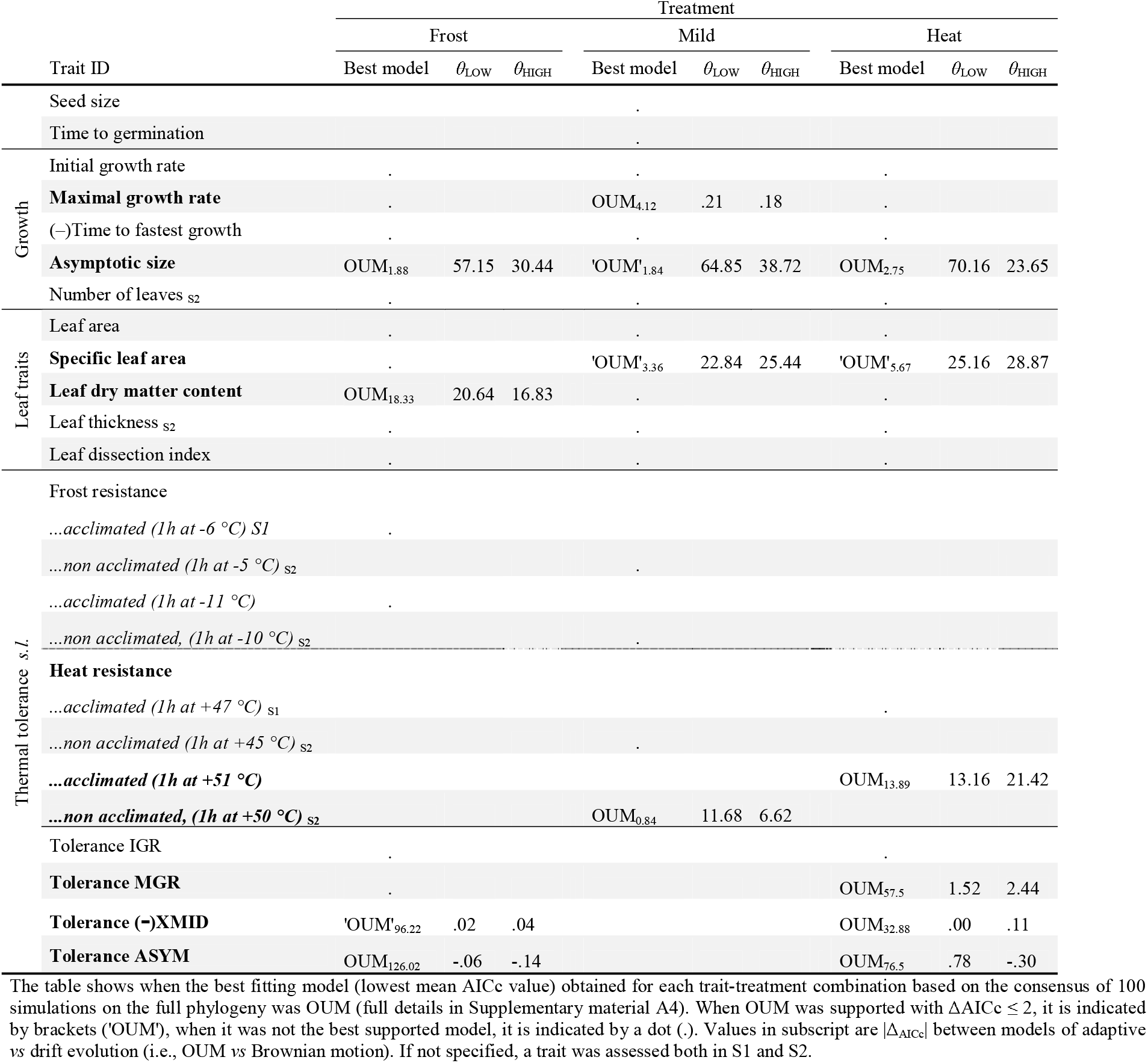
List of traits measured in the three growth environments (regular frost, mild and regular heat) for which the best supported evolutionary model was Ornstein-Uhlenbeck with two optima (OUM), and the suggested trait optima (θ) for low- and high-elevation species

Several of the traits found to differ between low- and high-elevation species in mixed-model analyses were confirmed to support a scenario of adaptive evolution with two optima. These traits included: maximal growth rate, asymptotic size, leaf dry matter content, heat resistance and tolerances in growth parameters (Tab. 2, A4). The optimum for high-elevation species was at a lower MGR under control conditions, at a smaller asymptotic size under all growth conditions and at a lower LDMC under regular frost. Furthermore, high-elevation species had an optimum at lower heat resistance when raised under mild conditions, but at higher heat resistance when raised under regular heat. Finally, high-elevation species had optima at higher tolerance values to heat based on MGR and -XMID; they had been selected to accelerate the speed of growth more under heat stress. But they had optima for tolerance to frost and heat based on asymptotic size that were lower. These results appear to be robust, as they did not deviate significantly from the results obtained from bootstrap simulations (A4).

Simulations performed on the phylogeny but with synthetic data (A4 Fig. 2) revealed that adaptive divergence between low- and high-elevation species was identified correctly when trait variance was low (<100) and the difference between optima (thetas) large. False positives for the adaptive model were rare, while false negatives in favour of OU or WN were frequent. Simulations that randomly removed a third of the species generally resulted in increased support for OUM (A4). Specifically, to the traits already mentioned above, high-compared to low-elevation species also differed in having optima at slower initial growth under control conditions, but at higher maximum growth rate under stress (in frost and heat treatments). Furthermore, optima differed for leaf size under frost (i.e., at smaller leaf size for high-elevation species) and leaf dry matter content under heat (i.e., at lower LDMC for high-elevation species). No further differences were found for traits related to resistance, while tolerance to regular frost based on maximal growth rate had separate optima, with the one of high-elevation species being at lower tolerance to frost.

**Figure 2.**
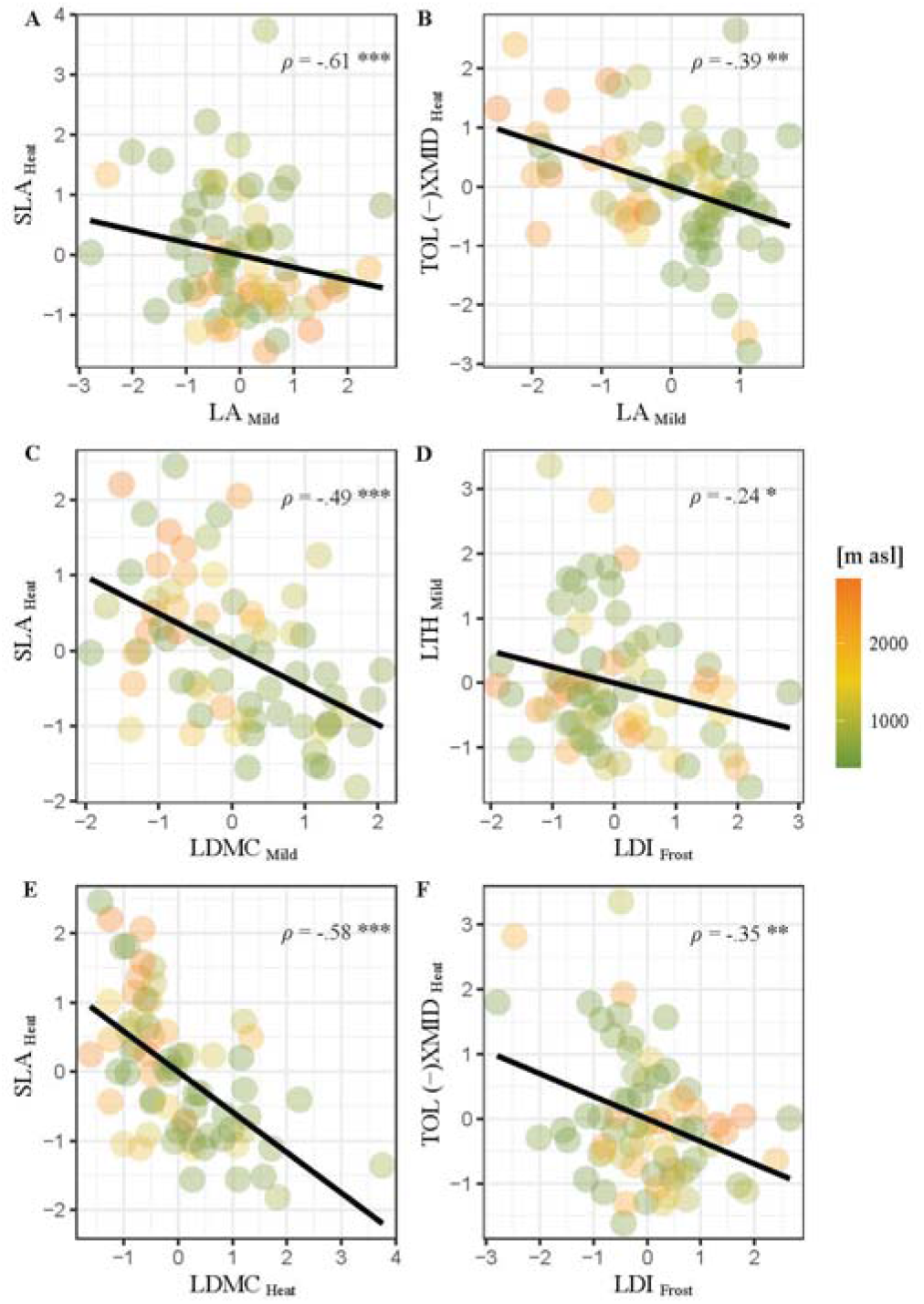
Trait differentiation between low- and high-elevation species, as revealed by discriminant analyses and multi-trait correlations. Each point represents a species. The median elevation of origin is represented by a colour scale ranging from green (low elevation) to brown (high elevation). The black line reflects the relationship between pairs of traits and the associated correlation coefficient is reported (full details in Supplementary material A5). Traits values are centred and scaled to unit variance.

Measures of phylogenetic half-life (i.e., ln(2) *alpha*^−1^; Tab. 3, A4) were rarely significantly larger than 0 (25-38% depending on treatment, Tab. 3), The most constrained traits were associated with size and morphology, e.g., ASYM with a half-life of 10-16 Mya, LA_.Heat_ with 25 Mya, LTh with 7-10 Mya and LDI_.Frost_ with 15 Mya. Mixed-effects analysis with bootstrap simulations revealed that the evolution of trait values under regular frost was less constrained compared to mild conditions or regular heat (i.e., Frost *vs* Mild: −.605 [−.616, −.593]; Heat *vs* Mild: .192 [.180, .204]), resulting in a reduction of average half-life of about 50% (A4 Fig. 3A, B).

**Table 3.**
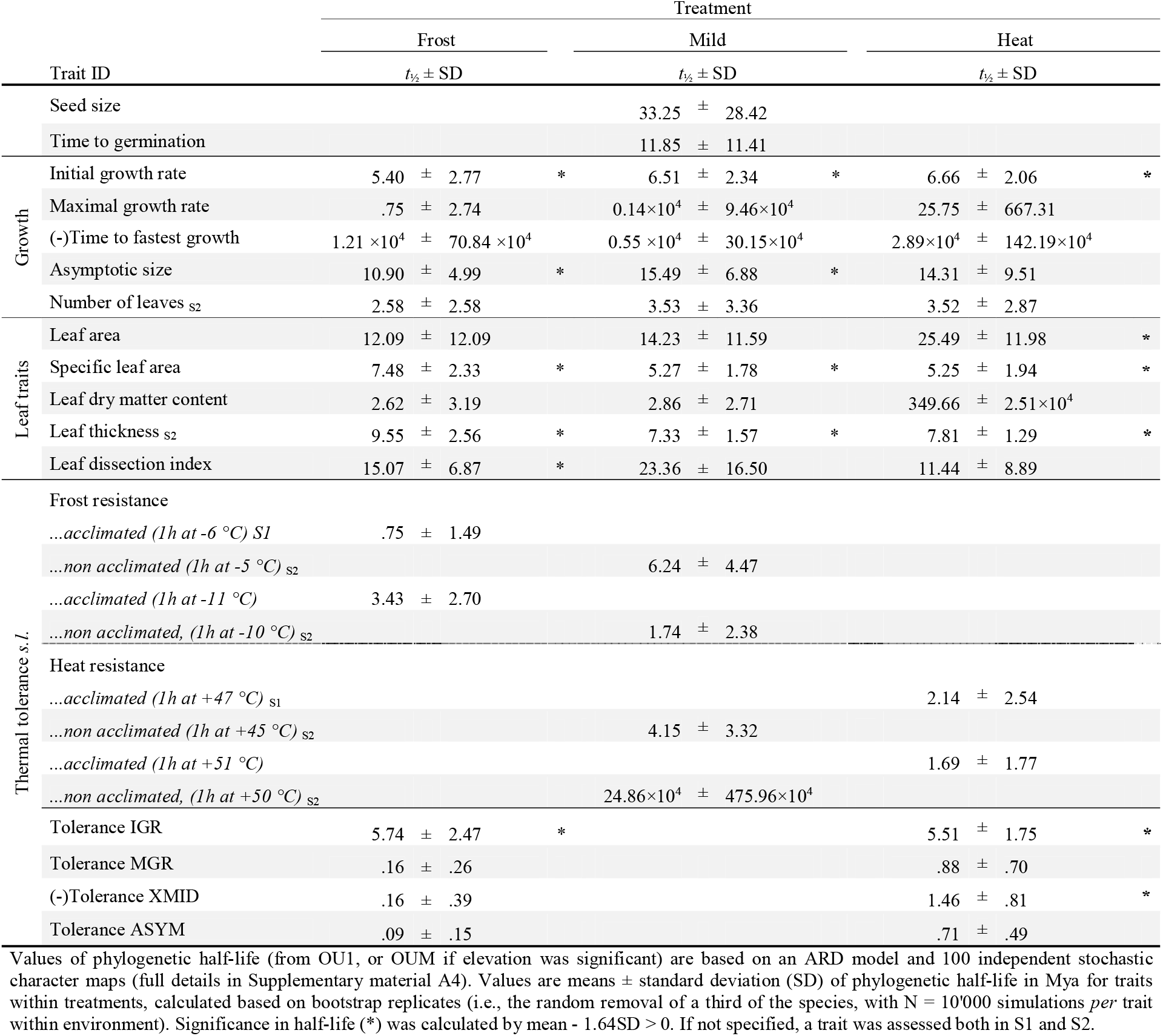
Half-life of trait evolution toward the optimum in Mya

### Multi-trait relationships and trade-offs

A principal component analysis on trait values of all trait-growth treatment combinations revealed their correlation structure. The first PC axis explained 15.7% of the total variance and depicted the relationship between timing of plant growth, especially in the heat treatment, and plant size under mild conditions. The second PC (10.5%) was primarily influenced by LTh, and to a lesser extent by basal resistance and tolerance components, depicting a distinction between these two strategies (Supplementary material A5). The optimal number of principal components to retain (i.e., lowest MSE and highest mean success) based on cross-validation was 35 (accounting for 99% of trait variation, A5).

With these PCs, taxa could be assigned to their elevation of origin, either low or high, with an accuracy of 98% and 94.4% respectively (A5). In multivariate space, the trait with the greatest weight was leaf area under mild conditions, while the other traits that contributed most to differentiating low- and high-elevation species were associated to leaf morphology under mild and warm conditions (i.e., LTh, LDMC, SLA), speed of growth under heat (i.e., TOL_IGR, TOL_-XMID) and tolerance under frost (i.e., LDI, TOL_ASYM, A5). Pearson correlations were significantly negative between specific leaf area under heat and leaf area (LA_Mild_, Fig 2A) or leaf dry matter content (LDMC_Mild_, Fig. 2C; LDMC_Heat_, Fig. 2E), with the latter correlation being likely driven by non-independence of calculating estimates. Furthermore, leaf area under mild conditions was negatively correlated with heat tolerance based on the mid-point of growth (TOL_-XMID, Fig. 2B), suggesting a trade-off between maintaining large size and speeding up growth under heat. Tolerance under warm based on midpoint of growth was also negatively associated with leaf dissection index under frost (Fig. 2F), which in turn was negatively associated with leaf thickness under mild conditions (Fig. 2D). However, these two correlations did not involve traits linked to elevational distribution.

## DISCUSSION

Past studies in ecology and biogeography have indicated that temperature is a limiting factor of species distribution, suggesting that there are ubiquitous constraints to the evolution of the climate niche. To improve our understanding of such constraints, we studied approximately 100 species differing in elevational distribution and presumably with different climate niches. More specifically, we investigated which traits differed with elevational distribution, whether those traits had been under divergent selection over the elevational gradient, and potential sources of constraints in their adaptive divergence. The species were found to systematically differ in few traits. Most importantly, higher-elevation plants were found to have smaller and less robust leaves. Further differences emerged when growing conditions included regular heat bouts. Then higher-elevation species accelerated growth more, at the cost of a considerable reduction in size. The same or similar traits were found to be under divergent selection over the elevational gradient, and some were involved in moderate trade-offs, notably the ability to speed up growth under heat and plant size. The discussion focuses on traits under divergent selection, evidence for evolutionary constraints, and hypotheses on the selection environment and adaptive strategies.

### Trait differences between low- and high-elevation species

Generalized linear models and evolutionary models mainly overlapped in pointing to differences in traits depending on whether species had low- or high elevation distributions (Table 4). The traits that were consistently different between low- and high-elevation species in the two types of models depicted plant size (i.e., ASYM, LA), leaf morphology (i.e., LDMC, SLA), the response of speed of growth to stress, and thermal resistance.

**Table 4.**
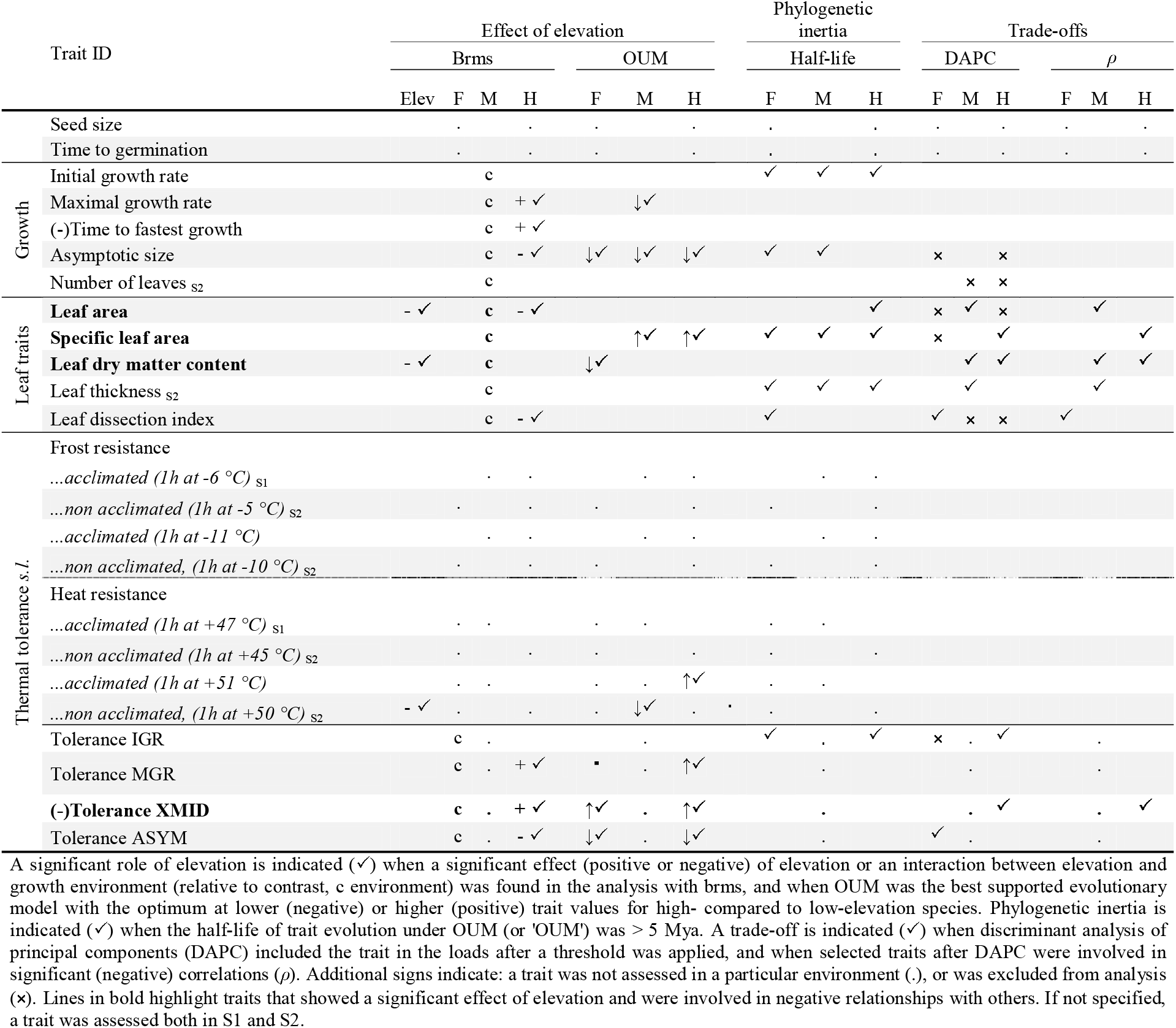
Summary of results on trait differences between low- and high-elevation species in the three growth treatments (regular frost [F], mild conditions [M], and regular heat [H]) across types of analyses (mixed models [brms], testing for two evolutionary optima [OUM], half-life of trait evolution, discriminant analysis of principal components [DAPC], and (negative) correlations [*ρ*])

Across growth environments, alpine species had smaller leaves and less dry matter content in leaves (Tab. 4, Fig. 1B, C and A3 Fig. 1H, J). Evolutionary models supported that optima for plant size were at smaller values for high-compared to low-elevation species under all growth conditions. Furthermore, they supported an optimum at lower LDMC under growth conditions with regular frost, and as a trend an optimum at higher SLA, which is typically inversely related to LDMC, under mild conditions or conditions with regular heat. Results for size are in line with previous studies on multi-species comparisons, which reported a reduction in leaf size with increasing elevational distribution (Qi et al. 2014; Zhong et al. 2014). In contrast, previous studies reported either higher LDMC and smaller SLA (Körner et al. 1986; Qi et al. 2014; Rosbakh et al. 2014; Midolo et al. 2019), or the contrary (Zhong et al. 2014) as found for the Brassicaceae. Lower LDMC and higher SLA are typically associated with a strategy of fast assimilation and growth but weak hardiness and short leaf life-span (Pérez-Harguindeguy et al. 2013).

The other type of trait that generally differed between low- and high-elevation species was the response to heat during the growth phase. Both heat and frost caused plants and their leaves to be smaller, indicating that conditions were generally stressful. Furthermore, plants speeded up growth under these conditions; the time to reach the midpoint of asymptotic size was shorter (-XMID), and under the regular occurrence of heat bouts, also the maximum growth rate was higher (MGR). An important finding of this study is that higher-compared to lower-elevation species could accelerate growth under conditions with regular heat bouts even more (MGR, -XMID; Fig. 1A, E, F), at the cost that their leaves were more reduced (Fig. 1B, G). Evolutionary models too provided evidence that tolerance for speeding up growth (TOL_MGR, TOL_-XMID) under heat had an optimum at higher values in high-elevation species. Evolutionary models pointed also to an optimum at higher values for tolerance of speeding up growth under frost (as a trend). Results suggest general selection for escape strategies under stress, and that high-elevation species seem to have adapted to exploit heat phases better by growing faster when they occur. The finding is novel and needs verification in more plant families.

Interestingly, low- and high-elevation species also differed in thermal resistance, though not in the direction that was previously advocated. Our mixed-effects analysis supported that heat resistance decreased with median elevational of species distribution. Evolutionary models supported a lower optimum for basal heat resistance in high-elevation species, but a higher optimum of acclimation-based heat resistance. However, increased frost resistance (after acclimation) in high-elevation species was only reported for the first round of sowing but not the second (−12 °C S1 *vs* - 10 °C S2, A3 Fig. 1R), and the result was significant only when phylogeny was not considered (A3, model 15). In contrast, a number of earlier studies documented rather consistently that high-elevation tree species were more frost resistant (Körner 2003; Taschler and Neuner 2004; Neuner 2014; Neuner et al. 2020; Schrieber et al. 2020). The discrepancy may have two potential reasons. First, the latter studies did not account for phylogeny in their analysis, which could have produced increased type I error (Li and Ives 2017). Second, there may be fundamental differences between trees and herbaceous plants in the role of frost resistance on distribution limits because of differences in the life history or the plant architecture and functioning.

In summary, the picture that emerges is that high-compared to low-elevation species are fast growers when it is warm, but have reduced size, have less hardy leaves and are neither particularly heat-nor frost-resistant.

### Trade-offs and evolutionary inertia

We detected trade-offs among traits that contributed most to the differentiation between low- and high-elevation species (Tab. 4, A5). Specific leaf area under heat was negatively related with leaf area (LA_Mild_, Fig 2A). In turn, leaf area under mild conditions was negatively correlated with heat tolerance based on the mid-point of growth (TOL_-XMID, Fig. 2B). Before discussing the two results in a more general context, it is to note that when analyses were done, a fraction of traits, which were actually calculated on the basis of trait-growth environment combinations, had already been excluded because of redundancy in information. The phenotypic aspect that the remaining traits represented was therefore probably larger. Based on this reasoning, we can say that an important trade-off was between assimilation efficiency combined with less leaf hardiness (high SLA) under heat and (plant) size. Another was between size and the capacity to speed up growth under heat. In other words, there is good macroevolutionary evidence that fast growth under heat, small size and assimilation potent leaves with less dry mass come as a syndrome of high-elevation species, shaped by trade-offs. Whether these trade-offs occur on a within-species level and may constrain adaptive evolution and niche expansion at range edges remains to be tested.

Trade-offs involving thermal resistance or tolerance were also found, but may have little impact on species distribution. Weak to moderate negative relationships were detected between non-acclimated resistances (to cold or heat) and assimilatory capacity (SLA, number of leaves; A5). But, resistance did not figure among the nine most relevant traits in differentiating low- and high-elevation species in a multivariate space (Tab. 4; Fig. 2; A5).

Considerable evolutionary half-lives of traits important in driving elevational distribution were found. The highest value of phylogenetic inertia was found for leaf area, one of the two most discriminating traits between low- and high-elevation species (Fig. 2, A5). The half-life was estimated to be ~26Mya when leaf area was expressed under the regular occurrence of heat (Tab. 3, A4). Also asymptotic size and leaf dissection index (under regular occurrence of frost) had considerable half-lives, between 11 and 15 Mya. The remaining traits (i.e., IGR, SLA, LTH, TOL_IGR and -TOL_XMID) had lower, but still considerable values ranging from ~1.5 Mya for heat tolerance based on the time until fastest growth, to 9.5 Mya for leaf thickness under cold conditions. This considerable half-lives generally indicate constraints to adaptive evolution.

### Selection environment, adaptive strategies and evolutionary constraints

Insights discussed above and further ones gained from analyses evoke novel hypotheses on the causes of limits to niche evolution and disparate elevational or climatic distribution.

Evidence for divergent adaptation between low- and high-elevation species was more common for traits recorded under mild and heat conditions compared to the regular occurrence of frost (Tab. 3, Tab. 4). This strongly supports that high-elevation species have adaptively diverged on exploiting warm conditions and not (so much) to resist the cold. This is a very important insight, and also – as a side note – warrants attention that the detection of traits under selection is environment-dependent. Comparative studies typically rely on measurements taken in the field or on collection material (e.g., Luxbacher and Knouft 2009; Edwards and Smith 2010), or after raising organisms under standard conditions (e.g., Kellermann et al. 2012; Mason and Donovan 2015). While the former brings the problem of the inability of separating the effects of genetics and the environment on trait differences, the latter has the flaw that the adaptive potential of a trait may not be detected as the environment is not the one in which divergence is expressed. Again, for Brassicaceae along the elevational gradient, it is mild and heat conditions that are likely to have played more of a role in adaptive divergence.

Several insights speak in favour that high-elevation conditions select for faster growth at the cost of small size and possibly a shorter life, with the environmental driver being the short growing season. On the one hand, our study showed that plants of high elevations were not better at coping with cold, but they had evolved to better exploit warm conditions for fast growth. In line, previous eco-physiological studies reported higher photosynthetic rate in alpine herbaceous species cultivated at warmer temperature (Mächler and Nösberger 1977) or during daily warm spells in the wild (Körner and Diemer 1987), pointing to faster resource acquisition under warm conditions. Furthermore, niche-modelling suggested that upper ranges were constrained not primarily by the direct effect of cool temperatures but the brevity of the growing season (Morin et al. 2007; Patsiou et al. 2021). These studies too pointed to speed of growth or development being under selection under higher-elevation conditions. Based on the two sets of insights, we propose that whether a species (of Brassicaceae) can live at high elevation depends on the ability to cope with the short growing season, which is achieved by maximising growth during short thermal windows when the temperature is relatively high. Superficially, the geographic pattern may resemble counter-gradient variation (Conover and Schultz 1995), where high-elevation genotypes grow faster while their environment may generally cause growth to be slow. One distinction is that the heightened acceleration of growth is expressed only under warmer conditions, and a second is that the relevant environmental difference seems to be the shorter growing season.

Analyses on trade-offs pointed to leaf morphology being coupled with fast growth under heat and reduced plant size. Speeding up growth under heat was negatively correlated with leaf size, while small life size implied higher SLA (related with lower LDMC) – generally thinner leaves with higher assimilation capacity (Fig. 2A, B). The importance of the leaf morphology in this context adds a further notion of a constraint of fast growth. According to the world-wide leaf economics spectrum (Wright et al. 2004), species either follow a strategy of quick return on investment, with nutrient-rich leaves, high photosynthetic rates, and short life-spans *versus* a strategy of slow return, with expensive but long-lived leaves. In a broader context, the continuum of fast production versus slowness is also reflected in the concept of r/K selection (Pianka 1970), where r-selected species grow more rapidly, but to a smaller size and they reproduce earlier, while K-selected species grow more slowly, but to larger size and they reproduce later. For plants, the concept was expanded, with now three strategies – stress-tolerant (S), competitive (C), ruderal (R) – being positioned along three axes of environmental gradients, of abiotic stress, competition and disturbance (Grime 1977). Pierce et al. (2013) showed how these strategies can be correctly attributed with the use of the same leaf-traits that show the main trade-offs in our work, i.e., leaf area, leaf dry matter content and specific leaf area. However, and in contrary to their reports, LDMC and LA did not form separate axes in our study. Nonetheless, following their sorting suggests that alpine (Brassicaceae) species primarily follow an r-strategy, whereas lowland species follow a C/S (or K) strategy, at least in relation to temperature responses.

Finally, also phylogenetic inertia of traits was found to depend on the environment in which they were expressed (Tab. 3, A4 Fig. 1A, B). In our study, the mild and heat treatments were not only the more discriminating among low- and high-elevation species, they were also those in which traits had on average higher phylogenetic inertia. The phylogenetic half-life of traits expressed under mild and heat was 50% higher compared to trait expression under the regular occurrence of frost. Results therefore suggest that adaptation to exploit or live under generally warmer conditions is more constrained. The result is in line with a recent large-scale phylogenetic analysis, showing that across plants and animals, the rate of adaptation to warm was much slower than to cold, both in endotherms and ectotherms (Bennet et al. 2021).

## CONCLUSION

Our study highlights that the most discriminating traits separating high-from low-elevation Brassicaceae species are their ability to speed up growth under conditions with heat bouts, at the cost of much reduced leaf and plant size, and possibly a more ephemeral lifestyle with less investment into leaves. Results suggest a general trade-off between exploiting the short vegetation period at high elevation and being less enduring in general or under certain thermal extremes or under competition. The trade-off could be a result of multivariate selection differing among low- and high-elevation sites and/or negative genetic correlations. In parallel, we found that thermal resistance did not play a strong role in differentiating species along the elevational gradient. Finally, we found evidence that divergent adaptation under conditions with regular heat was more pronounced compared to conditions with regular frost, and that adaptation to heat was more constrained.

## Declaration of authorship

AM and YW conceived the study and conducted the field work. AM executed the experimental work, run the statistical analyses and wrote the first draft of the manuscript. YW contributed to writing. All authors gave final approval for publication.

## Acknowledgements

We thank Lea Bona, Camilla Jenny, Adrian Möhl, Ramon Müller, Jens Paulsen, Florian Schreier, and Daniel Slodowicz,for collecting seeds in the field. Georg Ambruster, Olivier Bachmann, Markus Funk, Kay Lucek, Michela Meier, Jens Paulsen, Theofania-Sotiria Patsiou, Antoine Perrier, Susanna Riedl, Dario Sanchez-Castro, Martinez Sylvia and Nora Walden, helped with sowing seeds, taking plant measures and fruitful discussions. Finally, we would like to thank all the field botanists that contributed to the collection of occurrence data for the InfoFlora database.

## SUPPLEMENTARY MATERIAL

A1 – List of species and populations used in this study

A2 – Additional information on traits measurement and methods for validating evolutionary models

A3 – Mixed-effects model on the relationship between traits, treatment and elevational distribution and data distribution.

A4 – Model of trait evolution (WN, BM, BMM, OU, OUM), optimal trait values, AICc distribution, phylogenetic half-life and simulations

A5 – Multivariate analysis, and Pearson’s correlation

## REFERENCES

Addo-Bediako, A., S. L. Chown, and K. J. Gaston. 2000. Thermal tolerance, climatic variability and latitude. Proc. R. Soc. B. 267:739–745.

Aeschimann, D., K. Lauber, D. M. Moser, and J.-P. Theurillat. 2004. Flora alpina. Éditions Belin.

Agrawal, A. A., J. K. Conner, and J. R. Stinchcombe. 2004. Evolution of plant resistance and tolerance to frost damage. Ecol. Lett. 7:1199–1208.

Al-Shehbaz, I. A., M. A. Beilstein, and E. A. Kellogg. 2006. Systematics and phylogeny of the Brassicaceae (Cruciferae): an overview. Plant Syst. Evol. 259:89–120.

Angilletta, M. J. 2009. Thermal adaptation: a theoretical and empirical synthesis. Oxford Univ. Press, Oxford, U.K.

Beaulieu, J. M., D. C. Hwang C. Boettiger, and B. C. O’Meara. 20152. Modelling stabilizing selection: expanding the Ornstein–Uhlenbeck model of adaptive evolution. Evolution 66:2369–2383.

Bell, G. 1984. Measuring the cost of reproduction. I. The correlation structure of the life table of a plank rotifer. Evolution 38:300–313.

Bennett, A. F., and R. E. Lenski. 2007. An experimental test of evolutionary trade-offs during temperature adaptation. Proc. Natl. Acad. Sci. 104:8649–8654.

Bennett, J. M., Sunday, J., Calosi, P., Villalobos, F., Martínez, B., Molina-Venegas, R., … & Olalla-Tárraga, M. Á. (2021). The evolution of critical thermal limits of life on Earth. Nat. Commun., 12, 1–9.

Blom, M. P. K., P. Horner, and C. Moritz. 2016. Convergence across a continent: adaptive diversification in a recent radiation of Australian lizards. Proc. R. Soc. B. 283:20160181.

Boucher, F. C., W. Thuiller, C. Roquet, R. Douzet, S. Aubert, N. Alvarez, and S. Lavergne. 2012. Reconstructing the origins of high-alpine niches and cushion life form in the genus *Androsace s.l*. (Primulaceae). Evolution 66:1255–1268.

Bozinovic, F., M. J. M. Orellana, S. I. Martel, and J. M. Bogdanovich. 2014. Testing the heat-invariant and cold-variability tolerance hypotheses across geographic gradients. Comp. Biochem. Physiol. A. Mol. Integr. Physiol. 178:46–50.

Briceño, V. F., D. Harris-Pascal, A. B. Nicotra, E. Williams, and M. C. Ball. 2014. Variation in snow cover drives differences in frost resistance in seedlings of the alpine herb *Aciphylla glacialis*. Environ. Exp. Bot. 106:174–181.

Bucher, S. F., R. Feiler, O. Buchner, G. Neuner, S. Rosbakh, M. Leiterer, and C. Römermann. 2019. Temporal and spatial trade-offs between resistance and performance traits in herbaceous plant species. Environ. Exp. Bot. 157:187–196.

Bürkner, P.C. 2017. Advanced Bayesian multilevel modeling with the R package brms. ArXiv170511123 Stat.

Butler, M. A., and A. A. King. 2004. Phylogenetic comparative analysis: a modeling approach for adaptive evolution. Am. Nat. 164:683–695.

Cahill, A. E., M. E. Aiello-Lammens, M. C. Fisher-Reid, X. Hua, C. J. Karanewsky, H. Y. Ryu, G. C. Sbeglia, F. Spagnolo, J. B. Waldron, and J. J. Wiens. 2014. Causes of warm-edge range limits: systematic review, proximate factors and implications for climate change. J. Biogeogr. 41:429–442.

Casanueva, M. O., A. Burga, and B. Lehner. 2012. Fitness trade-offs and environmentally induced mutation buffering in isogenic *C. elegans*. Science 335:82–85.

Clavel J, Escarguel G, and Merceron G. 2015. mvMORPH: an R package for fitting multivariate evolutionary models to morphometric data. Methods Ecol. Evol. 6:1311–1319.

Connallon, T., and C. M. Sgrò. 2018. In search of a general theory of species’ range evolution. PLoS Biol. 16:e2006735.

Conover, D. O., and E. T. Schultz. 1995. Phenotypic similarity and the evolutionary significance of countergradient variation. Trends Ecol. Evol. 10:248–252.

Cooper, N., W. Jetz, and R. P. Freckleton. 2010. Phylogenetic comparative approaches for studying niche conservatism. J. Evol. Biol. 23:2529–2539.

Dahl, E. 1951. On the Relation between summer temperature and the distribution of alpine vascular plants in the lowlands of Fennoscandia. Oikos 3:22–52.

Donoghue, M. J. 2008. A phylogenetic perspective on the distribution of plant diversity. Proc. Natl. Acad. Sci. 105:11549–11555.

Edwards, E. J., and S. A. Smith. 2010. Phylogenetic analyses reveal the shady history of C4 grasses. Proc. Natl. Acad. Sci. 107:2532–2537.

Falconer, D. S., and T. F. C. Mackay. 1996. Introduction to quantitative genetics. Longmans Green, Harlow, Essex, U.K.

Futuyama, D. J., and G. Moreno. 1988. The evolution of ecological specialization. Annu. Rev. Ecol. Syst. 19:207–233.

Garamszegi, L. Z. 2014. Modern phylogenetic comparative methods and their application in evolutionary biology: concepts and practice. Springer, Berlin Heidelberg, Germany.

Gaston, K. J. 2003. The structure and dynamics of geographic ranges. Oxford Univ. Press, Oxford, U.K.

Gómez, C., E. A. Tenorio, P. Montoya, and C. D. Cadena. 2016. Niche-tracking migrants and niche-switching residents: evolution of climatic niches in New World warblers (Parulidae). Proc. R. Soc. B. 283:20152458.

Gratani, L. 2014. Plant phenotypic plasticity in response to environmental factors. Adv. Bot. 2014:208747.

Grime, J. P. 1977. Evidence for the existence of three primary strategies in plants and its relevance to ecological and evolutionary theory. Am. Nat. 111:1169–1194.

Harmon L. J, J. T Weir, C. D Brock, R. E Glor, and W. Challenger. 2008. GEIGER: investigating evolutionary radiations. Bioinformatics 24:129–131.

Harrell, J., Frank E., and with contribution from Charles Dupont and many others. 2019. Hmisc: Harrell Miscellaneous. R package version 4.2-0.

Hawkins, B. A., M. Rueda, T. F. Rangel, R. Field, and J. A. F. Diniz-Filho. 2014. Community phylogenetics at the biogeographical scale: cold tolerance, niche conservatism and the structure of North American forests. J. Biogeogr. 41:23–38.

Herrando-Pérez, S. 2013. Climate change heats matrix population models. J. Anim. Ecol. 82:1117–1119.

Hoffmann, A. A., R. Hallas, A. R. Anderson, and M. Telonis-Scott. 2005. Evidence for a robust sex-specific trade-off between cold resistance and starvation resistance in *Drosophila melanogaster*. J. Evol. Biol. 18:804–810.

Houle, D. 1992. Comparing evolvability and variability of quantitative traits. Genetics 130:195–204.

Hutchinson, G. E. 1957. Concluding Remarks. Cold Spring Harb. Symp. Quant. Biol. 22:415–427.

Iversen, J. 1944. *Viscum*, *Hedera* and *Ilex* as climate indicators. Geol. Fören. Stockh. Förh. 66:463–483.

Jombart, T. 2008. adegenet: a R package for the multivariate analysis of genetic markers. Bioinformatics 24:1403–1405.

Kappen, L. 1981. Ecological significance of resistance to high temperature. Pp. 439–474 *in* O. L. Lange, P. S. Nobel, C. B. Osmond, and H. Ziegler, eds. Physiological plant ecology I: responses to the physical environment. Springer, Berlin Heidelberg, Germany.

Kellermann, V., J. Overgaard, A. A. Hoffmann, C. Fløjgaard, J.-C. Svenning, and V. Loeschcke. 2012. Upper thermal limits of *Drosophila* are linked to species distributions and strongly constrained phylogenetically. Proc. Natl. Acad. Sci. 109:16228–16233.

Kelly, M. W., R. K. Grosberg, and E. Sanford. 2013. Trade-offs, geography, and limits to thermal adaptation in a Tide Pool Copepod. Am. Nat. 181:846–854.

Kingsolver, J. G. 2009. The well-temperatured biologist. Am. Nat. 174:755–768.

Kingsolver, J. G., and S. E. Diamond. 2011. Phenotypic selection in natural populations: what limits directional selection? Am. Nat. 177:346–357.

Koehler, K., A. Center, and J. Cavender-Bares. 2012. Evidence for a freezing tolerance–growth rate trade-off in the live oaks (*Quercus series* Virentes) across the tropical–temperate divide. New Phytol. 193:730–744.

Körner, C. 2003. Alpine plant life: functional plant ecology of high mountain ecosystems. 2nd ed. Springer, Berlin Heidelberg, Germany.

Körner, C., D. Basler, G. Hoch, C. Kollas, A. Lenz, C. F. Randin, Y. Vitasse, and N. E. Zimmermann. 2016. Where, why and how? Explaining the low-temperature range limits of temperate tree species. J. Ecol. 104:1076–1088.

Körner, C., P. Bannister, and A. F. Mark. 1986. Altitudinal variation in stomatal conductance, nitrogen content and leaf anatomy in different plant life forms in New Zealand. Oecologia 69:577–588.

Körner, C., and M. Diemer. 1987. *In situ* photosynthetic responses to light, temperature and carbon dioxide in herbaceous plants from low and high altitude. Funct. Ecol. 1:179–194.

Kostikova, A., G. Litsios, N. Salamin, and P. B. Pearman. 2013. Linking life-history traits, ecology, and niche breadth evolution in North American Eriogonoids (Polygonaceae). Am. Nat. 182:760–774.

Larcher, W. and J. Wagner. 1976. Temperaturgrenzen der CO2-Aufnahme und Temperaturresistenz der Blätter von Gebirgspflanzen im vegetationsaktiven Zustand. Oecol. Plant. 11:361–374.

Lee-Yaw, J. A., H. M. Kharouba, M. Bontrager, C. Mahony, A. M. Csergő, A. M. E. Noreen, Q. Li, R. Schuster, and A. L. Angert. 2016. A synthesis of transplant experiments and ecological niche models suggests that range limits are often niche limits. Ecol. Lett. 19:710–722.

Leibold, M. A. 1995. The niche concept revisited: mechanistic models and community context. Ecology 76:1371–1382.

Lenz, A., G. Hoch, Y. Vitasse, and C. Körner. 2013. European deciduous trees exhibit similar safety margins against damage by spring freeze events along elevational gradients. New Phytol. 200:1166–1175.

Li, D., and A. R. Ives. 2017. The statistical need to include phylogeny in trait-based analyses of community composition. Methods Ecol. Evol. 8:1192–1199.

Litsios, G., A. Kostikova, and N. Salamin. 2014. Host specialist clownfishes are environmental niche generalists. Proc. R. Soc. B. 281:20133220.

Loehle, C. 1998. Height growth rate tradeoffs determine northern and southern range limits for trees. J. Biogeogr. 25:735–742.

Louthan, A. M., D. F. Doak, and A. L. Angert. 2015. Where and when do species interactions set range limits? Trends Ecol. Evol. 30:780–792.

Luckinbill, L. S. 1998. Selection for longevity confers resistance to low-temperature stress in *Drosophila melanogaster*. J. Gerontol. Ser. A 53A:B147–B153.

Luxbacher, A. M., and J. H. Knouft. 2009. Assessing concurrent patterns of environmental niche and morphological evolution among species of horned lizards (*Phrynosoma*). J. Evol. Biol. 22:1669–1678.

MacArthur, R. H. 1972. Geographical ecology. Harper & Row, Princeton Univ. Press, Princton, NJ.

Macek, P., J. Macková, and F. de Bello. 2009. Morphological and ecophysiological traits shaping altitudinal distribution of three *Polylepis* treeline species in the dry tropical Andes. Acta Oecologica 35:778–785.

Mächler, F., and J. Nösberger. 1977. Effect of light intensity and temperature on apparent photosynthesis of altitudinal ecotypes of *Trifolium repens* L. Oecologia 31:73–78.

Makowski, D., Ben-Shachar, M. S., & Lüdecke, D. (2019). bayestestR: Describing effects and their uncertainty, existence and significance within the Bayesian framework. Journal of Open Source Software, 4, 1541.

Mason, C. M., and L. A. Donovan. 2015. Evolution of the leaf economics spectrum in herbs: evidence from environmental divergences in leaf physiology across *Helianthus* (Asteraceae). Evolution 69:2705–2720.

Midolo, G., P. D. Frenne, N. Hölzel, and C. Wellstein. 2019. Global patterns of intraspecific leaf trait responses to elevation. Glob. Change Biol. 25:2485–2498.

Mockett, R. J., and R. S. Sohal. 2006. Temperature-dependent trade-offs between longevity and fertility in the *Drosophila* mutant, methuselah. Exp. Gerontol. 41:566–573.

Molina-Montenegro, M. A., J. Gallardo-Cerda, T. S. M. Flores, and C. Atala. 2012. The trade-off between cold resistance and growth determines the *Nothofagus pumilio* treeline. Plant Ecol. 213:133–142.

Morin, X., C. Augspurger, and I. Chuine. 2007. Process-based modeling of species’ distributions: what limits temperate tree species’ range boundaries? Ecology 88:2280–2291.

Mousseau, T. A., D. A. Roff. 1987. Natural selection and the heritability of fitness components. Heredity 59:181–197.

Moser D., Gygax A., Bäumier B., Wyler N., Palese R. 2002. Rote Liste der gefährdeten Farn- und Blütenpflanzen der Schweiz. BAFU, Switzerland.

Neuner, G. 2014. Frost resistance in alpine woody plants. Front. Plant Sci. 5:654.

Neuner, G., B. Huber, A. Plangger, J.-M. Pohlin, and J. Walde. 2020. Low temperatures at higher elevations require plants to exhibit increased freezing resistance throughout the summer months. Environ. Exp. Bot. 169:103882.

Normand, S., U. A. Treier, C. Randin, P. Vittoz, A. Guisan, and J.-C. Svenning. 2009. Importance of abiotic stress as a range-limit determinant for European plants: insights from species responses to climatic gradients. Glob. Ecol. Biogeogr. 18:437–449.

Norry, F. M., F. H. Gomez, and V. Loeschcke. 2007. Knockdown resistance to heat stress and slow recovery from chill coma are genetically associated in a quantitative trait locus region of chromosome 2 in *Drosophila melanogaster*. Mol. Ecol. 16:3274–3284.

Norry, F. M., and V. Loeschcke. 2002. Temperature-induced shifts in associations of longevity with body size in *Drosophila melanogaster*. Evolution 56:299–306.

Orme, D., R. Freckleton, T. Gavin, T. Petzoldt, S. Fritz, N. Isaac, and W. Pearse. 2018. caper: comparative analyses of phylogenetics and evolution in R. R package version 1.0.1.

Patsiou, T. S., N. Walden, and Y. Willi. 2021. What drives species distribution along elevational gradients? Macroecological and -evolutionary insights from Brassicaceae species of central Alps. What drives species distribution along elevational gradients? Global Ecol. Biogeogr. 30:1030–1042

Pérez, F., L. F. Hinojosa, C. G. Ossa, F. Campano, and F. Orrego. 2014. Decoupled evolution of foliar freezing resistance, temperature niche and morphological leaf traits in Chilean *Myrceugenia*. J. Ecol. 102:972–980.

Pérez-Harguindeguy, N., S. Díaz, E. Garnier, S. Lavorel, H. Poorter, P. Jaureguiberry, et al and J. H. C. Cornelissen. 2013. New handbook for standardised measurement of plant functional traits worldwide. Aust. J. Bot. 61:167–234.

Pianka, E. R. 1970. On r- and K-selection. Am. Nat. 104:592–597.

Pigott, C. D., and J. P. Huntley. 1981. Factors controlling the distribution of *Tilia cordata* at the northern limits of its geographical range III. Nature and causes of seed sterility. New Phytol. 87:817–839.

Pierce, S., Brusa, G., Vagge, I., and Cerabolini, B. E. 2013. Allocating CSR plant functional types: the use of leaf economics and size traits to classify woody and herbaceous vascular plants. Funct. Ecol., 27:1002–1010.

Pither, J. 2003. Climate tolerance and interspecific variation in geographic range size. Proc. R. Soc. B. 270:475–481.

Qi, W., H. Bu, K. Liu, W. Li, J. M. H. Knops, J. Wang, W. Li, and G. Du. 2014. Biological traits are correlated with elevational distribution range of eastern Tibetan herbaceous species. Plant Ecol. 215:1187–1198.

R Core Team. 2014. A language and environment for statistical computing. R Foundation for Statistical Computing, Vienna. Vienna Austria.

Rizzo, M. L., and G. J. Székely. 2016. Energy distance. WIREs Comput. Stat. 8:27–38.

Rodrigues, J. F. M., F. Villalobos, J. B. Iverson, J. Alexandre, and J. A. F. Diniz-Filho. 2019. Climatic niche evolution in turtles is characterized by phylogenetic conservatism for both aquatic and terrestrial species. J. Evol. Biol. 32:66–75.

Roff, D. A., and D. J. Fairbairn. 2012. A test of the hypothesis that correlational selection generates genetic correlations. Evolution 66:2953–2960.

Root, T. 1988. Energy Constraints on avian distributions and abundances. Ecology 69:330–339.

Rosbakh, S., M. Bernhardt-Römermann, and P. Poschlod. 2014. Elevation matters: contrasting effects of climate change on the vegetation development at different elevations in the Bavarian Alps. Alp. Bot. 124:143–154.

Rose, M. R. 1983. Further models of selection with antagonistic pleiotropy. Pp. 47–53 *in* H. I. Freedman and C. Strobeck, eds. Population Biology. Springer, Berlin Heidelberg, Germany.

Rosenzweig, M. L., and R. M. L. 1995. Species diversity in space and time. Cambridge Univ. Press, Cambridge, U.K.

Sakai, A., and W. Larcher. 2012. Frost survival of plants: responses and adaptation to freezing stress. Springer Science & Business Media.

Salisbury, E. J. 1926. The geographical distribution of plants in relation to climatic factors. Geogr. J. 67:312–335.

Schluter, D. 2001. Ecology and the origin of species. Trends Ecol. Evol. 16:372–380.

Schrieber, K., Y. Cáceres, A. Engelmann, P. Marcora, D. Renison, I. Hensen, and C. Müller. 2020. Elevational differentiation in metabolic cold stress responses of an endemic mountain tree. Environ. Exp. Bot. 171:103918.

Sgrò, C. M., and A. A. Hoffmann. 2004. Genetic correlations, tradeoffs and environmental variation. Heredity 93:241–248.

Stoks, R., and M. De Block. 2011. Rapid growth reduces cold resistance: evidence from latitudinal variation in growth rate, cold resistance and stress proteins. PLoS ONE 6:e16935.

Sunday, J. M. Bennett, P. Calosi, S. Clusella-Trullas, S. Gravel, A. L. Hargreaves, F. P. Leiva, W. C. E. P. Verberk, M. Á. Olalla-Tárraga, and I. Morales-Castilla. 2019. Thermal tolerance patterns across latitude and elevation. Philos. Trans. R. Soc. B. 374:20190036.

Sunday, J. M., A. E. Bates, and N. K. Dulvy. 2012. Thermal tolerance and the global redistribution of animals. Nat. Clim. Change 2:686–690.

Sutinen, M.-L., R. Arora, M. Wisniewski, E. Ashworth, R. Strimbeck, and J. Palta. 2001. Mechanisms of frost survival and freeze-damage in nature. Pp. 89–120 *in* F. J. Bigras and S. J. Colombo, eds. Conifer cold hardiness. Springer, Dordrecht, Netherlands.

Taschler, D., and G. Neuner. 2004. Summer frost resistance and freezing patterns measured *in situ* in leaves of major alpine plant growth forms in relation to their upper distribution boundary. Plant Cell Environ. 27:737–746.

Thakur, D., N. Rathore, and A. Chawla. 2019. Increase in light interception cost and metabolic mass component of leaves are coupled for efficient resource use in the high altitude vegetation. Oikos 128:254–263.

Tonnabel, J., F. M. Schurr, F. Boucher, W. Thuiller, J. Renaud, E. J. P. Douzery, and O. Ronce. 2018. Life-history traits evolved jointly with climatic niche and disturbance regime in the genus *Leucadendron* (Proteaceae). Am. Nat. 191:220–234.

Urban, M. C. 2007. The growth–predation risk trade-off under a growing gape-limited predation threat. Ecology 88:2587–2597.

van Noordwijk, A. J., and G. de Jong. 1986. Acquisition and allocation of resources: their influence on variation in life history tactics. Am. Nat. 128:137–142.

Vetaas, O. R. 2002. Realized and potential climate niches: a comparison of four *Rhododendron* tree species. J. Biogeogr. 29:545–554.

Wagner, G. P., and K. Schwenk. 2000. Evolutionarily stable configurations: functional integration and the evolution of phenotypic stability. Pp. 155–217 *in* M. K. Hecht, R. J. Macintyre, and M. T. Clegg, eds. Evolutionary Biology. Springer US, Boston, MA.

Walker, J. A. 2007. A general model of functional constraints on phenotypic evolution. Am. Nat. 170:681–689.

Wen, Y., D. Qin, B. Leng, Y. Zhu, and K. Cao. 2018. The physiological cold tolerance of warm-climate plants is correlated with their latitudinal range limit. Biol. Lett. 14:20180277.

Willi, Y., and J. Van Buskirk. 2019. A practical guide to the study of distribution limits. Am. Nat. 193:773–785.

Williams, B. R., B. V. Heerwaarden, D. K. Dowling, and C. M. Sgrò. 2012. A multivariate test of evolutionary constraints for thermal tolerance in *Drosophila melanogaster*. J. Evol. Biol. 25:1415–1426.

Wright, I. J., P. B. Reich, M. Westoby, D. D. Ackerly, Z. Baruch, F. Bongers, J. et al. and R. Villar. 2004. The worldwide leaf economics spectrum. Nature 428:821–827.

Wos, G., and Y. Willi. 2015. Temperature-stress resistance and tolerance along a latitudinal cline in North American *Arabidopsis lyrata*. PLOS ONE 10:e0131808.

Zhong, M., J. Wang, K. Liu, R. Wu, Y. Liu, X. Wei, D. Pan, and X. Shao. 2014. Leaf morphology shift of three dominant species along altitudinal gradient in an alpine meadow of the Qinghai-Tibetan plateau. Pol. J. Ecol. 62:639–648.

