## Supplementary material for "Trait divergence and trade-offs among Brassicaceae species differing in elevational distribution": A2_SupplementaryMethods_And_GrowthTraj_ForSub.pdf

**Supplemental material A2: Supplementary methods**

**Trait assessment**

*Seed size (SSIZ)* - For all species, 10-20 seeds per field-collected maternal plant were haphazardly selected. When possible, the same maternal lines were used as for sowing. If not enough seeds were available, another maternal line was randomly picked. Seeds were then photographed under a stereomicroscope (Leica M205 C, Leica Microsystems GmbH, Wetzlar, Germany), and the area of each seed was measured (in $mm^2$) with the software ImageJ v.1.44 (Schneider et al. 2012).

*Time to germination (TGER)* - After stratification, seeds were checked daily for germination during the first week and every second day until the start of treatment. Germination was defined as when two cotyledons were fully open. Time to germination was the number of days between the end of stratification and the day of germination.

*Growth (IGR, MGR, XMID, ASYM, NLEA)* - Plant growth was measured once a week for 5 weeks, starting the week before the temperature treatment began. The two longest leaves of every plant were measured (in mm). Leaves with more than 25% damage or senescence were not considered. Means across leaves were used to estimate the growth trajectory by fitting seven alternative growth models (linear, exponential, power, two-, three-parameter logistic, Gompertz and Bertalanaffy). Models were fit with 'nlsLM' {minpack.lm} (Elzhov et al. 2016) using the base functions 'SSlogis', 'SSgompertz' and 'vbT' {FSA} (Ogle 2017), and AIC values were extracted with 'aictab' {AICcmodavg} (Mazreolle 2017). The Gompertz model and the three-parameter logistic model were the two best-supported based on AIC (Fig. A2.1). Parameters of the three-parameter logistic model were extracted as their interpretation is more intuitive: asymptotic size (ASYM), maximal growth rate (MGR, i.e., $scale^{-1}$) and the time to fastest growth (XMID). Since smaller values of XMID meant that a plant achieved mid-size faster, values were multiplied by -1 ([-1XMID]) to represent speed of growth. The initial growth rate (IGR) was calculated as the derivative with the 'deriv' function at the bend in the exponential phase of the curve located by the 'maxcurve' function {soilphysics} (da Silva and de Lima. 2017). The number of leaves was additionally counted once a week, but since only few species reached a final asymptote, the number of leaves on day 35 of treatment was used in analysis (NLEA).

*Leaf functional traits (LA, SLA, LDMC, LDI, LTh)* - During week 8 of treatment, 1-2 fully elongated leaves from the $2^{nd}$ to $3^{rd}$ whorl were harvested from each plant and used for leaf trait assessment. For very tiny leaves (< 25 $mm^2$) twice the amount of material was collected. Leaves were immediately weighed individually on microbalances (AT250, XA205 DualRange, Mettler Toledo, Columbus, USA) to the nearest 0.01 mg. Leaves were scanned (CanonScan, LiDe120, Canon, Tokyo, Japan) and analysed by ImageJ to obtain leaf area (LA, in $mm^2$) and perimeter (in mm). Leaves were then packed in paper bags and dried at 60 °C for 72 h in an oven (Termaks AS, Bergen, Norway), and weighed again using the same balance. Specific leaf area was calculated as area over dry weight (SLA, in $mm^2$ $mg^{-1}$) and leaf dry matter content as the ratio of dry weight over fresh weight (LDMC, in mg $g^{-1}$). Leaf dissection index (LDI) was calculated as the ratio of perimeter to area following Fourier's transformation (Kincaid and Schneider 1983); smaller values indicated less dissected leaves. Leaf thickness was estimated right after harvesting using a mini-digital thickness gauge (digitalmicrometers.co.uk) (LTh, in mm). Traits were assessed on individual leaves, and their mean per plant was calculated.

*Frost and heat resistance (electrolyte leakage, RES)* - During week 9 of treatment, 3 healthy and fully developed leaves from the $3^{rd}$ to $5^{th}$ whorl were picked and a circular disc of 6 mm diameter punched out on the tip of the blade, avoiding main nervures. Each leaf disc was placed in a 15 ml falcon tube (Sarstedt, PP, 120x17 mm, Nümbrecht, Germany) filled with 2 ml of $dH_2O$ and kept in there for 0.5-1 h to wash the sample. The water was discarded, and the leaf in each tube was exposed to short thermal stress. Based on a pilot study and reports in the literature (e.g., Levitt 1980; Kappen 1981; Gauslaa 1984), the temperatures of -10 °C (*minus*T2) and -5 °C (*minus*T1) were chosen to assess frost resistance, and +45 °C (*plus*T1) and +50 °C (*plus*T2) to assess heat resistance. Resistance to T1 was tested only on non-acclimated plants (i.e., plants of the mild growth treatment), while T2 was tested on non-acclimated and acclimated plants (i.e., plants pre-exposed to frost for assessing frost resistance, and plants pre-exposed to heat for assessing heat resistance).

Resistance was assessed based on a modified protocol of Pérez-Harguindeguy *et al.* (2013). Frost exposure was applied by first filling tubes into aluminium boxes (to buffer thermal variation), which were then placed in programmable freezers, one per negative temperature. After an initial 0.5 h at 5 °C for temperature equilibration, the target temperature was approached with a cooling rate of -3 K $h^{-1}$. Samples were kept at the target temperature for 1 h, and then temperature was increased to +5 °C. Heat exposure was done in a water-bath (Julabo TW20, HuberLab, Aesch, Switzerland) by submersing tubes for 5 min in the bath for temperature equilibration. Then tubes were kept in the bath for 1 h, in dark. After frost and heat exposure, all tubes received 3 ml of $dH_2O$ and were stored overnight in dark and at room temperature for electrolytes to dissolve in the water. Electrolyte concentration was measured with a calibrated conductivity meter (Fe30/EL30, Mettler Toledo, Columbus, USA). Then tubes were sealed and subjected to a boiling bath for 1 h. Tubes were again kept overnight, and total leakage was measured on the following day. Electrolyte leakage due to stress was calculated as the ratio of conductivity measured after stress to conductivity after the boiling bath, in per cent. Resistance was calculated by the formula: 100% - electrolyte leakage, with higher values indicating higher resistance.

*Frost and heat tolerance (TOL)* - Tolerance to repeated frost or heat during the growth phase was calculated as growth parameter (MGR, -XMID, ASYM) under frost or heat treatment minus the estimate in the mild treatment, divided by the estimate in the mild treatment (relative measure of tolerance). Negative values meant that a plant was unable to maintain size or speed of growth under frost or heat stress; it was less tolerant.

**Past evolutionary forces: ancestral state reconstruction and validation methods**

*Ancestral state reconstruction* - To perform ancestral state reconstruction, regimes were assigned to each species, and two models with increasing parameterization, i.e., one of equal rate of evolutionary switch (ER) and one of unequal rate (ARD), were run on the full phylogenetic tree with 'fitMk' function in phytools (Revell 2012). Models were compared based on AIC across 100 simulations. The best model was selected for ancestral state reconstruction on 100 independent stochastic character maps on the phylogenetic tree with 'make.simmap' in phytools (Revell 2012). The initial tree with 125 tips and 124 internal nodes was pruned with 'treedata' {geiger} and 'drop.tip.simmap'{phytools} and un-measured taxa (e.g., outgroups, not germinate) were removed.

*Past evolutionary forces and validation methods* - To validate whether we could distinguish accurately among evolutionary models given our phylogeny, we generated a synthetic distribution of trait data across all taxa by assuming one of four evolutionary scenarios, i.e., BM, BMM, OU, or OUM. Data were generated with the function 'mvSIM' {mvMORPH} and by using different initial parameterizations in the range of observed values based on our results, and on an increasing gradient of trait variance. For each data set, the four evolutionary scenarios plus a WN model were fit and the number of times the evolutionary process was correctly identified was counted on 1000 simulations for each data set (Supplementary material A4).

Furthermore, to assess consistency in results given that the species included in the study were a spare representation of a species-rich plant family, we additionally used a bootstrap approach of removing one third of the species randomly 100 times (keeping at least 50% of the alpine species to allow the fiting of multi-regime models). The process was repeated for each of the 100 stochastic character map reconstructions, resulting in 10'000 bootstrap simulations for each trait-treatment combination. Evolutionary models were again fit and compared based on these 10'000 simulations, and the model with the lowest mean AICc value was considered the best (Supplementary material A4). Trait divergence among species were run separately for the three temperature treatments, and by considering variance in trait means of species of the two rounds of sowing. Where species variance could not be calculated for a specific trait, because it was assessed only in one round of sowing, its value was derived from the mean variance across species within the treatment.

**References**


da Silva, A.R., and R. P., de Lima. 2017. Determination of maximum curvature point with the R package soilphysics. Int. J. Curr. Res. 9:45241–45245.

Elzhov, T. V., Mullen, K. M., Spiess, A. N., Bolker, B., and M. K. M, Mullen. 2016. Package 'minpack.lm' R interface Levenberg-Marquardt Nonlinear least-Sq. Algorithm Found MINPACK Plus Support Bounds.

Gauslaa, Y. 1984. Heat resistance and energy budget in different scandinavian plants. Ecography 7:5–6.

Kappen, L. 1981. Ecological significance of resistance to high temperature. Pp. 439–474 *in* O. L. Lange, P. S. Nobel, C. B. Osmond, and H. Ziegler, eds. *Physiological plant ecology I: responses to the physical environment*. Springer, Berlin Heidelberg, Germany.

Kincaid, D. T., and R. B. Schneider. 1983. Quantification of leaf shape with a microcomputer and Fourier transform. Can. J. Bot. 61:2333–2342.

Levitt, J. 1980. Responses of plants to environmental stresses. Vol 1. Chilling, freezing, and high temperature stresses. Academic Press, New York.

Mazerolle, M. J. 2017. Package 'AICcmodavg'. R package.

Ogle, D. 2017. Package 'FSA'. CRAN Repos, 1-206.

Pérez-Harguindeguy, N., Diaz, S., Gamier, E., Lavorel, S., Poorter, H., Jaureguiberry, P., ... & Cornelissen, J. H. C. Cornelissen (2013). New handbook for standardised measurement of plant functional traits worldwide. Aus-tralian Journal of Botany 61: 167-234.

Revell, L. J. 2012. phytools: an R package for phylogenetic comparative biology (and other things). Methods Ecol. Evol. 3:217–223.

Schneider, C. A., W. S. Rasband, and K. W. Eliceiri. 2012. NIH Image to ImageJ: 25 years of image analysis. Nat. Methods 9:671–675.

***Next page.*** **Figure 1.** AIC comparisons across models describing the growth trajectory of plants

Comparison of growth models across all plants raised (S1, across treatment)

Based on leaf length

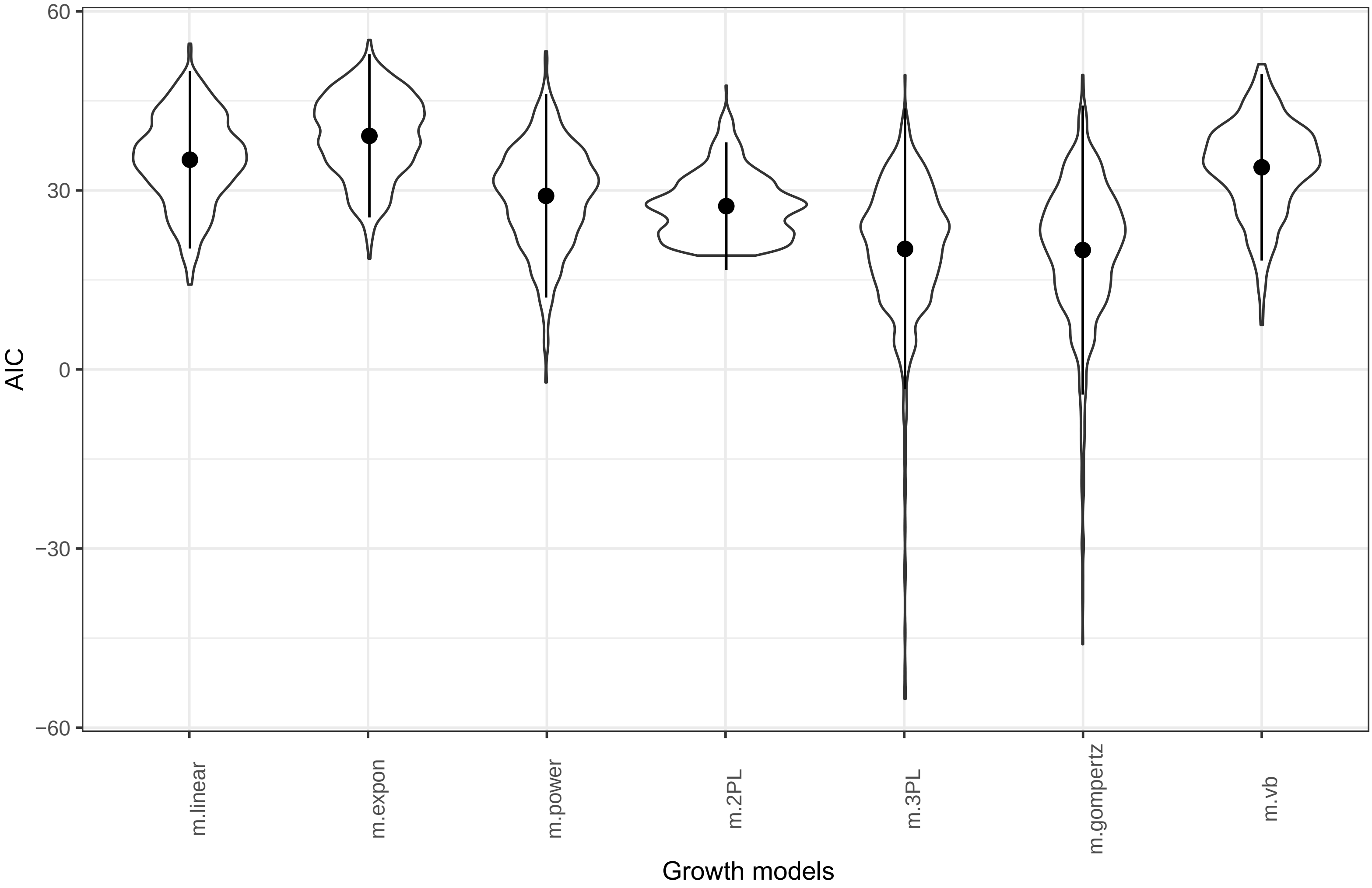


Comparison of growth models across all plants raised (S1, split by treatment)

Based on leaf length

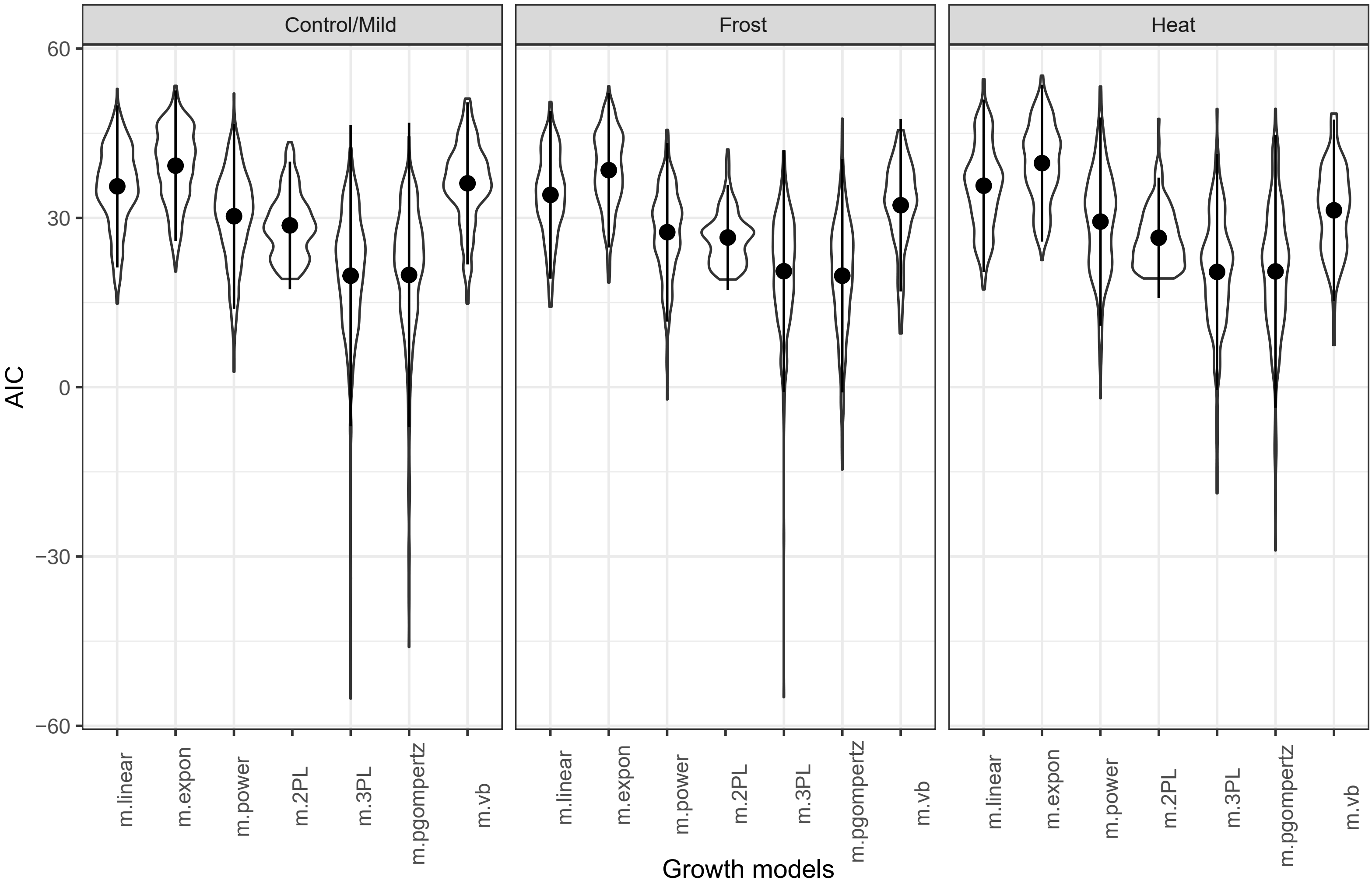

### Comparison of growth models across all plants raised (S2, across treatments)

Based on leaf length

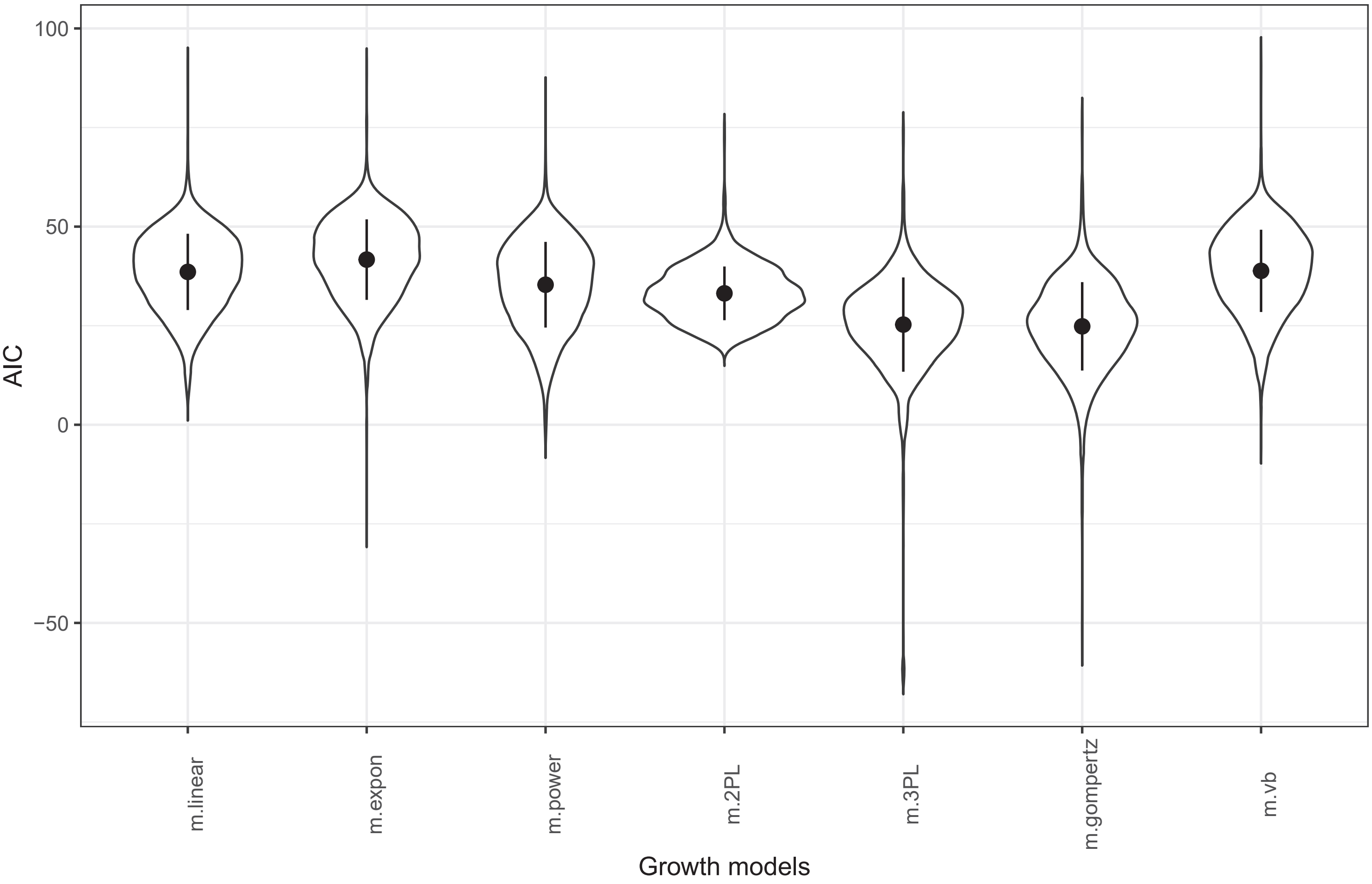


### Comparison of growth models across all plants raised (S2, split by treatment)

Based on leaf length

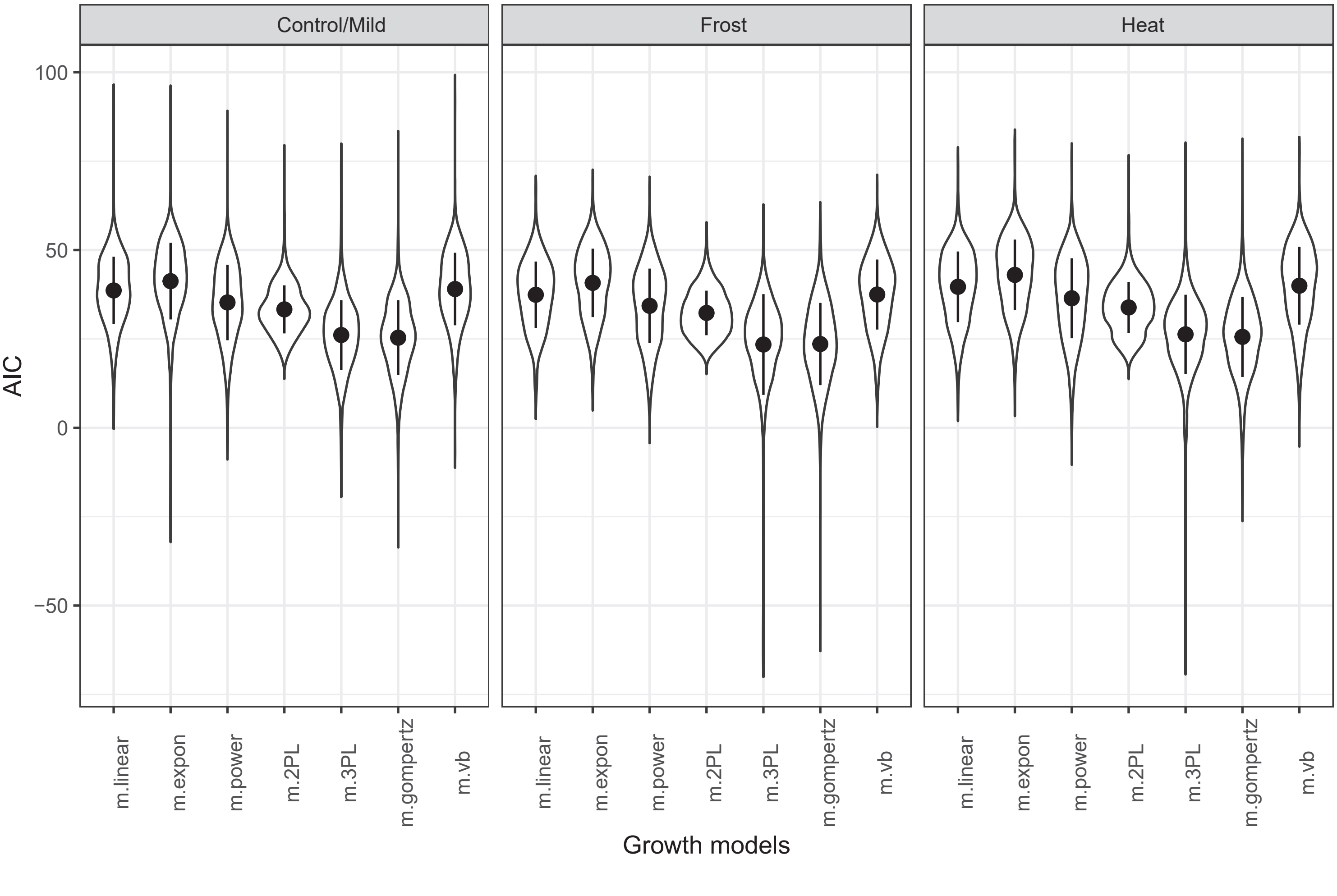
