## Supplementary material for "Trait divergence and trade-offs among Brassicaceae species differing in elevational distribution": A3_M01_MixedEffectsModels_ForSub.pdf

### A3 – Supplementary material: Mixed-effects models

Traits that showed a significant effect of elevation or an interaction effect of elevation-by-treatment are written in red. Effects and their model estimates are highlighted in yellow if there were differences between the model accounting for phylogeny (left) and the model that did not (right). Effects that were significant are written in bold (95%ETI not containing 0). Posterior medians of fixed effects are reported, relative to the baseline of average elevation and mild growth conditions and, for tolerance traits, the coefficients express differences between estimates under heat compared to those under frost. For random effects, reporting includes the within-group variance (sigma-squared), the between-group variance (tau-zero-zero), and the number of levels, N (i.e., species and/or round of sowing).

##### 1. Bayesian mixed-effects model for: SSIZ

| *Predictors* | *Median* | *CI (95%)* | *Median* | *CI (95%)* |
|---|---|---|---|---|
| **Intercept** | **0.9933** | **0.1471 – 1.8284** | **0.3614** | **0.0911 – 0.6306** |
| Elevation | 0.0583 | -0.1659 – 0.2802 | -0.1134 | -0.3844 – 0.1520 |
| **Random effects** | | | | |
| $\sigma^2$ | 1795.1641 | | | |
| $\tau_{00}$ | 1.1661 | | | |
| N | 95 taxa | | | |
| Observations | 95 | | 95 | |

##### 2. Bayesian mixed-effects model for: TGER

| *Predictors* | *Median* | *CI (95%)* | *Median* | *CI (95%)* |
|---|---|---|---|---|
| **Intercept** | **2.5841** | **1.6294 – 3.1756** | **2.4456** | **0.8774 – 3.2305** |
| Elevation | 0.0426 | -0.0322 – 0.1142 | 0.0547 | -0.0060 – 0.1157 |
| **Random effects** | | | | |
| $\sigma^2$ | 0.3580 | | 0.3989 | |
| $\tau_{00}$ | 0.2033 sowing_id | | 0.5075 sowing_id | |
| | 0.0133 taxa | | | |
| N | 93 taxa | | 2 sowing_id | |
| | 2 sowing_id | | | |
| Observations | 168 | | 168 | |

**3. Bayesian mixed-effects model for: IGR**

| *Predictors* | *Median* | *CI (95%)* | *Median* | *CI (95%)* |
|---|---|---|---|---|
| Intercept | -0.5728 | -1.5335 – 0.1254 | -0.6115 | -1.9583 – 0.4005 |
| Elevation | -0.0088 | -0.0325 – 0.0143 | -0.0090 | -0.0316 – 0.0143 |
| Treatment (Frost vs Mild) | -0.0061 | -0.0377 – 0.0250 | -0.0062 | -0.0387 – 0.0265 |
| Treatment (Heat vs Mild) | -0.0190 | -0.0513 – 0.0125 | -0.0191 | -0.0505 – 0.0140 |
| Elevation x Frost treatment vs Mild | 0.0124 | -0.0185 – 0.0442 | 0.0125 | -0.0202 – 0.0449 |
| Elevation x Heat treatment vs Mild | 0.0149 | -0.0172 – 0.0468 | 0.0154 | -0.0169 – 0.0476 |
| **Random effects** | | | | |
| $\sigma^2$ | 0.0004 | | 0.0006 | |
| $\tau_{00}$ | 0.0039 | | 0.0036 | |
| N | 90 taxa | | 1 sowing_id | |
| | 1 sowing_id | | | |
| Observations | 263 | | 263 | |

**4. Bayesian mixed-effects model for: MGR**

| *Predictors* | *Median* | *CI (95%)* | *Median* | *CI (95%)* |
|---|---|---|---|---|
| **Intercept** | **-1.5334** | **-2.3955 – -0.6486** | **-1.5233** | **-2.4030 – -0.6429** |
| Elevation | -0.0296 | -0.0989 – 0.0392 | -0.0351 | -0.1026 – 0.0328 |
| Treatment (Frost vs Mild) | -0.0409 | -0.1394 – 0.0540 | -0.0419 | -0.1401 – 0.0556 |
| **Treatment (Heat vs Mild)** | **0.4227** | **0.3233 – 0.5221** | **0.4206** | **0.3201 – 0.5214** |
| Elevation x Frost treatment vs Mild | 0.0129 | -0.0801 – 0.1071 | 0.0123 | -0.0825 – 0.1084 |
| **Elevation x Heat treatment vs Mild** | **0.1292** | **0.0312 – 0.2240** | **0.1282** | **0.0331 – 0.2251** |
| **Random effects** | | | | |
| $\sigma^2$ | 0.0086 | | 0.0076 | |
| $\tau_{00}$ | 0.0312 | | 0.0318 | |
| N | 92 taxa | | 2 sowing_id | |
| | 2 sowing_id | | | |
| Observations | 467 | | 467 | |

**5. Bayesian mixed-effects model for: XMID**

| *Predictors* | *Median* | *CI (95%)* | *Median* | *CI (95%)* |
|---|---|---|---|---|
| **Intercept** | **3.1455** | **1.8422 – 3.7487** | **3.1763** | **1.9100 – 3.7803** |
| Elevation | 0.0189 | -0.0133 – 0.0514 | 0.0189 | -0.0080 – 0.0469 |
| Treatment (Frost vs Mild) | -0.0330 | -0.0684 – 0.0020 | -0.0332 | -0.0738 – 0.0050 |
| **Treatment (Heat vs Mild)** | **-0.1146** | **-0.1502 – -0.0785** | **-0.1131** | **-0.1534 – -0.0741** |
| Elevation x Frost treatment vs Mild | -0.0040 | -0.0393 – 0.0314 | -0.0053 | -0.0441 – 0.0340 |
| **Elevation x Heat treatment vs Mild** | **-0.0650** | **-0.1015 – -0.0282** | **-0.0641** | **-0.1042 – -0.0220** |
| **Random effects** | | | | |
| $\sigma^2$ | 29.3880 | | 24.1119 | |
| $\tau_{00}$ | 14.7235 | | 19.9767 | |
| N | 92 taxa | | 2 sowing_id | |
| | 2 sowing_id | | | |
| Observations | 460 | | 460 | |

**6. Bayesian mixed-effects model for: ASYM**

| *Predictors* | *Median* | *CI (95%)* | *Median* | *CI (95%)* |
|---|---|---|---|---|
| **Intercept** | **3.8519** | **3.4457 – 4.2379** | **3.8260** | **3.5206 – 4.1236** |
| **Elevation** | -0.0403 | -0.1234 – 0.0472 | **-0.2531** | **-0.3296 – -0.1783** |
| **Treatment (Frost vs Mild)** | **-0.1394** | **-0.1973 – -0.0845** | **-0.1383** | **-0.2511 – -0.0318** |
| **Treatment (Heat vs Mild)** | **-0.1526** | **-0.2118 – -0.0956** | **-0.1338** | **-0.2387 – -0.0257** |
| Elevation x Frost treatment vs Mild | -0.0256 | -0.0822 – 0.0314 | -0.0304 | -0.1393 – 0.0760 |
| **Elevation x Heat treatment vs Mild** | **-0.1349** | **-0.1931 – -0.0757** | **-0.1344** | **-0.2444 – -0.0248** |
| **Random effects** | | | | |
| $\sigma^2$ | 597.2925 | | 3.6393 | |
| $\tau_{00}$ | 166.9919 | | 1007.9378 | |
| N | 92 taxa | | 2 sowing_id | |
| | 2 sowing_id | | | |
| Observations | 460 | | 460 | |

**7. Bayesian mixed-effects model for: NLEA**

| *Predictors* | *Median* | *CI (95%)* | *Median* | *CI (95%)* |
|---|---|---|---|---|
| **Intercept** | **2.7193** | **1.9778 – 3.6339** | **2.7541** | **2.0872 – 3.7375** |
| Elevation | -0.0107 | -0.1272 – 0.1028 | 0.0287 | -0.0940 – 0.1511 |
| **Treatment (Frost vs Mild)** | **-0.0642** | **-0.1226 – -0.0054** | -0.0581 | -0.2304 – 0.1085 |
| Treatment (Heat vs Mild) | 0.0047 | -0.0544 – 0.0638 | 0.0137 | -0.1619 – 0.1813 |
| Elevation x Frost treatment vs Mild | -0.0089 | -0.0660 – 0.0498 | -0.0000 | -0.1678 – 0.1669 |
| Elevation x Heat treatment vs Mild | -0.0247 | -0.0815 – 0.0334 | -0.0179 | -0.1898 – 0.1539 |
| **Random effects** | | | | |
| $\sigma^2$ | 102.8772 | | -61.7739 | |
| $\tau_{00}$ | 10.2023 | | 150.9636 | |
| N | 89 taxa | | 1 sowing_id | |
| | 1 sowing_id | | | |
| Observations | 257 | | 257 | |

**8. Bayesian mixed-effects model for: LA**

| *Predictors* | *Median* | *CI (95%)* | *Median* | *CI (95%)* |
|---|---|---|---|---|
| **Intercept** | **3.5008** | **2.4944 – 4.5804** | **3.4708** | **2.4693 – 4.5139** |
| **Elevation** | **-0.3482** | **-0.5105 – -0.1898** | **-0.6721** | **-0.8112 – -0.5295** |
| **Treatment (Frost vs Mild)** | **-0.1618** | **-0.2617 – -0.0619** | -0.1320 | -0.3331 – 0.0668 |
| **Treatment (Heat vs Mild)** | -0.2548 | -0.3541 – -0.1543 | **-0.2125** | **-0.4113 – -0.0073** |
| Elevation x Frost treatment vs Mild | 0.0325 | -0.0713 – 0.1322 | 0.0257 | -0.1805 – 0.2276 |
| **Elevation x Heat treatment vs Mild** | **-0.1171** | **-0.2216 – -0.0125** | -0.1269 | -0.3312 – 0.0784 |
| **Random effects** | | | | |
| $\sigma^2$ | 180945.8460 | | 334942.1002 | |
| $\tau_{00}$ | 395.7070 | | 4761.9862 | |
| N | 93 taxa | | 2 sowing_id | |
| | 2 sowing_id | | | |
| Observations | 434 | | 434 | |

**9. Bayesian mixed-effects model for: SLA**

| *Predictors* | *Median* | *CI (95%)* | *Median* | *CI (95%)* |
|---|---|---|---|---|
| **Intercept** | **3.0456** | **2.8726 – 3.2001** | **3.0949** | **2.8723 – 3.2247** |
| Elevation | 0.0182 | -0.0474 – 0.0854 | 0.0511 | -0.0032 – 0.1033 |
| **Treatment (Frost vs Mild)** | **-0.1301** | **-0.1948 – -0.0641** | **-0.1262** | **-0.2035 – -0.0481** |
| **Treatment (Heat vs Mild)** | **0.1121** | **0.0457 – 0.1789** | **0.1151** | **0.0341 – 0.1941** |
| Elevation x Frost treatment vs Mild | -0.0295 | -0.0942 – 0.0359 | -0.0291 | -0.1044 – 0.0498 |
| Elevation x Heat treatment vs Mild | 0.0175 | -0.0495 – 0.0842 | 0.0355 | -0.0435 – 0.1153 |
| **Random effects** | | | | |
| $\sigma^2$ | 63.5872 | | 20.6432 | |
| $\tau_{00}$ | 53.3399 | | 97.3756 | |
| N | 93 $_{taxa}$ | | 2 $_{sowing\_id}$ | |
| | 2 $_{sowing\_id}$ | | | |
| Observations | 434 | | 434 | |

**10. Bayesian mixed-effects model for: LDMC**

| *Predictors* | *Median* | *CI (95%)* | *Median* | *CI (95%)* |
|---|---|---|---|---|
| **Intercept** | **2.9769** | **2.8141 – 3.1553** | **2.9575** | **2.8152 – 3.1424** |
| **Elevation** | **-0.0665** | **-0.1148 – -0.0181** | **-0.0864** | **-0.1241 – -0.0486** |
| Treatment (Frost vs Mild) | 0.0419 | -0.0001 – 0.0833 | 0.0432 | -0.0120 – 0.0989 |
| **Treatment (Heat vs Mild)** | **-0.1818** | **-0.2242 – -0.1403** | **-0.1727** | **-0.2279 – -0.1180** |
| Elevation x Frost treatment vs Mild | 0.0083 | -0.0339 – 0.0502 | 0.0031 | -0.0524 – 0.0570 |
| Elevation x Heat treatment vs Mild | -0.0198 | -0.0624 – 0.0237 | -0.0276 | -0.0838 – 0.0277 |
| **Random effects** | | | | |
| $\sigma^2$ | 11.6147 | | -1.3698 | |
| $\tau_{00}$ | 17.7749 | | 30.5715 | |
| N | 92 $_{taxa}$ | | 2 $_{sowing\_id}$ | |
| | 2 $_{sowing\_id}$ | | | |
| Observations | 423 | | 423 | |

**11. Bayesian mixed-effects model for: LTH**

| *Predictors* | *Median* | *CI (95%)* | *Median* | *CI (95%)* |
|---|---|---|---|---|
| **Intercept** | **-1.8316** | **-2.4362 – -1.2884** | **-1.9051** | **-2.5051 – -1.3061** |
| Elevation | 0.0256 | -0.0225 – 0.0750 | 0.0141 | -0.0342 – 0.0610 |
| **Treatment (Frost vs Mild)** | **0.0770** | **0.0178 – 0.1346** | **0.0764** | **0.0094 – 0.1423** |
| Treatment (Heat vs Mild) | -0.0167 | -0.0730 – 0.0407 | -0.0194 | -0.0858 – 0.0463 |
| Elevation x Frost treatment vs Mild | -0.0108 | -0.0690 – 0.0464 | -0.0120 | -0.0774 – 0.0553 |
| Elevation x Heat treatment vs Mild | 0.0004 | -0.0586 – 0.0600 | 0.0021 | -0.0639 – 0.0699 |
| **Random effects** | | | | |
| $\sigma^2$ | 0.0029 | | 0.0027 | |
| $\tau_{00}$ | 0.0011 | | 0.0012 | |
| N | 86 taxa | | 1 sowing_id | |
| | 1 sowing_id | | | |
| Observations | 241 | | 241 | |

**12. Bayesian mixed-effects model for: LDI**

| *Predictors* | *Median* | *CI (95%)* | *Median* | *CI (95%)* |
|---|---|---|---|---|
| Intercept | 0.4964 | -0.1029 – 0.8456 | 0.5314 | -0.0641 – 0.8177 |
| Elevation | 0.0393 | -0.0058 – 0.0863 | -0.0206 | -0.0612 – 0.0204 |
| Treatment (Frost vs Mild) | -0.0097 | -0.0360 – 0.0169 | -0.0045 | -0.0632 – 0.0536 |
| Treatment (Heat vs Mild) | -0.0170 | -0.0437 – 0.0099 | -0.0278 | -0.0875 – 0.0324 |
| Elevation x Frost treatment vs Mild | 0.0188 | -0.0078 – 0.0455 | 0.0128 | -0.0448 – 0.0718 |
| **Elevation x Heat treatment vs Mild** | **-0.0451** | **-0.0724 – -0.0171** | -0.0525 | -0.1117 – 0.0075 |
| **Random effects** | | | | |
| $\sigma^2$ | 0.2810 | | 0.0787 | |
| $\tau_{00}$ | 0.0420 | | 0.2207 | |
| N | 93 taxa | | 2 sowing_id | |
| | 2 sowing_id | | | |
| Observations | 434 | | 434 | |

**13. Bayesian mixed-effects model for: RESmT1_S1**

| *Predictors* | *Median* | *CI (95%)* | *Median* | *CI (95%)* |
|---|---|---|---|---|
| **Intercept** | **0.4235** | **0.2291 – 0.7446** | **0.3710** | **0.2409 – 0.4994** |
| Elevation | -0.0792 | -0.2351 – 0.0819 | -0.0854 | -0.2239 – 0.0523 |
| **Random effects** | | | | |
| $\sigma^2$ | 0.0028 | | | |
| $\tau_{00}$ | 0.0135 | | | |
| N | 62 $_{\text{taxa}}$ | | | |
| Observations | 63 | | 63 | |

**14. Bayesian mixed-effects model for: RESmT1_S2**

| *Predictors* | *Median* | *CI (95%)* | *Median* | *CI (95%)* |
|---|---|---|---|---|
| **Intercept** | **1.1050** | **0.9060 – 1.3166** | **1.0639** | **0.9863 – 1.1438** |
| Elevation | 0.0317 | -0.0543 – 0.1214 | 0.0072 | -0.0741 – 0.0907 |
| **Random effects** | | | | |
| $\sigma^2$ | 0.0022 | | | |
| $\tau_{00}$ | 0.0031 | | | |
| N | 87 $_{\text{taxa}}$ | | | |
| Observations | 87 | | 87 | |

**15. Bayesian mixed-effects model for: RESmT2_S1S2**

| *Predictors* | *Median* | *CI (95%)* | *Median* | *CI (95%)* |
|---|---|---|---|---|
| **Intercept** | **-0.9520** | **-1.4950 – -0.4063** | **-0.9403** | **-1.5043 – -0.3974** |
| **Elevation** | 0.1230 | -0.0064 – 0.2488 | **0.1271** | **0.0183 – 0.2375** |
| **Random effects** | | | | |
| $\sigma^2$ | 0.0476 | | 0.0451 | |
| $\tau_{00}$ | 0.0173 | | 0.0195 | |
| N | 91 $_{\text{taxa}}$ | | 2 $_{\text{sowing\_id}}$ | |
| | 2 $_{\text{sowing\_id}}$ | | | |
| Observations | 146 | | 146 | |

**16. Bayesian mixed-effects model for: RESmT2**

| *Predictors* | *Median* | *CI (95%)* | *Median* | *CI (95%)* |
|---|---|---|---|---|
| **Intercept** | 0.0288 | -0.4267 – 0.3972 | **0.4109** | **0.2950 – 0.5227** |
| Elevation | 0.0433 | -0.0933 – 0.1789 | 0.0179 | -0.0934 – 0.1285 |
| **Random effects** | | | | |
| $\sigma^2$ | 0.0065 | | | |
| $\tau_{00}$ | 0.0112 | | | |
| N | 87 $_{taxa}$ | | | |
| Observations | 87 | | 87 | |

**17. Bayesian mixed-effects model for: RESpT1_S1**

| *Predictors* | *Median* | *CI (95%)* | *Median* | *CI (95%)* |
|---|---|---|---|---|
| **Intercept** | **0.5506** | **0.1594 – 0.9724** | **0.3080** | **0.1530 – 0.4645** |
| Elevation | -0.0292 | -0.2017 – 0.1452 | 0.0327 | -0.1240 – 0.1884 |
| **Random effects** | | | | |
| $\sigma^2$ | 0.0140 | | | |
| $\tau_{00}$ | 0.0029 | | | |
| N | 46 $_{taxa}$ | | | |
| Observations | 46 | | 46 | |

**18. Bayesian mixed-effects model for: RESpT1_S2**

| *Predictors* | *Median* | *CI (95%)* | *Median* | *CI (95%)* |
|---|---|---|---|---|
| **Intercept** | **0.7355** | **0.5232 – 0.9764** | **0.7053** | **0.6137 – 0.8008** |
| Elevation | 0.0436 | -0.0625 – 0.1525 | -0.0095 | -0.1065 – 0.0881 |
| **Random effects** | | | | |
| $\sigma^2$ | 0.0035 | | | |
| $\tau_{00}$ | 0.0067 | | | |
| N | 87 $_{taxa}$ | | | |
| Observations | 87 | | 87 | |

**19. Bayesian mixed-effects model for: RESpT2_S1S2**

| *Predictors* | *Median* | *CI (95%)* | *Median* | *CI (95%)* |
|---|---|---|---|---|
| **Intercept** | **-1.2419** | **-1.6665 – -0.6740** | **-1.2997** | **-1.6170 – -0.7160** |
| Elevation | 0.0259 | -0.1764 – 0.2121 | 0.0924 | -0.0575 – 0.2325 |
| **Random effects** | | | | |
| $\sigma^2$ | 0.0030 | | -0.0019 | |
| $\tau_{00}$ | 0.0202 | | 0.0260 | |
| N | 86 $_{\text{taxa}}$ | | 2 $_{\text{sowing\_id}}$ | |
| | 2 $_{\text{sowing\_id}}$ | | | |
| Observations | 132 | | 132 | |

**20. Bayesian mixed-effects model for: RESpT2_S2**

| *Predictors* | *Median* | *CI (95%)* | *Median* | *CI (95%)* |
|---|---|---|---|---|
| **Intercept** | **-1.7528** | **-2.1135 – -1.3187** | **-2.0883** | **-2.2421 – -1.9197** |
| **Elevation** | **-0.1760** | **-0.3477 – -0.0061** | **-0.1899** | **-0.3311 – -0.0498** |
| **Random effects** | | | | |
| $\sigma^2$ | 0.0014 | | | |
| $\tau_{00}$ | 0.0041 | | | |
| N | 87 $_{\text{taxa}}$ | | | |
| Observations | 87 | | 87 | |

**21. Bayesian mixed-effects model for: TOL_IGR**

| *Predictors* | *Median* | *CI (95%)* | *Median* | *CI (95%)* |
|---|---|---|---|---|
| Intercept | 0.0119 | -0.3223 – 0.3207 | 0.0131 | -0.3311 – 0.3387 |
| Elevation | 0.0125 | -0.0219 – 0.0474 | 0.0115 | -0.0224 – 0.0446 |
| Treatment (Heat vs Frost) | 0.0024 | -0.0449 – 0.0498 | 0.0031 | -0.0439 – 0.0513 |
| Elevation x Heat treatment vs Frost | 0.0122 | -0.0343 – 0.0607 | 0.0131 | -0.0333 – 0.0624 |
| **Random effects** | | | | |
| $\sigma^2$ | 0.0008 | | -0.0000 | |
| $\tau_{00}$ | 0.0223 | | 0.0228 | |
| N | 80 taxa | | 1 sowing_id | |
| | 1 sowing_id | | | |
| Observations | 150 | | 150 | |

**22. Bayesian mixed-effects model for: TOL_MGR**

| *Predictors* | *Median* | *CI (95%)* | *Median* | *CI (95%)* |
|---|---|---|---|---|
| Intercept | 0.0157 | -0.3524 – 0.3723 | 0.1655 | -0.2196 – 0.4161 |
| Elevation | 0.0827 | -0.1707 – 0.3372 | 0.0358 | -0.1618 – 0.2359 |
| **Treatment (Heat vs Frost)** | **1.4953** | **1.2338 – 1.7401** | **1.3725** | **1.0969 – 1.6343** |
| **Elevation x Heat treatment vs Frost** | **0.3153** | **0.0721 – 0.5639** | **0.3057** | **0.0222 – 0.5661** |
| **Random effects** | | | | |
| $\sigma^2$ | 0.5017 | | 0.0089 | |
| $\tau_{00}$ | 2.0030 | | 2.4691 | |
| N | 80 taxa | | 2 sowing_id | |
| | 2 sowing_id | | | |
| Observations | 300 | | 300 | |

**23. Bayesian mixed-effects model for: TOL_XMID**

| *Predictors* | *Median* | *CI (95%)* | *Median* | *CI (95%)* |
|---|---|---|---|---|
| Intercept | -0.0044 | -0.1870 – 0.1980 | -0.0256 | -0.2053 – 0.1654 |
| Elevation | 0.0080 | -0.0230 – 0.0384 | 0.0036 | -0.0230 – 0.0295 |
| Treatment (Heat vs Frost) | -0.0175 | -0.0536 – 0.0186 | -0.0134 | -0.0508 – 0.0237 |
| **Elevation x Heat treatment vs Frost** | **-0.0475** | **-0.0819 – -0.0126** | **-0.0502** | **-0.0874 – -0.0135** |
| **Random effects** | | | | |
| $\sigma^2$ | 0.0042 | | 0.0001 | |
| $\tau_{00}$ | 0.0254 | | 0.0296 | |
| N | 80 $_{\text{taxa}}$<br>2 $_{\text{sowing\_id}}$ | | 2 $_{\text{sowing\_id}}$ | |
| Observations | 300 | | 300 | |

**24. Bayesian mixed-effects model for: TOL_ASYM**

| *Predictors* | *Median* | *CI (95%)* | *Median* | *CI (95%)* |
|---|---|---|---|---|
| Intercept | -0.0289 | -0.5527 – 0.5166 | -0.0351 | -0.4709 – 0.3984 |
| Elevation | 0.0134 | -0.1312 – 0.1627 | -0.0601 | -0.1776 – 0.0580 |
| **Treatment (Heat vs Frost)** | **0.4121** | **0.2602 – 0.5589** | **0.4036** | **0.2326 – 0.5726** |
| **Elevation x Heat treatment vs Frost** | **-0.3033** | **-0.4485 – -0.1560** | **-0.3018** | **-0.4668 – -0.1374** |
| **Random effects** | | | | |
| $\sigma^2$ | 0.2109 | | 0.0034 | |
| $\tau_{00}$ | 0.5894 | | 0.7897 | |
| N | 80 $_{\text{taxa}}$<br>2 $_{\text{sowing\_id}}$ | | 2 $_{\text{sowing\_id}}$ | |
| Observations | 300 | | 300 | |

**Comparison of models (phylogenetic effect)**

Comparison of models with *vs*. without consideration of phylogenetic relationships between species (i.e., phylogeny as random effect). Indicated in bold is when one model was preferred over the other (expected log-predictive density, ELPD ± SE ≠ 0). Negative values of ELPD indicate that the model including the phylogeny performed better, positive values that the model without the phylogeny performed better. In red are the traits for which there was a significant effect of elevation or its interaction. Comparison is performed via the expected log-predictive density (i.e., ELPD) using eave-one-out cross validation (LOO).

| Trait | Trait ID | ELPD_diff ± SE | | | |
|---|---|---|---|---|---|
| **Seed size** | **SSIZ** | **-** | **81.1** | **±** | **13.1** |
| **Time to germinate** | **TGER** | **-** | **6** | **±** | **3.4** |
| Initial growth rate | IGR | + | 0.6 | ± | 0.8 |
| Maximal growth rate | MGR | + | 0.5 | ± | 0.9 |
| **Time to fastest growth** | **XMID** | **-** | **40** | **±** | **9.2** |
| **Asymptotic size** | **ASYM** | **-** | **279.3** | **±** | **22.1** |
| **Number of leaves** s2 | **NLEA** | **-** | **239.6** | **±** | **24.4** |
| **Leaf area** | **LA** | **-** | **266.1** | **±** | **27.2** |
| **Specific leaf area** | **SLA** | **-** | **66.8** | **±** | **17.5** |
| **Leaf dry matter content** | **LDMC** | **-** | **98** | **±** | **14.1** |
| **Leaf thickness** s2 | **LTh** | **-** | **21.5** | **±** | **6.8** |
| **Leaf dissection index** | **LDI** | **-** | **305.6** | **±** | **21.5** |
| Frost resistance | | | | | |
| *...acclimated (1h at -6 °C)* | RES(-)T1 | + | 0.5 | ± | 1.1 |
| ***...non acclimated (1h at -5 °C)*** s2 | **RES(-)T1** | **-** | **8** | **±** | **4.8** |
| *...acclimated (1h at -11 °C)* | RES(-)T2 | - | 0.4 | ± | 2.2 |
| *...non acclimated, (1h at -10 °C)* s2 | RES(-)T2 | - | 1.8 | ± | 3.3 |
| Heat resistance | | | | | |
| ***...acclimated (1h at +47 °C)*** | **RES(+)T1** | **-** | **20.7** | **±** | **4.7** |
| ***...non acclimated (1h at +45 °C)*** s2 | **RES(+)T1** | **-** | **6.3** | **±** | **3.2** |
| ***...acclimated (1h at +51 °C)*** | **RES(+)T2** | **+** | **6.6** | **±** | **3.3** |
| ***...non acclimated, (1h at +50 °C)*** s2 | **RES(+)T2** | **-** | **7.5** | **±** | **4.1** |
| **Tolerance IGR** | **TOL_IGR** | **+** | **0.9** | **±** | **0.8** |
| **Tolerance MGR** | **TOL_MGR** | **-** | **31.3** | **±** | **7.6** |
| **Tolerance XMID** | **TOL_XMID** | **-** | **7.2** | **±** | **4.5** |
| **Tolerance ASYM** | **TOL_ASYM** | **-** | **27.9** | **±** | **8.8** |

***Next page.*** **Figure 1.** Boxplots showing the distribution of species-mean trait values for which species differed depending on their median elevation (low- vs high-elevation), either across growth treatments or in a particular growth treatment (frost, mild, or heat). Colours inside boxes represent the treatments (blue for frost, greyscale for control and red for heat), while the colour intensity represents median elevation of species occurrence (darker colours for low elevation and lighter colours for high elevation). Colours of the frames of boxes, whiskers and outliers represents the two rounds of sowing (grey S1, black S2). The thick horizontal line is the median, the lower and upper hinges are the 25th and 75th percentiles; whiskers extends from the hinges to the smallest (largest) value at most (no further than) 1.5 * IQR of the hinges, and dots are values beyond that range.

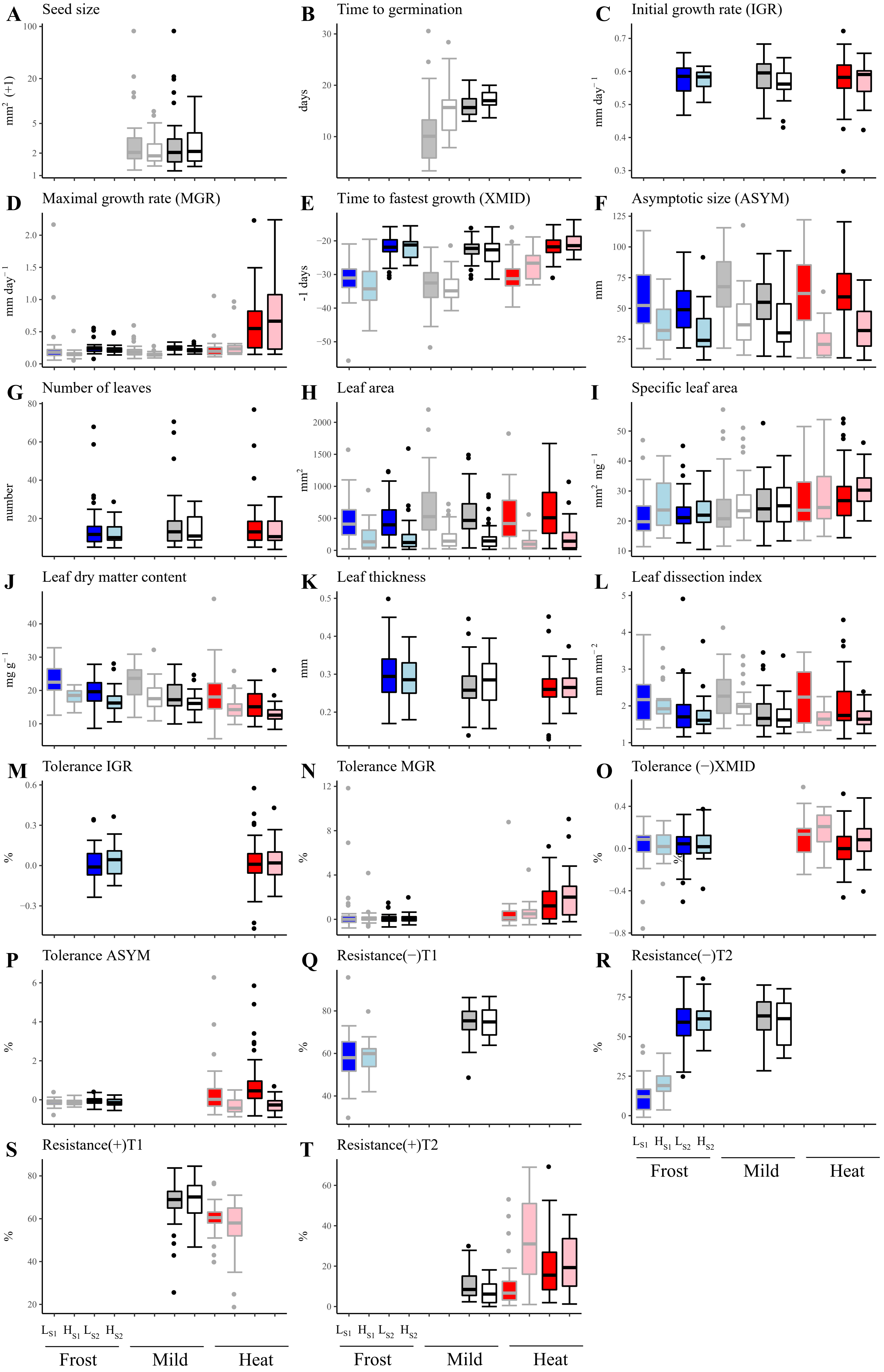
A Seed size
mm² (+1)
B Time to germination
days
C Initial growth rate (IGR)
mm day⁻¹
D Maximal growth rate (MGR)
mm day⁻¹
E Time to fastest growth (XMID)
-1 days
F Asymptotic size (ASYM)
mm
G Number of leaves
number
H Leaf area
mm²
I Specific leaf area
mm² mg⁻¹
J Leaf dry matter content
mg g⁻¹
K Leaf thickness
mm
L Leaf dissection index
mm mm⁻²
M Tolerance IGR
%
N Tolerance MGR
%
O Tolerance (−)XMID
%
P Tolerance ASYM
%
Q Resistance(−)T1
%
R Resistance(−)T2
%
S Resistance(+)T1
%
T Resistance(+)T2
%
LS1 HS1 LS2 HS2
Frost
Mild
Heat
