## Supplementary material for "Trait divergence and trade-offs among Brassicaceae species differing in elevational distribution": A4_M01_PhylogeneticEvolutionaryAnalysis_ForSub.pdf

### A4 - Supplementary material: evolutionary models and simulations

AICc values are based on 100 simulated trees with 2 regimes (high vs. low elevation; with simmMap); mapping was done with the best model (i.e., ARD: all rates different). Evolutionary models were fitted either on the entire dataset ("full phylogeny") or after random removal of one third of the species ("bootstrapped phylogeny"). Random removal of species (whit a maximal removal of one half of the alpine species, to allow the fitting of multiple-regimes models) was performed 100 times, and for each of these data sets, 100 simmMap simulations were performed. Mean ΔAICc value was compared. 'OUM' is highlighted in yellow when it was the best supported model or when ΔAICc < 2. Graphs of ΔAICc value distributions and simulation analyses are shown on the following pages.

#### Frost (full phylogeny)

| Result | model | LogLik | AIC | AICc | ΔAICc | num_parai | root_theta( | theta_high | theta_low | sigma | sigma_higl | sigma_low | alpha | half_life |
|---|---|---|---|---|---|---|---|---|---|---|---|---|---|---|
| IGR | BM1 | 136.9133 | -269.827 | -269.687 | 0 | 2 | 0.576348 | NA | NA | 1.00E-21 | NA | NA | NA | NA |
| IGR | BMM | 136.9927 | -267.985 | -267.703 | 1.98 | 3 | 0.575174 | NA | NA | NA | 6.29E-06 | 8.50E-15 | NA | NA |
| IGR | WN | 136.9133 | -267.827 | -267.544 | 2.14 | 3 | 0.576348 | NA | NA | 2.70E-03 | NA | NA | NA | NA |
| IGR | OU1 | 136.9133 | -267.827 | -267.544 | 2.14 | 3 | 0.576348 | NA | NA | 1.04E-16 | NA | NA | 0.092943 | 7.457772 |
| IGR | OUM | 137.9211 | -267.842 | -267.366 | 2.32 | 4 | 0.583925 | 0.58975 | 0.568263 | 1.33E-13 | NA | NA | 0.067116 | 10.33138 |

| Result | model | LogLik | AIC | AICc | ΔAICc | num_parai | root_theta( | theta_high | theta_low | sigma | sigma_higl | sigma_low | alpha | half_life |
|---|---|---|---|---|---|---|---|---|---|---|---|---|---|---|
| MGR | OU1 | 86.39636 | -166.793 | -166.52 | 0 | 3 | 0.211569 | NA | NA | 1.76578 | NA | NA | 263.0424 | 0.002635 |
| MGR | OUM | 86.44107 | -164.882 | -164.422 | 2.1 | 4 | 0.215226 | 0.215295 | 0.209507 | 2.790793 | NA | NA | 416.3716 | 0.001665 |
| MGR | BMM | 78.25206 | -150.504 | -150.231 | 16.29 | 3 | 0.210154 | NA | NA | NA | 0.001345 | 5.78E-05 | NA | NA |
| MGR | BM1 | 64.2299 | -124.46 | -124.325 | 42.2 | 2 | 0.213372 | NA | NA | 0.000454 | NA | NA | NA | NA |
| MGR | WN | 46.91431 | -87.8286 | -87.5559 | 78.96 | 3 | 0.247124 | NA | NA | 0.021115 | NA | NA | NA | NA |

| Result | model | LogLik | AIC | AICc | ΔAICc | num_parai | root_theta( | theta_high | theta_low | sigma | sigma_higl | sigma_low | alpha | half_life |
|---|---|---|---|---|---|---|---|---|---|---|---|---|---|---|
| XMID | WN | -265.105 | 536.21 | 536.4859 | 0 | 3 | 25.9942 | NA | NA | 19.85799 | NA | NA | NA | NA |
| XMID | OU1 | -280.292 | 566.5836 | 566.8595 | 30.37 | 3 | 25.14033 | NA | NA | 46.53219 | NA | NA | 3.357091 | 0.206473 |
| XMID | OUM | -279.383 | 566.7663 | 567.2314 | 30.75 | 4 | 24.18382 | 24.03991 | 25.76687 | 31.77518 | NA | NA | 2.759281 | 0.25128 |
| XMID | BM1 | -283.609 | 571.2178 | 571.3542 | 34.87 | 2 | 24.8974 | NA | NA | 0.411843 | NA | NA | NA | NA |
| XMID | BMM | -283.415 | 572.8299 | 573.1057 | 36.62 | 3 | 24.77125 | NA | NA | NA | 0.263968 | 0.463529 | NA | NA |

| Result | model | LogLik | AIC | AICc | ΔAICc | num_parai | root_theta( | theta_high | theta_low | sigma | sigma_higl | sigma_low | alpha | half_life |
|---|---|---|---|---|---|---|---|---|---|---|---|---|---|---|
| ASYM | OUM | -387.674 | 783.3481 | 783.8132 | 0 | 4 | 39.1842 | 30.44055 | 57.14994 | 35.67186 | NA | NA | 0.081748 | 8.479141 |
| ASYM | BM1 | -390.778 | 785.5556 | 785.692 | 1.88 | 2 | 47.264 | NA | NA | 17.52332 | NA | NA | NA | NA |
| ASYM | BMM | -390.66 | 787.32 | 787.5958 | 3.78 | 3 | 46.95896 | NA | NA | NA | 15.19637 | 19.25918 | NA | NA |
| ASYM | OU1 | -390.677 | 787.3549 | 787.6308 | 3.82 | 3 | 46.82432 | NA | NA | 29.39115 | NA | NA | 0.04472 | 15.49974 |
| ASYM | WN | -408.932 | 823.8643 | 824.1402 | 40.33 | 3 | 46.3118 | NA | NA | 468.5501 | NA | NA | NA | NA |

| Result | model | LogLik | AIC | AICc | ΔAICc | num_parai | root_theta( | theta_high | theta_low | sigma | sigma_higl | sigma_low | alpha | half_life |
|---|---|---|---|---|---|---|---|---|---|---|---|---|---|---|
| NLEA | BM1 | -319.848 | 643.6967 | 643.8413 | 0 | 2 | 13.21461 | NA | NA | 0.472307 | NA | NA | NA | NA |
| NLEA | BMM | -319.531 | 645.0622 | 645.3549 | 1.51 | 3 | 13.21589 | NA | NA | NA | 0.631794 | 0.168338 | NA | NA |
| NLEA | OU1 | -319.571 | 645.1418 | 645.4345 | 1.59 | 3 | 13.41738 | NA | NA | 2.850573 | NA | NA | 0.104984 | 6.602385 |
| NLEA | WN | -320.173 | 646.3452 | 646.6379 | 2.8 | 3 | 13.459 | NA | NA | 100.2831 | NA | NA | NA | NA |
| NLEA | OUM | -319.711 | 647.4213 | 647.9151 | 4.07 | 4 | 12.25009 | 12.04814 | 14.22028 | 2.999854 | NA | NA | 1.509313 | 0.459249 |

| Result | model | LogLik | AIC | AICc | ΔAICc | num_parai | root_theta( | theta_high | theta_low | sigma | sigma_higl | sigma_low | alpha | half_life |
|---|---|---|---|---|---|---|---|---|---|---|---|---|---|---|
| LA | BMM | -630.618 | 1267.237 | 1267.516 | 0 | 3 | 345.9843 | NA | NA | NA | 836.438 | 5382.327 | NA | NA |
| LA | OUM | -630.926 | 1269.851 | 1270.321 | 2.8 | 4 | 269.9758 | 169.3345 | 495.4844 | 7533.059 | NA | NA | 0.089755 | 7.722677 |
| LA | BM1 | -633.914 | 1271.828 | 1271.966 | 4.45 | 2 | 391.5096 | NA | NA | 3595.044 | NA | NA | NA | NA |
| LA | OU1 | -633.728 | 1273.457 | 1273.736 | 6.22 | 3 | 384.2648 | NA | NA | 6311.318 | NA | NA | 0.05322 | 13.0242 |
| LA | WN | -651.802 | 1309.605 | 1309.884 | 42.37 | 3 | 396.1312 | NA | NA | 114304.3 | NA | NA | NA | NA |

| Result | model | LogLik | AIC | AICc | ΔAICc | num_parai | root_theta( | theta_high | theta_low | sigma | sigma_higl | sigma_low | alpha | half_life |
|---|---|---|---|---|---|---|---|---|---|---|---|---|---|---|
| SLA | BMM | -290.494 | 586.9872 | 587.2662 | 0 | 3 | 20.45231 | NA | NA | NA | 2.394457 | 0.237694 | NA | NA |
| SLA | BM1 | -294.371 | 592.7428 | 592.8807 | 5.61 | 2 | 20.73033 | NA | NA | 1.097213 | NA | NA | NA | NA |
| SLA | OU1 | -293.686 | 593.3719 | 593.651 | 6.38 | 3 | 21.0041 | NA | NA | 2.647908 | NA | NA | 0.07989 | 8.676306 |
| SLA | OUM | -293.562 | 595.1237 | 595.5943 | 8.33 | 4 | 20.96078 | 21.54552 | 20.70129 | 2.656552 | NA | NA | 0.082308 | 8.421357 |
| SLA | WN | -307.153 | 620.3065 | 620.5856 | 33.32 | 3 | 22.66615 | NA | NA | 53.93346 | NA | NA | NA | NA |

| Result | model | LogLik | AIC | AICc | ΔAICc | num_parai | root_theta( | theta_high | theta_low | sigma | sigma_higl | sigma_low | alpha | half_life |
|---|---|---|---|---|---|---|---|---|---|---|---|---|---|---|
| LDMC | OUM | -248.867 | 505.7331 | 506.2093 | 0 | 4 | 17.10058 | 16.82926 | 20.64002 | 8.461489 | NA | NA | 0.561422 | 1.234772 |
| LDMC | OU1 | -255.082 | 516.1631 | 516.4454 | 10.24 | 3 | 19.42341 | NA | NA | 3.353703 | NA | NA | 0.148259 | 4.675233 |
| LDMC | WN | -257.294 | 520.588 | 520.8703 | 14.66 | 3 | 19.37002 | NA | NA | 18.99185 | NA | NA | NA | NA |
| LDMC | BMM | -258.772 | 523.5442 | 523.8265 | 17.62 | 3 | 19.01253 | NA | NA | NA | 0.454242 | 1.423746 | NA | NA |
| LDMC | BM1 | -260.201 | 524.402 | 524.5415 | 18.33 | 2 | 19.48248 | NA | NA | 1.046212 | NA | NA | NA | NA |

| Result | model | LogLik | AIC | AICc | ΔAICc | num_parai | root_theta( | theta_high | theta_low | sigma | sigma_higl | sigma_low | alpha | half_life |
|---|---|---|---|---|---|---|---|---|---|---|---|---|---|---|
| LTH | BM1 | 109.3944 | -214.789 | -214.633 | 0 | 2 | 0.294246 | NA | NA | 3.14E-05 | NA | NA | NA | NA |
| LTH | BMM | 109.5762 | -213.152 | -212.837 | 1.8 | 3 | 0.293711 | NA | NA | NA | 5.19E-05 | 1.61E-05 | NA | NA |
| LTH | OU1 | 108.9659 | -211.932 | -211.616 | 3.02 | 3 | 0.295221 | NA | NA | 7.69E-05 | NA | NA | 0.058095 | 11.93118 |
| LTH | OUM | 109.1373 | -210.275 | -209.741 | 4.89 | 4 | 0.292959 | 0.304362 | 0.290665 | 7.32E-05 | NA | NA | 0.060326 | 11.48995 |
| LTH | WN | 107.9976 | -209.995 | -209.679 | 4.95 | 3 | 0.295544 | NA | NA | 3.94E-03 | NA | NA | NA | NA |

| Result | model | LogLik | AIC | AICc | ΔAICc | num_parai | root_theta( | theta_high | theta_low | sigma | sigma_higl | sigma_low | alpha | half_life |
|---|---|---|---|---|---|---|---|---|---|---|---|---|---|---|
| LDI | BMM | -61.5391 | 129.0782 | 129.3573 | 0 | 3 | 1.944179 | NA | NA | NA | 0.003536 | 0.013709 | NA | NA |
| LDI | BM1 | -63.4802 | 130.9605 | 131.0984 | 1.74 | 2 | 1.984863 | NA | NA | 0.009776 | NA | NA | NA | NA |
| LDI | OU1 | -63.8981 | 133.7962 | 134.0753 | 4.72 | 3 | 1.941361 | NA | NA | 0.015857 | NA | NA | 0.039924 | 17.3617 |
| LDI | OUM | -63.505 | 135.01 | 135.4806 | 6.12 | 4 | 1.926488 | 1.760417 | 2.023022 | 0.016819 | NA | NA | 0.047699 | 14.53174 |
| LDI | WN | -79.3083 | 164.6167 | 164.8957 | 35.54 | 3 | 1.978486 | NA | NA | 0.341137 | NA | NA | NA | NA |

| Result | model | LogLik | AIC | AICc | ΔAICc | num_parai | root_theta( | theta_high | theta_low | sigma | sigma_higl | sigma_low | alpha | half_life |
|---|---|---|---|---|---|---|---|---|---|---|---|---|---|---|
| RESmT1 | BM1 | -241.466 | 486.9327 | 487.1361 | 0 | 2 | 58.34153 | NA | NA | 2.26E-23 | NA | NA | NA | NA |
| RESmT1 | WN | -241.462 | 488.9246 | 489.3384 | 2.2 | 3 | 58.34153 | NA | NA | 1.41E+02 | NA | NA | NA | NA |
| RESmT1 | BMM | -241.466 | 488.9327 | 489.3465 | 2.21 | 3 | 58.34153 | NA | NA | NA | 9.56E-13 | 4.17E-11 | NA | NA |
| RESmT1 | OU1 | -241.466 | 488.9327 | 489.3465 | 2.21 | 3 | 58.34153 | NA | NA | 8.20E-12 | NA | NA | 16.67878 | 0.041559 |
| RESmT1 | OUM | -241.07 | 490.1394 | 490.8411 | 3.7 | 4 | 56.76667 | 56.29585 | 59.37356 | 3.46E-08 | NA | NA | 1.029879 | 0.673253 |

| Result | model | LogLik | AIC | AICc | ΔAICc | num_parai | root_theta( | theta_high | theta_low | sigma | sigma_higl | sigma_low | alpha | half_life |
|---|---|---|---|---|---|---|---|---|---|---|---|---|---|---|
| RESmT2 | WN | -393.078 | 792.1565 | 792.4324 | 0 | 3 | 43.15285 | NA | NA | 330.698 | NA | NA | NA | NA |
| RESmT2 | BM1 | -415.688 | 835.3759 | 835.5123 | 43.08 | 2 | 44.06042 | NA | NA | 2.036066 | NA | NA | NA | NA |
| RESmT2 | OU1 | -415.035 | 836.0699 | 836.3458 | 43.91 | 3 | 44.72562 | NA | NA | 31.12075 | NA | NA | 0.168389 | 4.116335 |
| RESmT2 | OUM | -413.973 | 835.9451 | 836.4102 | 43.98 | 4 | 49.33254 | 50.98705 | 41.58934 | 40.66669 | NA | NA | 0.286043 | 2.425961 |
| RESmT2 | BMM | -415.297 | 836.5929 | 836.8688 | 44.44 | 3 | 45.09444 | NA | NA | NA | 1.46E-11 | 6.27949 | NA | NA |

| Result | model | LogLik | AIC | AICc | ΔAICc | num_parai | root_theta( | theta_high | theta_low | sigma | sigma_higl | sigma_low | alpha | half_life |
|---|---|---|---|---|---|---|---|---|---|---|---|---|---|---|
| TOL_IGR | BM1 | 50.50212 | -97.0042 | -96.8352 | 0 | 2 | 0.020221 | NA | NA | 6.86E-25 | NA | NA | NA | NA |
| TOL_IGR | WN | 50.50329 | -95.0066 | -94.6637 | 2.17 | 3 | 0.020221 | NA | NA | 1.50E-02 | NA | NA | NA | NA |
| TOL_IGR | BMM | 50.50212 | -95.0042 | -94.6614 | 2.17 | 3 | 0.020221 | NA | NA | NA | 4.61E-14 | 2.47E-15 | NA | NA |
| TOL_IGR | OU1 | 50.50212 | -95.0042 | -94.6614 | 2.17 | 3 | 0.020221 | NA | NA | 3.51E-24 | NA | NA | 0.09603 | 7.217997 |
| TOL_IGR | OUM | 51.46369 | -94.9274 | -94.3477 | 2.49 | 4 | 0.041059 | 0.052486 | 0.001254 | 3.32E-14 | NA | NA | 0.043834 | 15.83698 |

| Result | model | LogLik | AIC | AICc | ΔAICc | num_param | root_theta | theta_high | theta_low | sigma | sigma_high | sigma_low | alpha | half_life |
|---|---|---|---|---|---|---|---|---|---|---|---|---|---|---|
| TOL_MGR | OU1 | -39.8947 | 85.78939 | 86.13225 | 0 | 3 | 0.103091 | NA | NA | 2.14791 | NA | NA | 6.238566 | 0.111107 |
| TOL_MGR | WN | -39.9234 | 85.84679 | 86.18965 | 0.06 | 3 | 0.102006 | NA | NA | 0.17224 | NA | NA | NA | NA |
| TOL_MGR | OUM | -39.7992 | 87.59829 | 88.17801 | 2.05 | 4 | 0.105233 | 0.135642 | 0.084846 | 3.251017 | NA | NA | 9.476518 | 0.07316 |
| TOL_MGR | BMM | -94.3368 | 194.6737 | 195.0166 | 108.88 | 3 | 0.045423 | NA | NA | NA | 0.048732 | 0.162991 | NA | NA |
| TOL_MGR | BM1 | -98.3356 | 200.6712 | 200.8403 | 114.71 | 2 | 0.045983 | NA | NA | 0.122595 | NA | NA | NA | NA |

| Result | model | LogLik | AIC | AICc | ΔAICc | num_param | root_theta | theta_high | theta_low | sigma | sigma_high | sigma_low | alpha | half_life |
|---|---|---|---|---|---|---|---|---|---|---|---|---|---|---|
| TOL_XMID | WN | 33.35682 | -60.7136 | -60.3708 | 0 | 3 | -0.02616 | NA | NA | 0.023768 | NA | NA | NA | NA |
| TOL_XMID | OU1 | 33.35678 | -60.7136 | -60.3707 | 0 | 3 | -0.02616 | NA | NA | 1.33854 | NA | NA | 28.15952 | 0.024615 |
| TOL_XMID | OUM | 33.5052 | -59.0104 | -58.4307 | 1.94 | 4 | -0.03981 | -0.03981 | -0.01919 | 1.789862 | NA | NA | 37.80411 | 0.01834 |
| TOL_XMID | BM1 | -16.8106 | 37.62112 | 37.79014 | 98.16 | 2 | -0.02733 | NA | NA | 0.013538 | NA | NA | NA | NA |
| TOL_XMID | BMM | -16.6707 | 39.34148 | 39.68433 | 100.06 | 3 | -0.02715 | NA | NA | NA | 0.014916 | 0.012705 | NA | NA |

| Result | model | LogLik | AIC | AICc | ΔAICc | num_param | root_theta | theta_high | theta_low | sigma | sigma_high | sigma_low | alpha | half_life |
|---|---|---|---|---|---|---|---|---|---|---|---|---|---|---|
| TOL_ASYM | OUM | 11.85982 | -15.7196 | -15.1399 | 0 | 4 | -0.13586 | -0.13586 | -0.05452 | 3.365332 | NA | NA | 39.59847 | 0.017504 |
| TOL_ASYM | WN | 10.59275 | -15.1855 | -14.8427 | 0.3 | 3 | -0.082 | NA | NA | 0.043974 | NA | NA | NA | NA |
| TOL_ASYM | OU1 | 10.59272 | -15.1854 | -14.8426 | 0.3 | 3 | -0.082 | NA | NA | 2.748059 | NA | NA | 31.24768 | 0.022182 |
| TOL_ASYM | BM1 | -53.3544 | 110.7088 | 110.8779 | 126.02 | 2 | -0.06396 | NA | NA | 0.036349 | NA | NA | NA | NA |
| TOL_ASYM | BMM | -53.2101 | 112.4202 | 112.7631 | 127.9 | 3 | -0.06439 | NA | NA | NA | 0.032822 | 0.038413 | NA | NA |

#### Mild (control, full phylogeny)

| Result | model | LogLik | AIC | AICc | ΔAICc | num_param | root_theta | theta_high | theta_low | sigma | sigma_high | sigma_low | alpha | half_life |
|---|---|---|---|---|---|---|---|---|---|---|---|---|---|---|
| SSIZ | BMM | -20.4678 | 46.93551 | 47.19925 | 0 | 3 | 0.669946 | NA | NA | NA | 0.024395 | 0.009018 | NA | NA |
| SSIZ | BM1 | -25.1854 | 54.37081 | 54.50124 | 7.3 | 2 | 0.656344 | NA | NA | 0.014617 | NA | NA | NA | NA |
| SSIZ | OU1 | -25.1173 | 56.23465 | 56.49839 | 9.3 | 3 | 0.629671 | NA | NA | 0.018905 | NA | NA | 0.031041 | 22.3302 |
| SSIZ | OUM | -24.991 | 57.98191 | 58.42635 | 11.23 | 4 | 0.638447 | 0.548111 | 0.675005 | 0.018903 | NA | NA | 0.031322 | 22.1296 |
| SSIZ | WN | -55.9642 | 117.9284 | 118.1922 | 70.99 | 3 | 0.457714 | NA | NA | 0.190199 | NA | NA | NA | NA |

| Result | model | LogLik | AIC | AICc | ΔAICc | num_param | root_theta | theta_high | theta_low | sigma | sigma_high | sigma_low | alpha | half_life |
|---|---|---|---|---|---|---|---|---|---|---|---|---|---|---|
| TGER | BM1 | -251.035 | 506.0709 | 506.2057 | 0 | 2 | 16.81124 | NA | NA | 0.142634 | NA | NA | NA | NA |
| TGER | BMM | -250.065 | 506.1295 | 506.4022 | 0.2 | 3 | 16.81227 | NA | NA | NA | 3.39E-11 | 0.245739 | NA | NA |
| TGER | OU1 | -252.244 | 510.4873 | 510.76 | 4.55 | 3 | 16.64445 | NA | NA | 0.244388 | NA | NA | 0.029812 | 23.25063 |
| TGER | OUM | -252.111 | 512.2212 | 512.6809 | 6.48 | 4 | 16.73431 | 17.12934 | 15.78375 | 0.276189 | NA | NA | 0.045587 | 15.20533 |
| TGER | WN | -256.651 | 519.3013 | 519.5741 | 13.37 | 3 | 15.01955 | NA | NA | 15.51 | NA | NA | NA | NA |

| Result | model | LogLik | AIC | AICc | ΔAICc | num_param | root_theta | theta_high | theta_low | sigma | sigma_high | sigma_low | alpha | half_life |
|---|---|---|---|---|---|---|---|---|---|---|---|---|---|---|
| IGR | BM1 | 140.0523 | -276.105 | -275.964 | 0 | 2 | 0.581831 | NA | NA | 6.40E-06 | NA | NA | NA | NA |
| IGR | BMM | 140.2149 | -274.43 | -274.144 | 1.82 | 3 | 0.582011 | NA | NA | NA | 1.45E-05 | 4.51E-07 | NA | NA |
| IGR | OUM | 141.1832 | -274.366 | -273.884 | 2.08 | 4 | 0.579382 | 0.568177 | 0.58744 | 1.44E-10 | NA | NA | 0.092125 | 7.523963 |
| IGR | OU1 | 140.0205 | -274.041 | -273.755 | 2.21 | 3 | 0.580798 | NA | NA | 2.34E-05 | NA | NA | 0.092712 | 7.47635 |
| IGR | WN | 139.9259 | -273.852 | -273.566 | 2.4 | 3 | 0.580302 | NA | NA | 2.43E-03 | NA | NA | NA | NA |

| Result | model | LogLik | AIC | AICc | ΔAICc | num_param | root_theta | theta_high | theta_low | sigma | sigma_high | sigma_low | alpha | half_life |
|---|---|---|---|---|---|---|---|---|---|---|---|---|---|---|
| MGR | OUM | 145.3458 | -282.692 | -282.232 | 0 | 4 | 0.184456 | 0.184103 | 0.209015 | 1.59E-02 | NA | NA | 42.84511 | 0.016178 |
| MGR | BM1 | 141.124 | -278.248 | -278.113 | 4.12 | 2 | 0.200128 | NA | NA | 6.63E-06 | NA | NA | NA | NA |
| MGR | OU1 | 141.4922 | -276.985 | -276.712 | 5.52 | 3 | 0.201061 | NA | NA | 8.49E-05 | NA | NA | 0.237975 | 2.912688 |
| MGR | BMM | 141.3953 | -276.791 | -276.518 | 5.71 | 3 | 0.199597 | NA | NA | NA | 1.82E-08 | 1.04E-05 | NA | NA |
| MGR | WN | 129.0734 | -252.147 | -251.874 | 30.36 | 3 | 0.220296 | NA | NA | 3.54E-03 | NA | NA | NA | NA |

| Result | model | LogLik | AIC | AICc | ΔAICc | num_param | root_theta | theta_high | theta_low | sigma | sigma_high | sigma_low | alpha | half_life |
|---|---|---|---|---|---|---|---|---|---|---|---|---|---|---|
| XMID | WN | -259.685 | 525.369 | 525.6418 | 0 | 3 | 27.42776 | NA | NA | 16.56743 | NA | NA | NA | NA |
| XMID | BM1 | -286.127 | 576.2534 | 576.3883 | 50.75 | 2 | 25.6311 | NA | NA | 0.361368 | NA | NA | NA | NA |
| XMID | OU1 | -285.189 | 576.3783 | 576.651 | 51.01 | 3 | 25.93681 | NA | NA | 1.603342 | NA | NA | 0.122333 | 5.666056 |
| XMID | BMM | -285.624 | 577.248 | 577.5208 | 51.88 | 3 | 25.69992 | NA | NA | NA | 0.08954 | 0.473444 | NA | NA |
| XMID | OUM | -285.134 | 578.2687 | 578.7285 | 53.09 | 4 | 25.93366 | 26.27692 | 25.73551 | 1.571034 | NA | NA | 0.121887 | 5.686825 |

| Result | model | LogLik | AIC | AICc | ΔAICc | num_param | root_theta | theta_high | theta_low | sigma | sigma_high | sigma_low | alpha | half_life |
|---|---|---|---|---|---|---|---|---|---|---|---|---|---|---|
| ASYM | BM1 | -397.657 | 799.3138 | 799.4486 | 0 | 2 | 53.21985 | NA | NA | 20.87169 | NA | NA | NA | NA |
| ASYM | OUM | -396.414 | 800.8278 | 801.2876 | 1.84 | 4 | 47.47171 | 38.72143 | 64.84732 | 33.14599 | NA | NA | 0.047669 | 14.54077 |
| ASYM | BMM | -397.568 | 801.1354 | 801.4081 | 1.96 | 3 | 52.91117 | NA | NA | NA | 18.37153 | 22.54203 | NA | NA |
| ASYM | OU1 | -398.05 | 802.0991 | 802.3718 | 2.92 | 3 | 53.22831 | NA | NA | 31.10405 | NA | NA | 0.035327 | 19.62092 |
| ASYM | WN | -424.724 | 855.4474 | 855.7202 | 56.27 | 3 | 53.78076 | NA | NA | 598.9915 | NA | NA | NA | NA |

| Result | model | LogLik | AIC | AICc | ΔAICc | num_param | root_theta | theta_high | theta_low | sigma | sigma_high | sigma_low | alpha | half_life |
|---|---|---|---|---|---|---|---|---|---|---|---|---|---|---|
| NLEA | BM1 | -335.478 | 674.9556 | 675.0985 | 0 | 2 | 14.76192 | NA | NA | 1.47E+00 | NA | NA | NA | NA |
| NLEA | BMM | -335.042 | 676.0844 | 676.3735 | 1.28 | 3 | 14.77484 | NA | NA | NA | 0.477514 | 2.216113 | NA | NA |
| NLEA | OU1 | -335.651 | 677.3025 | 677.5916 | 2.49 | 3 | 15.0625 | NA | NA | 4.26E+00 | NA | NA | 0.079584 | 8.709611 |
| NLEA | WN | -336.672 | 679.3449 | 679.634 | 4.54 | 3 | 14.97174 | NA | NA | 1.35E+02 | NA | NA | NA | NA |
| NLEA | OUM | -336.021 | 680.0418 | 680.5296 | 5.43 | 4 | 13.04028 | 12.98981 | 16.00744 | 2.17E-04 | NA | NA | 5.1666 | 0.134548 |

| Result | model | LogLik | AIC | AICc | ΔAICc | num_param | root_theta | theta_high | theta_low | sigma | sigma_high | sigma_low | alpha | half_life |
|---|---|---|---|---|---|---|---|---|---|---|---|---|---|---|
| LA | BMM | -637.28 | 1280.56 | 1280.836 | 0 | 3 | 388.7511 | NA | NA | NA | 1698.266 | 6984.076 | NA | NA |
| LA | OUM | -638.106 | 1284.212 | 1284.677 | 3.84 | 4 | 311.217 | 180.8324 | 628.2547 | 7368.694 | NA | NA | 0.055139 | 12.57102 |
| LA | BM1 | -640.464 | 1284.928 | 1285.065 | 4.23 | 2 | 410.9441 | NA | NA | 4490.969 | NA | NA | NA | NA |
| LA | OU1 | -640.779 | 1287.558 | 1287.834 | 7 | 3 | 419.0167 | NA | NA | 6747.16 | NA | NA | 0.03846 | 18.02263 |
| LA | WN | -670.276 | 1346.552 | 1346.827 | 65.99 | 3 | 468.5414 | NA | NA | 146306.5 | NA | NA | NA | NA |

| Result | model | LogLik | AIC | AICc | ΔAICc | num_param | root_theta | theta_high | theta_low | sigma | sigma_high | sigma_low | alpha | half_life |
|---|---|---|---|---|---|---|---|---|---|---|---|---|---|---|
| SLA | OU1 | -313.379 | 632.7576 | 633.0335 | 0 | 3 | 23.71782 | NA | NA | 7.854961 | NA | NA | 0.123995 | 5.590134 |
| SLA | OUM | -312.959 | 633.917 | 634.3821 | 1.35 | 4 | 23.8729 | 25.44093 | 22.83967 | 7.947909 | NA | NA | 0.130964 | 5.292672 |
| SLA | BM1 | -316.801 | 637.6022 | 637.7385 | 4.71 | 2 | 23.6234 | NA | NA | 2.518904 | NA | NA | NA | NA |
| SLA | BMM | -316.71 | 639.4204 | 639.6963 | 6.66 | 3 | 23.57882 | NA | NA | NA | 2.859758 | 2.326576 | NA | NA |
| SLA | WN | -321.09 | 648.1791 | 648.455 | 15.42 | 3 | 25.84099 | NA | NA | 67.96794 | NA | NA | NA | NA |

| Result | model | LogLik | AIC | AICc | ΔAICc | num_param | root_theta | theta_high | theta_low | sigma | sigma_high | sigma_low | alpha | half_life |
|---|---|---|---|---|---|---|---|---|---|---|---|---|---|---|
| LDMC | WN | -266.061 | 538.1217 | 538.4008 | 0 | 3 | 19.06962 | NA | NA | 21.64093 | NA | NA | NA | NA |
| LDMC | OUM | -266.431 | 540.8614 | 541.332 | 2.93 | 4 | 16.52525 | 16.00058 | 19.90072 | 8.629902 | NA | NA | 0.316647 | 2.189046 |
| LDMC | OU1 | -269.897 | 545.7931 | 546.0722 | 7.67 | 3 | 18.67183 | NA | NA | 6.354504 | NA | NA | 0.196708 | 3.523733 |
| LDMC | BMM | -272.437 | 550.8741 | 551.1531 | 12.75 | 3 | 18.09034 | NA | NA | NA | 0.326468 | 2.054706 | NA | NA |
| LDMC | BM1 | -275.737 | 555.4749 | 555.6128 | 17.21 | 2 | 18.69359 | NA | NA | 1.356884 | NA | NA | NA | NA |

| Result | model | LogLik | AIC | AICc | ΔAICc | num_param | root_theta | theta_high | theta_low | sigma | sigma_high | sigma_low | alpha | half_life |
|---|---|---|---|---|---|---|---|---|---|---|---|---|---|---|
| LTH | BM1 | 110.4539 | -216.908 | -216.752 | 0 | 2 | 0.277357 | NA | NA | 3.56E-05 | NA | NA | NA | NA |
| LTH | BMM | 111.4953 | -216.991 | -216.675 | 0.08 | 3 | 0.273853 | NA | NA | NA | 9.42E-05 | 5.50E-12 | NA | NA |
| LTH | OU1 | 110.4862 | -214.973 | -214.657 | 2.1 | 3 | 0.275823 | NA | NA | 1.27E-04 | NA | NA | 0.084283 | 8.224015 |

| | | | | | | | | | | | | | | |
|---|---|---|---|---|---|---|---|---|---|---|---|---|---|---|
| LTH | OUM | 111.5845 | -215.169 | -214.636 | 2.12 | 4 | 0.287276 | 0.298834 | 0.262601 | 1.46E-04 | NA | NA | 0.106951 | 6.4811 |
| LTH | WN | 109.2522 | -212.504 | -212.189 | 4.56 | 3 | 0.2743 | NA | NA | 3.81E-03 | NA | NA | NA | NA |

| Result | model | LogLik | AIC | AICc | ΔAICc | num_para | root_theta( | theta_high | theta_low | sigma | sigma_hig | sigma_low | alpha | half_life |
|---|---|---|---|---|---|---|---|---|---|---|---|---|---|---|
| LDI | BMM | -56.8881 | 119.7762 | 120.0521 | 0 | 3 | 2.007273 | NA | NA | NA | 0.004889 | 0.013685 | NA | NA |
| LDI | BM1 | -58.248 | 120.496 | 120.6324 | 0.58 | 2 | 2.021529 | NA | NA | 0.01047 | NA | NA | NA | NA |
| LDI | OU1 | -58.1999 | 122.3998 | 122.6756 | 2.62 | 3 | 1.982371 | NA | NA | 0.016602 | NA | NA | 0.039292 | 17.64072 |
| LDI | OUM | -57.9923 | 123.9847 | 124.4498 | 4.4 | 4 | 1.988012 | 1.879668 | 2.01889 | 0.017116 | NA | NA | 0.042695 | 16.23578 |
| LDI | WN | -74.9858 | 155.9715 | 156.2474 | 36.2 | 3 | 2.014365 | NA | NA | 0.30427 | NA | NA | NA | NA |

| Result | model | LogLik | AIC | AICc | ΔAICc | num_para | root_theta( | theta_high | theta_low | sigma | sigma_hig | sigma_low | alpha | half_life |
|---|---|---|---|---|---|---|---|---|---|---|---|---|---|---|
| RESmT1 | BM1 | -295.396 | 594.792 | 594.9349 | 0 | 2 | 74.98387 | NA | NA | 0.519498 | NA | NA | NA | NA |
| RESmT1 | BMM | -294.799 | 595.5979 | 595.8871 | 0.95 | 3 | 74.83885 | NA | NA | NA | 1.137984 | 0.114445 | NA | NA |
| RESmT1 | OU1 | -295.573 | 597.1464 | 597.4355 | 2.5 | 3 | 74.89086 | NA | NA | 1.582793 | NA | NA | 0.06646 | 10.42957 |
| RESmT1 | OUM | -295.197 | 598.3933 | 598.8811 | 3.95 | 4 | 75.21219 | 76.62251 | 73.80983 | 1.639094 | NA | NA | 0.071014 | 9.760778 |
| RESmT1 | WN | -297.733 | 601.4668 | 601.7559 | 6.82 | 3 | 74.6774 | NA | NA | 54.95796 | NA | NA | NA | NA |

| Result | model | LogLik | AIC | AICc | ΔAICc | num_para | root_theta( | theta_high | theta_low | sigma | sigma_hig | sigma_low | alpha | half_life |
|---|---|---|---|---|---|---|---|---|---|---|---|---|---|---|
| RESmT2 | BM1 | -346.472 | 696.9435 | 697.0864 | 0 | 2 | 61.62727 | NA | NA | 1.75E-21 | NA | NA | NA | NA |
| RESmT2 | OU1 | -346.331 | 698.6614 | 698.9506 | 1.86 | 3 | 61.54437 | NA | NA | 3.49E+00 | NA | NA | 0.150054 | 4.619316 |
| RESmT2 | BMM | -346.418 | 698.8367 | 699.1259 | 2.04 | 3 | 61.62727 | NA | NA | NA | 0.657367 | 6.39E-12 | NA | NA |
| RESmT2 | WN | -346.471 | 698.9423 | 699.2315 | 2.15 | 3 | 61.62727 | NA | NA | 1.69E+02 | NA | NA | NA | NA |
| RESmT2 | OUM | -346.177 | 700.3538 | 700.8416 | 3.76 | 4 | 61.51365 | 60.19197 | 62.30646 | 2.65E+00 | NA | NA | 0.14756 | 4.697383 |

| Result | model | LogLik | AIC | AICc | ΔAICc | num_para | root_theta( | theta_high | theta_low | sigma | sigma_hig | sigma_low | alpha | half_life |
|---|---|---|---|---|---|---|---|---|---|---|---|---|---|---|
| RESpT1 | BMM | -323.669 | 653.3373 | 653.6264 | 0 | 3 | 67.27832 | NA | NA | NA | 2.446103 | 1.07E-11 | NA | NA |
| RESpT1 | BM1 | -324.906 | 653.8121 | 653.955 | 0.33 | 2 | 67.03622 | NA | NA | 0.788725 | NA | NA | NA | NA |
| RESpT1 | OU1 | -324.72 | 655.4395 | 655.7286 | 2.1 | 3 | 67.1015 | NA | NA | 3.239108 | NA | NA | 0.096021 | 7.218669 |
| RESpT1 | OUM | -324.635 | 657.2695 | 657.7573 | 4.13 | 4 | 67.05412 | 66.34745 | 67.51739 | 3.124135 | NA | NA | 0.094829 | 7.309437 |
| RESpT1 | WN | -325.865 | 657.7309 | 658.0201 | 4.39 | 3 | 67.10914 | NA | NA | 104.9289 | NA | NA | NA | NA |

| Result | model | LogLik | AIC | AICc | ΔAICc | num_para | root_theta( | theta_high | theta_low | sigma | sigma_hig | sigma_low | alpha | half_life |
|---|---|---|---|---|---|---|---|---|---|---|---|---|---|---|
| RESpT2 | OUM | -294.446 | 596.891 | 597.3789 | 0 | 4 | 6.682961 | 6.618584 | 11.68404 | 1.69E-08 | NA | NA | 6.628699 | 0.104594 |
| RESpT2 | BM1 | -297.038 | 598.0759 | 598.2188 | 0.84 | 2 | 10.68056 | NA | NA | 5.76E-01 | NA | NA | NA | NA |
| RESpT2 | BMM | -296.592 | 599.1846 | 599.4738 | 2.09 | 3 | 10.45806 | NA | NA | NA | 1.26E-10 | 0.94786 | NA | NA |
| RESpT2 | OU1 | -297.742 | 601.4843 | 601.7735 | 4.39 | 3 | 10.31878 | NA | NA | 1.19E+00 | NA | NA | 0.056186 | 12.33662 |
| RESpT2 | WN | -298.767 | 603.5335 | 603.8226 | 6.44 | 3 | 9.919536 | NA | NA | 5.63E+01 | NA | NA | NA | NA |

#### Heat (full phylogeny)

| Result | model | LogLik | AIC | AICc | ΔAICc | num_para | root_theta( | theta_high | theta_low | sigma | sigma_hig | sigma_low | alpha | half_life |
|---|---|---|---|---|---|---|---|---|---|---|---|---|---|---|
| IGR | BM1 | 105.4037 | -206.807 | -206.663 | 0 | 2 | 0.571841 | NA | NA | 2.55E-26 | NA | NA | NA | NA |
| IGR | WN | 105.4038 | -204.808 | -204.515 | 2.15 | 3 | 0.571841 | NA | NA | 5.05E-03 | NA | NA | NA | NA |
| IGR | BMM | 105.4037 | -204.807 | -204.515 | 2.15 | 3 | 0.571841 | NA | NA | NA | 4.97E-18 | 3.30E-18 | NA | NA |
| IGR | OU1 | 105.4037 | -204.807 | -204.515 | 2.15 | 3 | 0.571841 | NA | NA | 3.84E-27 | NA | NA | 0.093341 | 7.425997 |
| IGR | OUM | 105.5204 | -203.041 | -202.547 | 4.12 | 4 | 0.570295 | 0.576863 | 0.569346 | 4.96E-14 | NA | NA | 0.000468 | 1481.402 |

| Result | model | LogLik | AIC | AICc | ΔAICc | num_para | root_theta( | theta_high | theta_low | sigma | sigma_hig | sigma_low | alpha | half_life |
|---|---|---|---|---|---|---|---|---|---|---|---|---|---|---|
| MGR | OU1 | -36.4714 | 78.94274 | 79.22181 | 0 | 3 | 0.370747 | NA | NA | 0.013027 | NA | NA | 0.230802 | 3.003209 |
| MGR | OUM | -36.4049 | 80.80974 | 81.28033 | 2.06 | 4 | 0.369848 | 0.393342 | 0.361426 | 0.013039 | NA | NA | 0.231217 | 2.997822 |
| MGR | BM1 | -41.3129 | 86.62579 | 86.76372 | 7.54 | 2 | 0.386679 | NA | NA | 0.002485 | NA | NA | NA | NA |
| MGR | BMM | -40.5094 | 87.01886 | 87.29793 | 8.08 | 3 | 0.377967 | NA | NA | NA | 0.000659 | 0.003646 | NA | NA |
| MGR | WN | -40.8208 | 87.64165 | 87.92072 | 8.7 | 3 | 0.539052 | NA | NA | 0.14504 | NA | NA | NA | NA |

| Result | model | LogLik | AIC | AICc | ΔAICc | num_para | root_theta( | theta_high | theta_low | sigma | sigma_hig | sigma_low | alpha | half_life |
|---|---|---|---|---|---|---|---|---|---|---|---|---|---|---|
| XMID | WN | -254.33 | 514.6596 | 514.942 | 0 | 3 | 23.75042 | NA | NA | 17.768 | NA | NA | NA | NA |
| XMID | OUM | -261.272 | 530.5446 | 531.0208 | 16.08 | 4 | 21.8053 | 21.79405 | 23.65611 | 116.6036 | NA | NA | 14.59505 | 0.047495 |
| XMID | OU1 | -262.98 | 531.9608 | 532.2432 | 17.3 | 3 | 22.80657 | NA | NA | 5.21299 | NA | NA | 0.592061 | 1.170735 |
| XMID | BM1 | -264.885 | 533.7698 | 533.9094 | 18.97 | 2 | 22.34494 | NA | NA | 0.197241 | NA | NA | NA | NA |
| XMID | BMM | -264.539 | 535.0773 | 535.3597 | 20.42 | 3 | 22.42109 | NA | NA | NA | 0.299126 | 0.052178 | NA | NA |

| Result | model | LogLik | AIC | AICc | ΔAICc | num_para | root_theta( | theta_high | theta_low | sigma | sigma_hig | sigma_low | alpha | half_life |
|---|---|---|---|---|---|---|---|---|---|---|---|---|---|---|
| ASYM | OUM | -396.629 | 801.2572 | 801.7334 | 0 | 4 | 34.97924 | 23.65422 | 70.15951 | 54.74676 | NA | NA | 0.064409 | 10.76164 |
| ASYM | BMM | -398.906 | 803.811 | 804.0933 | 2.36 | 3 | 45.99845 | NA | NA | NA | 16.25319 | 40.24193 | NA | NA |
| ASYM | BM1 | -400.174 | 804.3481 | 804.4876 | 2.75 | 2 | 48.35494 | NA | NA | 30.845 | NA | NA | NA | NA |
| ASYM | OU1 | -400.877 | 807.7537 | 808.0361 | 6.3 | 3 | 48.8826 | NA | NA | 45.40588 | NA | NA | 0.03091 | 22.4245 |
| ASYM | WN | -420.022 | 846.0447 | 846.327 | 44.59 | 3 | 49.19182 | NA | NA | 735.7035 | NA | NA | NA | NA |

| Result | model | LogLik | AIC | AICc | ΔAICc | num_para | root_theta( | theta_high | theta_low | sigma | sigma_hig | sigma_low | alpha | half_life |
|---|---|---|---|---|---|---|---|---|---|---|---|---|---|---|
| NLEA | BM1 | -318.693 | 641.3861 | 641.5343 | 0 | 2 | 14.16497 | NA | NA | 8.64E-01 | NA | NA | NA | NA |
| NLEA | OU1 | -318.385 | 642.7709 | 643.0709 | 1.54 | 3 | 14.44845 | NA | NA | 3.84E+00 | NA | NA | 0.095826 | 7.233413 |
| NLEA | BMM | -318.407 | 642.814 | 643.114 | 1.58 | 3 | 14.17775 | NA | NA | NA | 0.617522 | 1.177774 | NA | NA |
| NLEA | WN | -319.181 | 644.3611 | 644.6611 | 3.13 | 3 | 14.55893 | NA | NA | 5.85E+01 | NA | NA | NA | NA |
| NLEA | OUM | -318.668 | 645.3364 | 645.8427 | 4.31 | 4 | 13.0431 | 12.89013 | 15.43508 | 3.25E-05 | NA | NA | 1.879465 | 0.368853 |

| Result | model | LogLik | AIC | AICc | ΔAICc | num_para | root_theta( | theta_high | theta_low | sigma | sigma_hig | sigma_low | alpha | half_life |
|---|---|---|---|---|---|---|---|---|---|---|---|---|---|---|
| LA | BMM | -605.469 | 1216.937 | 1217.23 | 0 | 3 | 358.8248 | NA | NA | NA | 1398.481 | 7435.237 | NA | NA |
| LA | BM1 | -608.794 | 1221.587 | 1221.732 | 4.5 | 2 | 393.5505 | NA | NA | 4672.391 | NA | NA | NA | NA |
| LA | OUM | -608.826 | 1225.652 | 1226.145 | 8.91 | 4 | 294.2134 | 109.2145 | 949.4656 | 5594.209 | NA | NA | 0.019644 | 35.28589 |
| LA | OU1 | -610.496 | 1226.992 | 1227.285 | 10.06 | 3 | 395.085 | NA | NA | 5619.748 | NA | NA | 0.013578 | 51.0497 |
| LA | WN | -630.635 | 1267.271 | 1267.563 | 50.33 | 3 | 409.6372 | NA | NA | 137044.4 | NA | NA | NA | NA |

| Result | model | LogLik | AIC | AICc | ΔAICc | num_para | root_theta( | theta_high | theta_low | sigma | sigma_hig | sigma_low | alpha | half_life |
|---|---|---|---|---|---|---|---|---|---|---|---|---|---|---|
| SLA | OU1 | -299.817 | 605.6339 | 605.9266 | 0 | 3 | 26.4019 | NA | NA | 8.448543 | NA | NA | 0.122807 | 5.644212 |
| SLA | OUM | -299.072 | 606.1446 | 606.6385 | 0.71 | 4 | 27.1726 | 28.87014 | 25.15555 | 8.933052 | NA | NA | 0.136861 | 5.064616 |
| SLA | BMM | -302.558 | 611.1161 | 611.4087 | 5.48 | 3 | 25.25685 | NA | NA | NA | 4.005508 | 1.488057 | NA | NA |
| SLA | WN | -302.607 | 611.2145 | 611.5072 | 5.58 | 3 | 29.23419 | NA | NA | 66.65304 | NA | NA | NA | NA |
| SLA | BM1 | -304.083 | 612.1657 | 612.3103 | 6.38 | 2 | 25.70928 | NA | NA | 2.708231 | NA | NA | NA | NA |

| Result | model | LogLik | AIC | AICc | ΔAICc | num_para | root_theta( | theta_high | theta_low | sigma | sigma_hig | sigma_low | alpha | half_life |
|---|---|---|---|---|---|---|---|---|---|---|---|---|---|---|
| LDMC | BMM | -235.06 | 476.1196 | 476.4159 | 0 | 3 | 15.34228 | NA | NA | NA | 0.044915 | 0.646601 | NA | NA |
| LDMC | BM1 | -237.064 | 478.1285 | 478.2749 | 1.86 | 2 | 15.88268 | NA | NA | 0.434275 | NA | NA | NA | NA |
| LDMC | OUM | -234.962 | 477.9239 | 478.4239 | 2.01 | 4 | 13.8642 | 12.4577 | 17.38341 | 1.013005 | NA | NA | 0.08939 | 7.75566 |
| LDMC | OU1 | -237.932 | 481.863 | 482.1593 | 5.74 | 3 | 15.77833 | NA | NA | 0.713337 | NA | NA | 0.034255 | 20.23479 |
| LDMC | WN | -247.673 | 501.3457 | 501.642 | 25.23 | 3 | 15.59853 | NA | NA | 19.88011 | NA | NA | NA | NA |

| Result | model | LogLik | AIC | AICc | ΔAICc | num_para | root_theta( | theta_high | theta_low | sigma | sigma_hig | sigma_low | alpha | half_life |
|---|---|---|---|---|---|---|---|---|---|---|---|---|---|---|

| | | | | | | | | | | | | | | |
|---|---|---|---|---|---|---|---|---|---|---|---|---|---|---|
| LTH | BM1 | 122.4799 | -240.96 | -240.806 | 0 | 2 | 0.266539 | NA | NA | 6.60E-06 | NA | NA | NA | NA |
| LTH | BMM | 123.0806 | -240.161 | -239.85 | 0.96 | 3 | 0.26388 | NA | NA | NA | 2.75E-05 | 1.71E-15 | NA | NA |
| LTH | OUM | 123.6463 | -239.293 | -238.766 | 2.04 | 4 | 0.275536 | 0.284685 | 0.256445 | 7.22E-13 | NA | NA | 0.001148 | 611.6538 |
| LTH | OU1 | 122.4317 | -238.863 | -238.552 | 2.25 | 3 | 0.267006 | NA | NA | 3.45E-05 | NA | NA | 0.093 | 7.453163 |
| LTH | WN | 122.2344 | -238.469 | -238.157 | 2.65 | 3 | 0.267296 | NA | NA | 2.86E-03 | NA | NA | NA | NA |

| Result | model | LogLik | AIC | AICc | ΔAICc | num_parar | root_theta( | theta_high | theta_low | sigma | sigma_higl | sigma_low | alpha | half_life |
|---|---|---|---|---|---|---|---|---|---|---|---|---|---|---|
| LDI | BMM | -63.4825 | 132.9649 | 133.2576 | 0 | 3 | 1.903958 | NA | NA | NA | 0.005258 | 0.023265 | NA | NA |
| LDI | OUM | -63.7317 | 135.4635 | 135.9573 | 2.7 | 4 | 1.762806 | 1.565796 | 2.089806 | 0.034159 | NA | NA | 0.088064 | 7.870929 |
| LDI | OU1 | -65.2842 | 136.5685 | 136.8612 | 3.6 | 3 | 1.92259 | NA | NA | 0.030039 | NA | NA | 0.062654 | 11.06308 |
| LDI | BM1 | -66.7635 | 137.527 | 137.6716 | 4.41 | 2 | 1.965588 | NA | NA | 0.015894 | NA | NA | NA | NA |
| LDI | WN | -77.9445 | 161.889 | 162.1817 | 28.92 | 3 | 1.942379 | NA | NA | 0.35872 | NA | NA | NA | NA |

| Result | model | LogLik | AIC | AICc | ΔAICc | num_parar | root_theta( | theta_high | theta_low | sigma | sigma_higl | sigma_low | alpha | half_life |
|---|---|---|---|---|---|---|---|---|---|---|---|---|---|---|
| RESpT1 | BM1 | -183.369 | 370.7384 | 371.0175 | 0 | 2 | 57.02283 | NA | NA | 5.34E-16 | NA | NA | NA | NA |
| RESpT1 | WN | -183.364 | 372.7274 | 373.2988 | 2.28 | 3 | 57.02283 | NA | NA | 1.70E+02 | NA | NA | NA | NA |
| RESpT1 | BMM | -183.369 | 372.7376 | 373.3091 | 2.29 | 3 | 57.02283 | NA | NA | NA | 1.32E-10 | 1.86E-12 | NA | NA |
| RESpT1 | OU1 | -183.369 | 372.7384 | 373.3098 | 2.29 | 3 | 57.02283 | NA | NA | 4.43E-09 | NA | NA | 20.7317 | 0.033434 |
| RESpT1 | OUM | -182.95 | 373.9004 | 374.876 | 3.86 | 4 | 54.46667 | 54.46667 | 58.25968 | 3.40E-07 | NA | NA | 36.49379 | 0.018994 |

| Result | model | LogLik | AIC | AICc | ΔAICc | num_parar | root_theta( | theta_high | theta_low | sigma | sigma_higl | sigma_low | alpha | half_life |
|---|---|---|---|---|---|---|---|---|---|---|---|---|---|---|
| RESpT2 | OUM | -346.703 | 701.4052 | 701.899 | 0 | 4 | 19.90238 | 21.41821 | 13.157 | 133.9275 | NA | NA | 0.705141 | 0.982997 |
| RESpT2 | OU1 | -348.594 | 703.1872 | 703.4799 | 1.58 | 3 | 15.3512 | NA | NA | 66.17052 | NA | NA | 0.312603 | 2.217338 |
| RESpT2 | WN | -349.079 | 704.158 | 704.4507 | 2.55 | 3 | 18.45934 | NA | NA | 196.4169 | NA | NA | NA | NA |
| RESpT2 | BMM | -354.558 | 715.1153 | 715.408 | 13.51 | 3 | 15.39927 | NA | NA | NA | 17.69318 | 5.686797 | NA | NA |
| RESpT2 | BM1 | -355.823 | 715.6456 | 715.7902 | 13.89 | 2 | 15.48957 | NA | NA | 8.48579 | NA | NA | NA | NA |

| Result | model | LogLik | AIC | AICc | ΔAICc | num_parar | root_theta( | theta_high | theta_low | sigma | sigma_higl | sigma_low | alpha | half_life |
|---|---|---|---|---|---|---|---|---|---|---|---|---|---|---|
| TOL_IGR | BM1 | 27.75295 | -51.5059 | -51.3415 | 0 | 2 | 0.021701 | NA | NA | 5.55E-26 | NA | NA | NA | NA |
| TOL_IGR | WN | 27.75295 | -49.5059 | -49.1726 | 2.17 | 3 | 0.021701 | NA | NA | 2.82E-02 | NA | NA | NA | NA |
| TOL_IGR | BMM | 27.75295 | -49.5059 | -49.1726 | 2.17 | 3 | 0.021701 | NA | NA | NA | 3.22E-17 | 1.14E-17 | NA | NA |
| TOL_IGR | OU1 | 27.75295 | -49.5059 | -49.1726 | 2.17 | 3 | 0.021701 | NA | NA | 1.11E-20 | NA | NA | 0.100985 | 6.863867 |
| TOL_IGR | OUM | 27.83178 | -47.6636 | -47.1002 | 4.24 | 4 | 0.019368 | 0.028279 | 0.018902 | 1.02E-13 | NA | NA | 0.079606 | 8.707239 |

| Result | model | LogLik | AIC | AICc | ΔAICc | num_parar | root_theta( | theta_high | theta_low | sigma | sigma_higl | sigma_low | alpha | half_life |
|---|---|---|---|---|---|---|---|---|---|---|---|---|---|---|
| TOL_MGR | OUM | -155.298 | 318.5952 | 319.1586 | 0 | 4 | 2.288214 | 2.443176 | 1.517032 | 6.139177 | NA | NA | 0.846862 | 0.818489 |
| TOL_MGR | OU1 | -156.812 | 319.6247 | 319.958 | 0.8 | 3 | 1.811094 | NA | NA | 5.910328 | NA | NA | 0.780298 | 0.888311 |
| TOL_MGR | WN | -158.482 | 322.9632 | 323.2965 | 4.14 | 3 | 1.797652 | NA | NA | 3.791232 | NA | NA | NA | NA |
| TOL_MGR | BMM | -176.546 | 359.092 | 359.4253 | 40.27 | 3 | 1.601739 | NA | NA | NA | 2.025604 | 0.468728 | NA | NA |
| TOL_MGR | BM1 | -186.249 | 376.4978 | 376.6622 | 57.5 | 2 | 1.82147 | NA | NA | 1.083437 | NA | NA | NA | NA |

| Result | model | LogLik | AIC | AICc | ΔAICc | num_parar | root_theta( | theta_high | theta_low | sigma | sigma_higl | sigma_low | alpha | half_life |
|---|---|---|---|---|---|---|---|---|---|---|---|---|---|---|
| TOL_XMID | OUM | 24.99031 | -41.9806 | -41.4172 | 0 | 4 | -0.08504 | -0.10829 | 0.002202 | 0.026493 | NA | NA | 0.395385 | 1.753093 |
| TOL_XMID | OU1 | 23.20334 | -40.4067 | -40.0733 | 1.34 | 3 | -0.03101 | NA | NA | 0.024677 | NA | NA | 0.344247 | 2.013519 |
| TOL_XMID | WN | 21.04655 | -36.0931 | -35.7598 | 5.66 | 3 | -0.03682 | NA | NA | 0.03365 | NA | NA | NA | NA |
| TOL_XMID | BM1 | 6.352536 | -8.70507 | -8.54069 | 32.88 | 2 | -0.00095 | NA | NA | 0.006817 | NA | NA | NA | NA |
| TOL_XMID | BMM | 6.91611 | -7.83222 | -7.49889 | 33.92 | 3 | 0.002371 | NA | NA | NA | 0.008421 | 0.005901 | NA | NA |

| Result | model | LogLik | AIC | AICc | ΔAICc | num_parar | root_theta( | theta_high | theta_low | sigma | sigma_higl | sigma_low | alpha | half_life |
|---|---|---|---|---|---|---|---|---|---|---|---|---|---|---|
| TOL_ASYM | OUM | -111.756 | 231.5123 | 232.0757 | 0 | 4 | -0.18282 | -0.30392 | 0.781733 | 1.988704 | NA | NA | 0.854097 | 0.811555 |
| TOL_ASYM | OU1 | -117.825 | 241.65 | 241.9833 | 9.91 | 3 | 0.433554 | NA | NA | 1.795807 | NA | NA | 0.653398 | 1.060834 |
| TOL_ASYM | WN | -121.203 | 248.4066 | 248.7399 | 16.66 | 3 | 0.438219 | NA | NA | 1.421459 | NA | NA | NA | NA |
| TOL_ASYM | BMM | -146.888 | 299.7764 | 300.1097 | 68.03 | 3 | 0.252076 | NA | NA | NA | 0.173269 | 0.56718 | NA | NA |
| TOL_ASYM | BM1 | -152.204 | 308.4072 | 308.5716 | 76.5 | 2 | 0.389545 | NA | NA | 0.44229 | NA | NA | NA | NA |

### Frost (bootstrapped phylogeny)

| Result | model | LogLik | AIC | AICc | ΔAICc | num_params | root_theta0 | theta_high | theta_low | sigma | sigma_high | sigma_low | alpha | half_life |
|---|---|---|---|---|---|---|---|---|---|---|---|---|---|---|
| IGR | BM1 | 95.34349 | -186.687 | -186.4903 | 0.00 | 2 | 0.5771454 | NA | NA | 5.45E-21 | NA | NA | NA | NA |
| IGR | BMM | 95.64448 | -185.289 | -184.8949 | 1.60 | 3 | 0.5756188 | NA | NA | NA | 3.02E-07 | 1.53E-15 | NA | NA |
| IGR | OU1 | 95.41899 | -184.838 | -184.438 | 2.05 | 3 | 0.5771454 | NA | NA | 3.89E-11 | NA | NA | 0.09307075 | 7.44753 |
| IGR | WN | 95.37817 | -184.7563 | -184.3458 | 2.14 | 3 | 0.5772459 | NA | NA | 2.34E-03 | NA | NA | NA | NA |
| IGR | OUM | 96.49793 | -184.9959 | -184.2891 | 2.20 | 4 | 0.5792877 | 0.5904825 | 0.5685765 | 7.16E-13 | NA | NA | 0.12723178 | 5.44791 |

| Result | model | LogLik | AIC | AICc | ΔAICc | num_params | root_theta0 | theta_high | theta_low | sigma | sigma_high | sigma_low | alpha | half_life |
|---|---|---|---|---|---|---|---|---|---|---|---|---|---|---|
| MGR | OU1 | 59.59174 | -113.18347 | -112.78654 | 0.00 | 3 | 0.2128647 | NA | NA | 0.020538807 | NA | NA | 2.64465 | 0.2621029 |
| MGR | OUM | 59.9329 | -111.8658 | -111.18184 | 1.60 | 4 | 0.2131173 | 0.2172296 | 0.2108829 | 0.01682384 | NA | NA | 2.655911 | 0.2609829 |
| MGR | BMM | 56.56991 | -107.13982 | -106.74627 | 6.04 | 3 | 0.2091986 | NA | NA | NA | 0.001221568 | 4.89E-05 | NA | NA |
| MGR | BM1 | 43.71264 | -83.42528 | -83.23327 | 29.55 | 2 | 0.2138872 | NA | NA | 0.000486681 | NA | NA | NA | NA |
| MGR | WN | 29.19296 | -52.38593 | -51.98856 | 60.80 | 3 | 0.2459143 | NA | NA | 0.012019988 | NA | NA | NA | NA |

| Result | model | LogLik | AIC | AICc | ΔAICc | num_params | root_theta0 | theta_high | theta_low | sigma | sigma_high | sigma_low | alpha | half_life |
|---|---|---|---|---|---|---|---|---|---|---|---|---|---|---|
| XMID | WN | -184.7462 | 375.4924 | 375.8958 | 0.00 | 3 | 26.00079 | NA | NA | 18.295458 | NA | NA | NA | NA |
| XMID | OU1 | -194.775 | 395.5501 | 395.957 | 20.06 | 3 | 25.18824 | NA | NA | 28.685312 | NA | NA | 3.129154 | 0.2215517 |
| XMID | OUM | -193.7666 | 395.5332 | 396.2207 | 20.32 | 4 | 24.84619 | 24.03057 | 25.84682 | 15.102729 | NA | NA | 1.578458 | 0.4391294 |
| XMID | BM1 | -197.2119 | 398.4238 | 398.6238 | 22.73 | 2 | 25.08986 | NA | NA | 0.371684 | NA | NA | NA | NA |
| XMID | BMM | -196.4502 | 398.9004 | 399.3072 | 23.41 | 3 | 25.03914 | NA | NA | NA | 0.2996011 | 0.4038964 | NA | NA |

| Result | model | LogLik | AIC | AICc | ΔAICc | num_params | root_theta0 | theta_high | theta_low | sigma | sigma_high | sigma_low | alpha | half_life |
|---|---|---|---|---|---|---|---|---|---|---|---|---|---|---|
| ASYM | OUM | -271.101 | 550.2019 | 550.8904 | 0.00 | 4 | 41.17834 | 30.02935 | 56.11459 | 32.37123 | NA | NA | 0.07053518 | 9.826971 |
| ASYM | BM1 | -273.4649 | 550.9298 | 551.1282 | 0.24 | 2 | 46.86057 | NA | NA | 16.25364 | NA | NA | NA | NA |
| ASYM | BMM | -273.0433 | 552.0866 | 552.4934 | 1.60 | 3 | 46.17249 | NA | NA | NA | 14.47669 | 17.78111 | NA | NA |
| ASYM | OU1 | -273.7363 | 553.4727 | 553.8795 | 2.99 | 3 | 46.42162 | NA | NA | 27.9215 | NA | NA | 0.04306619 | 16.096378 |
| ASYM | WN | -283.6866 | 573.3732 | 573.78 | 22.89 | 3 | 46.0312 | NA | NA | 431.11361 | NA | NA | NA | NA |

| Result | model | LogLik | AIC | AICc | ΔAICc | num_params | root_theta0 | theta_high | theta_low | sigma | sigma_high | sigma_low | alpha | half_life |
|---|---|---|---|---|---|---|---|---|---|---|---|---|---|---|
| NLEA | BM1 | -223.967 | 451.934 | 452.1448 | 0.00 | 2 | 13.38796 | NA | NA | 0.2852644 | NA | NA | NA | NA |
| NLEA | OU1 | -223.0608 | 452.1217 | 452.5661 | 0.42 | 3 | 13.54488 | NA | NA | 3.987463 | NA | NA | 0.2464543 | 2.8168227 |
| NLEA | BMM | -223.2705 | 452.5409 | 452.9854 | 0.84 | 3 | 13.15935 | NA | NA | NA | 3.07E-10 | 0.4865884 | NA | NA |
| NLEA | WN | -223.8138 | 453.6277 | 454.0721 | 1.93 | 3 | 13.47826 | NA | NA | 74.7612487 | NA | NA | NA | NA |
| NLEA | OUM | -222.7742 | 453.5485 | 454.3029 | 2.16 | 4 | 12.78486 | 11.94412 | 14.24377 | 3.0631068 | NA | NA | 0.7510527 | 0.9229008 |

| Result | model | LogLik | AIC | AICc | ΔAICc | num_params | root_theta0 | theta_high | theta_low | sigma | sigma_high | sigma_low | alpha | half_life |
|---|---|---|---|---|---|---|---|---|---|---|---|---|---|---|
| LA | BMM | -438.8465 | 883.6931 | 884.1002 | 0.00 | 3 | 347.1932 | NA | NA | NA | 885.1515 | 4725.532 | NA | NA |
| LA | BM1 | -440.5301 | 885.0601 | 885.2636 | 1.16 | 2 | 381.9863 | NA | NA | 3308.932 | NA | NA | NA | NA |
| LA | OUM | -438.5992 | 885.1984 | 885.8997 | 1.80 | 4 | 291.1999 | 161.5481 | 474.2916 | 8021.005 | NA | NA | 0.09182253 | 7.54877 |
| LA | OU1 | -440.9904 | 887.9808 | 888.3946 | 4.29 | 3 | 375.3102 | NA | NA | 6411.603 | NA | NA | 0.05334056 | 12.99745 |
| LA | WN | -452.6265 | 911.2529 | 911.6667 | 27.57 | 3 | 394.7183 | NA | NA | 101525.311 | NA | NA | NA | NA |

| Result | model | LogLik | AIC | AICc | ΔAICc | num_params | root_theta0 | theta_high | theta_low | sigma | sigma_high | sigma_low | alpha | half_life |
|---|---|---|---|---|---|---|---|---|---|---|---|---|---|---|
| SLA | BMM | -204.6006 | 415.2012 | 415.6174 | 0.00 | 3 | 20.63452 | NA | NA | NA | 2.021628 | 0.3064765 | NA | NA |
| SLA | BM1 | -206.4877 | 416.9753 | 417.1787 | 1.56 | 2 | 20.91267 | NA | NA | 1.06516 | NA | NA | NA | NA |
| SLA | OU1 | -205.4735 | 416.947 | 417.3539 | 1.74 | 3 | 21.15704 | NA | NA | 3.077535 | NA | NA | 0.09397477 | 7.376014 |
| SLA | OUM | -205.1545 | 418.309 | 419.0085 | 3.39 | 4 | 21.22938 | 21.57654 | 20.84345 | 2.998925 | NA | NA | 0.09517908 | 7.282558 |
| SLA | WN | -213.602 | 433.2039 | 433.6142 | 18.00 | 3 | 22.69825 | NA | NA | 49.394064 | NA | NA | NA | NA |

| Result | model | LogLik | AIC | AICc | ΔAICc | num_params | root_theta0 | theta_high | theta_low | sigma | sigma_high | sigma_low | alpha | half_life |
|---|---|---|---|---|---|---|---|---|---|---|---|---|---|---|
| LDMC | OUM | -173.4647 | 354.9294 | 355.639 | 0.00 | 4 | 17.48517 | 16.75738 | 20.60613 | 8.4979723 | NA | NA | 0.6066543 | 1.142574 |
| LDMC | OU1 | -177.3278 | 360.6557 | 361.0625 | 5.42 | 3 | 19.29139 | NA | NA | 3.3249597 | NA | NA | 0.1219474 | 5.684231 |
| LDMC | WN | -179.1471 | 364.2943 | 364.7081 | 9.07 | 3 | 19.32963 | NA | NA | 17.8045724 | NA | NA | NA | NA |
| LDMC | BMM | -179.2225 | 364.445 | 364.8489 | 9.21 | 3 | 19.1751 | NA | NA | NA | 0.4383984 | 1.227271 | NA | NA |
| LDMC | BM1 | -180.5748 | 365.1496 | 365.3583 | 9.72 | 2 | 19.48723 | NA | NA | 0.9629076 | NA | NA | NA | NA |

| Result | model | LogLik | AIC | AICc | ΔAICc | num_params | root_theta0 | theta_high | theta_low | sigma | sigma_high | sigma_low | alpha | half_life |
|---|---|---|---|---|---|---|---|---|---|---|---|---|---|---|
| LTH | BM1 | 75.79719 | -147.5944 | -147.3614 | 0.00 | 2 | 0.2952818 | NA | NA | 2.13E-05 | NA | NA | NA | NA |
| LTH | BMM | 75.97914 | -145.9583 | -145.4973 | 1.86 | 3 | 0.2945349 | NA | NA | NA | 2.96E-05 | 1.29E-05 | NA | NA |
| LTH | OU1 | 75.71585 | -145.4317 | -144.97 | 2.39 | 3 | 0.2957558 | NA | NA | 6.32E-05 | NA | NA | 0.07650615 | 9.060031 |
| LTH | WN | 75.46099 | -144.922 | -144.4456 | 2.92 | 3 | 0.2957183 | NA | NA | 3.49E-03 | NA | NA | NA | NA |
| LTH | OUM | 76.06535 | -144.1307 | -143.3484 | 4.01 | 4 | 0.2959055 | 0.3000083 | 0.2929576 | 4.70E-05 | NA | NA | 0.06524476 | 10.623798 |

| Result | model | LogLik | AIC | AICc | ΔAICc | num_params | root_theta0 | theta_high | theta_low | sigma | sigma_high | sigma_low | alpha | half_life |
|---|---|---|---|---|---|---|---|---|---|---|---|---|---|---|
| LDI | BMM | -43.89934 | 93.79867 | 94.21398 | 0.00 | 3 | 1.935254 | NA | NA | NA | 0.003349727 | 0.0126504 | NA | NA |
| LDI | BM1 | -45.31416 | 94.62833 | 94.83172 | 0.62 | 2 | 1.969478 | NA | NA | 0.009306049 | NA | NA | NA | NA |
| LDI | OU1 | -45.85312 | 97.70624 | 98.12004 | 3.91 | 3 | 1.929377 | NA | NA | 0.016515866 | NA | NA | 0.04421579 | 15.67766 |
| LDI | OUM | -45.04839 | 98.09678 | 98.79527 | 4.58 | 4 | 1.904452 | 1.782163 | 2.006012 | 0.017099287 | NA | NA | 0.05127144 | 13.51917 |
| LDI | WN | -55.16038 | 116.32075 | 116.72753 | 22.51 | 3 | 1.969931 | NA | NA | 0.302859123 | NA | NA | NA | NA |

| Result | model | LogLik | AIC | AICc | ΔAICc | num_params | root_theta0 | theta_high | theta_low | sigma | sigma_high | sigma_low | alpha | half_life |
|---|---|---|---|---|---|---|---|---|---|---|---|---|---|---|
| RESmT1 | BM1 | -167.3045 | 338.6089 | 338.9128 | 0.00 | 2 | 58.08702 | NA | NA | 4.27E-17 | NA | NA | NA | NA |
| RESmT1 | BMM | -167.0855 | 340.171 | 340.7904 | 1.88 | 3 | 58.20784 | NA | NA | NA | 5.01E-12 | 1.02E-10 | NA | NA |
| RESmT1 | OU1 | -167.1599 | 340.3198 | 340.9513 | 2.04 | 3 | 58.08689 | NA | NA | 2.84E-06 | NA | NA | 15.403701 | 0.04499875 |
| RESmT1 | WN | -167.171 | 340.342 | 340.9655 | 2.05 | 3 | 58.07602 | NA | NA | 1.29E+02 | NA | NA | NA | NA |
| RESmT1 | OUM | -166.7825 | 341.565 | 342.6139 | 3.70 | 4 | 57.35257 | 56.14156 | 59.32276 | 1.75E-06 | NA | NA | 1.916014 | 0.36176526 |

| Result | model | LogLik | AIC | AICc | ΔAICc | num_params | root_theta0 | theta_high | theta_low | sigma | sigma_high | sigma_low | alpha | half_life |
|---|---|---|---|---|---|---|---|---|---|---|---|---|---|---|
| RESmT2 | WN | -272.3494 | 550.6989 | 551.1023 | 0.00 | 3 | 43.31033 | NA | NA | 300.801297 | NA | NA | NA | NA |
| RESmT2 | BM1 | -288.245 | 580.49 | 580.6901 | 29.59 | 2 | 43.81127 | NA | NA | 2.408543 | NA | NA | NA | NA |
| RESmT2 | BMM | -287.6373 | 581.2746 | 581.6802 | 30.58 | 3 | 44.5422 | NA | NA | NA | 1.34E-10 | 4.93284 | NA | NA |
| RESmT2 | OUM | -286.5864 | 581.1728 | 581.8624 | 30.76 | 4 | 46.50177 | 50.50145 | 40.16165 | 22.551743 | NA | NA | 0.3833366 | 1.808195 |
| RESmT2 | OU1 | -287.8868 | 581.7735 | 582.1769 | 31.07 | 3 | 44.65604 | NA | NA | 38.716924 | NA | NA | 0.2279655 | 3.042265 |

| Result | model | LogLik | AIC | AICc | ΔAICc | num_params | root_theta0 | theta_high | theta_low | sigma | sigma_high | sigma_low | alpha | half_life |
|---|---|---|---|---|---|---|---|---|---|---|---|---|---|---|
| TOL_IGR | BM1 | 35.08086 | -66.16171 | -65.91671 | 0.00 | 2 | 0.02165361 | NA | NA | 9.26E-21 | NA | NA | NA | NA |
| TOL_IGR | BMM | 35.22113 | -64.44227 | -63.9534 | 1.96 | 3 | 0.02134057 | NA | NA | NA | 8.86E-15 | 1.42E-15 | NA | NA |
| TOL_IGR | OU1 | 35.048 | -64.09601 | -63.60621 | 2.31 | 3 | 0.02172189 | NA | NA | 1.96E-18 | NA | NA | 0.0964923 | 7.183445 |
| TOL_IGR | WN | 34.99609 | -63.99219 | -63.51199 | 2.40 | 3 | 0.02165361 | NA | NA | 1.35E-02 | NA | NA | NA | NA |
| TOL_IGR | OUM | 36.16864 | -64.33728 | -63.49171 | 2.43 | 4 | 0.0303166 | 0.05764406 | -0.000603225 | 2.63E-13 | NA | NA | 0.1372543 | 5.050093 |

| Result | model | LogLik | AIC | AICc | ΔAICc | num_params | root_theta0 | theta_high | theta_low | sigma | sigma_high | sigma_low | alpha | half_life |
|---|---|---|---|---|---|---|---|---|---|---|---|---|---|---|
| TOL_MGR | OU1 | -30.01415 | 66.0283 | 66.52202 | 0.00 | 3 | 0.10922493 | NA | NA | 2.5833227 | NA | NA | 6.780388 | 0.10227851 |
| TOL_MGR | WN | -30.01547 | 66.03094 | 66.54713 | 0.03 | 3 | 0.10821106 | NA | NA | 0.1522989 | NA | NA | NA | NA |

| | | | | | | | | | | | | | | |
|---|---|---|---|---|---|---|---|---|---|---|---|---|---|---|
| TOL_MGR | OUM | -29.62994 | 67.25988 | 68.07159 | 1.55 | 4 | 0.11216866 | 0.1527408 | 0.08479186 | 3.5546958 | NA | NA | 9.869934 | 0.07022815 |
| TOL_MGR | BMM | -67.30789 | 140.61579 | 141.12891 | 74.61 | 3 | 0.0531556 | NA | NA | NA | 0.04697763 | 0.06113335 | NA | NA |
| TOL_MGR | BM1 | -74.35189 | 152.70377 | 152.94682 | 86.42 | 2 | 0.06044816 | NA | NA | 0.1324689 | NA | NA | NA | NA |

| **Result** | **model** | **LogLik** | **AIC** | **AICc** | **ΔAICc** | **num_params** | **root_theta0** | **theta_high** | **theta_low** | **sigma** | **sigma_high** | **sigma_low** | **alpha** | **half_life** |
|---|---|---|---|---|---|---|---|---|---|---|---|---|---|---|
| TOL_XMID | WN | 23.538994 | -41.07799 | -40.55625 | 0.00 | 3 | -0.02524179 | NA | NA | 0.019858819 | NA | NA | NA | NA |
| TOL_XMID | OU1 | 23.538993 | -41.07799 | -40.55625 | 0.00 | 3 | -0.02524178 | NA | NA | 0.869945184 | NA | NA | 18.09495 | 0.03830636 |
| TOL_XMID | OUM | 23.922551 | -39.8451 | -38.9761 | 1.58 | 4 | -0.03082673 | -0.04039956 | -0.01780265 | 0.778969403 | NA | NA | 17.228 | 0.04023375 |
| TOL_XMID | BMM | -5.499421 | 16.99884 | 17.51182 | 58.07 | 3 | -0.02423231 | NA | NA | NA | 0.01187382 | 0.007980026 | NA | NA |
| TOL_XMID | BM1 | -7.766353 | 19.53271 | 19.78016 | 60.34 | 2 | -0.02551727 | NA | NA | 0.009613321 | NA | NA | NA | NA |

| **Result** | **model** | **LogLik** | **AIC** | **AICc** | **ΔAICc** | **num_params** | **root_theta0** | **theta_high** | **theta_low** | **sigma** | **sigma_high** | **sigma_low** | **alpha** | **half_life** |
|---|---|---|---|---|---|---|---|---|---|---|---|---|---|---|
| TOL_ASYM | OU1 | 7.620806 | -9.241611 | -8.735916 | 0.00 | 3 | -0.08212485 | NA | NA | 1.49628874 | NA | NA | 18.98417 | 0.03652459 |
| TOL_ASYM | OUM | 8.763316 | -9.526633 | -8.637742 | 0.10 | 4 | -0.10612584 | -0.1344253 | -0.05230526 | 1.81383604 | NA | NA | 22.29539 | 0.03108926 |
| TOL_ASYM | WN | 7.55507 | -9.11014 | -8.604821 | 0.13 | 3 | -0.08197218 | NA | NA | 0.04000776 | NA | NA | NA | NA |
| TOL_ASYM | BMM | -32.795331 | 71.590663 | 72.103443 | 80.84 | 3 | -0.06702275 | NA | NA | NA | 0.02577003 | 0.025933 | NA | NA |
| TOL_ASYM | BM1 | -35.309737 | 74.619474 | 74.872133 | 83.61 | 2 | -0.0664359 | NA | NA | 0.02658426 | NA | NA | NA | NA |

#### Mild (control) (bootstrapped phylogeny)

| Result | model | LogLik | AIC | AICc | ΔAICc | num_params | root_theta0 | theta_high | theta_low | sigma | sigma_high | sigma_low | alpha | half_life |
|---|---|---|---|---|---|---|---|---|---|---|---|---|---|---|
| SSIZ | BMM | -19.45832 | 44.91664 | 45.30337 | 0.00 | 3 | 0.6447267 | NA | NA | NA | 0.0217687 | 0.0091142 | NA | NA |
| SSIZ | BM1 | -22.10004 | 48.20007 | 48.3906 | 3.09 | 2 | 0.6393159 | NA | NA | 0.0136394 | NA | NA | NA | NA |
| SSIZ | OU1 | -22.68605 | 51.3721 | 51.7592 | 6.46 | 3 | 0.610894 | NA | NA | 0.0180907 | NA | NA | 0.0283544 | 24.44639 |
| SSIZ | OUM | -22.23456 | 52.46913 | 53.12703 | 7.82 | 4 | 0.5964919 | 0.4967265 | 0.6548367 | 0.0180192 | NA | NA | 0.0291963 | 23.7409 |
| SSIZ | WN | -37.77636 | 81.55272 | 81.93675 | 36.63 | 3 | 0.4559634 | NA | NA | 0.1592683 | NA | NA | NA | NA |

| Result | model | LogLik | AIC | AICc | ΔAICc | num_params | root_theta0 | theta_high | theta_low | sigma | sigma_high | sigma_low | alpha | half_life |
|---|---|---|---|---|---|---|---|---|---|---|---|---|---|---|
| TGER | BM1 | -173.6364 | 351.2728 | 351.4679 | 0.00 | 2 | 16.61919 | NA | NA | 0.1313361 | NA | NA | NA | NA |
| TGER | BMM | -173.0042 | 352.0084 | 352.4057 | 0.94 | 3 | 16.60441 | NA | NA | NA | 2.96E-10 | 0.2066662 | NA | NA |
| TGER | OU1 | -174.787 | 355.574 | 355.9774 | 4.51 | 3 | 16.41958 | NA | NA | 0.3569506 | NA | NA | 0.058496 | 11.852678 |
| TGER | OUM | -174.1618 | 356.3236 | 357.0013 | 5.53 | 4 | 16.57606 | 17.08659 | 15.6942 | 0.5294165 | NA | NA | 0.2178009 | 3.182488 |
| TGER | WN | -177.8502 | 361.7003 | 362.1037 | 10.64 | 3 | 15.09396 | NA | NA | 14.526194 | NA | NA | NA | NA |

| Result | model | LogLik | AIC | AICc | ΔAICc | num_params | root_theta0 | theta_high | theta_low | sigma | sigma_high | sigma_low | alpha | half_life |
|---|---|---|---|---|---|---|---|---|---|---|---|---|---|---|
| IGR | BM1 | 97.70284 | -191.4057 | -191.1987 | 0.00 | 2 | 0.5808454 | NA | NA | 6.32E-18 | NA | NA | NA | NA |
| IGR | BMM | 97.95363 | -189.9073 | -189.4787 | 1.72 | 3 | 0.5812548 | NA | NA | NA | 2.52E-06 | 3.24E-13 | NA | NA |
| IGR | OUM | 99.05231 | -190.1046 | -189.3909 | 1.81 | 4 | 0.5780019 | 0.5657411 | 0.5888455 | 6.98E-12 | NA | NA | 0.0997758 | 6.947049 |
| IGR | WN | 97.79075 | -189.5815 | -189.1529 | 2.05 | 3 | 0.5795695 | NA | NA | 2.25E-03 | NA | NA | NA | NA |
| IGR | OU1 | 97.73291 | -189.4658 | -189.0446 | 2.15 | 3 | 0.580042 | NA | NA | 2.01E-05 | NA | NA | 0.0927235 | 7.475424 |

| Result | model | LogLik | AIC | AICc | ΔAICc | num_params | root_theta0 | theta_high | theta_low | sigma | sigma_high | sigma_low | alpha | half_life |
|---|---|---|---|---|---|---|---|---|---|---|---|---|---|---|
| MGR | OUM | 101.9327 | -195.8653 | -195.2033 | 0.00 | 4 | 0.1894353 | 0.1832986 | 0.2097854 | 4.30E-05 | NA | NA | 4.2091561 | 0.1646762 |
| MGR | BM1 | 99.2405 | -194.481 | -194.2842 | 0.92 | 2 | 0.2002217 | NA | NA | 6.70E-06 | NA | NA | NA | NA |
| MGR | OU1 | 99.6319 | -193.2638 | -192.8671 | 2.34 | 3 | 0.2010702 | NA | NA | 4.14E-05 | NA | NA | 0.0932645 | 7.432061 |
| MGR | BMM | 99.48067 | -192.9613 | -192.5475 | 2.66 | 3 | 0.1996722 | NA | NA | NA | 8.50E-09 | 8.43E-06 | NA | NA |
| MGR | WN | 90.39422 | -174.7884 | -174.395 | 20.81 | 3 | 0.2195547 | NA | NA | 3.03E-03 | NA | NA | NA | NA |

| Result | model | LogLik | AIC | AICc | ΔAICc | num_params | root_theta0 | theta_high | theta_low | sigma | sigma_high | sigma_low | alpha | half_life |
|---|---|---|---|---|---|---|---|---|---|---|---|---|---|---|
| XMID | WN | -180.6756 | 367.3512 | 367.7546 | 0.00 | 3 | 27.49705 | NA | NA | 15.210341 | NA | NA | NA | NA |
| XMID | BM1 | -198.6216 | 401.2433 | 401.4416 | 33.69 | 2 | 25.79103 | NA | NA | 0.3255048 | NA | NA | NA | NA |
| XMID | OU1 | -197.8775 | 401.7551 | 402.1585 | 34.40 | 3 | 26.0345 | NA | NA | 1.8211187 | NA | NA | 0.1414793 | 4.899701 |
| XMID | BMM | -198.0435 | 402.0869 | 402.4896 | 34.74 | 3 | 25.86572 | NA | NA | NA | 0.0594225 | 0.4040632 | NA | NA |
| XMID | OUM | -197.3981 | 402.7962 | 403.4815 | 35.73 | 4 | 26.05989 | 26.62388 | 25.66961 | 1.7659281 | NA | NA | 0.1429443 | 4.849073 |

| Result | model | LogLik | AIC | AICc | ΔAICc | num_params | root_theta0 | theta_high | theta_low | sigma | sigma_high | sigma_low | alpha | half_life |
|---|---|---|---|---|---|---|---|---|---|---|---|---|---|---|
| ASYM | BM1 | -278.9869 | 561.9739 | 562.1722 | 0.00 | 2 | 52.4008 | NA | NA | 19.80736 | NA | NA | NA | NA |
| ASYM | BMM | -278.7973 | 563.5946 | 563.9946 | 1.82 | 3 | 52.25478 | NA | NA | NA | 18.24642 | 21.71689 | NA | NA |
| ASYM | OUM | -278.1372 | 564.2743 | 564.9484 | 2.78 | 4 | 49.14824 | 37.14129 | 62.41582 | 33.57655 | NA | NA | 0.049621 | 13.96883 |
| ASYM | OU1 | -279.5514 | 565.1027 | 565.5028 | 3.33 | 3 | 52.36206 | NA | NA | 31.18928 | NA | NA | 0.0366645 | 18.90539 |
| ASYM | WN | -295.9178 | 597.8356 | 598.2356 | 36.06 | 3 | 53.32011 | NA | NA | 565.40335 | NA | NA | NA | NA |

| Result | model | LogLik | AIC | AICc | ΔAICc | num_params | root_theta0 | theta_high | theta_low | sigma | sigma_high | sigma_low | alpha | half_life |
|---|---|---|---|---|---|---|---|---|---|---|---|---|---|---|
| NLEA | BM1 | -234.5406 | 473.0811 | 473.2954 | 0.00 | 2 | 14.95811 | NA | NA | 1.011076 | NA | NA | NA | NA |
| NLEA | BMM | -233.9768 | 473.9536 | 474.38 | 1.08 | 3 | 14.72139 | NA | NA | NA | 1.24E-10 | 1.852219 | NA | NA |
| NLEA | OU1 | -234.288 | 474.5761 | 475.0048 | 1.71 | 3 | 15.10998 | NA | NA | 4.302551 | NA | NA | 0.1328213 | 5.2191232 |
| NLEA | WN | -235.0156 | 476.0313 | 476.4638 | 3.17 | 3 | 15.08092 | NA | NA | 115.0359 | NA | NA | NA | NA |
| NLEA | OUM | -233.9733 | 475.9467 | 476.6729 | 3.38 | 4 | 14.07814 | 12.69079 | 16.31771 | 3.471675 | NA | NA | 1.4140594 | 0.4901825 |

| Result | model | LogLik | AIC | AICc | ΔAICc | num_params | root_theta0 | theta_high | theta_low | sigma | sigma_high | sigma_low | alpha | half_life |
|---|---|---|---|---|---|---|---|---|---|---|---|---|---|---|
| LA | BMM | -445.7384 | 897.4768 | 897.8802 | 0.00 | 3 | 381.0261 | NA | NA | NA | 1736.631 | 6584.507 | NA | NA |
| LA | BM1 | -447.6875 | 899.375 | 899.5718 | 1.69 | 2 | 412.9088 | NA | NA | 4404.365 | NA | NA | NA | NA |
| LA | OUM | -445.6428 | 899.2857 | 899.9753 | 2.10 | 4 | 315.2056 | 174.3254 | 556.719 | 8356.038 | NA | NA | 0.0647941 | 10.69769 |
| LA | OU1 | -447.7474 | 901.4949 | 901.9051 | 4.02 | 3 | 418.3806 | NA | NA | 7379.884 | NA | NA | 0.0426372 | 16.26436 |
| LA | WN | -465.4556 | 936.9112 | 937.3079 | 39.43 | 3 | 466.1906 | NA | NA | 129234.02 | NA | NA | NA | NA |

| Result | model | LogLik | AIC | AICc | ΔAICc | num_params | root_theta0 | theta_high | theta_low | sigma | sigma_high | sigma_low | alpha | half_life |
|---|---|---|---|---|---|---|---|---|---|---|---|---|---|---|
| SLA | OU1 | -219.1387 | 444.2774 | 444.6774 | 0.00 | 3 | 23.8297 | NA | NA | 8.121187 | NA | NA | 0.124598 | 5.563255 |
| SLA | OUM | -218.8241 | 445.6481 | 446.3371 | 1.66 | 4 | 24.00895 | 25.30733 | 22.9385 | 8.307619 | NA | NA | 0.1325828 | 5.228031 |
| SLA | BM1 | -221.7629 | 447.5258 | 447.7241 | 3.05 | 2 | 23.78233 | NA | NA | 2.350677 | NA | NA | NA | NA |
| SLA | BMM | -221.3584 | 448.7167 | 449.1217 | 4.44 | 3 | 23.67024 | NA | NA | NA | 2.593968 | 2.21786 | NA | NA |
| SLA | WN | -223.405 | 452.8099 | 453.2133 | 8.54 | 3 | 25.99099 | NA | NA | 61.439309 | NA | NA | NA | NA |

| Result | model | LogLik | AIC | AICc | ΔAICc | num_params | root_theta0 | theta_high | theta_low | sigma | sigma_high | sigma_low | alpha | half_life |
|---|---|---|---|---|---|---|---|---|---|---|---|---|---|---|
| LDMC | WN | -185.1792 | 376.3583 | 376.7722 | 0.00 | 3 | 19.00759 | NA | NA | 20.468561 | NA | NA | NA | NA |
| LDMC | OUM | -185.7658 | 379.5316 | 380.2234 | 3.45 | 4 | 17.00866 | 16.071 | 19.93343 | 9.503794 | NA | NA | 0.3563438 | 1.945164 |
| LDMC | OU1 | -187.8936 | 381.7872 | 382.2011 | 5.43 | 3 | 18.64541 | NA | NA | 6.620264 | NA | NA | 0.192894 | 3.595274 |
| LDMC | BMM | -189.9357 | 385.8714 | 386.2826 | 9.51 | 3 | 18.14165 | NA | NA | NA | 0.310857 | 1.750417 | NA | NA |

| Result | model | LogLik | AIC | AICc | ΔAICc | num_params | root_theta0 | theta_high | theta_low | sigma | sigma_high | sigma_low | alpha | half_life |
|---|---|---|---|---|---|---|---|---|---|---|---|---|---|---|
| LDMC | BM1 | -191.8504 | 387.7007 | 387.9026 | 11.13 | 2 | 18.73593 | NA | NA | 1.139782 | NA | NA | NA | NA |

| Result | model | LogLik | AIC | AICc | ΔAICc | num_params | root_theta0 | theta_high | theta_low | sigma | sigma_high | sigma_low | alpha | half_life |
|---|---|---|---|---|---|---|---|---|---|---|---|---|---|---|
| LTH | BM1 | 77.13259 | -150.2652 | -150.043 | 0.00 | 2 | 0.2754979 | NA | NA | 2.21E-05 | NA | NA | NA | NA |
| LTH | BMM | 77.63941 | -149.2788 | -148.8223 | 1.22 | 3 | 0.2734668 | NA | NA | NA | 7.59E-05 | 4.60E-13 | NA | NA |
| LTH | OU1 | 77.03313 | -148.0663 | -147.591 | 2.45 | 3 | 0.2749684 | NA | NA | 1.11E-04 | NA | NA | 0.0973316 | 7.12154 |
| LTH | OUM | 78.09724 | -148.1945 | -147.4103 | 2.63 | 4 | 0.2799208 | 0.2983985 | 0.2617695 | 9.21E-05 | NA | NA | 0.1165921 | 5.945062 |
| LTH | WN | 76.34497 | -146.6899 | -146.2413 | 3.80 | 3 | 0.2742119 | NA | NA | 3.47E-03 | NA | NA | NA | NA |

| Result | model | LogLik | AIC | AICc | ΔAICc | num_params | root_theta0 | theta_high | theta_low | sigma | sigma_high | sigma_low | alpha | half_life |
|---|---|---|---|---|---|---|---|---|---|---|---|---|---|---|
| LDI | BM1 | -39.4581 | 82.9162 | 83.11965 | 0.00 | 2 | 2.011905 | NA | NA | 0.0091909 | NA | NA | NA | NA |
| LDI | BMM | -38.85476 | 83.70952 | 84.113 | 0.99 | 3 | 1.979714 | NA | NA | NA | 0.0049016 | 0.0114898 | NA | NA |
| LDI | OU1 | -40.46224 | 86.92447 | 87.32458 | 4.20 | 3 | 1.980032 | NA | NA | 0.0135414 | NA | NA | 0.034729 | 20.01845 |
| LDI | OUM | -39.89674 | 87.79347 | 88.4818 | 5.36 | 4 | 1.953612 | 1.851168 | 2.016643 | 0.0139886 | NA | NA | 0.0390942 | 17.73016 |
| LDI | WN | -51.53685 | 109.0737 | 109.4771 | 26.36 | 3 | 2.008907 | NA | NA | 0.2835623 | NA | NA | NA | NA |

| Result | model | LogLik | AIC | AICc | ΔAICc | num_params | root_theta0 | theta_high | theta_low | sigma | sigma_high | sigma_low | alpha | half_life |
|---|---|---|---|---|---|---|---|---|---|---|---|---|---|---|
| RESmT1 | BM1 | -206.3503 | 416.7007 | 416.908 | 0.00 | 2 | 74.87469 | NA | NA | 0.4844531 | NA | NA | NA | NA |
| RESmT1 | BMM | -205.9395 | 417.8791 | 418.3036 | 1.40 | 3 | 74.8426 | NA | NA | NA | 1.003427 | 0.1832012 | NA | NA |
| RESmT1 | OU1 | -206.523 | 419.0461 | 419.4746 | 2.57 | 3 | 74.77912 | NA | NA | 1.8028053 | NA | NA | 0.0859286 | 8.06673 |
| RESmT1 | OUM | -205.9813 | 419.9626 | 420.6899 | 3.78 | 4 | 75.02304 | 76.21255 | 74.00905 | 1.7464161 | NA | NA | 0.090066 | 7.695991 |
| RESmT1 | WN | -207.8171 | 421.6342 | 422.0627 | 5.15 | 3 | 74.73304 | NA | NA | 50.876766 | NA | NA | NA | NA |

| Result | model | LogLik | AIC | AICc | ΔAICc | num_params | root_theta0 | theta_high | theta_low | sigma | sigma_high | sigma_low | alpha | half_life |
|---|---|---|---|---|---|---|---|---|---|---|---|---|---|---|
| RESmT2 | BM1 | -240.8091 | 485.6183 | 485.8251 | 0.00 | 2 | 61.45403 | NA | NA | 1.46E-15 | NA | NA | NA | NA |
| RESmT2 | WN | -240.7381 | 487.4762 | 487.901 | 2.08 | 3 | 61.57294 | NA | NA | 1.57E+02 | NA | NA | NA | NA |
| RESmT2 | OU1 | -240.7739 | 487.5478 | 487.9726 | 2.15 | 3 | 61.54905 | NA | NA | 4.38E+00 | NA | NA | 15.459472 | 0.0448808 |
| RESmT2 | BMM | -240.7808 | 487.5617 | 487.9865 | 2.16 | 3 | 61.52119 | NA | NA | NA | 1.66E-09 | 3.08E-11 | NA | NA |
| RESmT2 | OUM | -240.5685 | 489.137 | 489.8538 | 4.03 | 4 | 61.45508 | 61.15972 | 61.80885 | 8.19E-03 | NA | NA | 1.347892 | 0.5142453 |

| Result | model | LogLik | AIC | AICc | ΔAICc | num_params | root_theta0 | theta_high | theta_low | sigma | sigma_high | sigma_low | alpha | half_life |
|---|---|---|---|---|---|---|---|---|---|---|---|---|---|---|
| RESpT1 | BM1 | -226.3677 | 456.7354 | 456.9423 | 0.00 | 2 | 66.97155 | NA | NA | 0.609361 | NA | NA | NA | NA |
| RESpT1 | BMM | -225.701 | 457.402 | 457.8268 | 0.88 | 3 | 67.15271 | NA | NA | NA | 1.818953 | 1.69E-11 | NA | NA |
| RESpT1 | OU1 | -226.5135 | 459.0269 | 459.4481 | 2.51 | 3 | 66.99978 | NA | NA | 3.207346 | NA | NA | 0.1259797 | 5.50451 |
| RESpT1 | WN | -226.7529 | 459.5058 | 459.9345 | 2.99 | 3 | 66.97834 | NA | NA | 93.047862 | NA | NA | NA | NA |
| RESpT1 | OUM | -225.9382 | 459.8763 | 460.6036 | 3.66 | 4 | 66.95102 | 66.54454 | 67.48756 | 2.447321 | NA | NA | 0.1439117 | 4.816476 |

| Result | model | LogLik | AIC | AICc | ΔAICc | num_params | root_theta0 | theta_high | theta_low | sigma | sigma_high | sigma_low | alpha | half_life |
|---|---|---|---|---|---|---|---|---|---|---|---|---|---|---|
| RESpT2 | OUM | -204.7279 | 417.4558 | 418.1619 | 0.00 | 4 | 7.529571 | 6.564833 | 11.96549 | 3.05E-06 | NA | NA | 5.8041063 | 0.1194237 |
| RESpT2 | BM1 | -207.1167 | 418.2334 | 418.4421 | 0.28 | 2 | 10.738474 | NA | NA | 6.15E-01 | NA | NA | NA | NA |
| RESpT2 | BMM | -206.6071 | 419.2141 | 419.6411 | 1.48 | 3 | 10.191173 | NA | NA | NA | 3.40E-11 | 0.8361094 | NA | NA |
| RESpT2 | OU1 | -207.6178 | 421.2356 | 421.6604 | 3.50 | 3 | 10.263111 | NA | NA | 1.52E+00 | NA | NA | 0.0858421 | 8.0748794 |
| RESpT2 | WN | -208.2611 | 422.5222 | 422.9437 | 4.78 | 3 | 10.024294 | NA | NA | 4.96E+01 | NA | NA | NA | NA |

#### Heat (bootstrapped phylogeny)

| **Result** | **model** | **LogLik** | **AIC** | **AICc** | **ΔAICc** | **num_params** | **root_theta0** | **theta_high** | **theta_low** | **sigma** | **sigma_high** | **sigma_low** | **alpha** | **half_life** |
|---|---|---|---|---|---|---|---|---|---|---|---|---|---|---|
| IGR | BM1 | 74.16213 | -144.3243 | -144.1119 | 0.00 | 2 | 0.5721737 | NA | NA | 5.82E-24 | NA | NA | NA | NA |
| IGR | WN | 74.19817 | -142.3963 | -141.9639 | 2.15 | 3 | 0.5721737 | NA | NA | 4.29E-03 | NA | NA | NA | NA |
| IGR | BMM | 74.164 | -142.328 | -141.8919 | 2.22 | 3 | 0.5724156 | NA | NA | NA | 4.22E-17 | 1.00E-16 | NA | NA |
| IGR | OU1 | 74.16213 | -142.3243 | -141.8918 | 2.22 | 3 | 0.5721737 | NA | NA | 2.08E-21 | NA | NA | 0.09339962 | 7.421306 |
| IGR | OUM | 74.42052 | -140.841 | -140.1025 | 4.01 | 4 | 0.5728028 | 0.5718358 | 0.5728255 | 1.72E-13 | NA | NA | 0.09340315 | 7.421026 |

| **Result** | **model** | **LogLik** | **AIC** | **AICc** | **ΔAICc** | **num_params** | **root_theta0** | **theta_high** | **theta_low** | **sigma** | **sigma_high** | **sigma_low** | **alpha** | **half_life** |
|---|---|---|---|---|---|---|---|---|---|---|---|---|---|---|
| MGR | OU1 | -25.96633 | 57.93265 | 58.34294 | 0.00 | 3 | 0.3691191 | NA | NA | 0.014647056 | NA | NA | 0.2850325 | 2.432062 |
| MGR | OUM | -25.61975 | 59.2395 | 59.92916 | 1.59 | 4 | 0.378012 | 0.4171866 | 0.3367523 | 0.015614694 | NA | NA | 0.2909374 | 2.382462 |
| MGR | BMM | -28.32663 | 62.65325 | 63.06354 | 4.72 | 3 | 0.3464606 | NA | NA | NA | 0.000314802 | 0.002307534 | NA | NA |
| MGR | BM1 | -29.51623 | 63.03247 | 63.23592 | 4.89 | 2 | 0.3691001 | NA | NA | 0.001767268 | NA | NA | NA | NA |
| MGR | WN | -29.31426 | 64.62853 | 65.03881 | 6.70 | 3 | 0.5404512 | NA | NA | 0.119555461 | NA | NA | NA | NA |

| **Result** | **model** | **LogLik** | **AIC** | **AICc** | **ΔAICc** | **num_params** | **root_theta0** | **theta_high** | **theta_low** | **sigma** | **sigma_high** | **sigma_low** | **alpha** | **half_life** |
|---|---|---|---|---|---|---|---|---|---|---|---|---|---|---|
| XMID | WN | -176.7067 | 359.4135 | 359.8238 | 0.00 | 3 | 23.66996 | NA | NA | 16.5486763 | NA | NA | NA | NA |
| XMID | OUM | -180.7213 | 369.4426 | 370.1447 | 10.32 | 4 | 22.51808 | 21.78934 | 23.58541 | 27.342628 | NA | NA | 6.3070883 | 0.1098997 |
| XMID | OU1 | -182.3585 | 370.7171 | 371.1345 | 11.31 | 3 | 22.67566 | NA | NA | 5.702002 | NA | NA | 0.8318017 | 0.833333 |
| XMID | BM1 | -183.6424 | 371.2848 | 371.4865 | 11.66 | 2 | 22.42437 | NA | NA | 0.1824042 | NA | NA | NA | NA |
| XMID | BMM | -183.1724 | 372.3447 | 372.7585 | 12.93 | 3 | 22.52074 | NA | NA | NA | 0.2600501 | 0.03882942 | NA | NA |

| **Result** | **model** | **LogLik** | **AIC** | **AICc** | **ΔAICc** | **num_params** | **root_theta0** | **theta_high** | **theta_low** | **sigma** | **sigma_high** | **sigma_low** | **alpha** | **half_life** |
|---|---|---|---|---|---|---|---|---|---|---|---|---|---|---|
| ASYM | OUM | -278.785 | 565.57 | 566.2665 | 0.00 | 4 | 37.68803 | 23.17763 | 65.3371 | 52.39409 | NA | NA | 0.05911481 | 11.72544 |
| ASYM | BMM | -279.9563 | 565.9125 | 566.3276 | 0.06 | 3 | 46.13897 | NA | NA | NA | 16.22286 | 37.1929 | NA | NA |
| ASYM | BM1 | -281.1326 | 566.2653 | 566.467 | 0.20 | 2 | 48.46633 | NA | NA | 29.51575 | NA | NA | NA | NA |
| ASYM | OU1 | -282.198 | 570.396 | 570.8028 | 4.54 | 3 | 48.81529 | NA | NA | 43.32991 | NA | NA | 0.02918887 | 23.74709 |
| ASYM | WN | -292.0697 | 590.1394 | 590.5497 | 24.28 | 3 | 48.80384 | NA | NA | 676.50096 | NA | NA | NA | NA |

| **Result** | **model** | **LogLik** | **AIC** | **AICc** | **ΔAICc** | **num_params** | **root_theta0** | **theta_high** | **theta_low** | **sigma** | **sigma_high** | **sigma_low** | **alpha** | **half_life** |
|---|---|---|---|---|---|---|---|---|---|---|---|---|---|---|
| NLEA | BM1 | -223.4508 | 450.9016 | 451.1218 | 0.00 | 2 | 14.36814 | NA | NA | 0.5590851 | NA | NA | NA | NA |
| NLEA | BMM | -222.8278 | 451.6557 | 452.0986 | 0.98 | 3 | 14.11951 | NA | NA | NA | 1.28E-10 | 0.9727474 | NA | NA |
| NLEA | OU1 | -223.1594 | 452.3188 | 452.7554 | 1.63 | 3 | 14.47276 | NA | NA | 5.6963527 | NA | NA | 0.1318141 | 5.258539 |
| NLEA | WN | -223.2271 | 452.4542 | 452.9029 | 1.78 | 3 | 14.58343 | NA | NA | 83.273899 | NA | NA | NA | NA |
| NLEA | OUM | -222.5743 | 453.1485 | 453.8996 | 2.78 | 4 | 13.75 | 12.6329 | 15.72531 | 6.8024292 | NA | NA | 0.3287143 | 2.108661 |

| **Result** | **model** | **LogLik** | **AIC** | **AICc** | **ΔAICc** | **num_params** | **root_theta0** | **theta_high** | **theta_low** | **sigma** | **sigma_high** | **sigma_low** | **alpha** | **half_life** |
|---|---|---|---|---|---|---|---|---|---|---|---|---|---|---|
| LA | BMM | -424.3892 | 854.7784 | 855.2109 | 0.00 | 3 | 346.2086 | NA | NA | NA | 1507.324 | 7235.175 | NA | NA |
| LA | BM1 | -426.2072 | 856.4145 | 856.6214 | 1.41 | 2 | 389.5389 | NA | NA | 5005.154 | NA | NA | NA | NA |
| LA | OUM | -426.1399 | 860.2798 | 860.9931 | 5.78 | 4 | 317.7456 | 107.1259 | 588.9605 | 6455.264 | NA | NA | 0.02945695 | 23.53086 |
| LA | OU1 | -427.5524 | 861.1048 | 861.5223 | 6.31 | 3 | 390.2264 | NA | NA | 6403.324 | NA | NA | 0.02038059 | 34.02562 |
| LA | WN | -436.5248 | 879.0496 | 879.4781 | 24.27 | 3 | 400.9644 | NA | NA | 114908.153 | NA | NA | NA | NA |

| **Result** | **model** | **LogLik** | **AIC** | **AICc** | **ΔAICc** | **num_params** | **root_theta0** | **theta_high** | **theta_low** | **sigma** | **sigma_high** | **sigma_low** | **alpha** | **half_life** |
|---|---|---|---|---|---|---|---|---|---|---|---|---|---|---|
| SLA | OU1 | -208.693 | 423.386 | 423.8146 | 0.00 | 3 | 26.5804 | NA | NA | 8.11543 | NA | NA | 0.1170439 | 5.922777 |
| SLA | OUM | -208.1042 | 424.2085 | 424.9358 | 1.12 | 4 | 27.07384 | 28.93464 | 25.30845 | 8.386539 | NA | NA | 0.1281949 | 5.40698 |

| | | | | | | | | | | | | | | |
|---|---|---|---|---|---|---|---|---|---|---|---|---|---|---|
| SLA | BMM | -210.1071 | 426.2141 | 426.6484 | 2.83 | 3 | 25.56737 | NA | NA | NA | 3.735193 | 1.43889 | NA | NA |
| SLA | WN | -210.4397 | 426.8793 | 427.3157 | 3.50 | 3 | 29.19803 | NA | NA | 60.557698 | NA | NA | NA | NA |
| SLA | BM1 | -211.8498 | 427.6995 | 427.91 | 4.10 | 2 | 25.95833 | NA | NA | 2.528676 | NA | NA | NA | NA |

| **Result** | **model** | **LogLik** | **AIC** | **AICc** | **ΔAICc** | **num_params** | **root_theta0** | **theta_high** | **theta_low** | **sigma** | **sigma_high** | **sigma_low** | **alpha** | **half_life** |
|---|---|---|---|---|---|---|---|---|---|---|---|---|---|---|
| LDMC | BMM | -163.6049 | 333.2099 | 333.6528 | 0.00 | 3 | 15.43553 | NA | NA | NA | 0.03662674 | 0.6139352 | NA | NA |
| LDMC | BM1 | -164.835 | 333.67 | 333.8882 | 0.24 | 2 | 15.85599 | NA | NA | 0.4379742 | NA | NA | NA | NA |
| LDMC | OUM | -163.3566 | 334.7132 | 335.4569 | 1.80 | 4 | 14.04467 | 12.63153 | 17.16862 | 1.1741601 | NA | NA | 0.09710076 | 7.138432 |
| LDMC | OU1 | -165.7332 | 337.4665 | 337.8956 | 4.24 | 3 | 15.77801 | NA | NA | 0.7870584 | NA | NA | 0.03622046 | 19.136899 |
| LDMC | WN | -173.4465 | 352.8929 | 353.3216 | 19.67 | 3 | 15.58739 | NA | NA | 17.5226803 | NA | NA | NA | NA |

| **Result** | **model** | **LogLik** | **AIC** | **AICc** | **ΔAICc** | **num_params** | **root_theta0** | **theta_high** | **theta_low** | **sigma** | **sigma_high** | **sigma_low** | **alpha** | **half_life** |
|---|---|---|---|---|---|---|---|---|---|---|---|---|---|---|
| LTH | BM1 | 85.92467 | -167.8493 | -167.6331 | 0.00 | 2 | 0.2660282 | NA | NA | 1.09E-06 | NA | NA | NA | NA |
| LTH | BMM | 86.49433 | -166.9887 | -166.5181 | 1.12 | 3 | 0.2647142 | NA | NA | NA | 4.57E-06 | 6.26E-14 | NA | NA |
| LTH | OU1 | 85.91156 | -165.8231 | -165.3745 | 2.26 | 3 | 0.2667098 | NA | NA | 1.11E-05 | NA | NA | 0.09282081 | 7.467584 |
| LTH | WN | 85.59336 | -165.1867 | -164.7463 | 2.89 | 3 | 0.2667548 | NA | NA | 2.47E-03 | NA | NA | NA | NA |
| LTH | OUM | 86.67114 | -165.3423 | -164.5686 | 3.06 | 4 | 0.2697478 | 0.2818866 | 0.2570028 | 6.56E-13 | NA | NA | 0.07283875 | 9.516187 |

| **Result** | **model** | **LogLik** | **AIC** | **AICc** | **ΔAICc** | **num_params** | **root_theta0** | **theta_high** | **theta_low** | **sigma** | **sigma_high** | **sigma_low** | **alpha** | **half_life** |
|---|---|---|---|---|---|---|---|---|---|---|---|---|---|---|
| LDI | BMM | -42.7938 | 91.58759 | 92.00507 | 0.00 | 3 | 1.881344 | NA | NA | NA | 0.004546867 | 0.01890251 | NA | NA |
| LDI | BM1 | -45.52559 | 95.05118 | 95.26741 | 3.26 | 2 | 1.935864 | NA | NA | 0.01360567 | NA | NA | NA | NA |
| LDI | OUM | -43.69718 | 95.39435 | 96.12513 | 4.12 | 4 | 1.846136 | 1.579857 | 2.070832 | 0.0288473 | NA | NA | 0.07747442 | 8.946787 |
| LDI | OU1 | -45.4832 | 96.96639 | 97.39497 | 5.39 | 3 | 1.913292 | NA | NA | 0.02482853 | NA | NA | 0.05591763 | 12.396677 |
| LDI | WN | -53.6634 | 113.3268 | 113.76331 | 21.76 | 3 | 1.930259 | NA | NA | 0.30376486 | NA | NA | NA | NA |

| **Result** | **model** | **LogLik** | **AIC** | **AICc** | **ΔAICc** | **num_params** | **root_theta0** | **theta_high** | **theta_low** | **sigma** | **sigma_high** | **sigma_low** | **alpha** | **half_life** |
|---|---|---|---|---|---|---|---|---|---|---|---|---|---|---|
| RESpT1 | BM1 | -125.3266 | 254.6533 | 255.0819 | 0.00 | 2 | 56.88129 | NA | NA | 3.34E-16 | NA | NA | NA | NA |
| RESpT1 | BMM | -125.0882 | 256.1764 | 257.0653 | 1.98 | 3 | 56.91484 | NA | NA | NA | 5.16E-10 | 1.26E-11 | NA | NA |
| RESpT1 | WN | -125.3621 | 256.7242 | 257.5972 | 2.52 | 3 | 56.94373 | NA | NA | 1.48E+02 | NA | NA | NA | NA |
| RESpT1 | OU1 | -125.4047 | 256.8093 | 257.6982 | 2.62 | 3 | 56.84541 | NA | NA | 4.09E-07 | NA | NA | 11.76126 | 0.05997688 |
| RESpT1 | OUM | -124.6496 | 257.2992 | 258.8366 | 3.75 | 4 | 56.54273 | 54.28846 | 58.30374 | 1.51E-05 | NA | NA | 14.9652 | 0.04631726 |

| **Result** | **model** | **LogLik** | **AIC** | **AICc** | **ΔAICc** | **num_params** | **root_theta0** | **theta_high** | **theta_low** | **sigma** | **sigma_high** | **sigma_low** | **alpha** | **half_life** |
|---|---|---|---|---|---|---|---|---|---|---|---|---|---|---|
| RESpT2 | OUM | -241.8009 | 491.6019 | 492.3291 | 0.00 | 4 | 17.30478 | 21.45835 | 13.13952 | 120.360087 | NA | NA | 0.6028516 | 1.149781 |
| RESpT2 | OU1 | -243.1017 | 492.2034 | 492.632 | 0.30 | 3 | 15.73253 | NA | NA | 63.145076 | NA | NA | 0.2886076 | 2.402099 |
| RESpT2 | WN | -243.2217 | 492.4435 | 492.872 | 0.54 | 3 | 18.64537 | NA | NA | 181.164602 | NA | NA | NA | NA |
| RESpT2 | BMM | -246.5566 | 499.1133 | 499.5418 | 7.21 | 3 | 15.26984 | NA | NA | NA | 14.52894 | 5.843217 | NA | NA |
| RESpT2 | BM1 | -247.9426 | 499.8853 | 500.0977 | 7.77 | 2 | 15.49823 | NA | NA | 7.949461 | NA | NA | NA | NA |

| **Result** | **model** | **LogLik** | **AIC** | **AICc** | **ΔAICc** | **num_params** | **root_theta0** | **theta_high** | **theta_low** | **sigma** | **sigma_high** | **sigma_low** | **alpha** | **half_life** |
|---|---|---|---|---|---|---|---|---|---|---|---|---|---|---|
| TOL_IGR | BM1 | 19.33953 | -34.67905 | -34.43896 | 0.00 | 2 | 0.0222101 | NA | NA | 1.52E-23 | NA | NA | NA | NA |
| TOL_IGR | OU1 | 19.53562 | -33.07124 | -32.57592 | 1.86 | 3 | 0.02204595 | NA | NA | 1.79E-17 | NA | NA | 0.1021202 | 6.787559 |
| TOL_IGR | BMM | 19.42989 | -32.85978 | -32.34968 | 2.09 | 3 | 0.02184819 | NA | NA | NA | 3.80E-16 | 8.95E-17 | NA | NA |
| TOL_IGR | WN | 19.33443 | -32.66887 | -32.16355 | 2.28 | 3 | 0.0222101 | NA | NA | 2.44E-02 | NA | NA | NA | NA |
| TOL_IGR | OUM | 19.84564 | -31.69128 | -30.83306 | 3.61 | 4 | 0.02175332 | 0.01898009 | 0.0249382 | 3.94E-13 | NA | NA | 0.1766803 | 3.923173 |

| **Result** | **model** | **LogLik** | **AIC** | **AICc** | **ΔAICc** | **num_params** | **root_theta0** | **theta_high** | **theta_low** | **sigma** | **sigma_high** | **sigma_low** | **alpha** | **half_life** |
|---|---|---|---|---|---|---|---|---|---|---|---|---|---|---|
| TOL_MGR | OUM | -107.9596 | 223.9192 | 224.7611 | 0.00 | 4 | 1.950833 | 2.48088 | 1.462942 | 7.742923 | NA | NA | 1.0246764 | 0.6764547 |
| TOL_MGR | OU1 | -109.8988 | 225.7977 | 226.2875 | 1.53 | 3 | 1.832108 | NA | NA | 7.734149 | NA | NA | 0.9370025 | 0.7397668 |
| TOL_MGR | WN | -110.5337 | 227.0673 | 227.5673 | 2.81 | 3 | 1.815046 | NA | NA | 3.40601 | NA | NA | NA | NA |
| TOL_MGR | BMM | -124.3463 | 254.6926 | 255.1796 | 30.42 | 3 | 1.596058 | NA | NA | NA | 1.959575 | 0.3721212 | NA | NA |
| TOL_MGR | BM1 | -133.2693 | 270.5386 | 270.7861 | 46.03 | 2 | 1.775863 | NA | NA | 1.100106 | NA | NA | NA | NA |

| **Result** | **model** | **LogLik** | **AIC** | **AICc** | **ΔAICc** | **num_params** | **root_theta0** | **theta_high** | **theta_low** | **sigma** | **sigma_high** | **sigma_low** | **alpha** | **half_life** |
|---|---|---|---|---|---|---|---|---|---|---|---|---|---|---|
| TOL_XMID | OUM | 17.211843 | -26.423686 | -25.599749 | 0.00 | 4 | -0.067293709 | -0.1086832 | -0.001932061 | 0.030737367 | NA | NA | 0.4825415 | 1.436451 |
| TOL_XMID | OU1 | 15.317504 | -24.635008 | -24.12924 | 1.47 | 3 | -0.03815613 | NA | NA | 0.028061768 | NA | NA | 0.3938685 | 1.759894 |
| TOL_XMID | WN | 14.336072 | -22.672144 | -22.19685 | 3.40 | 3 | -0.040427925 | NA | NA | 0.030583066 | NA | NA | NA | NA |
| TOL_XMID | BMM | 4.820712 | -3.641425 | -3.159893 | 22.44 | 3 | -0.007770368 | NA | NA | NA | 0.00753307 | 0.00495679 | NA | NA |
| TOL_XMID | BM1 | 3.637054 | -3.274109 | -3.02666 | 22.57 | 2 | -0.01002137 | NA | NA | 0.005882829 | NA | NA | NA | NA |

| **Result** | **model** | **LogLik** | **AIC** | **AICc** | **ΔAICc** | **num_params** | **root_theta0** | **theta_high** | **theta_low** | **sigma** | **sigma_high** | **sigma_low** | **alpha** | **half_life** |
|---|---|---|---|---|---|---|---|---|---|---|---|---|---|---|
| TOL_ASYM | OUM | -77.88186 | 163.7637 | 164.6198 | 0.00 | 4 | -0.2027545 | -0.3024569 | 0.7805902 | 2.2320002 | NA | NA | 0.9053213 | 0.7656366 |
| TOL_ASYM | OU1 | -82.48624 | 170.9725 | 171.4678 | 6.85 | 3 | 0.4273521 | NA | NA | 1.7952899 | NA | NA | 0.6642044 | 1.0435764 |
| TOL_ASYM | WN | -83.83223 | 173.6645 | 174.1606 | 9.54 | 3 | 0.4214921 | NA | NA | 1.1072331 | NA | NA | NA | NA |
| TOL_ASYM | BMM | -99.16004 | 204.3201 | 204.8201 | 40.20 | 3 | 0.2792002 | NA | NA | NA | 0.1543777 | 0.4141789 | NA | NA |
| TOL_ASYM | BM1 | -103.33586 | 210.6717 | 210.9142 | 46.29 | 2 | 0.404779 | NA | NA | 0.3270392 | NA | NA | NA | NA |

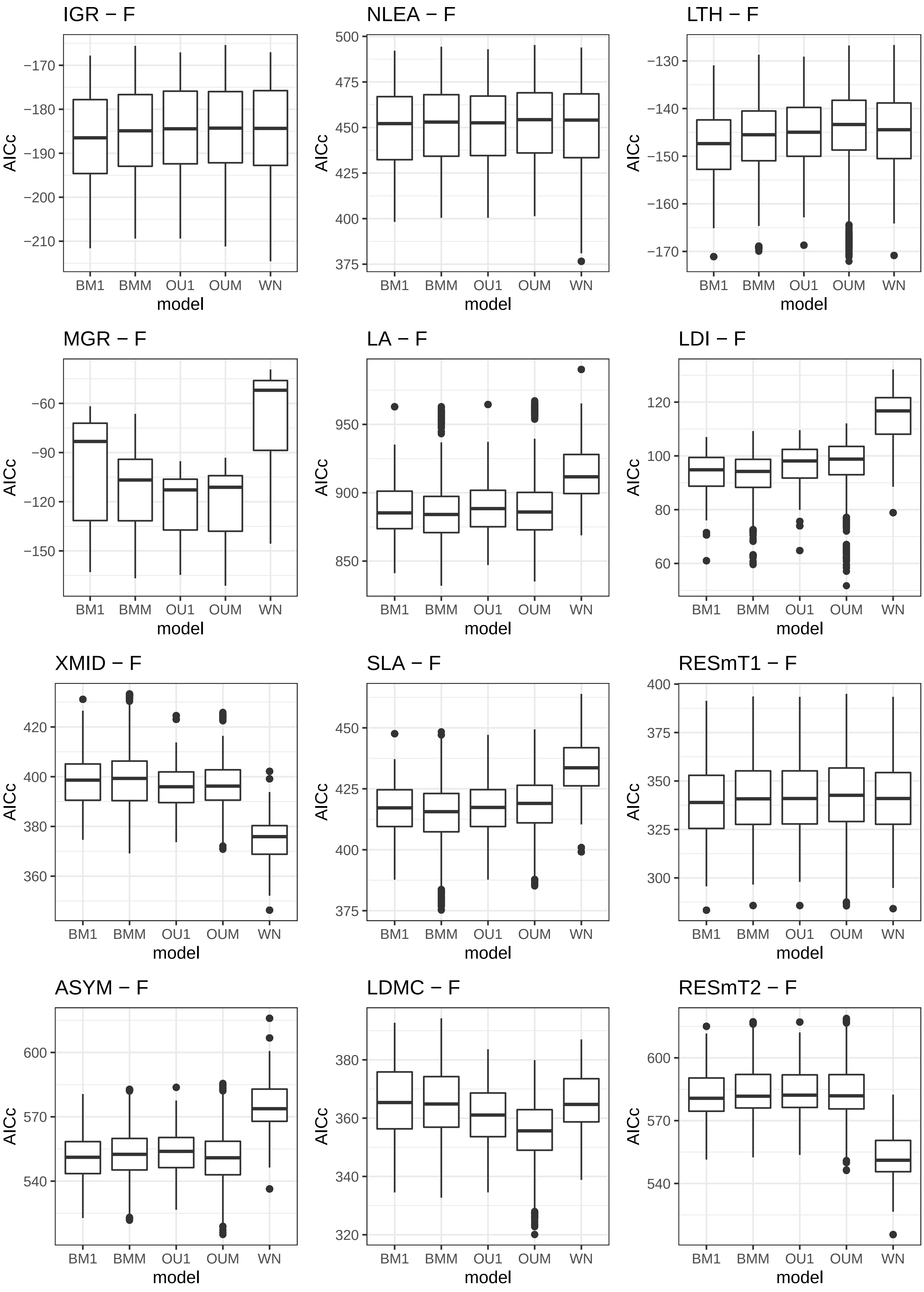
IGR − F
NLEA − F
LTH − F
MGR − F
LA − F
LDI − F
XMID − F
SLA − F
RESmT1 − F
ASYM − F
LDMC − F
RESmT2 − F
AICc
model
BM1
BMM
OU1
OUM
WN

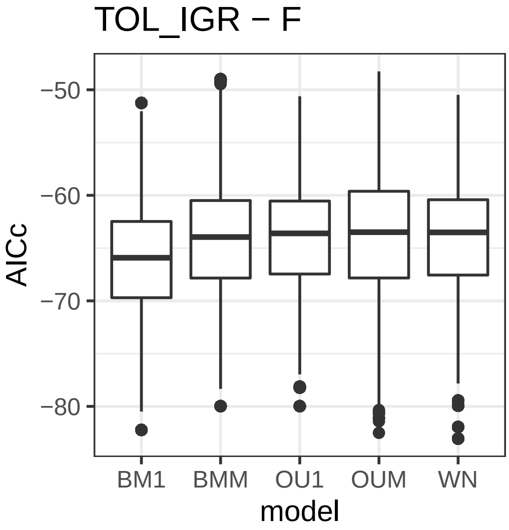
TOL_IGR − F
AICc
−50
−60
−70
−80
BM1
BMM
OU1
OUM
WN
model

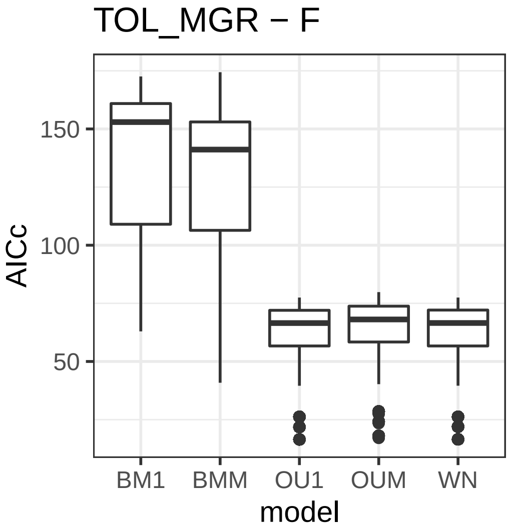
TOL_MGR − F
AICc
150
100
50
BM1
BMM
OU1
OUM
WN
model

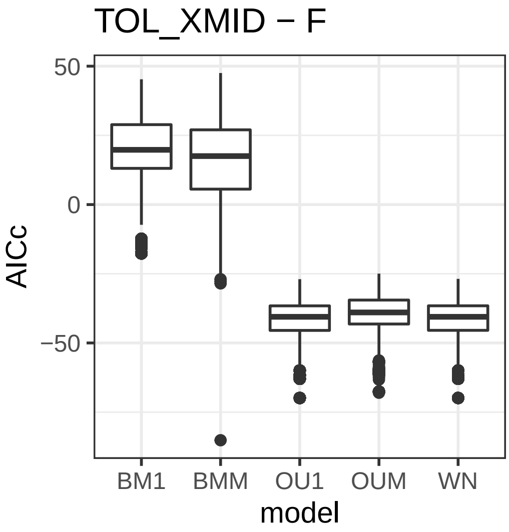
TOL_XMID − F
AICc
50
0
−50
BM1
BMM
OU1
OUM
WN
model

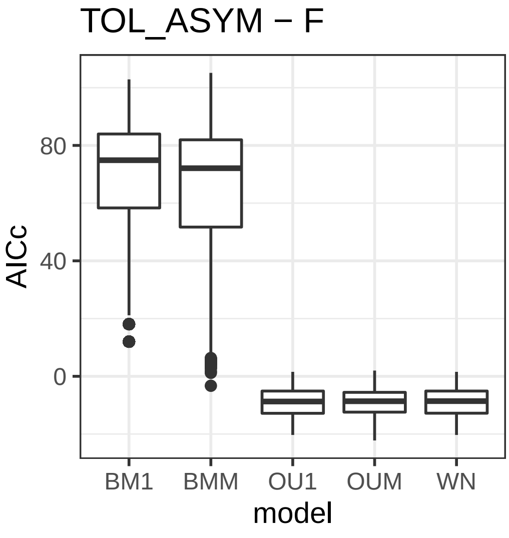
TOL_ASYM − F
AICc
80
40
0
BM1
BMM
OU1
OUM
WN
model

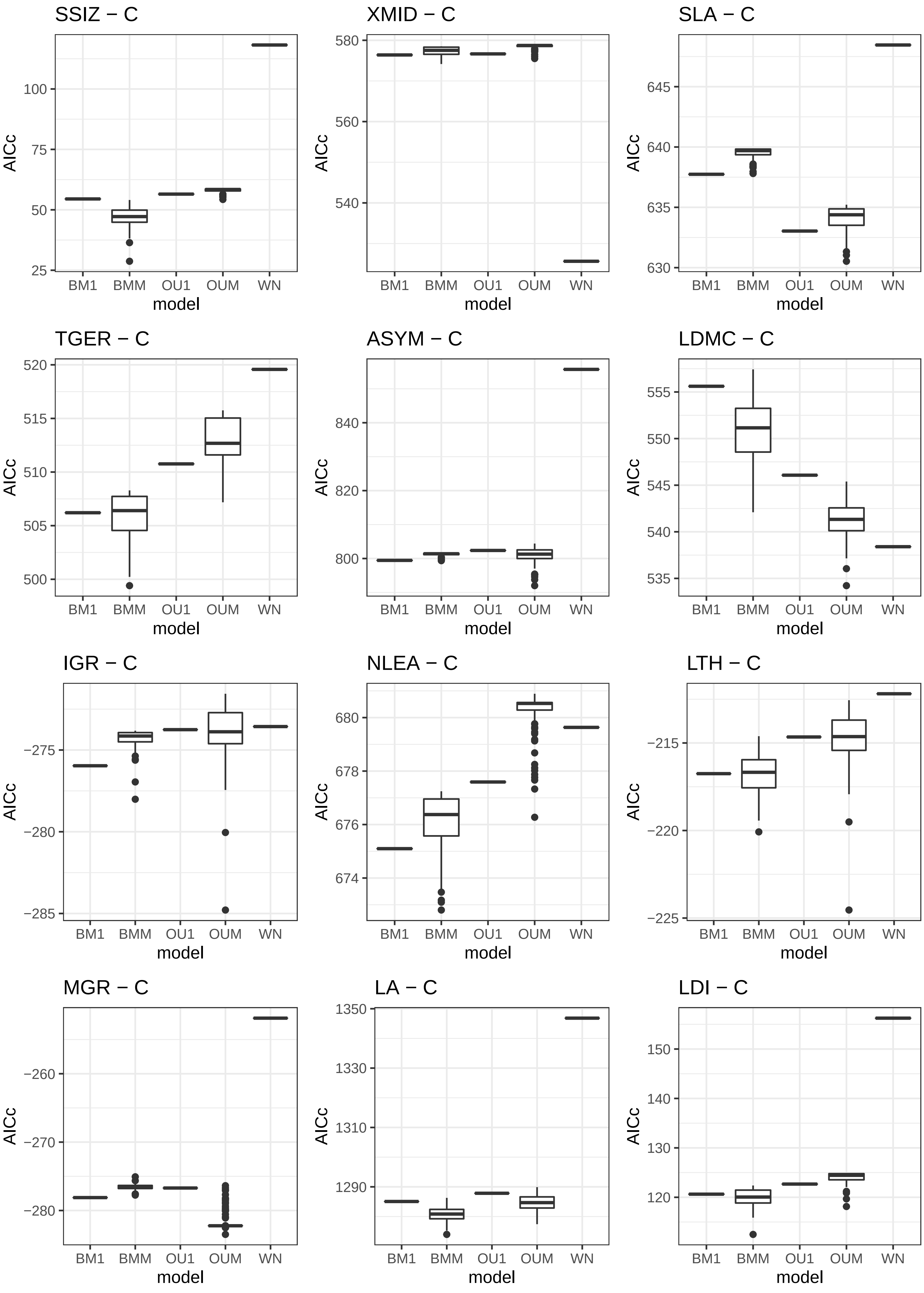
SSIZ − C
100
75
50
25
AICc
BM1 BMM OU1 OUM WN
model
XMID − C
580
560
540
AICc
BM1 BMM OU1 OUM WN
model
SLA − C
645
640
635
630
AICc
BM1 BMM OU1 OUM WN
model
TGER − C
520
515
510
505
500
AICc
BM1 BMM OU1 OUM WN
model
ASYM − C
840
820
800
AICc
BM1 BMM OU1 OUM WN
model
LDMC − C
555
550
545
540
535
AICc
BM1 BMM OU1 OUM WN
model
IGR − C
−275
−280
−285
AICc
BM1 BMM OU1 OUM WN
model
NLEA − C
680
678
676
674
AICc
BM1 BMM OU1 OUM WN
model
LTH − C
−215
−220
−225
AICc
BM1 BMM OU1 OUM WN
model
MGR − C
−260
−270
−280
AICc
BM1 BMM OU1 OUM WN
model
LA − C
1350
1330
1310
1290
AICc
BM1 BMM OU1 OUM WN
model
LDI − C
150
140
130
120
AICc
BM1 BMM OU1 OUM WN
model

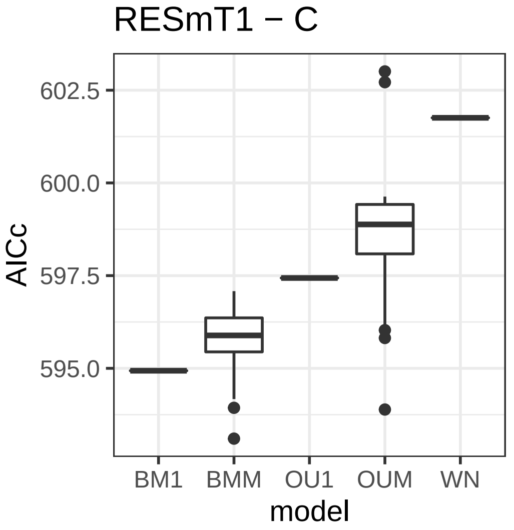
RESmT1 − C
AICc
602.5
600.0
597.5
595.0
BM1
BMM
OU1
OUM
WN
model

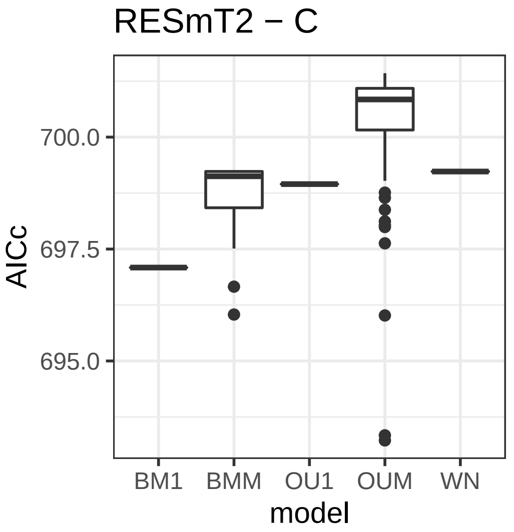
RESmT2 − C
AICc
700.0
697.5
695.0
BM1
BMM
OU1
OUM
WN
model

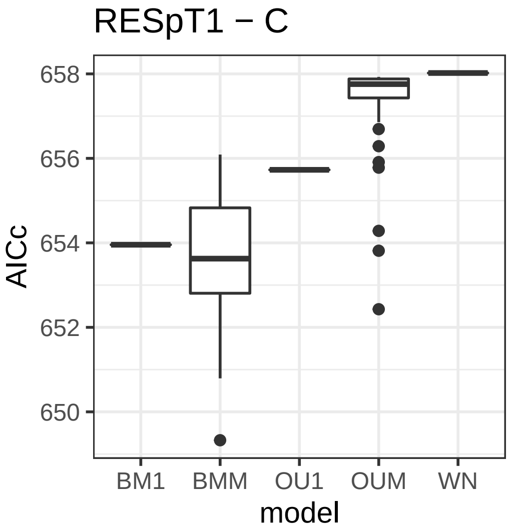
RESpT1 − C
AICc
658
656
654
652
650
BM1
BMM
OU1
OUM
WN
model

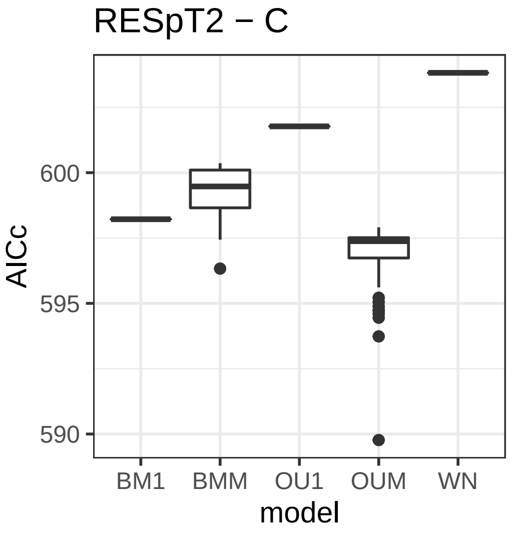
RESpT2 − C
AICc
600
595
590
BM1
BMM
OU1
OUM
WN
model

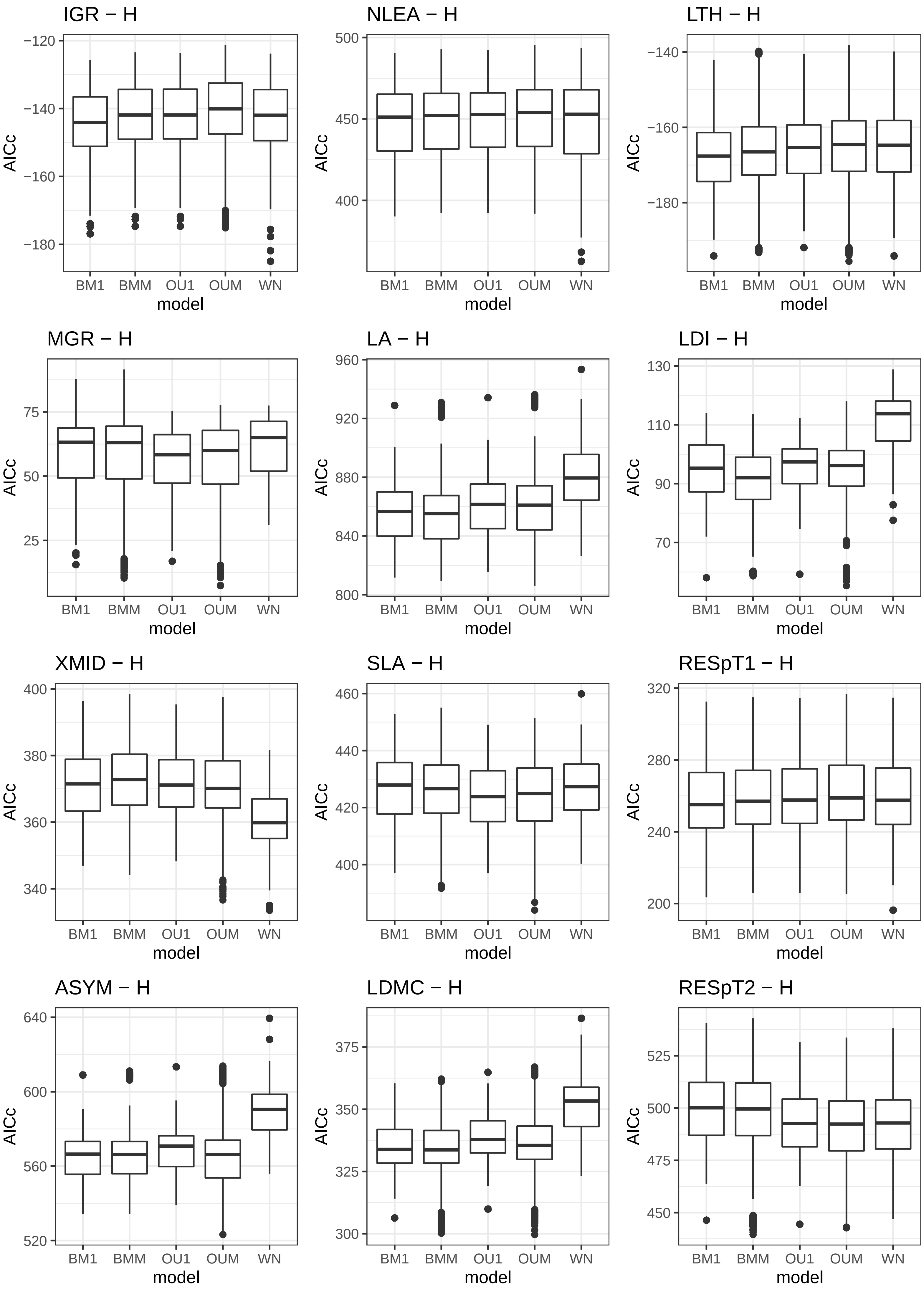
IGR − H
NLEA − H
LTH − H
MGR − H
LA − H
LDI − H
XMID − H
SLA − H
RESpT1 − H
ASYM − H
LDMC − H
RESpT2 − H
AICc
model
BM1
BMM
OU1
OUM
WN
−120
−140
−160
−180
500
450
400
960
920
880
840
800
75
50
25
130
110
90
70
400
380
360
340
460
440
420
320
280
240
200
640
600
560
520
375
350
325
300
525
500
475
450

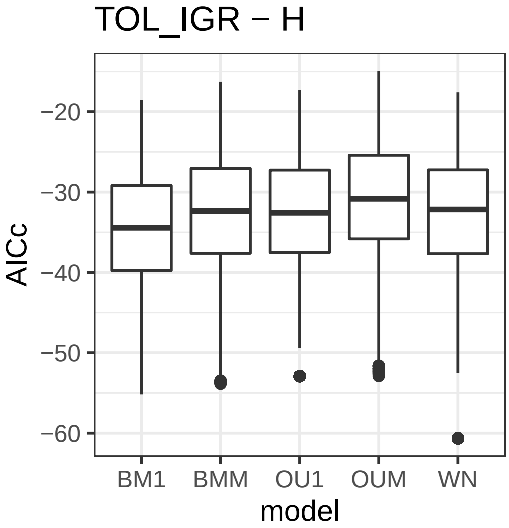
TOL_IGR − H
AICc
−20
−30
−40
−50
−60
BM1
BMM
OU1
OUM
WN
model

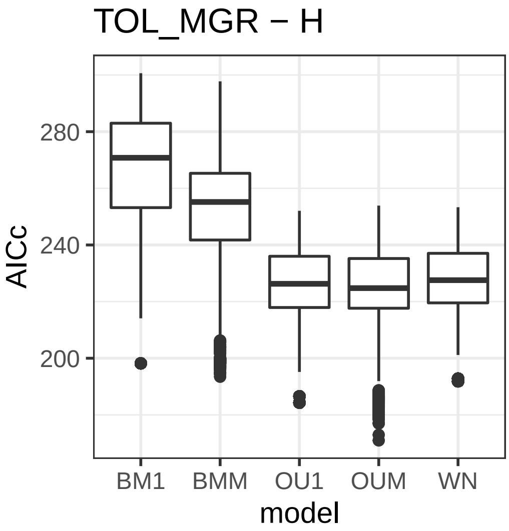
TOL_MGR − H
AICc
280
240
200
BM1
BMM
OU1
OUM
WN
model

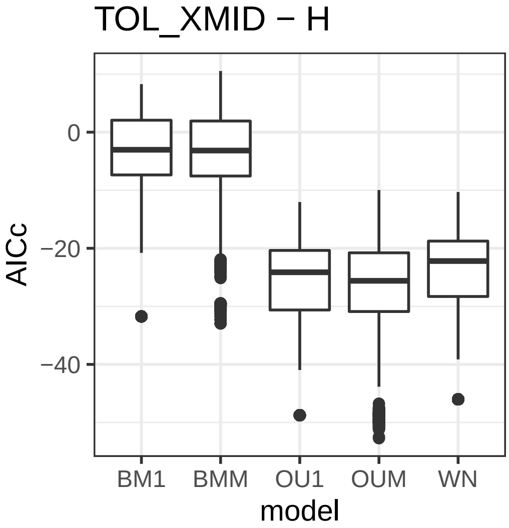
TOL_XMID − H
AICc
0
−20
−40
BM1
BMM
OU1
OUM
WN
model

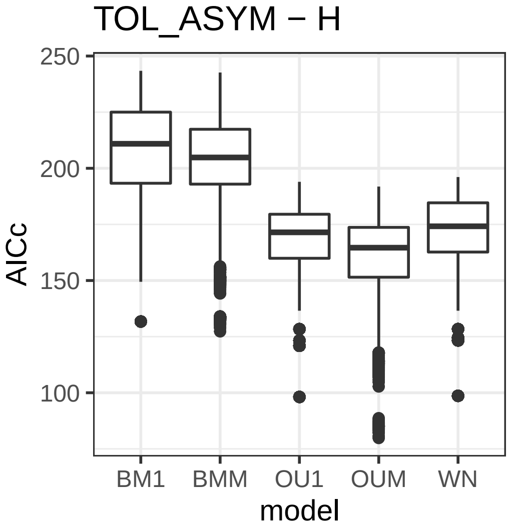
TOL_ASYM − H
AICc
250
200
150
100
BM1
BMM
OU1
OUM
WN
model

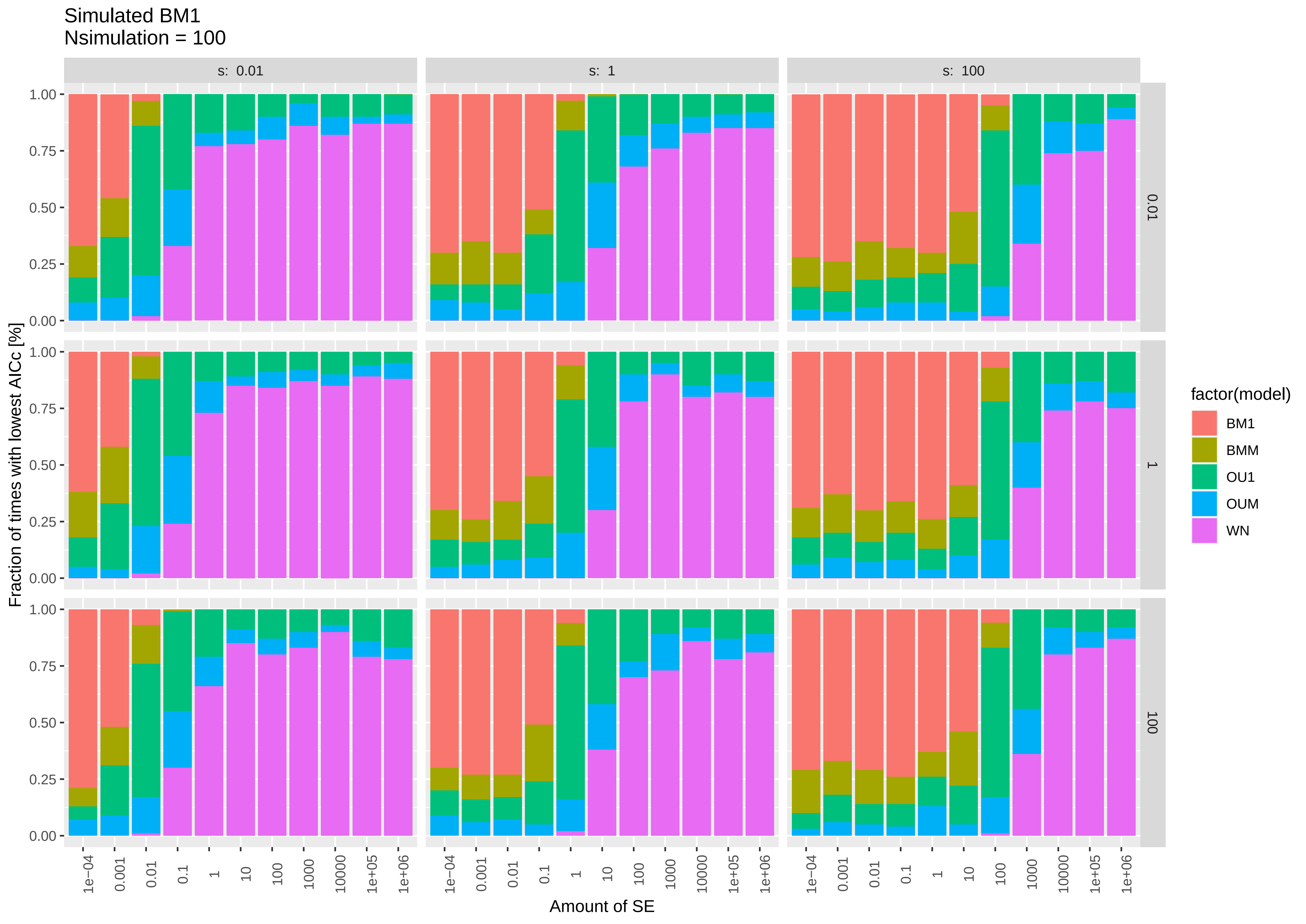

Simulated BM1
Nsimulation = 100
s: 0.01
s: 1
s: 100
0.01
1
100
Fraction of times with lowest AICc [%]
0.00
0.25
0.50
0.75
1.00
1e−04
0.001
0.01
0.1
1
10
100
1000
10000
1e+05
1e+06
Amount of SE
factor(model)
BM1
BMM
OU1
OUM
WN

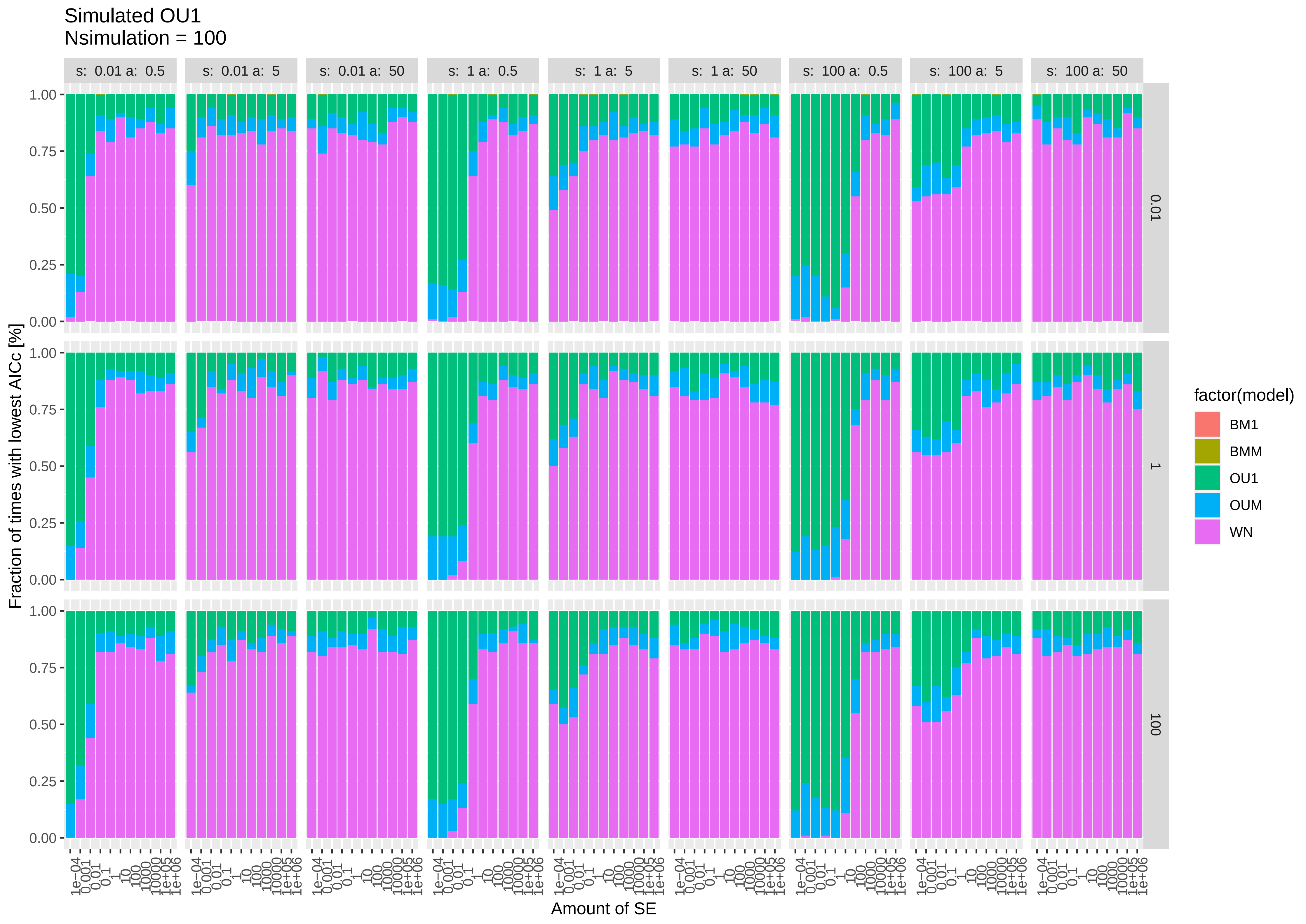

Simulated OU1
Nsimulation = 100
s: 0.01 a: 0.5
s: 0.01 a: 5
s: 0.01 a: 50
s: 1 a: 0.5
s: 1 a: 5
s: 1 a: 50
s: 100 a: 0.5
s: 100 a: 5
s: 100 a: 50
0.01
1
100
Fraction of times with lowest AICc [%]
0.00
0.25
0.50
0.75
1.00
1e−04
0.001
0.01
0.1
1
10
100
1000
10000
1e+05
1e+06
Amount of SE
factor(model)
BM1
BMM
OU1
OUM
WN

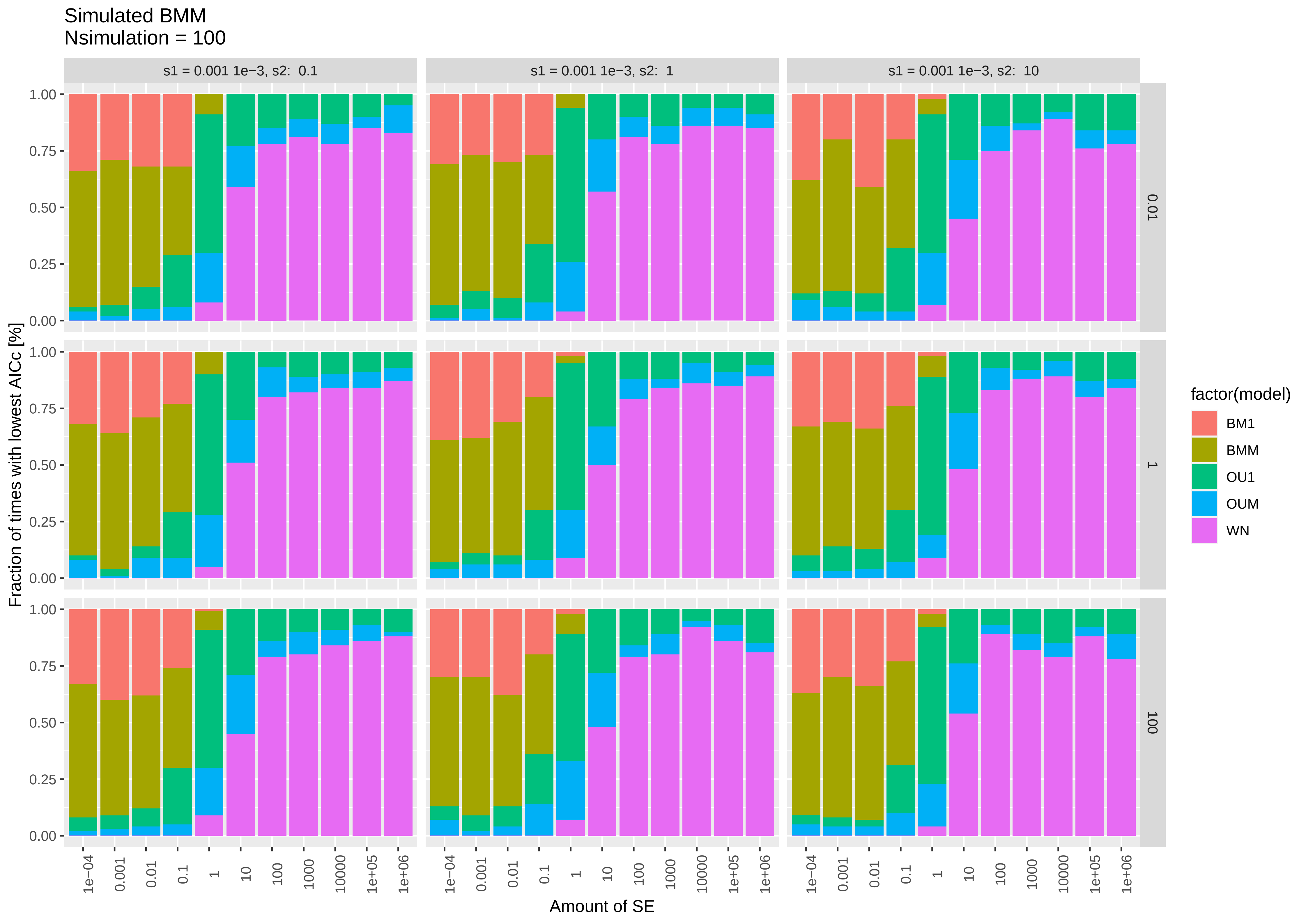
Simulated BMM
Nsimulation = 100
s1 = 0.001 1e−3, s2: 0.1
s1 = 0.001 1e−3, s2: 1
s1 = 0.001 1e−3, s2: 10
0.01
1
100
Fraction of times with lowest AICc [%]
1.00
0.75
0.50
0.25
0.00
1e−04
0.001
0.01
0.1
1
10
100
1000
10000
1e+05
1e+06
Amount of SE
factor(model)
BM1
BMM
OU1
OUM
WN

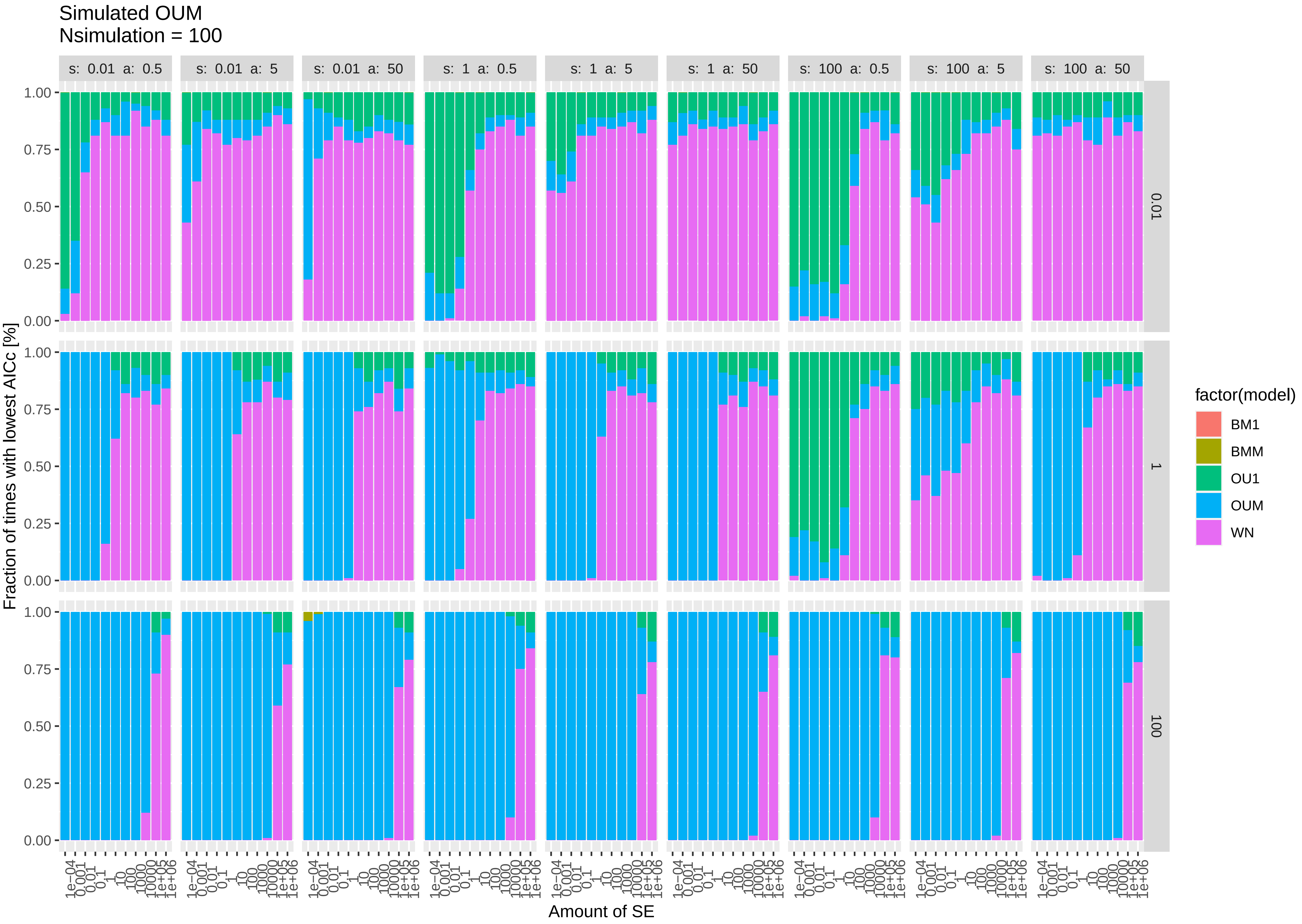
Simulated OUM
Nsimulation = 100
s: 0.01 a: 0.5
s: 0.01 a: 5
s: 0.01 a: 50
s: 1 a: 0.5
s: 1 a: 5
s: 1 a: 50
s: 100 a: 0.5
s: 100 a: 5
s: 100 a: 50
0.01
1
100
Fraction of times with lowest AICc [%]
0.00
0.25
0.50
0.75
1.00
1e−04
0.001
0.01
0.1
1
10
100
1000
10000
1e+05
1e+06
Amount of SE
factor(model)
BM1
BMM
OU1
OUM
WN

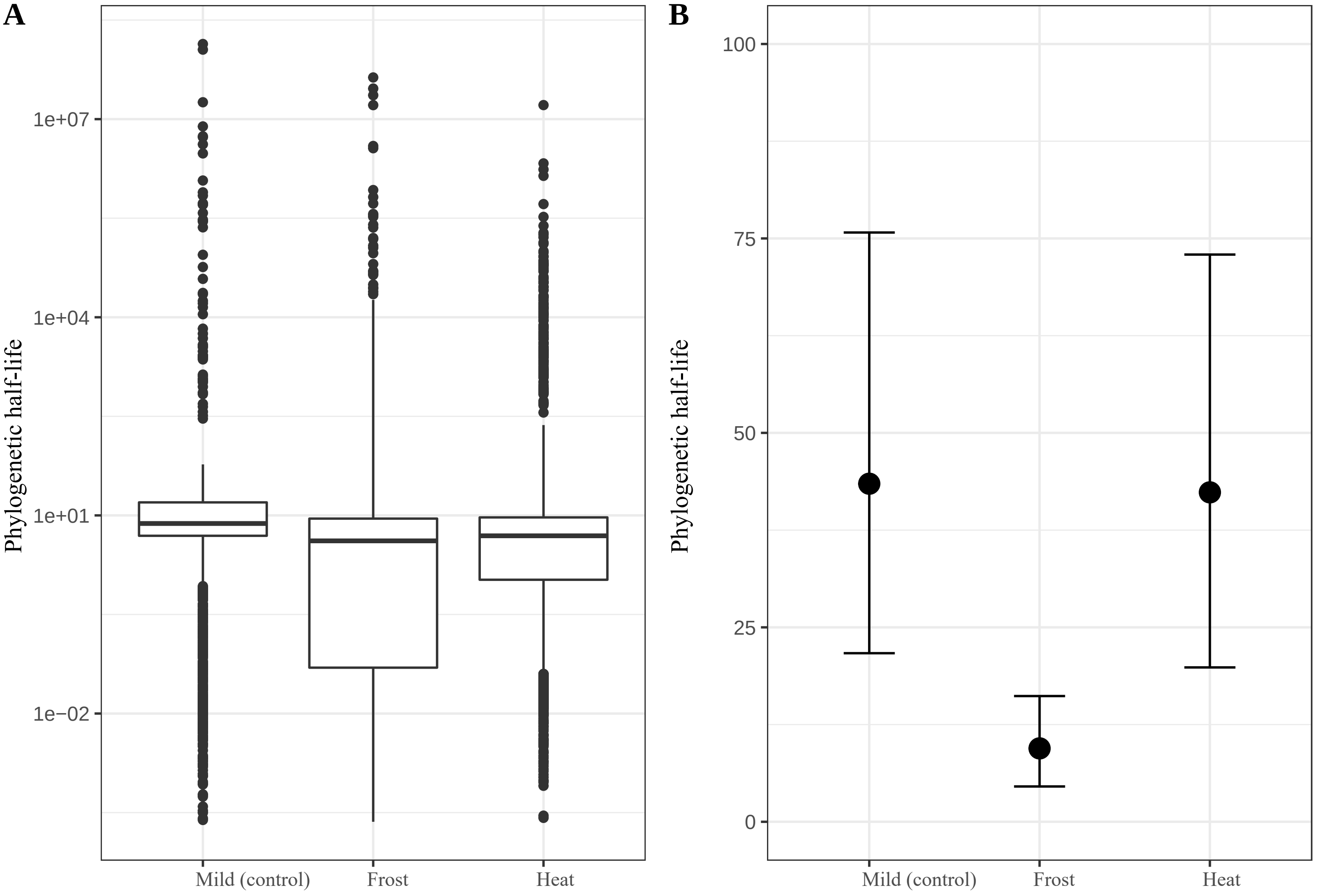

A
B
Phylogenetic half-life
1e+07
1e+04
1e+01
1e−02
Mild (control)
Frost
Heat
Phylogenetic half-life
100
75
50
25
0
Mild (control)
Frost
Heat

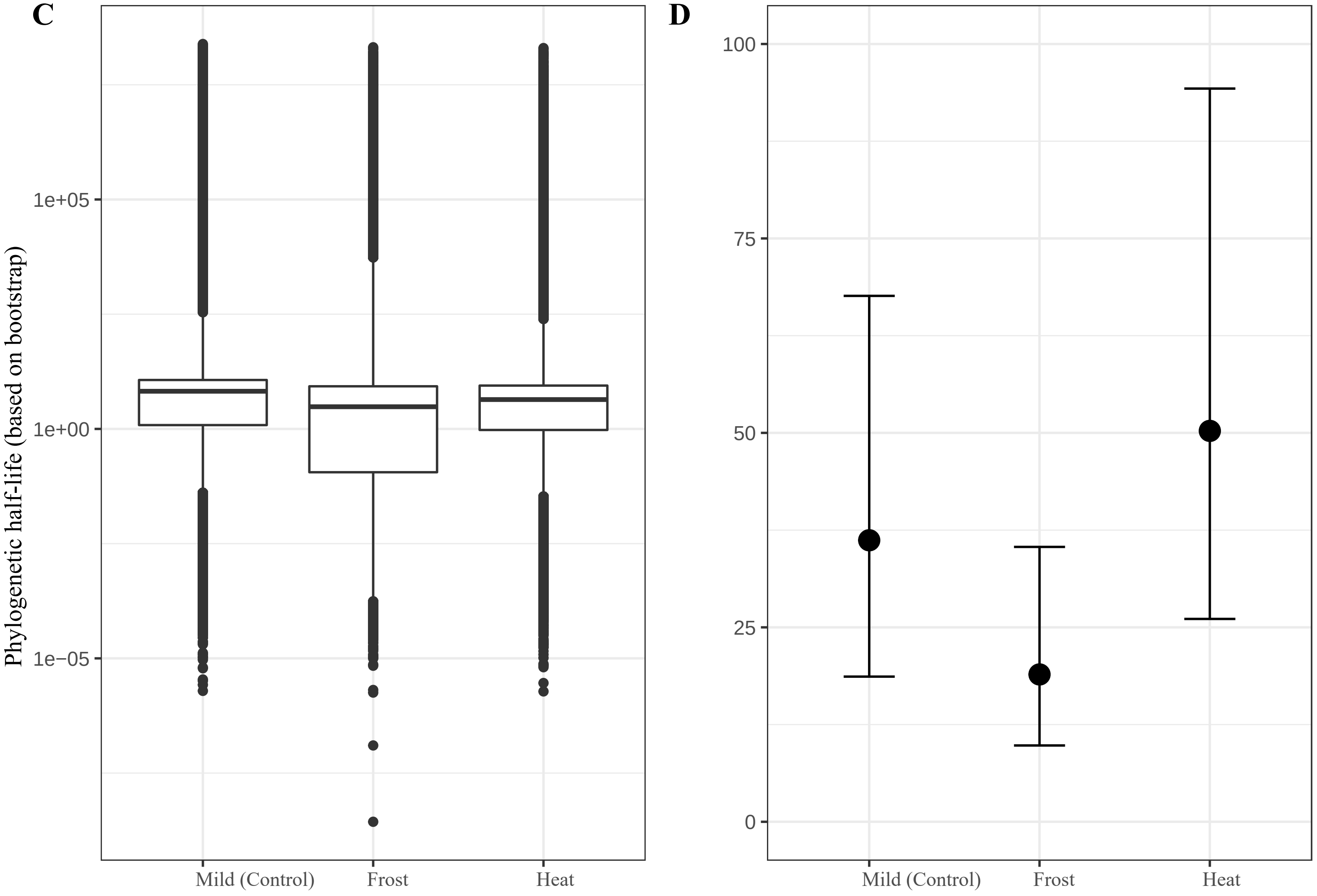

C
D
Phylogenetic half-life (based on bootstrap)
1e+05
1e+00
1e−05
100
75
50
25
0
Mild (Control)
Frost
Heat
Mild (Control)
Frost
Heat
