## Supplementary material for "Trait divergence and trade-offs among Brassicaceae species differing in elevational distribution": A5_M01_PhyloTraitCorr_ForSub.pdf

Multi-trait correlations among all traits expressed in the three growth treatments (based on mean between S1 and S2). Pearson's correlation coefficients and the associated p-values estimated with 'rcorr' {Hmisc}.
'C' : Mild (20°C); 'F' : Frost (-2°C, 1h); 'H' : Heat (+40°C, 1h). * Indicate sgnificant correlation (*p-value* < 0.05).

| Trait_1 | Trait_2 | corr.coef | p-value | |
|---|---|---|---|---|
| ASYM.C | LA.C | 0.810 | 0.000 | * |
| ASYM.C | LDI.C | 0.630 | 0.000 | * |
| ASYM.C | LDMC.C | 0.490 | 0.000 | * |
| ASYM.C | MGR.C | 0.210 | 0.080 | |
| ASYM.C | SLA.C | -0.320 | 0.007 | * |
| ASYM.C | SSIZ.C | 0.130 | 0.299 | |
| ASYM.C | TGER.C | -0.290 | 0.017 | * |
| ASYM.C | negXMID.C | -0.040 | 0.743 | |
| ASYM.C | ASYM.F | 0.900 | 0.000 | * |
| ASYM.C | LA.F | 0.770 | 0.000 | * |
| ASYM.C | LDI.F | 0.600 | 0.000 | * |
| ASYM.C | LDMC.F | 0.430 | 0.000 | * |
| ASYM.C | MGR.F | 0.060 | 0.640 | |
| ASYM.C | RESmT2.F | -0.260 | 0.031 | * |
| ASYM.C | SLA.F | -0.290 | 0.017 | * |
| ASYM.C | TOL_ASYM.F | -0.080 | 0.507 | |
| ASYM.C | TOL_MGR.F | -0.130 | 0.274 | |
| ASYM.C | TOL_negXMID.F | -0.010 | 0.929 | |
| ASYM.C | negXMID.F | -0.030 | 0.814 | |
| ASYM.C | ASYM.H | 0.880 | 0.000 | * |
| ASYM.C | LA.H | 0.780 | 0.000 | * |
| ASYM.C | LDI.H | 0.670 | 0.000 | * |
| ASYM.C | LDMC.H | 0.440 | 0.000 | * |
| ASYM.C | MGR.H | -0.050 | 0.659 | |
| ASYM.C | RESpT2.H | -0.220 | 0.076 | |
| ASYM.C | SLA.H | -0.390 | 0.001 | * |
| ASYM.C | TOL_ASYM.H | 0.570 | 0.000 | * |
| ASYM.C | TOL_MGR.H | 0.120 | 0.318 | |
| ASYM.C | TOL_negXMID.H | -0.280 | 0.021 | * |
| ASYM.C | negXMID.H | -0.320 | 0.007 | * |
| ASYM.C | IGR.C | 0.260 | 0.033 | * |
| ASYM.C | LTH.C | 0.120 | 0.324 | |
| ASYM.C | NLEA.C | -0.320 | 0.007 | * |
| ASYM.C | RESmT1.C | 0.050 | 0.697 | |
| ASYM.C | RESmT2.C | 0.270 | 0.022 | * |
| ASYM.C | RESpT1.C | 0.200 | 0.104 | |
| ASYM.C | RESpT2.C | 0.020 | 0.887 | |
| ASYM.C | IGR.F | 0.030 | 0.801 | |
| ASYM.C | LTH.F | -0.110 | 0.371 | |
| ASYM.C | NLEA.F | -0.360 | 0.003 | * |
| ASYM.C | TOL_IGR.F | -0.200 | 0.100 | |
| ASYM.C | IGR.H | -0.010 | 0.933 | |
| ASYM.C | LTH.H | 0.040 | 0.720 | |

| Trait_1 | Trait_2 | corr.coef | p-value | |
|---|---|---|---|---|
| ASYM.C | NLEA.H | -0.290 | 0.015 | * |
| ASYM.C | TOL_IGR.H | 0.050 | 0.694 | |
| ASYM.C | LA.C | 0.810 | 0.000 | * |
| LA.C | LDI.C | 0.500 | 0.000 | * |
| LA.C | LDMC.C | 0.500 | 0.000 | * |
| LA.C | MGR.C | 0.200 | 0.102 | |
| LA.C | SLA.C | -0.400 | 0.001 | * |
| LA.C | SSIZ.C | 0.180 | 0.138 | |
| LA.C | TGER.C | -0.180 | 0.141 | |
| LA.C | negXMID.C | 0.100 | 0.429 | |
| ASYM.F | LA.C | 0.840 | 0.000 | * |
| LA.C | LA.F | 0.930 | 0.000 | * |
| LA.C | LDI.F | 0.480 | 0.000 | * |
| LA.C | LDMC.F | 0.420 | 0.000 | * |
| LA.C | MGR.F | 0.050 | 0.674 | |
| LA.C | RESmT2.F | -0.300 | 0.012 | * |
| LA.C | SLA.F | -0.420 | 0.000 | * |
| LA.C | TOL_ASYM.F | 0.070 | 0.575 | |
| LA.C | TOL_MGR.F | -0.070 | 0.544 | |
| LA.C | TOL_negXMID.F | -0.180 | 0.137 | |
| LA.C | negXMID.F | -0.070 | 0.593 | |
| ASYM.H | LA.C | 0.870 | 0.000 | * |
| LA.C | LA.H | 0.930 | 0.000 | * |
| LA.C | LDI.H | 0.610 | 0.000 | * |
| LA.C | LDMC.H | 0.480 | 0.000 | * |
| LA.C | MGR.H | -0.070 | 0.557 | |
| LA.C | RESpT2.H | -0.190 | 0.126 | |
| LA.C | SLA.H | -0.520 | 0.000 | * |
| LA.C | TOL_ASYM.H | 0.560 | 0.000 | * |
| LA.C | TOL_MGR.H | 0.070 | 0.546 | |
| LA.C | TOL_negXMID.H | -0.360 | 0.003 | * |
| LA.C | negXMID.H | -0.360 | 0.003 | * |
| IGR.C | LA.C | 0.250 | 0.041 | * |
| LA.C | LTH.C | 0.180 | 0.141 | |
| LA.C | NLEA.C | -0.450 | 0.000 | * |
| LA.C | RESmT1.C | 0.180 | 0.146 | |
| LA.C | RESmT2.C | 0.160 | 0.195 | |
| LA.C | RESpT1.C | 0.210 | 0.091 | |
| LA.C | RESpT2.C | 0.070 | 0.593 | |
| IGR.F | LA.C | 0.060 | 0.604 | |
| LA.C | LTH.F | -0.030 | 0.782 | |
| LA.C | NLEA.F | -0.430 | 0.000 | * |
| LA.C | TOL_IGR.F | -0.200 | 0.099 | |

| Trait_1 | Trait_2 | corr.coef | p-value | |
|---|---|---|---|---|
| IGR.H | LA.C | 0.040 | 0.721 | |
| LA.C | LTH.H | 0.120 | 0.312 | |
| LA.C | NLEA.H | -0.420 | 0.000 | * |
| LA.C | TOL_IGR.H | 0.080 | 0.536 | |
| ASYM.C | LDI.C | 0.630 | 0.000 | * |
| LA.C | LDI.C | 0.500 | 0.000 | * |
| LDI.C | LDMC.C | 0.300 | 0.013 | * |
| LDI.C | MGR.C | 0.110 | 0.349 | |
| LDI.C | SLA.C | 0.020 | 0.894 | |
| LDI.C | SSIZ.C | -0.150 | 0.205 | |
| LDI.C | TGER.C | -0.120 | 0.336 | |
| LDI.C | negXMID.C | -0.030 | 0.837 | |
| ASYM.F | LDI.C | 0.660 | 0.000 | * |
| LA.F | LDI.C | 0.410 | 0.000 | * |
| LDI.C | LDI.F | 0.950 | 0.000 | * |
| LDI.C | LDMC.F | 0.320 | 0.008 | * |
| LDI.C | MGR.F | -0.020 | 0.882 | |
| LDI.C | RESmT2.F | -0.060 | 0.633 | |
| LDI.C | SLA.F | 0.060 | 0.624 | |
| LDI.C | TOL_ASYM.F | 0.020 | 0.855 | |
| LDI.C | TOL_MGR.F | -0.080 | 0.527 | |
| LDI.C | TOL_negXMID.F | 0.020 | 0.843 | |
| LDI.C | negXMID.F | -0.060 | 0.648 | |
| ASYM.H | LDI.C | 0.650 | 0.000 | * |
| LA.H | LDI.C | 0.440 | 0.000 | * |
| LDI.C | LDI.H | 0.920 | 0.000 | * |
| LDI.C | LDMC.H | 0.340 | 0.004 | * |
| LDI.C | MGR.H | -0.020 | 0.839 | |
| LDI.C | RESpT2.H | -0.100 | 0.424 | |
| LDI.C | SLA.H | -0.110 | 0.352 | |
| LDI.C | TOL_ASYM.H | 0.430 | 0.000 | * |
| LDI.C | TOL_MGR.H | 0.170 | 0.155 | |
| LDI.C | TOL_negXMID.H | -0.330 | 0.006 | * |
| LDI.C | negXMID.H | -0.230 | 0.058 | |
| IGR.C | LDI.C | 0.200 | 0.108 | |
| LDI.C | LTH.C | -0.260 | 0.034 | * |
| LDI.C | NLEA.C | -0.020 | 0.845 | |
| LDI.C | RESmT1.C | -0.190 | 0.109 | |
| LDI.C | RESmT2.C | 0.140 | 0.266 | |
| LDI.C | RESpT1.C | -0.110 | 0.353 | |
| LDI.C | RESpT2.C | 0.030 | 0.780 | |
| IGR.F | LDI.C | 0.040 | 0.723 | |
| LDI.C | LTH.F | -0.390 | 0.001 | * |
| LDI.C | NLEA.F | 0.000 | 0.975 | |
| LDI.C | TOL_IGR.F | -0.210 | 0.086 | |
| IGR.H | LDI.C | -0.140 | 0.242 | |
| LDI.C | LTH.H | 0.040 | 0.728 | |
| LDI.C | NLEA.H | 0.020 | 0.898 | |
| LDI.C | TOL_IGR.H | 0.030 | 0.803 | |

| Trait_1 | Trait_2 | corr.coef | p-value | |
|---|---|---|---|---|
| ASYM.C | LDMC.C | 0.490 | 0.000 | * |
| LA.C | LDMC.C | 0.500 | 0.000 | * |
| LDI.C | LDMC.C | 0.300 | 0.013 | * |
| LDMC.C | MGR.C | 0.200 | 0.097 | |
| LDMC.C | SLA.C | -0.490 | 0.000 | * |
| LDMC.C | SSIZ.C | 0.190 | 0.124 | |
| LDMC.C | TGER.C | -0.140 | 0.240 | |
| LDMC.C | negXMID.C | 0.140 | 0.235 | |
| ASYM.F | LDMC.C | 0.560 | 0.000 | * |
| LA.F | LDMC.C | 0.530 | 0.000 | * |
| LDI.F | LDMC.C | 0.310 | 0.009 | * |
| LDMC.C | LDMC.F | 0.790 | 0.000 | * |
| LDMC.C | MGR.F | -0.020 | 0.862 | |
| LDMC.C | RESmT2.F | -0.320 | 0.008 | * |
| LDMC.C | SLA.F | -0.460 | 0.000 | * |
| LDMC.C | TOL_ASYM.F | 0.090 | 0.468 | |
| LDMC.C | TOL_MGR.F | -0.030 | 0.827 | |
| LDMC.C | TOL_negXMID.F | 0.030 | 0.805 | |
| LDMC.C | negXMID.F | 0.080 | 0.515 | |
| ASYM.H | LDMC.C | 0.530 | 0.000 | * |
| LA.H | LDMC.C | 0.540 | 0.000 | * |
| LDI.H | LDMC.C | 0.390 | 0.001 | * |
| LDMC.C | LDMC.H | 0.790 | 0.000 | * |
| LDMC.C | MGR.H | -0.200 | 0.095 | |
| LDMC.C | RESpT2.H | 0.150 | 0.221 | |
| LDMC.C | SLA.H | -0.510 | 0.000 | * |
| LDMC.C | TOL_ASYM.H | 0.350 | 0.003 | * |
| LDMC.C | TOL_MGR.H | -0.130 | 0.304 | |
| LDMC.C | TOL_negXMID.H | -0.140 | 0.258 | |
| LDMC.C | negXMID.H | -0.220 | 0.064 | |
| IGR.C | LDMC.C | 0.130 | 0.292 | |
| LDMC.C | LTH.C | 0.050 | 0.706 | |
| LDMC.C | NLEA.C | -0.180 | 0.147 | |
| LDMC.C | RESmT1.C | -0.090 | 0.454 | |
| LDMC.C | RESmT2.C | 0.200 | 0.095 | |
| LDMC.C | RESpT1.C | 0.190 | 0.128 | |
| LDMC.C | RESpT2.C | 0.230 | 0.059 | |
| IGR.F | LDMC.C | -0.150 | 0.223 | |
| LDMC.C | LTH.F | -0.250 | 0.035 | * |
| LDMC.C | NLEA.F | -0.110 | 0.375 | |
| LDMC.C | TOL_IGR.F | -0.160 | 0.186 | |
| IGR.H | LDMC.C | 0.130 | 0.296 | |
| LDMC.C | LTH.H | 0.020 | 0.889 | |
| LDMC.C | NLEA.H | -0.060 | 0.602 | |
| LDMC.C | TOL_IGR.H | 0.110 | 0.369 | |
| ASYM.C | MGR.C | 0.210 | 0.080 | |
| LA.C | MGR.C | 0.200 | 0.102 | |
| LDI.C | MGR.C | 0.110 | 0.349 | |
| LDMC.C | MGR.C | 0.200 | 0.097 | |

| Trait_1 | Trait_2 | corr.coef | p-value | |
|---|---|---|---|---|
| MGR.C | SLA.C | -0.230 | 0.062 | |
| MGR.C | SSIZ.C | 0.030 | 0.794 | |
| MGR.C | TGER.C | -0.030 | 0.830 | |
| MGR.C | negXMID.C | 0.170 | 0.153 | |
| ASYM.F | MGR.C | 0.270 | 0.025 | * |
| LA.F | MGR.C | 0.220 | 0.064 | |
| LDI.F | MGR.C | 0.150 | 0.220 | |
| LDMC.F | MGR.C | 0.100 | 0.415 | |
| MGR.C | MGR.F | 0.140 | 0.238 | |
| MGR.C | RESmT2.F | -0.100 | 0.403 | |
| MGR.C | SLA.F | -0.150 | 0.211 | |
| MGR.C | TOL_ASYM.F | 0.130 | 0.271 | |
| MGR.C | TOL_MGR.F | -0.350 | 0.003 | * |
| MGR.C | TOL_negXMID.F | -0.100 | 0.423 | |
| MGR.C | negXMID.F | -0.080 | 0.531 | |
| ASYM.H | MGR.C | 0.260 | 0.030 | * |
| LA.H | MGR.C | 0.150 | 0.230 | |
| LDI.H | MGR.C | 0.060 | 0.626 | |
| LDMC.H | MGR.C | 0.090 | 0.452 | |
| MGR.C | MGR.H | -0.070 | 0.576 | |
| MGR.C | RESpT2.H | -0.130 | 0.284 | |
| MGR.C | SLA.H | -0.270 | 0.027 | * |
| MGR.C | TOL_ASYM.H | 0.150 | 0.209 | |
| MGR.C | TOL_MGR.H | -0.190 | 0.128 | |
| MGR.C | TOL_negXMID.H | 0.030 | 0.836 | |
| MGR.C | negXMID.H | -0.150 | 0.226 | |
| IGR.C | MGR.C | 0.030 | 0.827 | |
| LTH.C | MGR.C | 0.060 | 0.649 | |
| MGR.C | NLEA.C | 0.040 | 0.739 | |
| MGR.C | RESmT1.C | 0.080 | 0.498 | |
| MGR.C | RESmT2.C | 0.020 | 0.890 | |
| MGR.C | RESpT1.C | 0.180 | 0.142 | |
| MGR.C | RESpT2.C | 0.120 | 0.320 | |
| IGR.F | MGR.C | 0.130 | 0.298 | |
| LTH.F | MGR.C | 0.020 | 0.896 | |
| MGR.C | NLEA.F | 0.050 | 0.656 | |
| MGR.C | TOL_IGR.F | -0.040 | 0.726 | |
| IGR.H | MGR.C | 0.110 | 0.376 | |
| LTH.H | MGR.C | 0.050 | 0.667 | |
| MGR.C | NLEA.H | 0.080 | 0.521 | |
| MGR.C | TOL_IGR.H | 0.180 | 0.149 | |
| ASYM.C | SLA.C | -0.320 | 0.007 | * |
| LA.C | SLA.C | -0.400 | 0.001 | * |
| LDI.C | SLA.C | 0.020 | 0.894 | |
| LDMC.C | SLA.C | -0.490 | 0.000 | * |
| MGR.C | SLA.C | -0.230 | 0.062 | |
| SLA.C | SSIZ.C | -0.470 | 0.000 | * |
| SLA.C | TGER.C | 0.090 | 0.447 | |
| negXMID.C | SLA.C | -0.010 | 0.941 | |

| Trait_1 | Trait_2 | corr.coef | p-value | |
|---|---|---|---|---|
| ASYM.F | SLA.C | -0.310 | 0.009 | * |
| LA.F | SLA.C | -0.360 | 0.002 | * |
| LDI.F | SLA.C | -0.010 | 0.933 | |
| LDMC.F | SLA.C | -0.320 | 0.007 | * |
| MGR.F | SLA.C | 0.100 | 0.392 | |
| RESmT2.F | SLA.C | 0.260 | 0.030 | * |
| SLA.C | SLA.F | 0.800 | 0.000 | * |
| SLA.C | TOL_ASYM.F | 0.080 | 0.528 | |
| SLA.C | TOL_MGR.F | 0.150 | 0.219 | |
| SLA.C | TOL_negXMID.F | 0.190 | 0.110 | |
| negXMID.F | SLA.C | 0.140 | 0.258 | |
| ASYM.H | SLA.C | -0.290 | 0.015 | * |
| LA.H | SLA.C | -0.350 | 0.003 | * |
| LDI.H | SLA.C | -0.080 | 0.512 | |
| LDMC.H | SLA.C | -0.300 | 0.012 | * |
| MGR.H | SLA.C | 0.040 | 0.752 | |
| RESpT2.H | SLA.C | 0.030 | 0.820 | |
| SLA.C | SLA.H | 0.680 | 0.000 | * |
| SLA.C | TOL_ASYM.H | -0.060 | 0.639 | |
| SLA.C | TOL_MGR.H | 0.020 | 0.886 | |
| SLA.C | TOL_negXMID.H | -0.100 | 0.404 | |
| negXMID.H | SLA.C | 0.220 | 0.064 | |
| IGR.C | SLA.C | -0.080 | 0.506 | |
| LTH.C | SLA.C | -0.560 | 0.000 | * |
| NLEA.C | SLA.C | 0.410 | 0.000 | * |
| RESmT1.C | SLA.C | -0.370 | 0.002 | * |
| RESmT2.C | SLA.C | -0.100 | 0.410 | |
| RESpT1.C | SLA.C | -0.460 | 0.000 | * |
| RESpT2.C | SLA.C | 0.200 | 0.107 | |
| IGR.F | SLA.C | -0.010 | 0.924 | |
| LTH.F | SLA.C | -0.330 | 0.005 | * |
| NLEA.F | SLA.C | 0.340 | 0.004 | * |
| SLA.C | TOL_IGR.F | 0.110 | 0.385 | |
| IGR.H | SLA.C | -0.260 | 0.032 | * |
| LTH.H | SLA.C | -0.180 | 0.133 | |
| NLEA.H | SLA.C | 0.400 | 0.001 | * |
| SLA.C | TOL_IGR.H | -0.060 | 0.633 | |
| ASYM.C | SSIZ.C | 0.130 | 0.299 | |
| LA.C | SSIZ.C | 0.180 | 0.138 | |
| LDI.C | SSIZ.C | -0.150 | 0.205 | |
| LDMC.C | SSIZ.C | 0.190 | 0.124 | |
| MGR.C | SSIZ.C | 0.030 | 0.794 | |
| SLA.C | SSIZ.C | -0.470 | 0.000 | * |
| SSIZ.C | TGER.C | 0.210 | 0.081 | |
| negXMID.C | SSIZ.C | 0.340 | 0.004 | * |
| ASYM.F | SSIZ.C | 0.150 | 0.211 | |
| LA.F | SSIZ.C | 0.210 | 0.083 | |
| LDI.F | SSIZ.C | -0.230 | 0.061 | |
| LDMC.F | SSIZ.C | 0.170 | 0.169 | |

| Trait_1 | Trait_2 | corr.coef | p-value | |
|---|---|---|---|---|
| MGR.F | SSIZ.C | 0.010 | 0.940 | |
| RESmT2.F | SSIZ.C | -0.170 | 0.150 | |
| SLA.F | SSIZ.C | -0.610 | 0.000 | * |
| SSIZ.C | TOL_ASYM.F | 0.000 | 0.995 | |
| SSIZ.C | TOL_MGR.F | -0.110 | 0.358 | |
| SSIZ.C | TOL_negXMID.F | -0.220 | 0.070 | |
| negXMID.F | SSIZ.C | 0.150 | 0.220 | |
| ASYM.H | SSIZ.C | 0.120 | 0.338 | |
| LA.H | SSIZ.C | 0.210 | 0.086 | |
| LDI.H | SSIZ.C | -0.150 | 0.228 | |
| LDMC.H | SSIZ.C | 0.170 | 0.153 | |
| MGR.H | SSIZ.C | -0.070 | 0.552 | |
| RESpT2.H | SSIZ.C | 0.040 | 0.726 | |
| SLA.H | SSIZ.C | -0.470 | 0.000 | * |
| SSIZ.C | TOL_ASYM.H | 0.050 | 0.686 | |
| SSIZ.C | TOL_MGR.H | 0.010 | 0.954 | |
| SSIZ.C | TOL_negXMID.H | 0.080 | 0.506 | |
| negXMID.H | SSIZ.C | 0.190 | 0.109 | |
| IGR.C | SSIZ.C | 0.150 | 0.232 | |
| LTH.C | SSIZ.C | 0.400 | 0.001 | * |
| NLEA.C | SSIZ.C | -0.430 | 0.000 | * |
| RESmT1.C | SSIZ.C | 0.270 | 0.026 | * |
| RESmT2.C | SSIZ.C | -0.030 | 0.806 | |
| RESpT1.C | SSIZ.C | 0.370 | 0.002 | * |
| RESpT2.C | SSIZ.C | -0.140 | 0.237 | |
| IGR.F | SSIZ.C | -0.170 | 0.159 | |
| LTH.F | SSIZ.C | 0.290 | 0.015 | * |
| NLEA.F | SSIZ.C | -0.430 | 0.000 | * |
| SSIZ.C | TOL_IGR.F | -0.260 | 0.031 | * |
| IGR.H | SSIZ.C | 0.110 | 0.348 | |
| LTH.H | SSIZ.C | 0.120 | 0.336 | |
| NLEA.H | SSIZ.C | -0.430 | 0.000 | * |
| SSIZ.C | TOL_IGR.H | 0.020 | 0.856 | |
| ASYM.C | TGER.C | -0.290 | 0.017 | * |
| LA.C | TGER.C | -0.180 | 0.141 | |
| LDI.C | TGER.C | -0.120 | 0.336 | |
| LDMC.C | TGER.C | -0.140 | 0.240 | |
| MGR.C | TGER.C | -0.030 | 0.830 | |
| SLA.C | TGER.C | 0.090 | 0.447 | |
| SSIZ.C | TGER.C | 0.210 | 0.081 | |
| negXMID.C | TGER.C | 0.390 | 0.001 | * |
| ASYM.F | TGER.C | -0.300 | 0.011 | * |
| LA.F | TGER.C | -0.220 | 0.076 | |
| LDI.F | TGER.C | -0.140 | 0.264 | |
| LDMC.F | TGER.C | -0.240 | 0.050 | |
| MGR.F | TGER.C | 0.030 | 0.776 | |
| RESmT2.F | TGER.C | 0.300 | 0.013 | * |
| SLA.F | TGER.C | 0.070 | 0.587 | |
| TGER.C | TOL_ASYM.F | 0.030 | 0.836 | |

| Trait_1 | Trait_2 | corr.coef | p-value | |
|---|---|---|---|---|
| TGER.C | TOL_MGR.F | 0.000 | 0.970 | |
| TGER.C | TOL_negXMID.F | -0.100 | 0.431 | |
| negXMID.F | TGER.C | 0.370 | 0.002 | * |
| ASYM.H | TGER.C | -0.260 | 0.033 | * |
| LA.H | TGER.C | -0.190 | 0.111 | |
| LDI.H | TGER.C | -0.170 | 0.151 | |
| LDMC.H | TGER.C | -0.140 | 0.251 | |
| MGR.H | TGER.C | 0.150 | 0.233 | |
| RESpT2.H | TGER.C | 0.180 | 0.132 | |
| SLA.H | TGER.C | 0.040 | 0.766 | |
| TGER.C | TOL_ASYM.H | -0.310 | 0.009 | * |
| TGER.C | TOL_MGR.H | -0.010 | 0.959 | |
| TGER.C | TOL_negXMID.H | 0.050 | 0.710 | |
| negXMID.H | TGER.C | 0.430 | 0.000 | * |
| IGR.C | TGER.C | -0.170 | 0.175 | |
| LTH.C | TGER.C | -0.150 | 0.222 | |
| NLEA.C | TGER.C | -0.310 | 0.009 | * |
| RESmT1.C | TGER.C | 0.000 | 0.990 | |
| RESmT2.C | TGER.C | -0.320 | 0.007 | * |
| RESpT1.C | TGER.C | -0.110 | 0.362 | |
| RESpT2.C | TGER.C | -0.100 | 0.392 | |
| IGR.F | TGER.C | 0.060 | 0.650 | |
| LTH.F | TGER.C | 0.000 | 0.969 | |
| NLEA.F | TGER.C | -0.290 | 0.017 | * |
| TGER.C | TOL_IGR.F | 0.170 | 0.168 | |
| IGR.H | TGER.C | 0.030 | 0.827 | |
| LTH.H | TGER.C | 0.060 | 0.642 | |
| NLEA.H | TGER.C | -0.250 | 0.042 | * |
| TGER.C | TOL_IGR.H | 0.020 | 0.871 | |
| ASYM.C | negXMID.C | -0.040 | 0.743 | |
| LA.C | negXMID.C | 0.100 | 0.429 | |
| LDI.C | negXMID.C | -0.030 | 0.837 | |
| LDMC.C | negXMID.C | 0.140 | 0.235 | |
| MGR.C | negXMID.C | 0.170 | 0.153 | |
| negXMID.C | SLA.C | -0.010 | 0.941 | |
| negXMID.C | SSIZ.C | 0.340 | 0.004 | * |
| negXMID.C | TGER.C | 0.390 | 0.001 | * |
| ASYM.F | negXMID.C | 0.050 | 0.688 | |
| LA.F | negXMID.C | 0.180 | 0.142 | |
| LDI.F | negXMID.C | -0.030 | 0.804 | |
| LDMC.F | negXMID.C | 0.040 | 0.756 | |
| MGR.F | negXMID.C | 0.170 | 0.171 | |
| negXMID.C | RESmT2.F | 0.150 | 0.205 | |
| negXMID.C | SLA.F | -0.100 | 0.405 | |
| negXMID.C | TOL_ASYM.F | 0.290 | 0.016 | * |
| negXMID.C | TOL_MGR.F | 0.030 | 0.837 | |
| negXMID.C | TOL_negXMID.F | -0.260 | 0.029 | * |
| negXMID.C | negXMID.F | 0.440 | 0.000 | * |
| ASYM.H | negXMID.C | 0.110 | 0.377 | |

| Trait_1 | Trait_2 | corr.coef | p-value | |
|---|---|---|---|---|
| LA.H | negXMID.C | 0.170 | 0.153 | |
| LDI.H | negXMID.C | -0.010 | 0.934 | |
| LDMC.H | negXMID.C | 0.080 | 0.509 | |
| MGR.H | negXMID.C | -0.190 | 0.125 | |
| negXMID.C | RESpT2.H | -0.080 | 0.534 | |
| negXMID.C | SLA.H | -0.050 | 0.689 | |
| negXMID.C | TOL_ASYM.H | -0.020 | 0.886 | |
| negXMID.C | TOL_MGR.H | -0.180 | 0.128 | |
| negXMID.C | TOL_negXMID.H | 0.170 | 0.162 | |
| negXMID.C | negXMID.H | 0.430 | 0.000 | * |
| IGR.C | negXMID.C | 0.030 | 0.787 | |
| LTH.C | negXMID.C | 0.060 | 0.644 | |
| negXMID.C | NLEA.C | -0.150 | 0.204 | |
| negXMID.C | RESmT1.C | 0.080 | 0.504 | |
| negXMID.C | RESmT2.C | 0.040 | 0.735 | |
| negXMID.C | RESpT1.C | 0.000 | 0.998 | |
| negXMID.C | RESpT2.C | -0.030 | 0.806 | |
| IGR.F | negXMID.C | -0.140 | 0.256 | |
| LTH.F | negXMID.C | 0.010 | 0.926 | |
| negXMID.C | NLEA.F | -0.090 | 0.487 | |
| negXMID.C | TOL_IGR.F | -0.120 | 0.337 | |
| IGR.H | negXMID.C | -0.010 | 0.910 | |
| LTH.H | negXMID.C | 0.090 | 0.464 | |
| negXMID.C | NLEA.H | -0.100 | 0.414 | |
| negXMID.C | TOL_IGR.H | 0.060 | 0.636 | |
| ASYM.C | ASYM.F | 0.900 | 0.000 | * |
| ASYM.F | LA.C | 0.840 | 0.000 | * |
| ASYM.F | LDI.C | 0.660 | 0.000 | * |
| ASYM.F | LDMC.C | 0.560 | 0.000 | * |
| ASYM.F | MGR.C | 0.270 | 0.025 | * |
| ASYM.F | SLA.C | -0.310 | 0.009 | * |
| ASYM.F | SSIZ.C | 0.150 | 0.211 | |
| ASYM.F | TGER.C | -0.300 | 0.011 | * |
| ASYM.F | negXMID.C | 0.050 | 0.688 | |
| ASYM.F | LA.F | 0.860 | 0.000 | * |
| ASYM.F | LDI.F | 0.650 | 0.000 | * |
| ASYM.F | LDMC.F | 0.510 | 0.000 | * |
| ASYM.F | MGR.F | -0.010 | 0.951 | |
| ASYM.F | RESmT2.F | -0.270 | 0.026 | * |
| ASYM.F | SLA.F | -0.350 | 0.003 | * |
| ASYM.F | TOL_ASYM.F | 0.210 | 0.081 | |
| ASYM.F | TOL_MGR.F | -0.100 | 0.410 | |
| ASYM.F | TOL_negXMID.F | -0.130 | 0.274 | |
| ASYM.F | negXMID.F | -0.130 | 0.294 | |
| ASYM.F | ASYM.H | 0.900 | 0.000 | * |
| ASYM.F | LA.H | 0.820 | 0.000 | * |
| ASYM.F | LDI.H | 0.710 | 0.000 | * |
| ASYM.F | LDMC.H | 0.510 | 0.000 | * |
| ASYM.F | MGR.H | -0.050 | 0.702 | |

| Trait_1 | Trait_2 | corr.coef | p-value | |
|---|---|---|---|---|
| ASYM.F | RESpT2.H | -0.210 | 0.083 | |
| ASYM.F | SLA.H | -0.460 | 0.000 | * |
| ASYM.F | TOL_ASYM.H | 0.610 | 0.000 | * |
| ASYM.F | TOL_MGR.H | 0.100 | 0.402 | |
| ASYM.F | TOL_negXMID.H | -0.320 | 0.007 | * |
| ASYM.F | negXMID.H | -0.350 | 0.003 | * |
| ASYM.F | IGR.C | 0.300 | 0.012 | * |
| ASYM.F | LTH.C | 0.120 | 0.332 | |
| ASYM.F | NLEA.C | -0.320 | 0.008 | * |
| ASYM.F | RESmT1.C | 0.050 | 0.669 | |
| ASYM.F | RESmT2.C | 0.260 | 0.033 | * |
| ASYM.F | RESpT1.C | 0.190 | 0.115 | |
| ASYM.F | RESpT2.C | 0.120 | 0.338 | |
| ASYM.F | IGR.F | 0.020 | 0.886 | |
| ASYM.F | LTH.F | -0.180 | 0.129 | |
| ASYM.F | NLEA.F | -0.300 | 0.013 | * |
| ASYM.F | TOL_IGR.F | -0.280 | 0.020 | * |
| ASYM.F | IGR.H | -0.040 | 0.725 | |
| ASYM.F | LTH.H | 0.070 | 0.591 | |
| ASYM.F | NLEA.H | -0.260 | 0.033 | * |
| ASYM.F | TOL_IGR.H | 0.080 | 0.534 | |
| ASYM.C | LA.F | 0.770 | 0.000 | * |
| LA.C | LA.F | 0.930 | 0.000 | * |
| LA.F | LDI.C | 0.410 | 0.000 | * |
| LA.F | LDMC.C | 0.530 | 0.000 | * |
| LA.F | MGR.C | 0.220 | 0.064 | |
| LA.F | SLA.C | -0.360 | 0.002 | * |
| LA.F | SSIZ.C | 0.210 | 0.083 | |
| LA.F | TGER.C | -0.220 | 0.076 | |
| LA.F | negXMID.C | 0.180 | 0.142 | |
| ASYM.F | LA.F | 0.860 | 0.000 | * |
| LA.F | LDI.F | 0.430 | 0.000 | * |
| LA.F | LDMC.F | 0.450 | 0.000 | * |
| LA.F | MGR.F | 0.040 | 0.723 | |
| LA.F | RESmT2.F | -0.230 | 0.061 | |
| LA.F | SLA.F | -0.440 | 0.000 | * |
| LA.F | TOL_ASYM.F | 0.160 | 0.194 | |
| LA.F | TOL_MGR.F | -0.040 | 0.738 | |
| LA.F | TOL_negXMID.F | -0.110 | 0.348 | |
| LA.F | negXMID.F | -0.010 | 0.903 | |
| ASYM.H | LA.F | 0.850 | 0.000 | * |
| LA.F | LA.H | 0.940 | 0.000 | * |
| LA.F | LDI.H | 0.540 | 0.000 | * |
| LA.F | LDMC.H | 0.520 | 0.000 | * |
| LA.F | MGR.H | -0.070 | 0.586 | |
| LA.F | RESpT2.H | -0.210 | 0.089 | |
| LA.F | SLA.H | -0.560 | 0.000 | * |
| LA.F | TOL_ASYM.H | 0.540 | 0.000 | * |
| LA.F | TOL_MGR.H | 0.050 | 0.663 | |

| Trait_1 | Trait_2 | corr.coef | p-value | |
|---|---|---|---|---|
| LA.F | TOL_negXMID.H | -0.310 | 0.009 | * |
| LA.F | negXMID.H | -0.290 | 0.017 | * |
| IGR.C | LA.F | 0.230 | 0.060 | |
| LA.F | LTH.C | 0.190 | 0.110 | |
| LA.F | NLEA.C | -0.490 | 0.000 | * |
| LA.F | RESmT1.C | 0.170 | 0.153 | |
| LA.F | RESmT2.C | 0.160 | 0.190 | |
| LA.F | RESpT1.C | 0.220 | 0.066 | |
| LA.F | RESpT2.C | 0.120 | 0.319 | |
| IGR.F | LA.F | 0.060 | 0.649 | |
| LA.F | LTH.F | -0.060 | 0.626 | |
| LA.F | NLEA.F | -0.470 | 0.000 | * |
| LA.F | TOL_IGR.F | -0.190 | 0.109 | |
| IGR.H | LA.F | 0.040 | 0.754 | |
| LA.F | LTH.H | 0.140 | 0.260 | |
| LA.F | NLEA.H | -0.440 | 0.000 | * |
| LA.F | TOL_IGR.H | 0.080 | 0.538 | |
| ASYM.C | LDI.F | 0.600 | 0.000 | * |
| LA.C | LDI.F | 0.480 | 0.000 | * |
| LDI.C | LDI.F | 0.950 | 0.000 | * |
| LDI.F | LDMC.C | 0.310 | 0.009 | * |
| LDI.F | MGR.C | 0.150 | 0.220 | |
| LDI.F | SLA.C | -0.010 | 0.933 | |
| LDI.F | SSIZ.C | -0.230 | 0.061 | |
| LDI.F | TGER.C | -0.140 | 0.264 | |
| LDI.F | negXMID.C | -0.030 | 0.804 | |
| ASYM.F | LDI.F | 0.650 | 0.000 | * |
| LA.F | LDI.F | 0.430 | 0.000 | * |
| LDI.F | LDMC.F | 0.300 | 0.013 | * |
| LDI.F | MGR.F | 0.000 | 0.976 | |
| LDI.F | RESmT2.F | -0.010 | 0.966 | |
| LDI.F | SLA.F | 0.070 | 0.572 | |
| LDI.F | TOL_ASYM.F | 0.090 | 0.442 | |
| LDI.F | TOL_MGR.F | -0.030 | 0.793 | |
| LDI.F | TOL_negXMID.F | 0.010 | 0.960 | |
| LDI.F | negXMID.F | -0.100 | 0.428 | |
| ASYM.H | LDI.F | 0.610 | 0.000 | * |
| LA.H | LDI.F | 0.430 | 0.000 | * |
| LDI.F | LDI.H | 0.890 | 0.000 | * |
| LDI.F | LDMC.H | 0.320 | 0.007 | * |
| LDI.F | MGR.H | -0.020 | 0.850 | |
| LDI.F | RESpT2.H | -0.070 | 0.575 | |
| LDI.F | SLA.H | -0.110 | 0.349 | |
| LDI.F | TOL_ASYM.H | 0.390 | 0.001 | * |
| LDI.F | TOL_MGR.H | 0.180 | 0.150 | |
| LDI.F | TOL_negXMID.H | -0.260 | 0.032 | * |
| LDI.F | negXMID.H | -0.210 | 0.080 | |
| IGR.C | LDI.F | 0.240 | 0.046 | * |
| LDI.F | LTH.C | -0.210 | 0.081 | |

| Trait_1 | Trait_2 | corr.coef | p-value | |
|---|---|---|---|---|
| LDI.F | NLEA.C | -0.030 | 0.835 | |
| LDI.F | RESmT1.C | -0.210 | 0.084 | |
| LDI.F | RESmT2.C | 0.200 | 0.104 | |
| LDI.F | RESpT1.C | -0.150 | 0.214 | |
| LDI.F | RESpT2.C | 0.030 | 0.831 | |
| IGR.F | LDI.F | 0.040 | 0.741 | |
| LDI.F | LTH.F | -0.320 | 0.008 | * |
| LDI.F | NLEA.F | 0.000 | 0.998 | |
| LDI.F | TOL_IGR.F | -0.250 | 0.035 | * |
| IGR.H | LDI.F | -0.120 | 0.319 | |
| LDI.F | LTH.H | 0.020 | 0.889 | |
| LDI.F | NLEA.H | 0.010 | 0.953 | |
| LDI.F | TOL_IGR.H | 0.040 | 0.763 | |
| ASYM.C | LDMC.F | 0.430 | 0.000 | * |
| LA.C | LDMC.F | 0.420 | 0.000 | * |
| LDI.C | LDMC.F | 0.320 | 0.008 | * |
| LDMC.C | LDMC.F | 0.790 | 0.000 | * |
| LDMC.F | MGR.C | 0.100 | 0.415 | |
| LDMC.F | SLA.C | -0.320 | 0.007 | * |
| LDMC.F | SSIZ.C | 0.170 | 0.169 | |
| LDMC.F | TGER.C | -0.240 | 0.050 | |
| LDMC.F | negXMID.C | 0.040 | 0.756 | |
| ASYM.F | LDMC.F | 0.510 | 0.000 | * |
| LA.F | LDMC.F | 0.450 | 0.000 | * |
| LDI.F | LDMC.F | 0.300 | 0.013 | * |
| LDMC.F | MGR.F | -0.110 | 0.368 | |
| LDMC.F | RESmT2.F | -0.440 | 0.000 | * |
| LDMC.F | SLA.F | -0.500 | 0.000 | * |
| LDMC.F | TOL_ASYM.F | 0.090 | 0.444 | |
| LDMC.F | TOL_MGR.F | 0.060 | 0.604 | |
| LDMC.F | TOL_negXMID.F | 0.030 | 0.828 | |
| LDMC.F | negXMID.F | -0.040 | 0.716 | |
| ASYM.H | LDMC.F | 0.500 | 0.000 | * |
| LA.H | LDMC.F | 0.460 | 0.000 | * |
| LDI.H | LDMC.F | 0.360 | 0.002 | * |
| LDMC.F | LDMC.H | 0.730 | 0.000 | * |
| LDMC.F | MGR.H | -0.210 | 0.076 | |
| LDMC.F | RESpT2.H | 0.030 | 0.788 | |
| LDMC.F | SLA.H | -0.430 | 0.000 | * |
| LDMC.F | TOL_ASYM.H | 0.380 | 0.001 | * |
| LDMC.F | TOL_MGR.H | -0.060 | 0.600 | |
| LDMC.F | TOL_negXMID.H | -0.160 | 0.181 | |
| LDMC.F | negXMID.H | -0.240 | 0.044 | * |
| IGR.C | LDMC.F | 0.030 | 0.831 | |
| LDMC.F | LTH.C | -0.080 | 0.502 | |
| LDMC.F | NLEA.C | -0.030 | 0.791 | |
| LDMC.F | RESmT1.C | -0.210 | 0.089 | |
| LDMC.F | RESmT2.C | 0.110 | 0.368 | |
| LDMC.F | RESpT1.C | 0.100 | 0.415 | |

| Trait_1 | Trait_2 | corr.coef | p-value | |
|---|---|---|---|---|
| LDMC.F | RESpT2.C | 0.300 | 0.012 | * |
| IGR.F | LDMC.F | -0.110 | 0.348 | |
| LDMC.F | LTH.F | -0.340 | 0.005 | * |
| LDMC.F | NLEA.F | 0.000 | 0.978 | |
| LDMC.F | TOL_IGR.F | -0.090 | 0.484 | |
| IGR.H | LDMC.F | 0.050 | 0.669 | |
| LDMC.F | LTH.H | -0.090 | 0.457 | |
| LDMC.F | NLEA.H | 0.070 | 0.595 | |
| LDMC.F | TOL_IGR.H | 0.120 | 0.321 | |
| ASYM.C | MGR.F | 0.060 | 0.640 | |
| LA.C | MGR.F | 0.050 | 0.674 | |
| LDI.C | MGR.F | -0.020 | 0.882 | |
| LDMC.C | MGR.F | -0.020 | 0.862 | |
| MGR.C | MGR.F | 0.140 | 0.238 | |
| MGR.F | SLA.C | 0.100 | 0.392 | |
| MGR.F | SSIZ.C | 0.010 | 0.940 | |
| MGR.F | TGER.C | 0.030 | 0.776 | |
| MGR.F | negXMID.C | 0.170 | 0.171 | |
| ASYM.F | MGR.F | -0.010 | 0.951 | |
| LA.F | MGR.F | 0.040 | 0.723 | |
| LDI.F | MGR.F | 0.000 | 0.976 | |
| LDMC.F | MGR.F | -0.110 | 0.368 | |
| MGR.F | RESmT2.F | -0.030 | 0.806 | |
| MGR.F | SLA.F | 0.190 | 0.117 | |
| MGR.F | TOL_ASYM.F | 0.010 | 0.960 | |
| MGR.F | TOL_MGR.F | 0.350 | 0.003 | * |
| MGR.F | TOL_negXMID.F | -0.010 | 0.940 | |
| MGR.F | negXMID.F | 0.210 | 0.089 | |
| ASYM.H | MGR.F | -0.010 | 0.927 | |
| LA.H | MGR.F | 0.060 | 0.648 | |
| LDI.H | MGR.F | -0.050 | 0.710 | |
| LDMC.H | MGR.F | 0.040 | 0.770 | |
| MGR.F | MGR.H | 0.090 | 0.462 | |
| MGR.F | RESpT2.H | 0.000 | 0.981 | |
| MGR.F | SLA.H | 0.120 | 0.320 | |
| MGR.F | TOL_ASYM.H | 0.000 | 0.992 | |
| MGR.F | TOL_MGR.H | 0.100 | 0.434 | |
| MGR.F | TOL_negXMID.H | 0.030 | 0.789 | |
| MGR.F | negXMID.H | 0.160 | 0.199 | |
| IGR.C | MGR.F | 0.130 | 0.296 | |
| LTH.C | MGR.F | -0.010 | 0.947 | |
| MGR.F | NLEA.C | 0.110 | 0.382 | |
| MGR.F | RESmT1.C | -0.120 | 0.327 | |
| MGR.F | RESmT2.C | 0.070 | 0.573 | |
| MGR.F | RESpT1.C | -0.070 | 0.587 | |
| MGR.F | RESpT2.C | -0.130 | 0.304 | |
| IGR.F | MGR.F | 0.120 | 0.324 | |
| LTH.F | MGR.F | 0.000 | 0.989 | |
| MGR.F | NLEA.F | 0.020 | 0.851 | |

| Trait_1 | Trait_2 | corr.coef | p-value | |
|---|---|---|---|---|
| MGR.F | TOL_IGR.F | 0.000 | 0.999 | |
| IGR.H | MGR.F | -0.040 | 0.766 | |
| LTH.H | MGR.F | -0.050 | 0.665 | |
| MGR.F | NLEA.H | 0.050 | 0.708 | |
| MGR.F | TOL_IGR.H | 0.060 | 0.609 | |
| ASYM.C | RESmT2.F | -0.260 | 0.031 | * |
| LA.C | RESmT2.F | -0.300 | 0.012 | * |
| LDI.C | RESmT2.F | -0.060 | 0.633 | |
| LDMC.C | RESmT2.F | -0.320 | 0.008 | * |
| MGR.C | RESmT2.F | -0.100 | 0.403 | |
| RESmT2.F | SLA.C | 0.260 | 0.030 | * |
| RESmT2.F | SSIZ.C | -0.170 | 0.150 | |
| RESmT2.F | TGER.C | 0.300 | 0.013 | * |
| negXMID.C | RESmT2.F | 0.150 | 0.205 | |
| ASYM.F | RESmT2.F | -0.270 | 0.026 | * |
| LA.F | RESmT2.F | -0.230 | 0.061 | |
| LDI.F | RESmT2.F | -0.010 | 0.966 | |
| LDMC.F | RESmT2.F | -0.440 | 0.000 | * |
| MGR.F | RESmT2.F | -0.030 | 0.806 | |
| RESmT2.F | SLA.F | 0.260 | 0.032 | * |
| RESmT2.F | TOL_ASYM.F | 0.070 | 0.595 | |
| RESmT2.F | TOL_MGR.F | 0.080 | 0.490 | |
| RESmT2.F | TOL_negXMID.F | -0.030 | 0.811 | |
| negXMID.F | RESmT2.F | 0.310 | 0.010 | * |
| ASYM.H | RESmT2.F | -0.190 | 0.123 | |
| LA.H | RESmT2.F | -0.230 | 0.059 | |
| LDI.H | RESmT2.F | -0.070 | 0.569 | |
| LDMC.H | RESmT2.F | -0.280 | 0.020 | * |
| MGR.H | RESmT2.F | 0.260 | 0.034 | * |
| RESmT2.F | RESpT2.H | 0.070 | 0.592 | |
| RESmT2.F | SLA.H | 0.230 | 0.062 | |
| RESmT2.F | TOL_ASYM.H | -0.280 | 0.021 | * |
| RESmT2.F | TOL_MGR.H | 0.050 | 0.664 | |
| RESmT2.F | TOL_negXMID.H | -0.010 | 0.955 | |
| negXMID.H | RESmT2.F | 0.340 | 0.004 | * |
| IGR.C | RESmT2.F | -0.100 | 0.409 | |
| LTH.C | RESmT2.F | -0.160 | 0.181 | |
| NLEA.C | RESmT2.F | 0.000 | 0.983 | |
| RESmT1.C | RESmT2.F | 0.220 | 0.066 | |
| RESmT2.C | RESmT2.F | 0.110 | 0.377 | |
| RESmT2.F | RESpT1.C | -0.010 | 0.925 | |
| RESmT2.F | RESpT2.C | 0.080 | 0.502 | |
| IGR.F | RESmT2.F | 0.050 | 0.662 | |
| LTH.F | RESmT2.F | 0.050 | 0.654 | |
| NLEA.F | RESmT2.F | 0.040 | 0.724 | |
| RESmT2.F | TOL_IGR.F | 0.130 | 0.288 | |
| IGR.H | RESmT2.F | 0.130 | 0.286 | |
| LTH.H | RESmT2.F | 0.150 | 0.234 | |
| NLEA.H | RESmT2.F | -0.030 | 0.786 | |

| Trait_1 | Trait_2 | corr.coef | p-value | |
|---|---|---|---|---|
| RESmT2.F | TOL_IGR.H | 0.090 | 0.452 | |
| ASYM.C | SLA.F | -0.290 | 0.017 | * |
| LA.C | SLA.F | -0.420 | 0.000 | * |
| LDI.C | SLA.F | 0.060 | 0.624 | |
| LDMC.C | SLA.F | -0.460 | 0.000 | * |
| MGR.C | SLA.F | -0.150 | 0.211 | |
| SLA.C | SLA.F | 0.800 | 0.000 | * |
| SLA.F | SSIZ.C | -0.610 | 0.000 | * |
| SLA.F | TGER.C | 0.070 | 0.587 | |
| negXMID.C | SLA.F | -0.100 | 0.405 | |
| ASYM.F | SLA.F | -0.350 | 0.003 | * |
| LA.F | SLA.F | -0.440 | 0.000 | * |
| LDI.F | SLA.F | 0.070 | 0.572 | |
| LDMC.F | SLA.F | -0.500 | 0.000 | * |
| MGR.F | SLA.F | 0.190 | 0.117 | |
| RESmT2.F | SLA.F | 0.260 | 0.032 | * |
| SLA.F | TOL_ASYM.F | 0.050 | 0.692 | |
| SLA.F | TOL_MGR.F | 0.150 | 0.226 | |
| SLA.F | TOL_negXMID.F | 0.140 | 0.249 | |
| negXMID.F | SLA.F | 0.090 | 0.481 | |
| ASYM.H | SLA.F | -0.320 | 0.007 | * |
| LA.H | SLA.F | -0.390 | 0.001 | * |
| LDI.H | SLA.F | -0.020 | 0.851 | |
| LDMC.H | SLA.F | -0.380 | 0.001 | * |
| MGR.H | SLA.F | 0.150 | 0.234 | |
| RESpT2.H | SLA.F | 0.060 | 0.651 | |
| SLA.F | SLA.H | 0.760 | 0.000 | * |
| SLA.F | TOL_ASYM.H | -0.110 | 0.389 | |
| SLA.F | TOL_MGR.H | 0.070 | 0.563 | |
| SLA.F | TOL_negXMID.H | 0.030 | 0.822 | |
| negXMID.H | SLA.F | 0.190 | 0.123 | |
| IGR.C | SLA.F | -0.030 | 0.838 | |
| LTH.C | SLA.F | -0.470 | 0.000 | * |
| NLEA.C | SLA.F | 0.430 | 0.000 | * |
| RESmT1.C | SLA.F | -0.410 | 0.001 | * |
| RESmT2.C | SLA.F | -0.070 | 0.579 | |
| RESpT1.C | SLA.F | -0.490 | 0.000 | * |
| RESpT2.C | SLA.F | 0.070 | 0.559 | |
| IGR.F | SLA.F | -0.020 | 0.858 | |
| LTH.F | SLA.F | -0.310 | 0.009 | * |
| NLEA.F | SLA.F | 0.380 | 0.001 | * |
| SLA.F | TOL_IGR.F | 0.060 | 0.647 | |
| IGR.H | SLA.F | -0.240 | 0.045 | * |
| LTH.H | SLA.F | -0.170 | 0.169 | |
| NLEA.H | SLA.F | 0.440 | 0.000 | * |
| SLA.F | TOL_IGR.H | -0.060 | 0.649 | |
| ASYM.C | TOL_ASYM.F | -0.080 | 0.507 | |
| LA.C | TOL_ASYM.F | 0.070 | 0.575 | |
| LDI.C | TOL_ASYM.F | 0.020 | 0.855 | |

| Trait_1 | Trait_2 | corr.coef | p-value | |
|---|---|---|---|---|
| LDMC.C | TOL_ASYM.F | 0.090 | 0.468 | |
| MGR.C | TOL_ASYM.F | 0.130 | 0.271 | |
| SLA.C | TOL_ASYM.F | 0.080 | 0.528 | |
| SSIZ.C | TOL_ASYM.F | 0.000 | 0.995 | |
| TGER.C | TOL_ASYM.F | 0.030 | 0.836 | |
| negXMID.C | TOL_ASYM.F | 0.290 | 0.016 | * |
| ASYM.F | TOL_ASYM.F | 0.210 | 0.081 | |
| LA.F | TOL_ASYM.F | 0.160 | 0.194 | |
| LDI.F | TOL_ASYM.F | 0.090 | 0.442 | |
| LDMC.F | TOL_ASYM.F | 0.090 | 0.444 | |
| MGR.F | TOL_ASYM.F | 0.010 | 0.960 | |
| RESmT2.F | TOL_ASYM.F | 0.070 | 0.595 | |
| SLA.F | TOL_ASYM.F | 0.050 | 0.692 | |
| TOL_ASYM.F | TOL_MGR.F | 0.090 | 0.444 | |
| TOL_ASYM.F | TOL_negXMID.F | -0.520 | 0.000 | * |
| negXMID.F | TOL_ASYM.F | -0.120 | 0.333 | |
| ASYM.H | TOL_ASYM.F | 0.030 | 0.789 | |
| LA.H | TOL_ASYM.F | 0.080 | 0.527 | |
| LDI.H | TOL_ASYM.F | 0.050 | 0.656 | |
| LDMC.H | TOL_ASYM.F | 0.020 | 0.858 | |
| MGR.H | TOL_ASYM.F | -0.050 | 0.697 | |
| RESpT2.H | TOL_ASYM.F | 0.090 | 0.456 | |
| SLA.H | TOL_ASYM.F | -0.010 | 0.963 | |
| TOL_ASYM.F | TOL_ASYM.H | 0.060 | 0.600 | |
| TOL_ASYM.F | TOL_MGR.H | -0.130 | 0.292 | |
| TOL_ASYM.F | TOL_negXMID.H | -0.040 | 0.737 | |
| negXMID.H | TOL_ASYM.F | 0.110 | 0.385 | |
| IGR.C | TOL_ASYM.F | 0.170 | 0.161 | |
| LTH.C | TOL_ASYM.F | 0.010 | 0.945 | |
| NLEA.C | TOL_ASYM.F | 0.090 | 0.447 | |
| RESmT1.C | TOL_ASYM.F | -0.100 | 0.424 | |
| RESmT2.C | TOL_ASYM.F | -0.070 | 0.566 | |
| RESpT1.C | TOL_ASYM.F | -0.110 | 0.364 | |
| RESpT2.C | TOL_ASYM.F | 0.250 | 0.041 | * |
| IGR.F | TOL_ASYM.F | -0.110 | 0.378 | |
| LTH.F | TOL_ASYM.F | -0.140 | 0.243 | |
| NLEA.F | TOL_ASYM.F | 0.150 | 0.228 | |
| TOL_ASYM.F | TOL_IGR.F | -0.200 | 0.097 | |
| IGR.H | TOL_ASYM.F | -0.210 | 0.086 | |
| LTH.H | TOL_ASYM.F | -0.070 | 0.553 | |
| NLEA.H | TOL_ASYM.F | 0.130 | 0.288 | |
| TOL_ASYM.F | TOL_IGR.H | -0.070 | 0.588 | |
| ASYM.C | TOL_MGR.F | -0.130 | 0.274 | |
| LA.C | TOL_MGR.F | -0.070 | 0.544 | |
| LDI.C | TOL_MGR.F | -0.080 | 0.527 | |
| LDMC.C | TOL_MGR.F | -0.030 | 0.827 | |
| MGR.C | TOL_MGR.F | -0.350 | 0.003 | * |
| SLA.C | TOL_MGR.F | 0.150 | 0.219 | |
| SSIZ.C | TOL_MGR.F | -0.110 | 0.358 | |

| Trait_1 | Trait_2 | corr.coef | p-value | |
|---|---|---|---|---|
| TGER.C | TOL_MGR.F | 0.000 | 0.970 | |
| negXMID.C | TOL_MGR.F | 0.030 | 0.837 | |
| ASYM.F | TOL_MGR.F | -0.100 | 0.410 | |
| LA.F | TOL_MGR.F | -0.040 | 0.738 | |
| LDI.F | TOL_MGR.F | -0.030 | 0.793 | |
| LDMC.F | TOL_MGR.F | 0.060 | 0.604 | |
| MGR.F | TOL_MGR.F | 0.350 | 0.003 | * |
| RESmT2.F | TOL_MGR.F | 0.080 | 0.490 | |
| SLA.F | TOL_MGR.F | 0.150 | 0.226 | |
| TOL_ASYM.F | TOL_MGR.F | 0.090 | 0.444 | |
| TOL_MGR.F | TOL_negXMID.F | 0.010 | 0.953 | |
| negXMID.F | TOL_MGR.F | 0.000 | 0.989 | |
| ASYM.H | TOL_MGR.F | -0.080 | 0.539 | |
| LA.H | TOL_MGR.F | -0.010 | 0.911 | |
| LDI.H | TOL_MGR.F | -0.040 | 0.739 | |
| LDMC.H | TOL_MGR.F | 0.060 | 0.651 | |
| MGR.H | TOL_MGR.F | 0.000 | 0.986 | |
| RESpT2.H | TOL_MGR.F | -0.080 | 0.493 | |
| SLA.H | TOL_MGR.F | 0.270 | 0.025 | * |
| TOL_ASYM.H | TOL_MGR.F | -0.090 | 0.468 | |
| TOL_MGR.F | TOL_MGR.H | 0.100 | 0.432 | |
| TOL_MGR.F | TOL_negXMID.H | 0.030 | 0.827 | |
| negXMID.H | TOL_MGR.F | 0.010 | 0.917 | |
| IGR.C | TOL_MGR.F | -0.070 | 0.554 | |
| LTH.C | TOL_MGR.F | -0.020 | 0.881 | |
| NLEA.C | TOL_MGR.F | 0.000 | 0.982 | |
| RESmT1.C | TOL_MGR.F | -0.190 | 0.113 | |
| RESmT2.C | TOL_MGR.F | 0.060 | 0.629 | |
| RESpT1.C | TOL_MGR.F | -0.070 | 0.553 | |
| RESpT2.C | TOL_MGR.F | -0.010 | 0.950 | |
| IGR.F | TOL_MGR.F | 0.060 | 0.643 | |
| LTH.F | TOL_MGR.F | -0.090 | 0.471 | |
| NLEA.F | TOL_MGR.F | -0.010 | 0.913 | |
| TOL_IGR.F | TOL_MGR.F | 0.060 | 0.619 | |
| IGR.H | TOL_MGR.F | -0.120 | 0.329 | |
| LTH.H | TOL_MGR.F | -0.030 | 0.796 | |
| NLEA.H | TOL_MGR.F | -0.060 | 0.637 | |
| TOL_IGR.H | TOL_MGR.F | -0.170 | 0.157 | |
| ASYM.C | TOL_negXMID.F | -0.010 | 0.929 | |
| LA.C | TOL_negXMID.F | -0.180 | 0.137 | |
| LDI.C | TOL_negXMID.F | 0.020 | 0.843 | |
| LDMC.C | TOL_negXMID.F | 0.030 | 0.805 | |
| MGR.C | TOL_negXMID.F | -0.100 | 0.423 | |
| SLA.C | TOL_negXMID.F | 0.190 | 0.110 | |
| SSIZ.C | TOL_negXMID.F | -0.220 | 0.070 | |
| TGER.C | TOL_negXMID.F | -0.100 | 0.431 | |
| negXMID.C | TOL_negXMID.F | -0.260 | 0.029 | * |
| ASYM.F | TOL_negXMID.F | -0.130 | 0.274 | |
| LA.F | TOL_negXMID.F | -0.110 | 0.348 | |

| Trait_1 | Trait_2 | corr.coef | p-value | |
|---|---|---|---|---|
| LDI.F | TOL_negXMID.F | 0.010 | 0.960 | |
| LDMC.F | TOL_negXMID.F | 0.030 | 0.828 | |
| MGR.F | TOL_negXMID.F | -0.010 | 0.940 | |
| RESmT2.F | TOL_negXMID.F | -0.030 | 0.811 | |
| SLA.F | TOL_negXMID.F | 0.140 | 0.249 | |
| TOL_ASYM.F | TOL_negXMID.F | -0.520 | 0.000 | * |
| TOL_MGR.F | TOL_negXMID.F | 0.010 | 0.953 | |
| negXMID.F | TOL_negXMID.F | 0.290 | 0.016 | * |
| ASYM.H | TOL_negXMID.F | -0.080 | 0.522 | |
| LA.H | TOL_negXMID.F | -0.130 | 0.277 | |
| LDI.H | TOL_negXMID.F | 0.020 | 0.868 | |
| LDMC.H | TOL_negXMID.F | 0.110 | 0.367 | |
| MGR.H | TOL_negXMID.F | 0.030 | 0.826 | |
| RESpT2.H | TOL_negXMID.F | -0.030 | 0.787 | |
| SLA.H | TOL_negXMID.F | 0.170 | 0.155 | |
| TOL_ASYM.H | TOL_negXMID.F | -0.090 | 0.480 | |
| TOL_MGR.H | TOL_negXMID.F | 0.130 | 0.296 | |
| TOL_negXMID.F | TOL_negXMID.H | -0.160 | 0.182 | |
| negXMID.H | TOL_negXMID.F | -0.190 | 0.126 | |
| IGR.C | TOL_negXMID.F | -0.030 | 0.835 | |
| LTH.C | TOL_negXMID.F | -0.080 | 0.526 | |
| NLEA.C | TOL_negXMID.F | 0.040 | 0.731 | |
| RESmT1.C | TOL_negXMID.F | -0.120 | 0.317 | |
| RESmT2.C | TOL_negXMID.F | -0.150 | 0.211 | |
| RESpT1.C | TOL_negXMID.F | -0.160 | 0.180 | |
| RESpT2.C | TOL_negXMID.F | 0.000 | 0.986 | |
| IGR.F | TOL_negXMID.F | 0.050 | 0.704 | |
| LTH.F | TOL_negXMID.F | -0.150 | 0.222 | |
| NLEA.F | TOL_negXMID.F | 0.050 | 0.683 | |
| TOL_IGR.F | TOL_negXMID.F | 0.110 | 0.381 | |
| IGR.H | TOL_negXMID.F | -0.060 | 0.618 | |
| LTH.H | TOL_negXMID.F | 0.070 | 0.551 | |
| NLEA.H | TOL_negXMID.F | 0.020 | 0.854 | |
| TOL_IGR.H | TOL_negXMID.F | -0.070 | 0.590 | |
| ASYM.C | negXMID.F | -0.030 | 0.814 | |
| LA.C | negXMID.F | -0.070 | 0.593 | |
| LDI.C | negXMID.F | -0.060 | 0.648 | |
| LDMC.C | negXMID.F | 0.080 | 0.515 | |
| MGR.C | negXMID.F | -0.080 | 0.531 | |
| negXMID.F | SLA.C | 0.140 | 0.258 | |
| negXMID.F | SSIZ.C | 0.150 | 0.220 | |
| negXMID.F | TGER.C | 0.370 | 0.002 | * |
| negXMID.C | negXMID.F | 0.440 | 0.000 | * |
| ASYM.F | negXMID.F | -0.130 | 0.294 | |
| LA.F | negXMID.F | -0.010 | 0.903 | |
| LDI.F | negXMID.F | -0.100 | 0.428 | |
| LDMC.F | negXMID.F | -0.040 | 0.716 | |
| MGR.F | negXMID.F | 0.210 | 0.089 | |
| negXMID.F | RESmT2.F | 0.310 | 0.010 | * |

| Trait_1 | Trait_2 | corr.coef | p-value | |
|---|---|---|---|---|
| negXMID.F | SLA.F | 0.090 | 0.481 | |
| negXMID.F | TOL_ASYM.F | -0.120 | 0.333 | |
| negXMID.F | TOL_MGR.F | 0.000 | 0.989 | |
| negXMID.F | TOL_negXMID.F | 0.290 | 0.016 | * |
| ASYM.H | negXMID.F | -0.020 | 0.889 | |
| LA.H | negXMID.F | 0.010 | 0.928 | |
| LDI.H | negXMID.F | -0.020 | 0.891 | |
| LDMC.H | negXMID.F | 0.030 | 0.832 | |
| MGR.H | negXMID.F | 0.030 | 0.816 | |
| negXMID.F | RESpT2.H | 0.080 | 0.494 | |
| negXMID.F | SLA.H | 0.170 | 0.161 | |
| negXMID.F | TOL_ASYM.H | -0.150 | 0.230 | |
| negXMID.F | TOL_MGR.H | -0.100 | 0.433 | |
| negXMID.F | TOL_negXMID.H | 0.040 | 0.723 | |
| negXMID.F | negXMID.H | 0.470 | 0.000 | * |
| IGR.C | negXMID.F | -0.130 | 0.276 | |
| LTH.C | negXMID.F | -0.040 | 0.727 | |
| negXMID.F | NLEA.C | -0.020 | 0.865 | |
| negXMID.F | RESmT1.C | -0.010 | 0.910 | |
| negXMID.F | RESmT2.C | -0.040 | 0.720 | |
| negXMID.F | RESpT1.C | -0.030 | 0.805 | |
| negXMID.F | RESpT2.C | -0.080 | 0.521 | |
| IGR.F | negXMID.F | -0.180 | 0.137 | |
| LTH.F | negXMID.F | -0.100 | 0.405 | |
| negXMID.F | NLEA.F | 0.020 | 0.891 | |
| negXMID.F | TOL_IGR.F | 0.090 | 0.463 | |
| IGR.H | negXMID.F | 0.120 | 0.346 | |
| LTH.H | negXMID.F | 0.020 | 0.900 | |
| negXMID.F | NLEA.H | 0.040 | 0.759 | |
| negXMID.F | TOL_IGR.H | 0.150 | 0.211 | |
| ASYM.C | ASYM.H | 0.880 | 0.000 | * |
| ASYM.H | LA.C | 0.870 | 0.000 | * |
| ASYM.H | LDI.C | 0.650 | 0.000 | * |
| ASYM.H | LDMC.C | 0.530 | 0.000 | * |
| ASYM.H | MGR.C | 0.260 | 0.030 | * |
| ASYM.H | SLA.C | -0.290 | 0.015 | * |
| ASYM.H | SSIZ.C | 0.120 | 0.338 | |
| ASYM.H | TGER.C | -0.260 | 0.033 | * |
| ASYM.H | negXMID.C | 0.110 | 0.377 | |
| ASYM.F | ASYM.H | 0.900 | 0.000 | * |
| ASYM.H | LA.F | 0.850 | 0.000 | * |
| ASYM.H | LDI.F | 0.610 | 0.000 | * |
| ASYM.H | LDMC.F | 0.500 | 0.000 | * |
| ASYM.H | MGR.F | -0.010 | 0.927 | |
| ASYM.H | RESmT2.F | -0.190 | 0.123 | |
| ASYM.H | SLA.F | -0.320 | 0.007 | * |
| ASYM.H | TOL_ASYM.F | 0.030 | 0.789 | |
| ASYM.H | TOL_MGR.F | -0.080 | 0.539 | |
| ASYM.H | TOL_negXMID.F | -0.080 | 0.522 | |

| Trait_1 | Trait_2 | corr.coef | p-value | |
|---|---|---|---|---|
| ASYM.H | negXMID.F | -0.020 | 0.889 | |
| ASYM.H | LA.H | 0.890 | 0.000 | * |
| ASYM.H | LDI.H | 0.730 | 0.000 | * |
| ASYM.H | LDMC.H | 0.480 | 0.000 | * |
| ASYM.H | MGR.H | -0.090 | 0.449 | |
| ASYM.H | RESpT2.H | -0.280 | 0.019 | * |
| ASYM.H | SLA.H | -0.430 | 0.000 | * |
| ASYM.H | TOL_ASYM.H | 0.700 | 0.000 | * |
| ASYM.H | TOL_MGR.H | 0.030 | 0.794 | |
| ASYM.H | TOL_negXMID.H | -0.380 | 0.001 | * |
| ASYM.H | negXMID.H | -0.380 | 0.001 | * |
| ASYM.H | IGR.C | 0.230 | 0.058 | |
| ASYM.H | LTH.C | 0.050 | 0.687 | |
| ASYM.H | NLEA.C | -0.360 | 0.003 | * |
| ASYM.H | RESmT1.C | 0.080 | 0.494 | |
| ASYM.H | RESmT2.C | 0.210 | 0.088 | |
| ASYM.H | RESpT1.C | 0.210 | 0.079 | |
| ASYM.H | RESpT2.C | 0.100 | 0.393 | |
| ASYM.H | IGR.F | 0.100 | 0.426 | |
| ASYM.H | LTH.F | -0.150 | 0.206 | |
| ASYM.H | NLEA.F | -0.330 | 0.005 | * |
| ASYM.H | TOL_IGR.F | -0.210 | 0.079 | |
| ASYM.H | IGR.H | 0.040 | 0.714 | |
| ASYM.H | LTH.H | 0.110 | 0.371 | |
| ASYM.H | NLEA.H | -0.300 | 0.013 | * |
| ASYM.H | TOL_IGR.H | 0.110 | 0.369 | |
| ASYM.C | LA.H | 0.780 | 0.000 | * |
| LA.C | LA.H | 0.930 | 0.000 | * |
| LA.H | LDI.C | 0.440 | 0.000 | * |
| LA.H | LDMC.C | 0.540 | 0.000 | * |
| LA.H | MGR.C | 0.150 | 0.230 | |
| LA.H | SLA.C | -0.350 | 0.003 | * |
| LA.H | SSIZ.C | 0.210 | 0.086 | |
| LA.H | TGER.C | -0.190 | 0.111 | |
| LA.H | negXMID.C | 0.170 | 0.153 | |
| ASYM.F | LA.H | 0.820 | 0.000 | * |
| LA.F | LA.H | 0.940 | 0.000 | * |
| LA.H | LDI.F | 0.430 | 0.000 | * |
| LA.H | LDMC.F | 0.460 | 0.000 | * |
| LA.H | MGR.F | 0.060 | 0.648 | |
| LA.H | RESmT2.F | -0.230 | 0.059 | |
| LA.H | SLA.F | -0.390 | 0.001 | * |
| LA.H | TOL_ASYM.F | 0.080 | 0.527 | |
| LA.H | TOL_MGR.F | -0.010 | 0.911 | |
| LA.H | TOL_negXMID.F | -0.130 | 0.277 | |
| LA.H | negXMID.F | 0.010 | 0.928 | |
| ASYM.H | LA.H | 0.890 | 0.000 | * |
| LA.H | LDI.H | 0.590 | 0.000 | * |
| LA.H | LDMC.H | 0.500 | 0.000 | * |

| Trait_1 | Trait_2 | corr.coef | p-value | |
|---|---|---|---|---|
| LA.H | MGR.H | -0.080 | 0.502 | |
| LA.H | RESpT2.H | -0.160 | 0.200 | |
| LA.H | SLA.H | -0.480 | 0.000 | * |
| LA.H | TOL_ASYM.H | 0.600 | 0.000 | * |
| LA.H | TOL_MGR.H | 0.030 | 0.783 | |
| LA.H | TOL_negXMID.H | -0.330 | 0.005 | * |
| LA.H | negXMID.H | -0.310 | 0.010 | * |
| IGR.C | LA.H | 0.290 | 0.014 | * |
| LA.H | LTH.C | 0.180 | 0.133 | |
| LA.H | NLEA.C | -0.470 | 0.000 | * |
| LA.H | RESmT1.C | 0.140 | 0.251 | |
| LA.H | RESmT2.C | 0.160 | 0.198 | |
| LA.H | RESpT1.C | 0.290 | 0.016 | * |
| LA.H | RESpT2.C | 0.060 | 0.650 | |
| IGR.F | LA.H | 0.050 | 0.680 | |
| LA.H | LTH.F | -0.060 | 0.641 | |
| LA.H | NLEA.F | -0.460 | 0.000 | * |
| LA.H | TOL_IGR.F | -0.260 | 0.033 | * |
| IGR.H | LA.H | 0.040 | 0.742 | |
| LA.H | LTH.H | 0.100 | 0.411 | |
| LA.H | NLEA.H | -0.430 | 0.000 | * |
| LA.H | TOL_IGR.H | 0.030 | 0.822 | |
| ASYM.C | LDI.H | 0.670 | 0.000 | * |
| LA.C | LDI.H | 0.610 | 0.000 | * |
| LDI.C | LDI.H | 0.920 | 0.000 | * |
| LDI.H | LDMC.C | 0.390 | 0.001 | * |
| LDI.H | MGR.C | 0.060 | 0.626 | |
| LDI.H | SLA.C | -0.080 | 0.512 | |
| LDI.H | SSIZ.C | -0.150 | 0.228 | |
| LDI.H | TGER.C | -0.170 | 0.151 | |
| LDI.H | negXMID.C | -0.010 | 0.934 | |
| ASYM.F | LDI.H | 0.710 | 0.000 | * |
| LA.F | LDI.H | 0.540 | 0.000 | * |
| LDI.F | LDI.H | 0.890 | 0.000 | * |
| LDI.H | LDMC.F | 0.360 | 0.002 | * |
| LDI.H | MGR.F | -0.050 | 0.710 | |
| LDI.H | RESmT2.F | -0.070 | 0.569 | |
| LDI.H | SLA.F | -0.020 | 0.851 | |
| LDI.H | TOL_ASYM.F | 0.050 | 0.656 | |
| LDI.H | TOL_MGR.F | -0.040 | 0.739 | |
| LDI.H | TOL_negXMID.F | 0.020 | 0.868 | |
| LDI.H | negXMID.F | -0.020 | 0.891 | |
| ASYM.H | LDI.H | 0.730 | 0.000 | * |
| LA.H | LDI.H | 0.590 | 0.000 | * |
| LDI.H | LDMC.H | 0.380 | 0.001 | * |
| LDI.H | MGR.H | -0.010 | 0.908 | |
| LDI.H | RESpT2.H | -0.100 | 0.425 | |
| LDI.H | SLA.H | -0.150 | 0.230 | |
| LDI.H | TOL_ASYM.H | 0.510 | 0.000 | * |

| Trait_1 | Trait_2 | corr.coef | p-value | |
|---|---|---|---|---|
| LDI.H | TOL_MGR.H | 0.130 | 0.305 | |
| LDI.H | TOL_negXMID.H | -0.430 | 0.000 | * |
| LDI.H | negXMID.H | -0.340 | 0.004 | * |
| IGR.C | LDI.H | 0.240 | 0.045 | * |
| LDI.H | LTH.C | -0.140 | 0.243 | |
| LDI.H | NLEA.C | -0.060 | 0.620 | |
| LDI.H | RESmT1.C | -0.130 | 0.305 | |
| LDI.H | RESmT2.C | 0.130 | 0.303 | |
| LDI.H | RESpT1.C | -0.030 | 0.777 | |
| LDI.H | RESpT2.C | 0.020 | 0.867 | |
| IGR.F | LDI.H | 0.030 | 0.827 | |
| LDI.H | LTH.F | -0.330 | 0.006 | * |
| LDI.H | NLEA.F | -0.040 | 0.746 | |
| LDI.H | TOL_IGR.F | -0.220 | 0.065 | |
| IGR.H | LDI.H | -0.150 | 0.227 | |
| LDI.H | LTH.H | 0.010 | 0.928 | |
| LDI.H | NLEA.H | -0.040 | 0.717 | |
| LDI.H | TOL_IGR.H | -0.050 | 0.703 | |
| ASYM.C | LDMC.H | 0.440 | 0.000 | * |
| LA.C | LDMC.H | 0.480 | 0.000 | * |
| LDI.C | LDMC.H | 0.340 | 0.004 | * |
| LDMC.C | LDMC.H | 0.790 | 0.000 | * |
| LDMC.H | MGR.C | 0.090 | 0.452 | |
| LDMC.H | SLA.C | -0.300 | 0.012 | * |
| LDMC.H | SSIZ.C | 0.170 | 0.153 | |
| LDMC.H | TGER.C | -0.140 | 0.251 | |
| LDMC.H | negXMID.C | 0.080 | 0.509 | |
| ASYM.F | LDMC.H | 0.510 | 0.000 | * |
| LA.F | LDMC.H | 0.520 | 0.000 | * |
| LDI.F | LDMC.H | 0.320 | 0.007 | * |
| LDMC.F | LDMC.H | 0.730 | 0.000 | * |
| LDMC.H | MGR.F | 0.040 | 0.770 | |
| LDMC.H | RESmT2.F | -0.280 | 0.020 | * |
| LDMC.H | SLA.F | -0.380 | 0.001 | * |
| LDMC.H | TOL_ASYM.F | 0.020 | 0.858 | |
| LDMC.H | TOL_MGR.F | 0.060 | 0.651 | |
| LDMC.H | TOL_negXMID.F | 0.110 | 0.367 | |
| LDMC.H | negXMID.F | 0.030 | 0.832 | |
| ASYM.H | LDMC.H | 0.480 | 0.000 | * |
| LA.H | LDMC.H | 0.500 | 0.000 | * |
| LDI.H | LDMC.H | 0.380 | 0.001 | * |
| LDMC.H | MGR.H | -0.030 | 0.817 | |
| LDMC.H | RESpT2.H | 0.220 | 0.067 | |
| LDMC.H | SLA.H | -0.600 | 0.000 | * |
| LDMC.H | TOL_ASYM.H | 0.340 | 0.004 | * |
| LDMC.H | TOL_MGR.H | 0.080 | 0.498 | |
| LDMC.H | TOL_negXMID.H | -0.240 | 0.046 | * |
| LDMC.H | negXMID.H | -0.200 | 0.104 | |
| IGR.C | LDMC.H | 0.030 | 0.790 | |

| **Trait_1** | **Trait_2** | **corr.coef** | **p-value** | | **Trait_1** | **Trait_2** | **corr.coef** | **p-value** | |
|---|---|---|---|---|---|---|---|---|---|
| LDMC.H | LTH.C | -0.120 | 0.344 | | MGR.H | RESpT1.C | -0.020 | 0.851 | |
| LDMC.H | NLEA.C | -0.220 | 0.067 | | MGR.H | RESpT2.C | 0.070 | 0.590 | |
| LDMC.H | RESmT1.C | -0.200 | 0.104 | | IGR.F | MGR.H | -0.130 | 0.280 | |
| LDMC.H | RESmT2.C | 0.120 | 0.321 | | LTH.F | MGR.H | 0.010 | 0.946 | |
| LDMC.H | RESpT1.C | 0.030 | 0.813 | | MGR.H | NLEA.F | -0.110 | 0.356 | |
| LDMC.H | RESpT2.C | 0.350 | 0.003 | * | MGR.H | TOL_IGR.F | -0.020 | 0.901 | |
| IGR.F | LDMC.H | -0.220 | 0.064 | | IGR.H | MGR.H | 0.030 | 0.789 | |
| LDMC.H | LTH.F | -0.310 | 0.010 | * | LTH.H | MGR.H | 0.110 | 0.385 | |
| LDMC.H | NLEA.F | -0.160 | 0.190 | | MGR.H | NLEA.H | -0.200 | 0.102 | |
| LDMC.H | TOL_IGR.F | -0.150 | 0.210 | | MGR.H | TOL_IGR.H | 0.150 | 0.222 | |
| IGR.H | LDMC.H | 0.010 | 0.932 | | ASYM.C | RESpT2.H | -0.220 | 0.076 | |
| LDMC.H | LTH.H | 0.100 | 0.418 | | LA.C | RESpT2.H | -0.190 | 0.126 | |
| LDMC.H | NLEA.H | -0.160 | 0.176 | | LDI.C | RESpT2.H | -0.100 | 0.424 | |
| LDMC.H | TOL_IGR.H | 0.030 | 0.836 | | LDMC.C | RESpT2.H | 0.150 | 0.221 | |
| ASYM.C | MGR.H | -0.050 | 0.659 | | MGR.C | RESpT2.H | -0.130 | 0.284 | |
| LA.C | MGR.H | -0.070 | 0.557 | | RESpT2.H | SLA.C | 0.030 | 0.820 | |
| LDI.C | MGR.H | -0.020 | 0.839 | | RESpT2.H | SSIZ.C | 0.040 | 0.726 | |
| LDMC.C | MGR.H | -0.200 | 0.095 | | RESpT2.H | TGER.C | 0.180 | 0.132 | |
| MGR.C | MGR.H | -0.070 | 0.576 | | negXMID.C | RESpT2.H | -0.080 | 0.534 | |
| MGR.H | SLA.C | 0.040 | 0.752 | | ASYM.F | RESpT2.H | -0.210 | 0.083 | |
| MGR.H | SSIZ.C | -0.070 | 0.552 | | LA.F | RESpT2.H | -0.210 | 0.089 | |
| MGR.H | TGER.C | 0.150 | 0.233 | | LDI.F | RESpT2.H | -0.070 | 0.575 | |
| MGR.H | negXMID.C | -0.190 | 0.125 | | LDMC.F | RESpT2.H | 0.030 | 0.788 | |
| ASYM.F | MGR.H | -0.050 | 0.702 | | MGR.F | RESpT2.H | 0.000 | 0.981 | |
| LA.F | MGR.H | -0.070 | 0.586 | | RESmT2.F | RESpT2.H | 0.070 | 0.592 | |
| LDI.F | MGR.H | -0.020 | 0.850 | | RESpT2.H | SLA.F | 0.060 | 0.651 | |
| LDMC.F | MGR.H | -0.210 | 0.076 | | RESpT2.H | TOL_ASYM.F | 0.090 | 0.456 | |
| MGR.F | MGR.H | 0.090 | 0.462 | | RESpT2.H | TOL_MGR.F | -0.080 | 0.493 | |
| MGR.H | RESmT2.F | 0.260 | 0.034 | * | RESpT2.H | TOL_negXMID.F | -0.030 | 0.787 | |
| MGR.H | SLA.F | 0.150 | 0.234 | | negXMID.F | RESpT2.H | 0.080 | 0.494 | |
| MGR.H | TOL_ASYM.F | -0.050 | 0.697 | | ASYM.H | RESpT2.H | -0.280 | 0.019 | * |
| MGR.H | TOL_MGR.F | 0.000 | 0.986 | | LA.H | RESpT2.H | -0.160 | 0.200 | |
| MGR.H | TOL_negXMID.F | 0.030 | 0.826 | | LDI.H | RESpT2.H | -0.100 | 0.425 | |
| MGR.H | negXMID.F | 0.030 | 0.816 | | LDMC.H | RESpT2.H | 0.220 | 0.067 | |
| ASYM.H | MGR.H | -0.090 | 0.449 | | MGR.H | RESpT2.H | 0.150 | 0.222 | |
| LA.H | MGR.H | -0.080 | 0.502 | | RESpT2.H | SLA.H | -0.080 | 0.520 | |
| LDI.H | MGR.H | -0.010 | 0.908 | | RESpT2.H | TOL_ASYM.H | -0.150 | 0.208 | |
| LDMC.H | MGR.H | -0.030 | 0.817 | | RESpT2.H | TOL_MGR.H | 0.040 | 0.717 | |
| MGR.H | RESpT2.H | 0.150 | 0.222 | | RESpT2.H | TOL_negXMID.H | 0.010 | 0.961 | |
| MGR.H | SLA.H | 0.000 | 0.969 | | negXMID.H | RESpT2.H | 0.150 | 0.228 | |
| MGR.H | TOL_ASYM.H | -0.210 | 0.087 | | IGR.C | RESpT2.H | 0.070 | 0.585 | |
| MGR.H | TOL_MGR.H | 0.710 | 0.000 | * | LTH.C | RESpT2.H | -0.100 | 0.429 | |
| MGR.H | TOL_negXMID.H | -0.130 | 0.288 | | NLEA.C | RESpT2.H | 0.040 | 0.754 | |
| MGR.H | negXMID.H | 0.120 | 0.332 | | RESmT1.C | RESpT2.H | -0.180 | 0.136 | |
| IGR.C | MGR.H | -0.060 | 0.619 | | RESmT2.C | RESpT2.H | -0.070 | 0.564 | |
| LTH.C | MGR.H | 0.100 | 0.412 | | RESpT1.C | RESpT2.H | -0.050 | 0.708 | |
| MGR.H | NLEA.C | -0.160 | 0.195 | | RESpT2.C | RESpT2.H | 0.180 | 0.129 | |
| MGR.H | RESmT1.C | 0.060 | 0.652 | | IGR.F | RESpT2.H | -0.260 | 0.029 | * |
| MGR.H | RESmT2.C | -0.210 | 0.088 | | LTH.F | RESpT2.H | -0.040 | 0.736 | |

| Trait_1 | Trait_2 | corr.coef | p-value | |
|---|---|---|---|---|
| NLEA.F | RESpT2.H | -0.040 | 0.748 | |
| RESpT2.H | TOL_IGR.F | -0.070 | 0.573 | |
| IGR.H | RESpT2.H | -0.040 | 0.748 | |
| LTH.H | RESpT2.H | -0.080 | 0.531 | |
| NLEA.H | RESpT2.H | 0.110 | 0.360 | |
| RESpT2.H | TOL_IGR.H | -0.100 | 0.415 | |
| ASYM.C | SLA.H | -0.390 | 0.001 | * |
| LA.C | SLA.H | -0.520 | 0.000 | * |
| LDI.C | SLA.H | -0.110 | 0.352 | |
| LDMC.C | SLA.H | -0.510 | 0.000 | * |
| MGR.C | SLA.H | -0.270 | 0.027 | * |
| SLA.C | SLA.H | 0.680 | 0.000 | * |
| SLA.H | SSIZ.C | -0.470 | 0.000 | * |
| SLA.H | TGER.C | 0.040 | 0.766 | |
| negXMID.C | SLA.H | -0.050 | 0.689 | |
| ASYM.F | SLA.H | -0.460 | 0.000 | * |
| LA.F | SLA.H | -0.560 | 0.000 | * |
| LDI.F | SLA.H | -0.110 | 0.349 | |
| LDMC.F | SLA.H | -0.430 | 0.000 | * |
| MGR.F | SLA.H | 0.120 | 0.320 | |
| RESmT2.F | SLA.H | 0.230 | 0.062 | |
| SLA.F | SLA.H | 0.760 | 0.000 | * |
| SLA.H | TOL_ASYM.F | -0.010 | 0.963 | |
| SLA.H | TOL_MGR.F | 0.270 | 0.025 | * |
| SLA.H | TOL_negXMID.F | 0.170 | 0.155 | |
| negXMID.F | SLA.H | 0.170 | 0.161 | |
| ASYM.H | SLA.H | -0.430 | 0.000 | * |
| LA.H | SLA.H | -0.480 | 0.000 | * |
| LDI.H | SLA.H | -0.150 | 0.230 | |
| LDMC.H | SLA.H | -0.600 | 0.000 | * |
| MGR.H | SLA.H | 0.000 | 0.969 | |
| RESpT2.H | SLA.H | -0.080 | 0.520 | |
| SLA.H | TOL_ASYM.H | -0.260 | 0.030 | * |
| SLA.H | TOL_MGR.H | -0.060 | 0.629 | |
| SLA.H | TOL_negXMID.H | 0.130 | 0.275 | |
| negXMID.H | SLA.H | 0.190 | 0.125 | |
| IGR.C | SLA.H | 0.010 | 0.960 | |
| LTH.C | SLA.H | -0.250 | 0.040 | * |
| NLEA.C | SLA.H | 0.530 | 0.000 | * |
| RESmT1.C | SLA.H | -0.230 | 0.059 | |
| RESmT2.C | SLA.H | 0.030 | 0.797 | |
| RESpT1.C | SLA.H | -0.280 | 0.018 | * |
| RESpT2.C | SLA.H | -0.080 | 0.522 | |
| IGR.F | SLA.H | -0.010 | 0.921 | |
| LTH.F | SLA.H | -0.150 | 0.226 | |
| NLEA.F | SLA.H | 0.470 | 0.000 | * |
| SLA.H | TOL_IGR.F | 0.060 | 0.646 | |
| IGR.H | SLA.H | -0.090 | 0.440 | |
| LTH.H | SLA.H | -0.240 | 0.046 | * |

| Trait_1 | Trait_2 | corr.coef | p-value | |
|---|---|---|---|---|
| NLEA.H | SLA.H | 0.490 | 0.000 | * |
| SLA.H | TOL_IGR.H | -0.020 | 0.892 | |
| ASYM.C | TOL_ASYM.H | 0.570 | 0.000 | * |
| LA.C | TOL_ASYM.H | 0.560 | 0.000 | * |
| LDI.C | TOL_ASYM.H | 0.430 | 0.000 | * |
| LDMC.C | TOL_ASYM.H | 0.350 | 0.003 | * |
| MGR.C | TOL_ASYM.H | 0.150 | 0.209 | |
| SLA.C | TOL_ASYM.H | -0.060 | 0.639 | |
| SSIZ.C | TOL_ASYM.H | 0.050 | 0.686 | |
| TGER.C | TOL_ASYM.H | -0.310 | 0.009 | * |
| negXMID.C | TOL_ASYM.H | -0.020 | 0.886 | |
| ASYM.F | TOL_ASYM.H | 0.610 | 0.000 | * |
| LA.F | TOL_ASYM.H | 0.540 | 0.000 | * |
| LDI.F | TOL_ASYM.H | 0.390 | 0.001 | * |
| LDMC.F | TOL_ASYM.H | 0.380 | 0.001 | * |
| MGR.F | TOL_ASYM.H | 0.000 | 0.992 | |
| RESmT2.F | TOL_ASYM.H | -0.280 | 0.021 | * |
| SLA.F | TOL_ASYM.H | -0.110 | 0.389 | |
| TOL_ASYM.F | TOL_ASYM.H | 0.060 | 0.600 | |
| TOL_ASYM.H | TOL_MGR.F | -0.090 | 0.468 | |
| TOL_ASYM.H | TOL_negXMID.F | -0.090 | 0.480 | |
| negXMID.F | TOL_ASYM.H | -0.150 | 0.230 | |
| ASYM.H | TOL_ASYM.H | 0.700 | 0.000 | * |
| LA.H | TOL_ASYM.H | 0.600 | 0.000 | * |
| LDI.H | TOL_ASYM.H | 0.510 | 0.000 | * |
| LDMC.H | TOL_ASYM.H | 0.340 | 0.004 | * |
| MGR.H | TOL_ASYM.H | -0.210 | 0.087 | |
| RESpT2.H | TOL_ASYM.H | -0.150 | 0.208 | |
| SLA.H | TOL_ASYM.H | -0.260 | 0.030 | * |
| TOL_ASYM.H | TOL_MGR.H | -0.020 | 0.898 | |
| TOL_ASYM.H | TOL_negXMID.H | -0.440 | 0.000 | * |
| negXMID.H | TOL_ASYM.H | -0.470 | 0.000 | * |
| IGR.C | TOL_ASYM.H | 0.320 | 0.007 | * |
| LTH.C | TOL_ASYM.H | -0.080 | 0.506 | |
| NLEA.C | TOL_ASYM.H | -0.140 | 0.255 | |
| RESmT1.C | TOL_ASYM.H | -0.030 | 0.838 | |
| RESmT2.C | TOL_ASYM.H | 0.280 | 0.022 | * |
| RESpT1.C | TOL_ASYM.H | 0.080 | 0.515 | |
| RESpT2.C | TOL_ASYM.H | 0.210 | 0.090 | |
| IGR.F | TOL_ASYM.H | 0.190 | 0.118 | |
| LTH.F | TOL_ASYM.H | -0.150 | 0.217 | |
| NLEA.F | TOL_ASYM.H | -0.160 | 0.200 | |
| TOL_ASYM.H | TOL_IGR.F | -0.240 | 0.044 | * |
| IGR.H | TOL_ASYM.H | -0.020 | 0.898 | |
| LTH.H | TOL_ASYM.H | 0.090 | 0.483 | |
| NLEA.H | TOL_ASYM.H | -0.070 | 0.557 | |
| TOL_ASYM.H | TOL_IGR.H | 0.060 | 0.603 | |
| ASYM.C | TOL_MGR.H | 0.120 | 0.318 | |
| LA.C | TOL_MGR.H | 0.070 | 0.546 | |

| Trait_1 | Trait_2 | corr.coef | p-value | |
|---|---|---|---|---|
| LDI.C | TOL_MGR.H | 0.170 | 0.155 | |
| LDMC.C | TOL_MGR.H | -0.130 | 0.304 | |
| MGR.C | TOL_MGR.H | -0.190 | 0.128 | |
| SLA.C | TOL_MGR.H | 0.020 | 0.886 | |
| SSIZ.C | TOL_MGR.H | 0.010 | 0.954 | |
| TGER.C | TOL_MGR.H | -0.010 | 0.959 | |
| negXMID.C | TOL_MGR.H | -0.180 | 0.128 | |
| ASYM.F | TOL_MGR.H | 0.100 | 0.402 | |
| LA.F | TOL_MGR.H | 0.050 | 0.663 | |
| LDI.F | TOL_MGR.H | 0.180 | 0.150 | |
| LDMC.F | TOL_MGR.H | -0.060 | 0.600 | |
| MGR.F | TOL_MGR.H | 0.100 | 0.434 | |
| RESmT2.F | TOL_MGR.H | 0.050 | 0.664 | |
| SLA.F | TOL_MGR.H | 0.070 | 0.563 | |
| TOL_ASYM.F | TOL_MGR.H | -0.130 | 0.292 | |
| TOL_MGR.F | TOL_MGR.H | 0.100 | 0.432 | |
| TOL_MGR.H | TOL_negXMID.F | 0.130 | 0.296 | |
| negXMID.F | TOL_MGR.H | -0.100 | 0.433 | |
| ASYM.H | TOL_MGR.H | 0.030 | 0.794 | |
| LA.H | TOL_MGR.H | 0.030 | 0.783 | |
| LDI.H | TOL_MGR.H | 0.130 | 0.305 | |
| LDMC.H | TOL_MGR.H | 0.080 | 0.498 | |
| MGR.H | TOL_MGR.H | 0.710 | 0.000 | * |
| RESpT2.H | TOL_MGR.H | 0.040 | 0.717 | |
| SLA.H | TOL_MGR.H | -0.060 | 0.629 | |
| TOL_ASYM.H | TOL_MGR.H | -0.020 | 0.898 | |
| TOL_MGR.H | TOL_negXMID.H | -0.180 | 0.128 | |
| negXMID.H | TOL_MGR.H | -0.080 | 0.518 | |
| IGR.C | TOL_MGR.H | 0.120 | 0.333 | |
| LTH.C | TOL_MGR.H | 0.180 | 0.147 | |
| NLEA.C | TOL_MGR.H | -0.110 | 0.359 | |
| RESmT1.C | TOL_MGR.H | 0.080 | 0.506 | |
| RESmT2.C | TOL_MGR.H | -0.130 | 0.269 | |
| RESpT1.C | TOL_MGR.H | -0.110 | 0.356 | |
| RESpT2.C | TOL_MGR.H | 0.010 | 0.925 | |
| IGR.F | TOL_MGR.H | -0.100 | 0.408 | |
| LTH.F | TOL_MGR.H | 0.070 | 0.550 | |
| NLEA.F | TOL_MGR.H | -0.080 | 0.515 | |
| TOL_IGR.F | TOL_MGR.H | -0.190 | 0.111 | |
| IGR.H | TOL_MGR.H | -0.040 | 0.752 | |
| LTH.H | TOL_MGR.H | 0.210 | 0.087 | |
| NLEA.H | TOL_MGR.H | -0.220 | 0.075 | |
| TOL_IGR.H | TOL_MGR.H | 0.070 | 0.593 | |
| ASYM.C | TOL_negXMID.H | -0.280 | 0.021 | * |
| LA.C | TOL_negXMID.H | -0.360 | 0.003 | * |
| LDI.C | TOL_negXMID.H | -0.330 | 0.006 | * |
| LDMC.C | TOL_negXMID.H | -0.140 | 0.258 | |
| MGR.C | TOL_negXMID.H | 0.030 | 0.836 | |
| SLA.C | TOL_negXMID.H | -0.100 | 0.404 | |

| Trait_1 | Trait_2 | corr.coef | p-value | |
|---|---|---|---|---|
| SSIZ.C | TOL_negXMID.H | 0.080 | 0.506 | |
| TGER.C | TOL_negXMID.H | 0.050 | 0.710 | |
| negXMID.C | TOL_negXMID.H | 0.170 | 0.162 | |
| ASYM.F | TOL_negXMID.H | -0.320 | 0.007 | * |
| LA.F | TOL_negXMID.H | -0.310 | 0.009 | * |
| LDI.F | TOL_negXMID.H | -0.260 | 0.032 | * |
| LDMC.F | TOL_negXMID.H | -0.160 | 0.181 | |
| MGR.F | TOL_negXMID.H | 0.030 | 0.789 | |
| RESmT2.F | TOL_negXMID.H | -0.010 | 0.955 | |
| SLA.F | TOL_negXMID.H | 0.030 | 0.822 | |
| TOL_ASYM.F | TOL_negXMID.H | -0.040 | 0.737 | |
| TOL_MGR.F | TOL_negXMID.H | 0.030 | 0.827 | |
| TOL_negXMID.F | TOL_negXMID.H | -0.160 | 0.182 | |
| negXMID.F | TOL_negXMID.H | 0.040 | 0.723 | |
| ASYM.H | TOL_negXMID.H | -0.380 | 0.001 | * |
| LA.H | TOL_negXMID.H | -0.330 | 0.005 | * |
| LDI.H | TOL_negXMID.H | -0.430 | 0.000 | * |
| LDMC.H | TOL_negXMID.H | -0.240 | 0.046 | * |
| MGR.H | TOL_negXMID.H | -0.130 | 0.288 | |
| RESpT2.H | TOL_negXMID.H | 0.010 | 0.961 | |
| SLA.H | TOL_negXMID.H | 0.130 | 0.275 | |
| TOL_ASYM.H | TOL_negXMID.H | -0.440 | 0.000 | * |
| TOL_MGR.H | TOL_negXMID.H | -0.180 | 0.128 | |
| negXMID.H | TOL_negXMID.H | 0.530 | 0.000 | * |
| IGR.C | TOL_negXMID.H | -0.080 | 0.504 | |
| LTH.C | TOL_negXMID.H | 0.100 | 0.409 | |
| NLEA.C | TOL_negXMID.H | 0.210 | 0.083 | |
| RESmT1.C | TOL_negXMID.H | -0.170 | 0.161 | |
| RESmT2.C | TOL_negXMID.H | 0.190 | 0.109 | |
| RESpT1.C | TOL_negXMID.H | -0.080 | 0.510 | |
| RESpT2.C | TOL_negXMID.H | -0.240 | 0.051 | |
| IGR.F | TOL_negXMID.H | -0.090 | 0.485 | |
| LTH.F | TOL_negXMID.H | 0.210 | 0.079 | |
| NLEA.F | TOL_negXMID.H | 0.100 | 0.404 | |
| TOL_IGR.F | TOL_negXMID.H | 0.060 | 0.596 | |
| IGR.H | TOL_negXMID.H | 0.080 | 0.523 | |
| LTH.H | TOL_negXMID.H | 0.010 | 0.929 | |
| NLEA.H | TOL_negXMID.H | 0.240 | 0.048 | * |
| TOL_IGR.H | TOL_negXMID.H | 0.040 | 0.724 | |
| ASYM.C | negXMID.H | -0.320 | 0.007 | * |
| LA.C | negXMID.H | -0.360 | 0.003 | * |
| LDI.C | negXMID.H | -0.230 | 0.058 | |
| LDMC.C | negXMID.H | -0.220 | 0.064 | |
| MGR.C | negXMID.H | -0.150 | 0.226 | |
| negXMID.H | SLA.C | 0.220 | 0.064 | |
| negXMID.H | SSIZ.C | 0.190 | 0.109 | |
| negXMID.H | TGER.C | 0.430 | 0.000 | * |
| negXMID.C | negXMID.H | 0.430 | 0.000 | * |
| ASYM.F | negXMID.H | -0.350 | 0.003 | * |

| Trait_1 | Trait_2 | corr.coef | p-value | |
|---|---|---|---|---|
| LA.F | negXMID.H | -0.290 | 0.017 | * |
| LDI.F | negXMID.H | -0.210 | 0.080 | |
| LDMC.F | negXMID.H | -0.240 | 0.044 | * |
| MGR.F | negXMID.H | 0.160 | 0.199 | |
| negXMID.H | RESmT2.F | 0.340 | 0.004 | * |
| negXMID.H | SLA.F | 0.190 | 0.123 | |
| negXMID.H | TOL_ASYM.F | 0.110 | 0.385 | |
| negXMID.H | TOL_MGR.F | 0.010 | 0.917 | |
| negXMID.H | TOL_negXMID.F | -0.190 | 0.126 | |
| negXMID.F | negXMID.H | 0.470 | 0.000 | * |
| ASYM.H | negXMID.H | -0.380 | 0.001 | * |
| LA.H | negXMID.H | -0.310 | 0.010 | * |
| LDI.H | negXMID.H | -0.340 | 0.004 | * |
| LDMC.H | negXMID.H | -0.200 | 0.104 | |
| MGR.H | negXMID.H | 0.120 | 0.332 | |
| negXMID.H | RESpT2.H | 0.150 | 0.228 | |
| negXMID.H | SLA.H | 0.190 | 0.125 | |
| negXMID.H | TOL_ASYM.H | -0.470 | 0.000 | * |
| negXMID.H | TOL_MGR.H | -0.080 | 0.518 | |
| negXMID.H | TOL_negXMID.H | 0.530 | 0.000 | * |
| IGR.C | negXMID.H | -0.140 | 0.242 | |
| LTH.C | negXMID.H | -0.170 | 0.157 | |
| negXMID.H | NLEA.C | 0.030 | 0.823 | |
| negXMID.H | RESmT1.C | -0.160 | 0.189 | |
| negXMID.H | RESmT2.C | 0.070 | 0.581 | |
| negXMID.H | RESpT1.C | -0.080 | 0.526 | |
| negXMID.H | RESpT2.C | -0.180 | 0.137 | |
| IGR.F | negXMID.H | -0.280 | 0.019 | * |
| LTH.F | negXMID.H | 0.000 | 0.988 | |
| negXMID.H | NLEA.F | 0.000 | 0.990 | |
| negXMID.H | TOL_IGR.F | -0.020 | 0.860 | |
| IGR.H | negXMID.H | 0.010 | 0.942 | |
| LTH.H | negXMID.H | 0.020 | 0.859 | |
| negXMID.H | NLEA.H | 0.100 | 0.416 | |
| negXMID.H | TOL_IGR.H | 0.130 | 0.291 | |
| ASYM.C | IGR.C | 0.260 | 0.033 | * |
| IGR.C | LA.C | 0.250 | 0.041 | * |
| IGR.C | LDI.C | 0.200 | 0.108 | |
| IGR.C | LDMC.C | 0.130 | 0.292 | |
| IGR.C | MGR.C | 0.030 | 0.827 | |
| IGR.C | SLA.C | -0.080 | 0.506 | |
| IGR.C | SSIZ.C | 0.150 | 0.232 | |
| IGR.C | TGER.C | -0.170 | 0.175 | |
| IGR.C | negXMID.C | 0.030 | 0.787 | |
| ASYM.F | IGR.C | 0.300 | 0.012 | * |
| IGR.C | LA.F | 0.230 | 0.060 | |
| IGR.C | LDI.F | 0.240 | 0.046 | * |
| IGR.C | LDMC.F | 0.030 | 0.831 | |
| IGR.C | MGR.F | 0.130 | 0.296 | |

| Trait_1 | Trait_2 | corr.coef | p-value | |
|---|---|---|---|---|
| IGR.C | RESmT2.F | -0.100 | 0.409 | |
| IGR.C | SLA.F | -0.030 | 0.838 | |
| IGR.C | TOL_ASYM.F | 0.170 | 0.161 | |
| IGR.C | TOL_MGR.F | -0.070 | 0.554 | |
| IGR.C | TOL_negXMID.F | -0.030 | 0.835 | |
| IGR.C | negXMID.F | -0.130 | 0.276 | |
| ASYM.H | IGR.C | 0.230 | 0.058 | |
| IGR.C | LA.H | 0.290 | 0.014 | * |
| IGR.C | LDI.H | 0.240 | 0.045 | * |
| IGR.C | LDMC.H | 0.030 | 0.790 | |
| IGR.C | MGR.H | -0.060 | 0.619 | |
| IGR.C | RESpT2.H | 0.070 | 0.585 | |
| IGR.C | SLA.H | 0.010 | 0.960 | |
| IGR.C | TOL_ASYM.H | 0.320 | 0.007 | * |
| IGR.C | TOL_MGR.H | 0.120 | 0.333 | |
| IGR.C | TOL_negXMID.H | -0.080 | 0.504 | |
| IGR.C | negXMID.H | -0.140 | 0.242 | |
| IGR.C | LTH.C | 0.090 | 0.446 | |
| IGR.C | NLEA.C | 0.020 | 0.845 | |
| IGR.C | RESmT1.C | 0.010 | 0.908 | |
| IGR.C | RESmT2.C | 0.230 | 0.056 | |
| IGR.C | RESpT1.C | 0.160 | 0.197 | |
| IGR.C | RESpT2.C | -0.140 | 0.242 | |
| IGR.C | IGR.F | -0.030 | 0.794 | |
| IGR.C | LTH.F | 0.180 | 0.143 | |
| IGR.C | NLEA.F | -0.070 | 0.564 | |
| IGR.C | TOL_IGR.F | -0.730 | 0.000 | * |
| IGR.C | IGR.H | -0.080 | 0.530 | |
| IGR.C | LTH.H | 0.120 | 0.333 | |
| IGR.C | NLEA.H | 0.030 | 0.816 | |
| IGR.C | TOL_IGR.H | -0.130 | 0.284 | |
| ASYM.C | LTH.C | 0.120 | 0.324 | |
| LA.C | LTH.C | 0.180 | 0.141 | |
| LDI.C | LTH.C | -0.260 | 0.034 | * |
| LDMC.C | LTH.C | 0.050 | 0.706 | |
| LTH.C | MGR.C | 0.060 | 0.649 | |
| LTH.C | SLA.C | -0.560 | 0.000 | * |
| LTH.C | SSIZ.C | 0.400 | 0.001 | * |
| LTH.C | TGER.C | -0.150 | 0.222 | |
| LTH.C | negXMID.C | 0.060 | 0.644 | |
| ASYM.F | LTH.C | 0.120 | 0.332 | |
| LA.F | LTH.C | 0.190 | 0.110 | |
| LDI.F | LTH.C | -0.210 | 0.081 | |
| LDMC.F | LTH.C | -0.080 | 0.502 | |
| LTH.C | MGR.F | -0.010 | 0.947 | |
| LTH.C | RESmT2.F | -0.160 | 0.181 | |
| LTH.C | SLA.F | -0.470 | 0.000 | * |
| LTH.C | TOL_ASYM.F | 0.010 | 0.945 | |
| LTH.C | TOL_MGR.F | -0.020 | 0.881 | |

| Trait_1 | Trait_2 | corr.coef | p-value | |
|---|---|---|---|---|
| LTH.C | TOL_negXMID.F | -0.080 | 0.526 | |
| LTH.C | negXMID.F | -0.040 | 0.727 | |
| ASYM.H | LTH.C | 0.050 | 0.687 | |
| LA.H | LTH.C | 0.180 | 0.133 | |
| LDI.H | LTH.C | -0.140 | 0.243 | |
| LDMC.H | LTH.C | -0.120 | 0.344 | |
| LTH.C | MGR.H | 0.100 | 0.412 | |
| LTH.C | RESpT2.H | -0.100 | 0.429 | |
| LTH.C | SLA.H | -0.250 | 0.040 | * |
| LTH.C | TOL_ASYM.H | -0.080 | 0.506 | |
| LTH.C | TOL_MGR.H | 0.180 | 0.147 | |
| LTH.C | TOL_negXMID.H | 0.100 | 0.409 | |
| LTH.C | negXMID.H | -0.170 | 0.157 | |
| IGR.C | LTH.C | 0.090 | 0.446 | |
| LTH.C | NLEA.C | -0.330 | 0.005 | * |
| LTH.C | RESmT1.C | 0.510 | 0.000 | * |
| LTH.C | RESmT2.C | -0.030 | 0.780 | |
| LTH.C | RESpT1.C | 0.400 | 0.001 | * |
| LTH.C | RESpT2.C | -0.160 | 0.183 | |
| IGR.F | LTH.C | 0.030 | 0.794 | |
| LTH.C | LTH.F | 0.560 | 0.000 | * |
| LTH.C | NLEA.F | -0.290 | 0.017 | * |
| LTH.C | TOL_IGR.F | -0.070 | 0.559 | |
| IGR.H | LTH.C | 0.100 | 0.411 | |
| LTH.C | LTH.H | 0.110 | 0.390 | |
| LTH.C | NLEA.H | -0.410 | 0.000 | * |
| LTH.C | TOL_IGR.H | -0.090 | 0.475 | |
| ASYM.C | NLEA.C | -0.320 | 0.007 | * |
| LA.C | NLEA.C | -0.450 | 0.000 | * |
| LDI.C | NLEA.C | -0.020 | 0.845 | |
| LDMC.C | NLEA.C | -0.180 | 0.147 | |
| MGR.C | NLEA.C | 0.040 | 0.739 | |
| NLEA.C | SLA.C | 0.410 | 0.000 | * |
| NLEA.C | SSIZ.C | -0.430 | 0.000 | * |
| NLEA.C | TGER.C | -0.310 | 0.009 | * |
| negXMID.C | NLEA.C | -0.150 | 0.204 | |
| ASYM.F | NLEA.C | -0.320 | 0.008 | * |
| LA.F | NLEA.C | -0.490 | 0.000 | * |
| LDI.F | NLEA.C | -0.030 | 0.835 | |
| LDMC.F | NLEA.C | -0.030 | 0.791 | |
| MGR.F | NLEA.C | 0.110 | 0.382 | |
| NLEA.C | RESmT2.F | 0.000 | 0.983 | |
| NLEA.C | SLA.F | 0.430 | 0.000 | * |
| NLEA.C | TOL_ASYM.F | 0.090 | 0.447 | |
| NLEA.C | TOL_MGR.F | 0.000 | 0.982 | |
| NLEA.C | TOL_negXMID.F | 0.040 | 0.731 | |
| negXMID.F | NLEA.C | -0.020 | 0.865 | |
| ASYM.H | NLEA.C | -0.360 | 0.003 | * |
| LA.H | NLEA.C | -0.470 | 0.000 | * |

| Trait_1 | Trait_2 | corr.coef | p-value | |
|---|---|---|---|---|
| LDI.H | NLEA.C | -0.060 | 0.620 | |
| LDMC.H | NLEA.C | -0.220 | 0.067 | |
| MGR.H | NLEA.C | -0.160 | 0.195 | |
| NLEA.C | RESpT2.H | 0.040 | 0.754 | |
| NLEA.C | SLA.H | 0.530 | 0.000 | * |
| NLEA.C | TOL_ASYM.H | -0.140 | 0.255 | |
| NLEA.C | TOL_MGR.H | -0.110 | 0.359 | |
| NLEA.C | TOL_negXMID.H | 0.210 | 0.083 | |
| negXMID.H | NLEA.C | 0.030 | 0.823 | |
| IGR.C | NLEA.C | 0.020 | 0.845 | |
| LTH.C | NLEA.C | -0.330 | 0.005 | * |
| NLEA.C | RESmT1.C | -0.280 | 0.019 | * |
| NLEA.C | RESmT2.C | 0.130 | 0.279 | |
| NLEA.C | RESpT1.C | -0.220 | 0.070 | |
| NLEA.C | RESpT2.C | 0.010 | 0.938 | |
| IGR.F | NLEA.C | -0.150 | 0.207 | |
| LTH.F | NLEA.C | -0.160 | 0.186 | |
| NLEA.C | NLEA.F | 0.870 | 0.000 | * |
| NLEA.C | TOL_IGR.F | -0.020 | 0.886 | |
| IGR.H | NLEA.C | -0.190 | 0.127 | |
| LTH.H | NLEA.C | -0.240 | 0.044 | * |
| NLEA.C | NLEA.H | 0.920 | 0.000 | * |
| NLEA.C | TOL_IGR.H | -0.130 | 0.276 | |
| ASYM.C | RESmT1.C | 0.050 | 0.697 | |
| LA.C | RESmT1.C | 0.180 | 0.146 | |
| LDI.C | RESmT1.C | -0.190 | 0.109 | |
| LDMC.C | RESmT1.C | -0.090 | 0.454 | |
| MGR.C | RESmT1.C | 0.080 | 0.498 | |
| RESmT1.C | SLA.C | -0.370 | 0.002 | * |
| RESmT1.C | SSIZ.C | 0.270 | 0.026 | * |
| RESmT1.C | TGER.C | 0.000 | 0.990 | |
| negXMID.C | RESmT1.C | 0.080 | 0.504 | |
| ASYM.F | RESmT1.C | 0.050 | 0.669 | |
| LA.F | RESmT1.C | 0.170 | 0.153 | |
| LDI.F | RESmT1.C | -0.210 | 0.084 | |
| LDMC.F | RESmT1.C | -0.210 | 0.089 | |
| MGR.F | RESmT1.C | -0.120 | 0.327 | |
| RESmT1.C | RESmT2.F | 0.220 | 0.066 | |
| RESmT1.C | SLA.F | -0.410 | 0.001 | * |
| RESmT1.C | TOL_ASYM.F | -0.100 | 0.424 | |
| RESmT1.C | TOL_MGR.F | -0.190 | 0.113 | |
| RESmT1.C | TOL_negXMID.F | -0.120 | 0.317 | |
| negXMID.F | RESmT1.C | -0.010 | 0.910 | |
| ASYM.H | RESmT1.C | 0.080 | 0.494 | |
| LA.H | RESmT1.C | 0.140 | 0.251 | |
| LDI.H | RESmT1.C | -0.130 | 0.305 | |
| LDMC.H | RESmT1.C | -0.200 | 0.104 | |
| MGR.H | RESmT1.C | 0.060 | 0.652 | |
| RESmT1.C | RESpT2.H | -0.180 | 0.136 | |

| Trait_1 | Trait_2 | corr.coef | p-value | |
|---|---|---|---|---|
| RESmT1.C | SLA.H | -0.230 | 0.059 | |
| RESmT1.C | TOL_ASYM.H | -0.030 | 0.838 | |
| RESmT1.C | TOL_MGR.H | 0.080 | 0.506 | |
| RESmT1.C | TOL_negXMID.H | -0.170 | 0.161 | |
| negXMID.H | RESmT1.C | -0.160 | 0.189 | |
| IGR.C | RESmT1.C | 0.010 | 0.908 | |
| LTH.C | RESmT1.C | 0.510 | 0.000 | * |
| NLEA.C | RESmT1.C | -0.280 | 0.019 | * |
| RESmT1.C | RESmT2.C | 0.030 | 0.799 | |
| RESmT1.C | RESpT1.C | 0.570 | 0.000 | * |
| RESmT1.C | RESpT2.C | -0.040 | 0.738 | |
| IGR.F | RESmT1.C | 0.260 | 0.031 | * |
| LTH.F | RESmT1.C | 0.490 | 0.000 | * |
| NLEA.F | RESmT1.C | -0.230 | 0.058 | |
| RESmT1.C | TOL_IGR.F | 0.090 | 0.454 | |
| IGR.H | RESmT1.C | 0.160 | 0.182 | |
| LTH.H | RESmT1.C | 0.250 | 0.038 | * |
| NLEA.H | RESmT1.C | -0.330 | 0.006 | * |
| RESmT1.C | TOL_IGR.H | -0.010 | 0.918 | |
| ASYM.C | RESmT2.C | 0.270 | 0.022 | * |
| LA.C | RESmT2.C | 0.160 | 0.195 | |
| LDI.C | RESmT2.C | 0.140 | 0.266 | |
| LDMC.C | RESmT2.C | 0.200 | 0.095 | |
| MGR.C | RESmT2.C | 0.020 | 0.890 | |
| RESmT2.C | SLA.C | -0.100 | 0.410 | |
| RESmT2.C | SSIZ.C | -0.030 | 0.806 | |
| RESmT2.C | TGER.C | -0.320 | 0.007 | * |
| negXMID.C | RESmT2.C | 0.040 | 0.735 | |
| ASYM.F | RESmT2.C | 0.260 | 0.033 | * |
| LA.F | RESmT2.C | 0.160 | 0.190 | |
| LDI.F | RESmT2.C | 0.200 | 0.104 | |
| LDMC.F | RESmT2.C | 0.110 | 0.368 | |
| MGR.F | RESmT2.C | 0.070 | 0.573 | |
| RESmT2.C | RESmT2.F | 0.110 | 0.377 | |
| RESmT2.C | SLA.F | -0.070 | 0.579 | |
| RESmT2.C | TOL_ASYM.F | -0.070 | 0.566 | |
| RESmT2.C | TOL_MGR.F | 0.060 | 0.629 | |
| RESmT2.C | TOL_negXMID.F | -0.150 | 0.211 | |
| negXMID.F | RESmT2.C | -0.040 | 0.720 | |
| ASYM.H | RESmT2.C | 0.210 | 0.088 | |
| LA.H | RESmT2.C | 0.160 | 0.198 | |
| LDI.H | RESmT2.C | 0.130 | 0.303 | |
| LDMC.H | RESmT2.C | 0.120 | 0.321 | |
| MGR.H | RESmT2.C | -0.210 | 0.088 | |
| RESmT2.C | RESpT2.H | -0.070 | 0.564 | |
| RESmT2.C | SLA.H | 0.030 | 0.797 | |
| RESmT2.C | TOL_ASYM.H | 0.280 | 0.022 | * |
| RESmT2.C | TOL_MGR.H | -0.130 | 0.269 | |
| RESmT2.C | TOL_negXMID.H | 0.190 | 0.109 | |

| Trait_1 | Trait_2 | corr.coef | p-value | |
|---|---|---|---|---|
| negXMID.H | RESmT2.C | 0.070 | 0.581 | |
| IGR.C | RESmT2.C | 0.230 | 0.056 | |
| LTH.C | RESmT2.C | -0.030 | 0.780 | |
| NLEA.C | RESmT2.C | 0.130 | 0.279 | |
| RESmT1.C | RESmT2.C | 0.030 | 0.799 | |
| RESmT2.C | RESpT1.C | 0.170 | 0.153 | |
| RESmT2.C | RESpT2.C | -0.110 | 0.369 | |
| IGR.F | RESmT2.C | -0.040 | 0.719 | |
| LTH.F | RESmT2.C | 0.050 | 0.656 | |
| NLEA.F | RESmT2.C | 0.100 | 0.412 | |
| RESmT2.C | TOL_IGR.F | -0.190 | 0.126 | |
| IGR.H | RESmT2.C | 0.290 | 0.015 | * |
| LTH.H | RESmT2.C | -0.100 | 0.434 | |
| NLEA.H | RESmT2.C | 0.140 | 0.238 | |
| RESmT2.C | TOL_IGR.H | 0.180 | 0.143 | |
| ASYM.C | RESpT1.C | 0.200 | 0.104 | |
| LA.C | RESpT1.C | 0.210 | 0.091 | |
| LDI.C | RESpT1.C | -0.110 | 0.353 | |
| LDMC.C | RESpT1.C | 0.190 | 0.128 | |
| MGR.C | RESpT1.C | 0.180 | 0.142 | |
| RESpT1.C | SLA.C | -0.460 | 0.000 | * |
| RESpT1.C | SSIZ.C | 0.370 | 0.002 | * |
| RESpT1.C | TGER.C | -0.110 | 0.362 | |
| negXMID.C | RESpT1.C | 0.000 | 0.998 | |
| ASYM.F | RESpT1.C | 0.190 | 0.115 | |
| LA.F | RESpT1.C | 0.220 | 0.066 | |
| LDI.F | RESpT1.C | -0.150 | 0.214 | |
| LDMC.F | RESpT1.C | 0.100 | 0.415 | |
| MGR.F | RESpT1.C | -0.070 | 0.587 | |
| RESmT2.F | RESpT1.C | -0.010 | 0.925 | |
| RESpT1.C | SLA.F | -0.490 | 0.000 | * |
| RESpT1.C | TOL_ASYM.F | -0.110 | 0.364 | |
| RESpT1.C | TOL_MGR.F | -0.070 | 0.553 | |
| RESpT1.C | TOL_negXMID.F | -0.160 | 0.180 | |
| negXMID.F | RESpT1.C | -0.030 | 0.805 | |
| ASYM.H | RESpT1.C | 0.210 | 0.079 | |
| LA.H | RESpT1.C | 0.290 | 0.016 | * |
| LDI.H | RESpT1.C | -0.030 | 0.777 | |
| LDMC.H | RESpT1.C | 0.030 | 0.813 | |
| MGR.H | RESpT1.C | -0.020 | 0.851 | |
| RESpT1.C | RESpT2.H | -0.050 | 0.708 | |
| RESpT1.C | SLA.H | -0.280 | 0.018 | * |
| RESpT1.C | TOL_ASYM.H | 0.080 | 0.515 | |
| RESpT1.C | TOL_MGR.H | -0.110 | 0.356 | |
| RESpT1.C | TOL_negXMID.H | -0.080 | 0.510 | |
| negXMID.H | RESpT1.C | -0.080 | 0.526 | |
| IGR.C | RESpT1.C | 0.160 | 0.197 | |
| LTH.C | RESpT1.C | 0.400 | 0.001 | * |
| NLEA.C | RESpT1.C | -0.220 | 0.070 | |

| Trait_1 | Trait_2 | corr.coef | p-value | |
|---|---|---|---|---|
| RESmT1.C | RESpT1.C | 0.570 | 0.000 | * |
| RESmT2.C | RESpT1.C | 0.170 | 0.153 | |
| RESpT1.C | RESpT2.C | -0.120 | 0.322 | |
| IGR.F | RESpT1.C | 0.030 | 0.821 | |
| LTH.F | RESpT1.C | 0.310 | 0.009 | * |
| NLEA.F | RESpT1.C | -0.240 | 0.047 | * |
| RESpT1.C | TOL_IGR.F | -0.190 | 0.127 | |
| IGR.H | RESpT1.C | 0.120 | 0.317 | |
| LTH.H | RESpT1.C | 0.020 | 0.867 | |
| NLEA.H | RESpT1.C | -0.250 | 0.039 | * |
| RESpT1.C | TOL_IGR.H | -0.050 | 0.700 | |
| ASYM.C | RESpT2.C | 0.020 | 0.887 | |
| LA.C | RESpT2.C | 0.070 | 0.593 | |
| LDI.C | RESpT2.C | 0.030 | 0.780 | |
| LDMC.C | RESpT2.C | 0.230 | 0.059 | |
| MGR.C | RESpT2.C | 0.120 | 0.320 | |
| RESpT2.C | SLA.C | 0.200 | 0.107 | |
| RESpT2.C | SSIZ.C | -0.140 | 0.237 | |
| RESpT2.C | TGER.C | -0.100 | 0.392 | |
| negXMID.C | RESpT2.C | -0.030 | 0.806 | |
| ASYM.F | RESpT2.C | 0.120 | 0.338 | |
| LA.F | RESpT2.C | 0.120 | 0.319 | |
| LDI.F | RESpT2.C | 0.030 | 0.831 | |
| LDMC.F | RESpT2.C | 0.300 | 0.012 | * |
| MGR.F | RESpT2.C | -0.130 | 0.304 | |
| RESmT2.F | RESpT2.C | 0.080 | 0.502 | |
| RESpT2.C | SLA.F | 0.070 | 0.559 | |
| RESpT2.C | TOL_ASYM.F | 0.250 | 0.041 | * |
| RESpT2.C | TOL_MGR.F | -0.010 | 0.950 | |
| RESpT2.C | TOL_negXMID.F | 0.000 | 0.986 | |
| negXMID.F | RESpT2.C | -0.080 | 0.521 | |
| ASYM.H | RESpT2.C | 0.100 | 0.393 | |
| LA.H | RESpT2.C | 0.060 | 0.650 | |
| LDI.H | RESpT2.C | 0.020 | 0.867 | |
| LDMC.H | RESpT2.C | 0.350 | 0.003 | * |
| MGR.H | RESpT2.C | 0.070 | 0.590 | |
| RESpT2.C | RESpT2.H | 0.180 | 0.129 | |
| RESpT2.C | SLA.H | -0.080 | 0.522 | |
| RESpT2.C | TOL_ASYM.H | 0.210 | 0.090 | |
| RESpT2.C | TOL_MGR.H | 0.010 | 0.925 | |
| RESpT2.C | TOL_negXMID.H | -0.240 | 0.051 | |
| negXMID.H | RESpT2.C | -0.180 | 0.137 | |
| IGR.C | RESpT2.C | -0.140 | 0.242 | |
| LTH.C | RESpT2.C | -0.160 | 0.183 | |
| NLEA.C | RESpT2.C | 0.010 | 0.938 | |
| RESmT1.C | RESpT2.C | -0.040 | 0.738 | |
| RESmT2.C | RESpT2.C | -0.110 | 0.369 | |
| RESpT1.C | RESpT2.C | -0.120 | 0.322 | |
| IGR.F | RESpT2.C | -0.070 | 0.545 | |

| Trait_1 | Trait_2 | corr.coef | p-value | |
|---|---|---|---|---|
| LTH.F | RESpT2.C | -0.290 | 0.016 | * |
| NLEA.F | RESpT2.C | 0.060 | 0.614 | |
| RESpT2.C | TOL_IGR.F | 0.120 | 0.320 | |
| IGR.H | RESpT2.C | 0.020 | 0.879 | |
| LTH.H | RESpT2.C | 0.030 | 0.804 | |
| NLEA.H | RESpT2.C | 0.080 | 0.489 | |
| RESpT2.C | TOL_IGR.H | 0.060 | 0.634 | |
| ASYM.C | IGR.F | 0.030 | 0.801 | |
| IGR.F | LA.C | 0.060 | 0.604 | |
| IGR.F | LDI.C | 0.040 | 0.723 | |
| IGR.F | LDMC.C | -0.150 | 0.223 | |
| IGR.F | MGR.C | 0.130 | 0.298 | |
| IGR.F | SLA.C | -0.010 | 0.924 | |
| IGR.F | SSIZ.C | -0.170 | 0.159 | |
| IGR.F | TGER.C | 0.060 | 0.650 | |
| IGR.F | negXMID.C | -0.140 | 0.256 | |
| ASYM.F | IGR.F | 0.020 | 0.886 | |
| IGR.F | LA.F | 0.060 | 0.649 | |
| IGR.F | LDI.F | 0.040 | 0.741 | |
| IGR.F | LDMC.F | -0.110 | 0.348 | |
| IGR.F | MGR.F | 0.120 | 0.324 | |
| IGR.F | RESmT2.F | 0.050 | 0.662 | |
| IGR.F | SLA.F | -0.020 | 0.858 | |
| IGR.F | TOL_ASYM.F | -0.110 | 0.378 | |
| IGR.F | TOL_MGR.F | 0.060 | 0.643 | |
| IGR.F | TOL_negXMID.F | 0.050 | 0.704 | |
| IGR.F | negXMID.F | -0.180 | 0.137 | |
| ASYM.H | IGR.F | 0.100 | 0.426 | |
| IGR.F | LA.H | 0.050 | 0.680 | |
| IGR.F | LDI.H | 0.030 | 0.827 | |
| IGR.F | LDMC.H | -0.220 | 0.064 | |
| IGR.F | MGR.H | -0.130 | 0.280 | |
| IGR.F | RESpT2.H | -0.260 | 0.029 | * |
| IGR.F | SLA.H | -0.010 | 0.921 | |
| IGR.F | TOL_ASYM.H | 0.190 | 0.118 | |
| IGR.F | TOL_MGR.H | -0.100 | 0.408 | |
| IGR.F | TOL_negXMID.H | -0.090 | 0.485 | |
| IGR.F | negXMID.H | -0.280 | 0.019 | * |
| IGR.C | IGR.F | -0.030 | 0.794 | |
| IGR.F | LTH.C | 0.030 | 0.794 | |
| IGR.F | NLEA.C | -0.150 | 0.207 | |
| IGR.F | RESmT1.C | 0.260 | 0.031 | * |
| IGR.F | RESmT2.C | -0.040 | 0.719 | |
| IGR.F | RESpT1.C | 0.030 | 0.821 | |
| IGR.F | RESpT2.C | -0.070 | 0.545 | |
| IGR.F | LTH.F | 0.070 | 0.579 | |
| IGR.F | NLEA.F | -0.180 | 0.135 | |
| IGR.F | TOL_IGR.F | 0.560 | 0.000 | * |
| IGR.F | IGR.H | 0.190 | 0.126 | |

| Trait_1 | Trait_2 | corr.coef | p-value | |
|---|---|---|---|---|
| IGR.F | LTH.H | 0.250 | 0.041 | * |
| IGR.F | NLEA.H | -0.150 | 0.207 | |
| IGR.F | TOL_IGR.H | 0.090 | 0.475 | |
| ASYM.C | LTH.F | -0.110 | 0.371 | |
| LA.C | LTH.F | -0.030 | 0.782 | |
| LDI.C | LTH.F | -0.390 | 0.001 | * |
| LDMC.C | LTH.F | -0.250 | 0.035 | * |
| LTH.F | MGR.C | 0.020 | 0.896 | |
| LTH.F | SLA.C | -0.330 | 0.005 | * |
| LTH.F | SSIZ.C | 0.290 | 0.015 | * |
| LTH.F | TGER.C | 0.000 | 0.969 | |
| LTH.F | negXMID.C | 0.010 | 0.926 | |
| ASYM.F | LTH.F | -0.180 | 0.129 | |
| LA.F | LTH.F | -0.060 | 0.626 | |
| LDI.F | LTH.F | -0.320 | 0.008 | * |
| LDMC.F | LTH.F | -0.340 | 0.005 | * |
| LTH.F | MGR.F | 0.000 | 0.989 | |
| LTH.F | RESmT2.F | 0.050 | 0.654 | |
| LTH.F | SLA.F | -0.310 | 0.009 | * |
| LTH.F | TOL_ASYM.F | -0.140 | 0.243 | |
| LTH.F | TOL_MGR.F | -0.090 | 0.471 | |
| LTH.F | TOL_negXMID.F | -0.150 | 0.222 | |
| LTH.F | negXMID.F | -0.100 | 0.405 | |
| ASYM.H | LTH.F | -0.150 | 0.206 | |
| LA.H | LTH.F | -0.060 | 0.641 | |
| LDI.H | LTH.F | -0.330 | 0.006 | * |
| LDMC.H | LTH.F | -0.310 | 0.010 | * |
| LTH.F | MGR.H | 0.010 | 0.946 | |
| LTH.F | RESpT2.H | -0.040 | 0.736 | |
| LTH.F | SLA.H | -0.150 | 0.226 | |
| LTH.F | TOL_ASYM.H | -0.150 | 0.217 | |
| LTH.F | TOL_MGR.H | 0.070 | 0.550 | |
| LTH.F | TOL_negXMID.H | 0.210 | 0.079 | |
| LTH.F | negXMID.H | 0.000 | 0.988 | |
| IGR.C | LTH.F | 0.180 | 0.143 | |
| LTH.C | LTH.F | 0.560 | 0.000 | * |
| LTH.F | NLEA.C | -0.160 | 0.186 | |
| LTH.F | RESmT1.C | 0.490 | 0.000 | * |
| LTH.F | RESmT2.C | 0.050 | 0.656 | |
| LTH.F | RESpT1.C | 0.310 | 0.009 | * |
| LTH.F | RESpT2.C | -0.290 | 0.016 | * |
| IGR.F | LTH.F | 0.070 | 0.579 | |
| LTH.F | NLEA.F | -0.230 | 0.063 | |
| LTH.F | TOL_IGR.F | -0.080 | 0.490 | |
| IGR.H | LTH.F | 0.110 | 0.390 | |
| LTH.F | LTH.H | 0.240 | 0.043 | * |
| LTH.F | NLEA.H | -0.270 | 0.022 | * |
| LTH.F | TOL_IGR.H | -0.010 | 0.912 | |
| ASYM.C | NLEA.F | -0.360 | 0.003 | * |

| Trait_1 | Trait_2 | corr.coef | p-value | |
|---|---|---|---|---|
| LA.C | NLEA.F | -0.430 | 0.000 | * |
| LDI.C | NLEA.F | 0.000 | 0.975 | |
| LDMC.C | NLEA.F | -0.110 | 0.375 | |
| MGR.C | NLEA.F | 0.050 | 0.656 | |
| NLEA.F | SLA.C | 0.340 | 0.004 | * |
| NLEA.F | SSIZ.C | -0.430 | 0.000 | * |
| NLEA.F | TGER.C | -0.290 | 0.017 | * |
| negXMID.C | NLEA.F | -0.090 | 0.487 | |
| ASYM.F | NLEA.F | -0.300 | 0.013 | * |
| LA.F | NLEA.F | -0.470 | 0.000 | * |
| LDI.F | NLEA.F | 0.000 | 0.998 | |
| LDMC.F | NLEA.F | 0.000 | 0.978 | |
| MGR.F | NLEA.F | 0.020 | 0.851 | |
| NLEA.F | RESmT2.F | 0.040 | 0.724 | |
| NLEA.F | SLA.F | 0.380 | 0.001 | * |
| NLEA.F | TOL_ASYM.F | 0.150 | 0.228 | |
| NLEA.F | TOL_MGR.F | -0.010 | 0.913 | |
| NLEA.F | TOL_negXMID.F | 0.050 | 0.683 | |
| negXMID.F | NLEA.F | 0.020 | 0.891 | |
| ASYM.H | NLEA.F | -0.330 | 0.005 | * |
| LA.H | NLEA.F | -0.460 | 0.000 | * |
| LDI.H | NLEA.F | -0.040 | 0.746 | |
| LDMC.H | NLEA.F | -0.160 | 0.190 | |
| MGR.H | NLEA.F | -0.110 | 0.356 | |
| NLEA.F | RESpT2.H | -0.040 | 0.748 | |
| NLEA.F | SLA.H | 0.470 | 0.000 | * |
| NLEA.F | TOL_ASYM.H | -0.160 | 0.200 | |
| NLEA.F | TOL_MGR.H | -0.080 | 0.515 | |
| NLEA.F | TOL_negXMID.H | 0.100 | 0.404 | |
| negXMID.H | NLEA.F | 0.000 | 0.990 | |
| IGR.C | NLEA.F | -0.070 | 0.564 | |
| LTH.C | NLEA.F | -0.290 | 0.017 | * |
| NLEA.C | NLEA.F | 0.870 | 0.000 | * |
| NLEA.F | RESmT1.C | -0.230 | 0.058 | |
| NLEA.F | RESmT2.C | 0.100 | 0.412 | |
| NLEA.F | RESpT1.C | -0.240 | 0.047 | * |
| NLEA.F | RESpT2.C | 0.060 | 0.614 | |
| IGR.F | NLEA.F | -0.180 | 0.135 | |
| LTH.F | NLEA.F | -0.230 | 0.063 | |
| NLEA.F | TOL_IGR.F | 0.010 | 0.954 | |
| IGR.H | NLEA.F | -0.140 | 0.239 | |
| LTH.H | NLEA.F | -0.180 | 0.144 | |
| NLEA.F | NLEA.H | 0.800 | 0.000 | * |
| NLEA.F | TOL_IGR.H | 0.000 | 0.984 | |
| ASYM.C | TOL_IGR.F | -0.200 | 0.100 | |
| LA.C | TOL_IGR.F | -0.200 | 0.099 | |
| LDI.C | TOL_IGR.F | -0.210 | 0.086 | |
| LDMC.C | TOL_IGR.F | -0.160 | 0.186 | |
| MGR.C | TOL_IGR.F | -0.040 | 0.726 | |

| Trait_1 | Trait_2 | corr.coef | p-value | |
|---|---|---|---|---|
| SLA.C | TOL_IGR.F | 0.110 | 0.385 | |
| SSIZ.C | TOL_IGR.F | -0.260 | 0.031 | * |
| TGER.C | TOL_IGR.F | 0.170 | 0.168 | |
| negXMID.C | TOL_IGR.F | -0.120 | 0.337 | |
| ASYM.F | TOL_IGR.F | -0.280 | 0.020 | * |
| LA.F | TOL_IGR.F | -0.190 | 0.109 | |
| LDI.F | TOL_IGR.F | -0.250 | 0.035 | * |
| LDMC.F | TOL_IGR.F | -0.090 | 0.484 | |
| MGR.F | TOL_IGR.F | 0.000 | 0.999 | |
| RESmT2.F | TOL_IGR.F | 0.130 | 0.288 | |
| SLA.F | TOL_IGR.F | 0.060 | 0.647 | |
| TOL_ASYM.F | TOL_IGR.F | -0.200 | 0.097 | |
| TOL_IGR.F | TOL_MGR.F | 0.060 | 0.619 | |
| TOL_IGR.F | TOL_negXMID.F | 0.110 | 0.381 | |
| negXMID.F | TOL_IGR.F | 0.090 | 0.463 | |
| ASYM.H | TOL_IGR.F | -0.210 | 0.079 | |
| LA.H | TOL_IGR.F | -0.260 | 0.033 | * |
| LDI.H | TOL_IGR.F | -0.220 | 0.065 | |
| LDMC.H | TOL_IGR.F | -0.150 | 0.210 | |
| MGR.H | TOL_IGR.F | -0.020 | 0.901 | |
| RESpT2.H | TOL_IGR.F | -0.070 | 0.573 | |
| SLA.H | TOL_IGR.F | 0.060 | 0.646 | |
| TOL_ASYM.H | TOL_IGR.F | -0.240 | 0.044 | * |
| TOL_IGR.F | TOL_MGR.H | -0.190 | 0.111 | |
| TOL_IGR.F | TOL_negXMID.H | 0.060 | 0.596 | |
| negXMID.H | TOL_IGR.F | -0.020 | 0.860 | |
| IGR.C | TOL_IGR.F | -0.730 | 0.000 | * |
| LTH.C | TOL_IGR.F | -0.070 | 0.559 | |
| NLEA.C | TOL_IGR.F | -0.020 | 0.886 | |
| RESmT1.C | TOL_IGR.F | 0.090 | 0.454 | |
| RESmT2.C | TOL_IGR.F | -0.190 | 0.126 | |
| RESpT1.C | TOL_IGR.F | -0.190 | 0.127 | |
| RESpT2.C | TOL_IGR.F | 0.120 | 0.320 | |
| IGR.F | TOL_IGR.F | 0.560 | 0.000 | * |
| LTH.F | TOL_IGR.F | -0.080 | 0.490 | |
| NLEA.F | TOL_IGR.F | 0.010 | 0.954 | |
| IGR.H | TOL_IGR.F | 0.150 | 0.212 | |
| LTH.H | TOL_IGR.F | 0.020 | 0.852 | |
| NLEA.H | TOL_IGR.F | 0.010 | 0.912 | |
| TOL_IGR.F | TOL_IGR.H | 0.150 | 0.211 | |
| ASYM.C | IGR.H | -0.010 | 0.933 | |
| IGR.H | LA.C | 0.040 | 0.721 | |
| IGR.H | LDI.C | -0.140 | 0.242 | |
| IGR.H | LDMC.C | 0.130 | 0.296 | |
| IGR.H | MGR.C | 0.110 | 0.376 | |
| IGR.H | SLA.C | -0.260 | 0.032 | * |
| IGR.H | SSIZ.C | 0.110 | 0.348 | |
| IGR.H | TGER.C | 0.030 | 0.827 | |
| IGR.H | negXMID.C | -0.010 | 0.910 | |

| Trait_1 | Trait_2 | corr.coef | p-value | |
|---|---|---|---|---|
| ASYM.F | IGR.H | -0.040 | 0.725 | |
| IGR.H | LA.F | 0.040 | 0.754 | |
| IGR.H | LDI.F | -0.120 | 0.319 | |
| IGR.H | LDMC.F | 0.050 | 0.669 | |
| IGR.H | MGR.F | -0.040 | 0.766 | |
| IGR.H | RESmT2.F | 0.130 | 0.286 | |
| IGR.H | SLA.F | -0.240 | 0.045 | * |
| IGR.H | TOL_ASYM.F | -0.210 | 0.086 | |
| IGR.H | TOL_MGR.F | -0.120 | 0.329 | |
| IGR.H | TOL_negXMID.F | -0.060 | 0.618 | |
| IGR.H | negXMID.F | 0.120 | 0.346 | |
| ASYM.H | IGR.H | 0.040 | 0.714 | |
| IGR.H | LA.H | 0.040 | 0.742 | |
| IGR.H | LDI.H | -0.150 | 0.227 | |
| IGR.H | LDMC.H | 0.010 | 0.932 | |
| IGR.H | MGR.H | 0.030 | 0.789 | |
| IGR.H | RESpT2.H | -0.040 | 0.748 | |
| IGR.H | SLA.H | -0.090 | 0.440 | |
| IGR.H | TOL_ASYM.H | -0.020 | 0.898 | |
| IGR.H | TOL_MGR.H | -0.040 | 0.752 | |
| IGR.H | TOL_negXMID.H | 0.080 | 0.523 | |
| IGR.H | negXMID.H | 0.010 | 0.942 | |
| IGR.C | IGR.H | -0.080 | 0.530 | |
| IGR.H | LTH.C | 0.100 | 0.411 | |
| IGR.H | NLEA.C | -0.190 | 0.127 | |
| IGR.H | RESmT1.C | 0.160 | 0.182 | |
| IGR.H | RESmT2.C | 0.290 | 0.015 | * |
| IGR.H | RESpT1.C | 0.120 | 0.317 | |
| IGR.H | RESpT2.C | 0.020 | 0.879 | |
| IGR.F | IGR.H | 0.190 | 0.126 | |
| IGR.H | LTH.F | 0.110 | 0.390 | |
| IGR.H | NLEA.F | -0.140 | 0.239 | |
| IGR.H | TOL_IGR.F | 0.150 | 0.212 | |
| IGR.H | LTH.H | 0.040 | 0.761 | |
| IGR.H | NLEA.H | -0.160 | 0.178 | |
| IGR.H | TOL_IGR.H | 0.600 | 0.000 | * |
| ASYM.C | LTH.H | 0.040 | 0.720 | |
| LA.C | LTH.H | 0.120 | 0.312 | |
| LDI.C | LTH.H | 0.040 | 0.728 | |
| LDMC.C | LTH.H | 0.020 | 0.889 | |
| LTH.H | MGR.C | 0.050 | 0.667 | |
| LTH.H | SLA.C | -0.180 | 0.133 | |
| LTH.H | SSIZ.C | 0.120 | 0.336 | |
| LTH.H | TGER.C | 0.060 | 0.642 | |
| LTH.H | negXMID.C | 0.090 | 0.464 | |
| ASYM.F | LTH.H | 0.070 | 0.591 | |
| LA.F | LTH.H | 0.140 | 0.260 | |
| LDI.F | LTH.H | 0.020 | 0.889 | |
| LDMC.F | LTH.H | -0.090 | 0.457 | |

| Trait_1 | Trait_2 | corr.coef | p-value | |
|---|---|---|---|---|
| LTH.H | MGR.F | -0.050 | 0.665 | |
| LTH.H | RESmT2.F | 0.150 | 0.234 | |
| LTH.H | SLA.F | -0.170 | 0.169 | |
| LTH.H | TOL_ASYM.F | -0.070 | 0.553 | |
| LTH.H | TOL_MGR.F | -0.030 | 0.796 | |
| LTH.H | TOL_negXMID.F | 0.070 | 0.551 | |
| LTH.H | negXMID.F | 0.020 | 0.900 | |
| ASYM.H | LTH.H | 0.110 | 0.371 | |
| LA.H | LTH.H | 0.100 | 0.411 | |
| LDI.H | LTH.H | 0.010 | 0.928 | |
| LDMC.H | LTH.H | 0.100 | 0.418 | |
| LTH.H | MGR.H | 0.110 | 0.385 | |
| LTH.H | RESpT2.H | -0.080 | 0.531 | |
| LTH.H | SLA.H | -0.240 | 0.046 | * |
| LTH.H | TOL_ASYM.H | 0.090 | 0.483 | |
| LTH.H | TOL_MGR.H | 0.210 | 0.087 | |
| LTH.H | TOL_negXMID.H | 0.010 | 0.929 | |
| LTH.H | negXMID.H | 0.020 | 0.859 | |
| IGR.C | LTH.H | 0.120 | 0.333 | |
| LTH.C | LTH.H | 0.110 | 0.390 | |
| LTH.H | NLEA.C | -0.240 | 0.044 | * |
| LTH.H | RESmT1.C | 0.250 | 0.038 | * |
| LTH.H | RESmT2.C | -0.100 | 0.434 | |
| LTH.H | RESpT1.C | 0.020 | 0.867 | |
| LTH.H | RESpT2.C | 0.030 | 0.804 | |
| IGR.F | LTH.H | 0.250 | 0.041 | * |
| LTH.F | LTH.H | 0.240 | 0.043 | * |
| LTH.H | NLEA.F | -0.180 | 0.144 | |
| LTH.H | TOL_IGR.F | 0.020 | 0.852 | |
| IGR.H | LTH.H | 0.040 | 0.761 | |
| LTH.H | NLEA.H | -0.190 | 0.110 | |
| LTH.H | TOL_IGR.H | 0.030 | 0.801 | |
| ASYM.C | NLEA.H | -0.290 | 0.015 | * |
| LA.C | NLEA.H | -0.420 | 0.000 | * |
| LDI.C | NLEA.H | 0.020 | 0.898 | |
| LDMC.C | NLEA.H | -0.060 | 0.602 | |
| MGR.C | NLEA.H | 0.080 | 0.521 | |
| NLEA.H | SLA.C | 0.400 | 0.001 | * |
| NLEA.H | SSIZ.C | -0.430 | 0.000 | * |
| NLEA.H | TGER.C | -0.250 | 0.042 | * |
| negXMID.C | NLEA.H | -0.100 | 0.414 | |
| ASYM.F | NLEA.H | -0.260 | 0.033 | * |
| LA.F | NLEA.H | -0.440 | 0.000 | * |
| LDI.F | NLEA.H | 0.010 | 0.953 | |
| LDMC.F | NLEA.H | 0.070 | 0.595 | |
| MGR.F | NLEA.H | 0.050 | 0.708 | |
| NLEA.H | RESmT2.F | -0.030 | 0.786 | |
| NLEA.H | SLA.F | 0.440 | 0.000 | * |
| NLEA.H | TOL_ASYM.F | 0.130 | 0.288 | |

| Trait_1 | Trait_2 | corr.coef | p-value | |
|---|---|---|---|---|
| NLEA.H | TOL_MGR.F | -0.060 | 0.637 | |
| NLEA.H | TOL_negXMID.F | 0.020 | 0.854 | |
| negXMID.F | NLEA.H | 0.040 | 0.759 | |
| ASYM.H | NLEA.H | -0.300 | 0.013 | * |
| LA.H | NLEA.H | -0.430 | 0.000 | * |
| LDI.H | NLEA.H | -0.040 | 0.717 | |
| LDMC.H | NLEA.H | -0.160 | 0.176 | |
| MGR.H | NLEA.H | -0.200 | 0.102 | |
| NLEA.H | RESpT2.H | 0.110 | 0.360 | |
| NLEA.H | SLA.H | 0.490 | 0.000 | * |
| NLEA.H | TOL_ASYM.H | -0.070 | 0.557 | |
| NLEA.H | TOL_MGR.H | -0.220 | 0.075 | |
| NLEA.H | TOL_negXMID.H | 0.240 | 0.048 | * |
| negXMID.H | NLEA.H | 0.100 | 0.416 | |
| IGR.C | NLEA.H | 0.030 | 0.816 | |
| LTH.C | NLEA.H | -0.410 | 0.000 | * |
| NLEA.C | NLEA.H | 0.920 | 0.000 | * |
| NLEA.H | RESmT1.C | -0.330 | 0.006 | * |
| NLEA.H | RESmT2.C | 0.140 | 0.238 | |
| NLEA.H | RESpT1.C | -0.250 | 0.039 | * |
| NLEA.H | RESpT2.C | 0.080 | 0.489 | |
| IGR.F | NLEA.H | -0.150 | 0.207 | |
| LTH.F | NLEA.H | -0.270 | 0.022 | * |
| NLEA.F | NLEA.H | 0.800 | 0.000 | * |
| NLEA.H | TOL_IGR.F | 0.010 | 0.912 | |
| IGR.H | NLEA.H | -0.160 | 0.178 | |
| LTH.H | NLEA.H | -0.190 | 0.110 | |
| NLEA.H | TOL_IGR.H | -0.060 | 0.623 | |
| ASYM.C | TOL_IGR.H | 0.050 | 0.694 | |
| LA.C | TOL_IGR.H | 0.080 | 0.536 | |
| LDI.C | TOL_IGR.H | 0.030 | 0.803 | |
| LDMC.C | TOL_IGR.H | 0.110 | 0.369 | |
| MGR.C | TOL_IGR.H | 0.180 | 0.149 | |
| SLA.C | TOL_IGR.H | -0.060 | 0.633 | |
| SSIZ.C | TOL_IGR.H | 0.020 | 0.856 | |
| TGER.C | TOL_IGR.H | 0.020 | 0.871 | |
| negXMID.C | TOL_IGR.H | 0.060 | 0.636 | |
| ASYM.F | TOL_IGR.H | 0.080 | 0.534 | |
| LA.F | TOL_IGR.H | 0.080 | 0.538 | |
| LDI.F | TOL_IGR.H | 0.040 | 0.763 | |
| LDMC.F | TOL_IGR.H | 0.120 | 0.321 | |
| MGR.F | TOL_IGR.H | 0.060 | 0.609 | |
| RESmT2.F | TOL_IGR.H | 0.090 | 0.452 | |
| SLA.F | TOL_IGR.H | -0.060 | 0.649 | |
| TOL_ASYM.F | TOL_IGR.H | -0.070 | 0.588 | |
| TOL_IGR.H | TOL_MGR.F | -0.170 | 0.157 | |
| TOL_IGR.H | TOL_negXMID.F | -0.070 | 0.590 | |
| negXMID.F | TOL_IGR.H | 0.150 | 0.211 | |
| ASYM.H | TOL_IGR.H | 0.110 | 0.369 | |

| Trait_1 | Trait_2 | corr.coef | p-value | |
|---|---|---|---|---|
| LA.H | TOL_IGR.H | 0.030 | 0.822 | |
| LDI.H | TOL_IGR.H | -0.050 | 0.703 | |
| LDMC.H | TOL_IGR.H | 0.030 | 0.836 | |
| MGR.H | TOL_IGR.H | 0.150 | 0.222 | |
| RESpT2.H | TOL_IGR.H | -0.100 | 0.415 | |
| SLA.H | TOL_IGR.H | -0.020 | 0.892 | |
| TOL_ASYM.H | TOL_IGR.H | 0.060 | 0.603 | |
| TOL_IGR.H | TOL_MGR.H | 0.070 | 0.593 | |
| TOL_IGR.H | TOL_negXMID.H | 0.040 | 0.724 | |
| negXMID.H | TOL_IGR.H | 0.130 | 0.291 | |
| IGR.C | TOL_IGR.H | -0.130 | 0.284 | |
| LTH.C | TOL_IGR.H | -0.090 | 0.475 | |
| NLEA.C | TOL_IGR.H | -0.130 | 0.276 | |
| RESmT1.C | TOL_IGR.H | -0.010 | 0.918 | |
| RESmT2.C | TOL_IGR.H | 0.180 | 0.143 | |
| RESpT1.C | TOL_IGR.H | -0.050 | 0.700 | |
| RESpT2.C | TOL_IGR.H | 0.060 | 0.634 | |
| IGR.F | TOL_IGR.H | 0.090 | 0.475 | |
| LTH.F | TOL_IGR.H | -0.010 | 0.912 | |
| NLEA.F | TOL_IGR.H | 0.000 | 0.984 | |
| TOL_IGR.F | TOL_IGR.H | 0.150 | 0.211 | |
| IGR.H | TOL_IGR.H | 0.600 | 0.000 | * |
| LTH.H | TOL_IGR.H | 0.030 | 0.801 | |
| NLEA.H | TOL_IGR.H | -0.060 | 0.623 | |

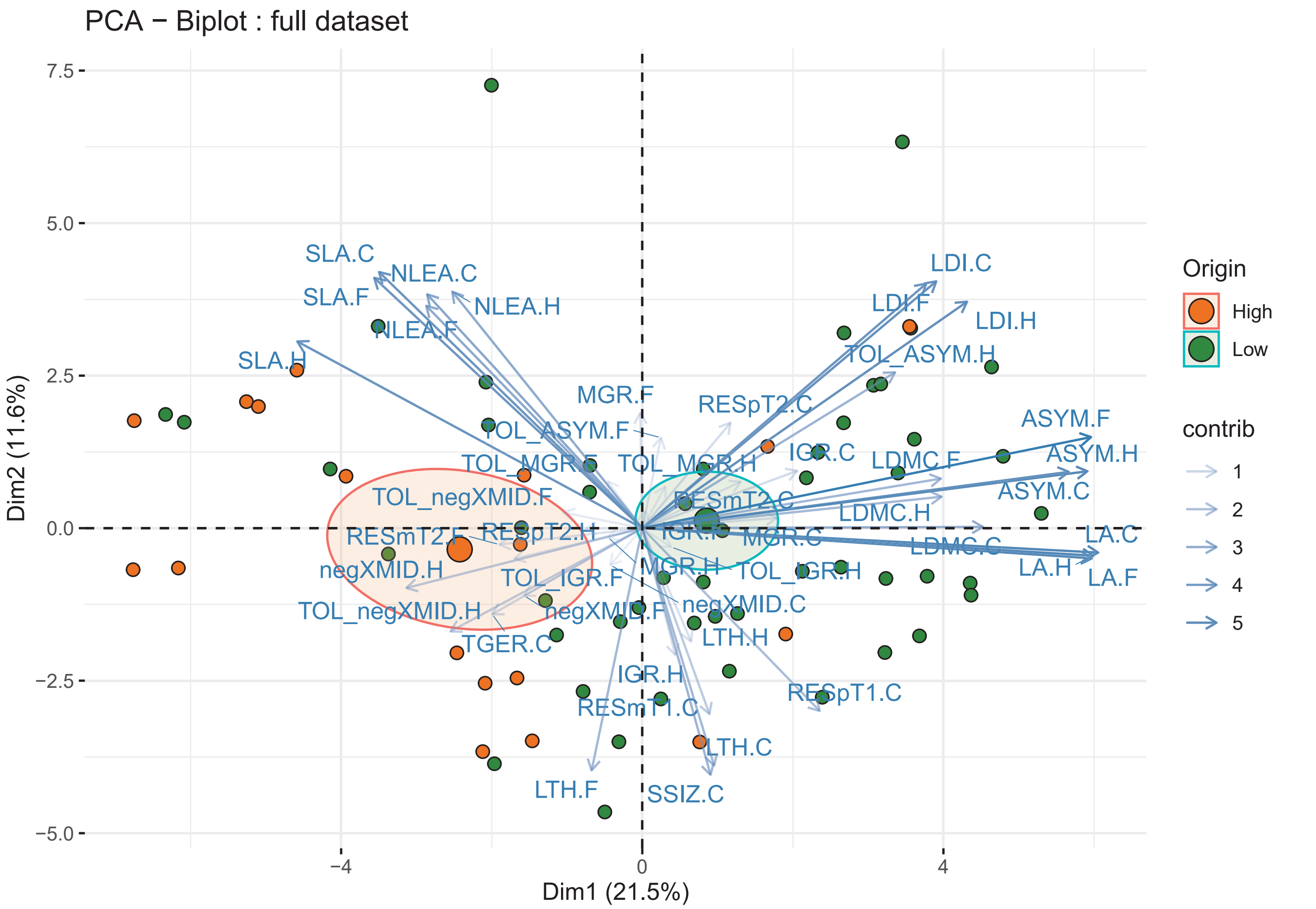
PCA − Biplot : full dataset
Dim1 (21.5%)
Dim2 (11.6%)
7.5
5.0
2.5
0.0
−2.5
−5.0
−4
0
4
SLA.C
NLEA.C
SLA.F
NLEA.H
NLEA.F
SLA.H
MGR.F
RESpT2.C
TOL_ASYM.F
TOL_MGR.F
TOL_MGR.H
IGR.C
TOL_negXMID.F
RESmT2.C
RESmT2.F
RESpT2.H
negXMID.H
TOL_IGR.F
MGR.H
TOL_IGR.H
TOL_negXMID.H
negXMID.F
negXMID.C
TGER.C
LTH.H
IGR.H
RESmT1.C
RESpT1.C
LTH.C
LTH.F
SSIZ.C
LDI.C
LDI.F
LDI.H
TOL_ASYM.H
ASYM.F
ASYM.H
LDMC.F
ASYM.C
LDMC.H
MGR.C
LDMC.C
LA.C
LA.H
LA.F
Origin
High
Low
contrib
1
2
3
4
5

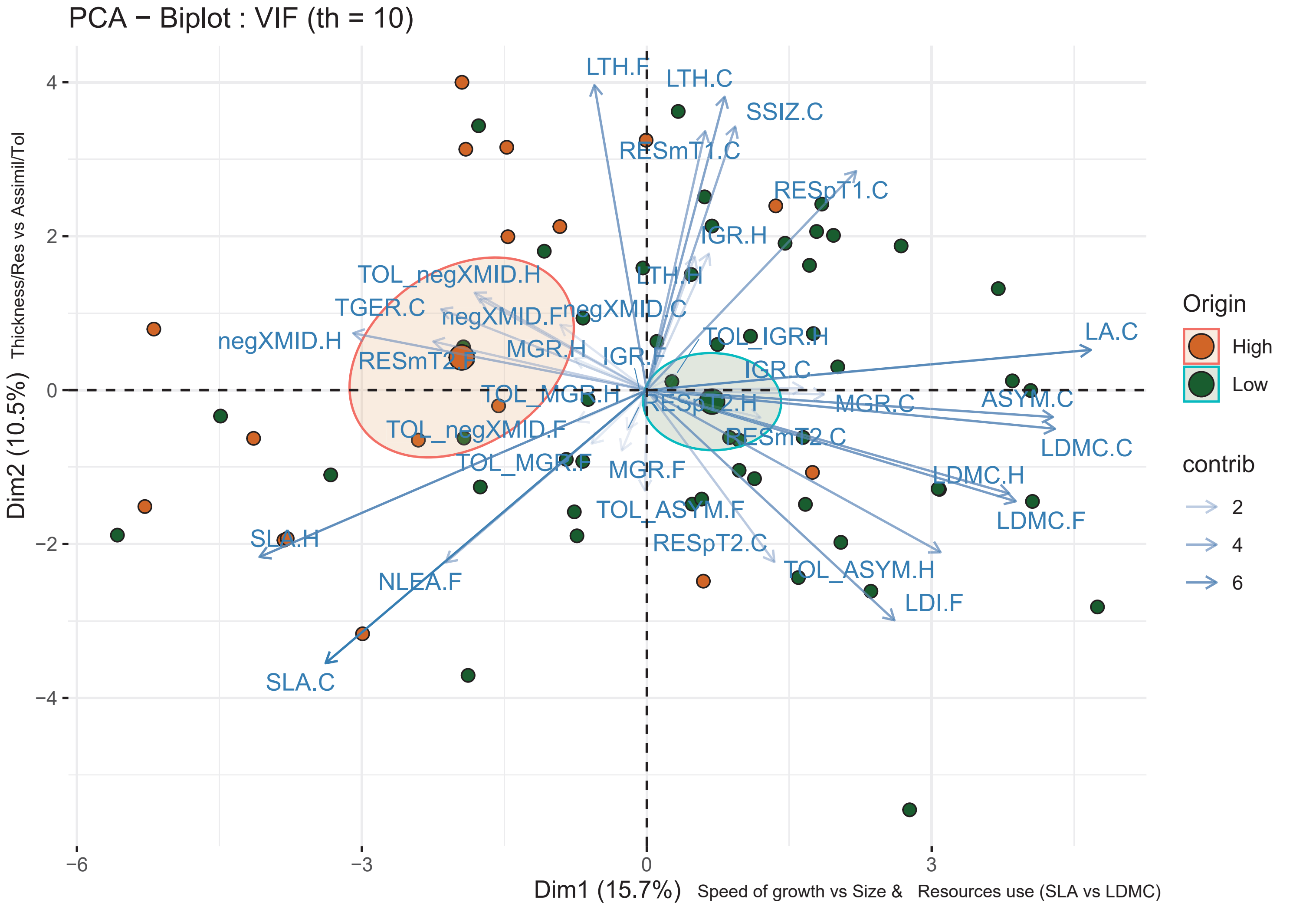
PCA − Biplot : VIF (th = 10)
Dim1 (15.7%)
Speed of growth vs Size & Resources use (SLA vs LDMC)
Dim2 (10.5%)
Thickness/Res vs Assimil/Tol
4
2
0
−2
−4
−6
−3
0
3
LTH.F
LTH.C
SSIZ.C
RESmT1.C
RESpT1.C
IGR.H
LTH.H
TOL_negXMID.H
TGER.C
negXMID.F
negXMID.C
negXMID.H
TOL_IGR.H
RESmT2.F
MGR.H
IGR.F
IGR.C
LA.C
TOL_MGR.H
RESpT2.H
MGR.C
ASYM.C
TOL_negXMID.F
RESmT2.C
LDMC.C
TOL_MGR.F
MGR.F
LDMC.H
TOL_ASYM.F
LDMC.F
SLA.H
RESpT2.C
TOL_ASYM.H
NLEA.F
LDI.F
SLA.C
Origin
High
Low
contrib
2
4
6

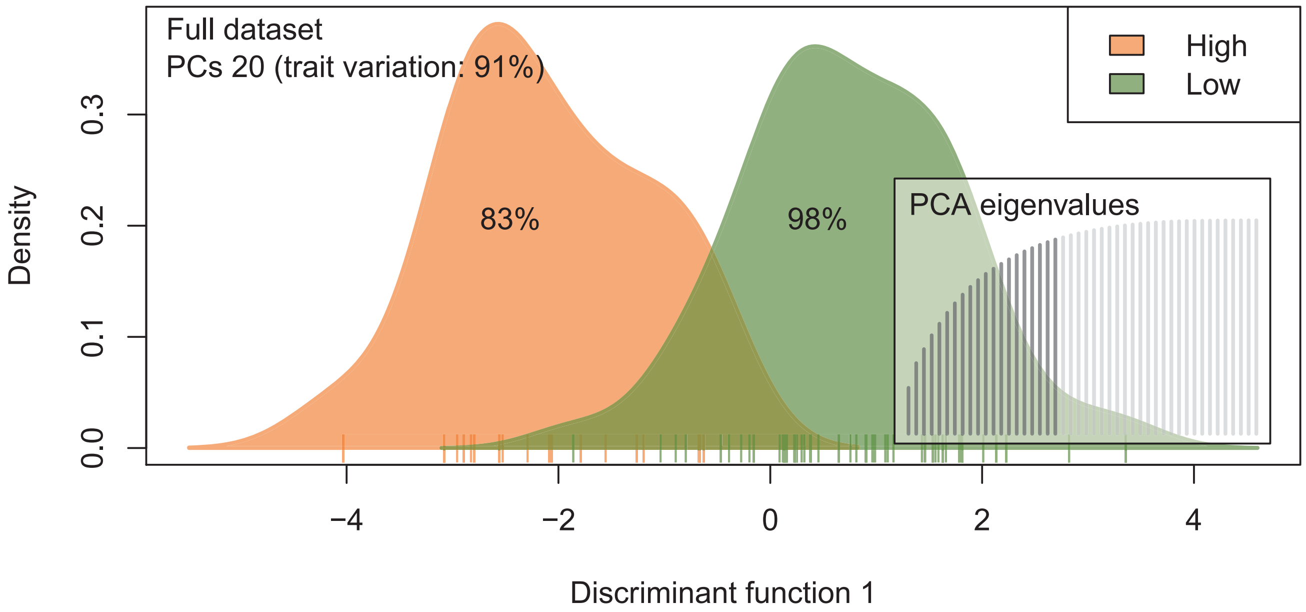
Full dataset
PCs 20 (trait variation: 91%)
High
Low
83%
98%
PCA eigenvalues
Density
0.0
0.1
0.2
0.3
−4
−2
0
2
4
Discriminant function 1

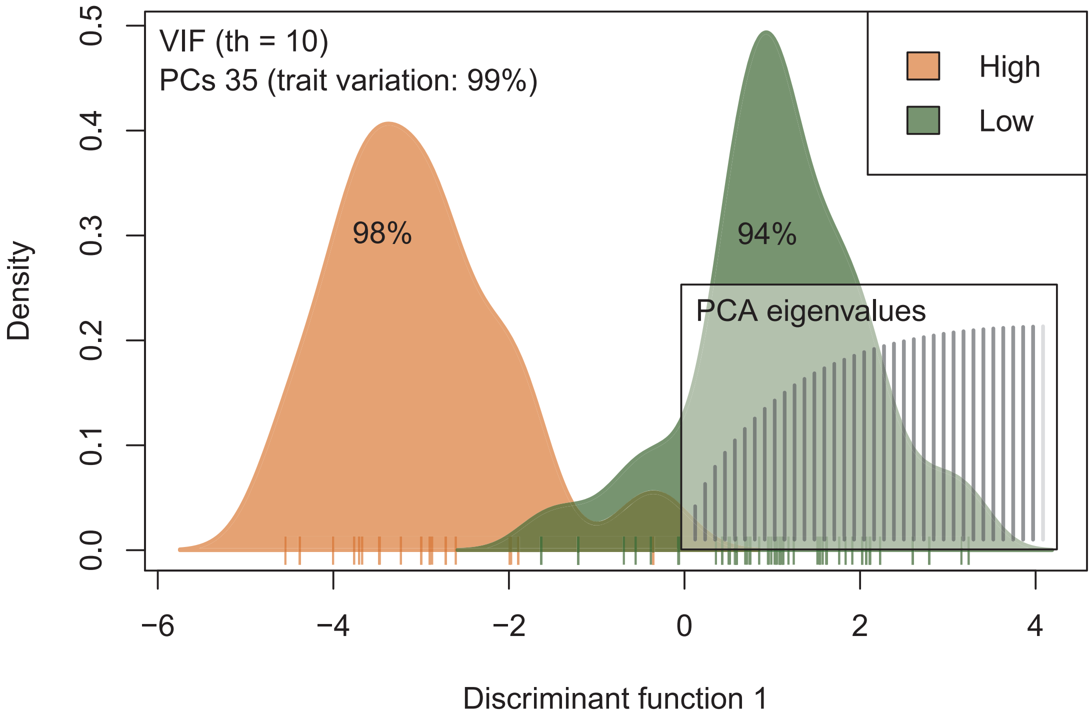
VIF (th = 10)
PCs 35 (trait variation: 99%)
High
Low
98%
94%
PCA eigenvalues
Density
0.0
0.1
0.2
0.3
0.4
0.5
−6
−4
−2
0
2
4
Discriminant function 1

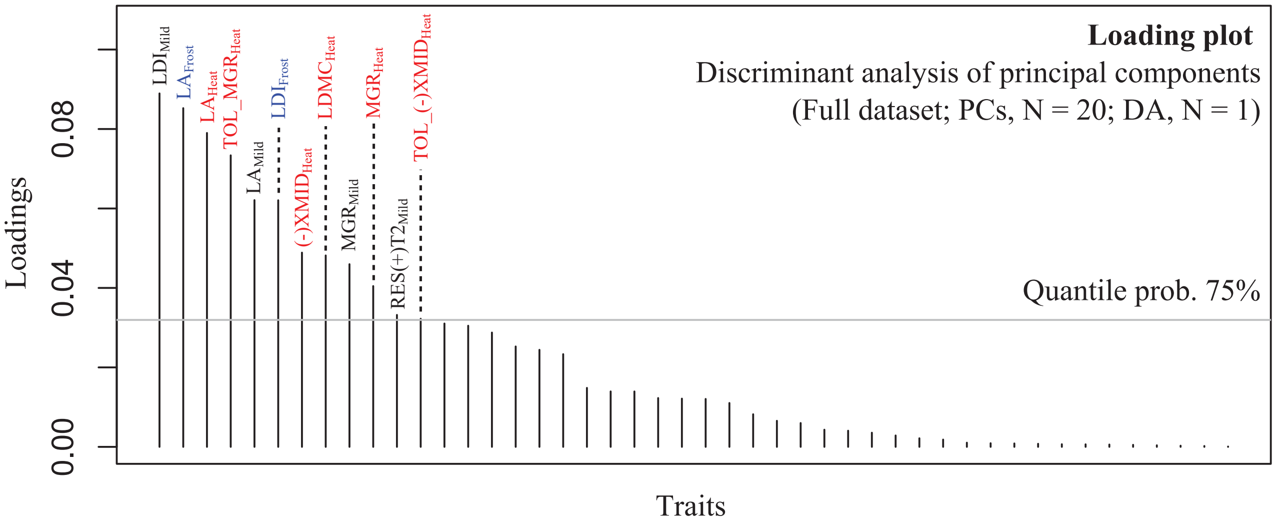
Loading plot
Discriminant analysis of principal components
(Full dataset; PCs, N = 20; DA, N = 1)
Quantile prob. 75%
Loadings
0.00
0.04
0.08
Traits
$LDI_{Mild}$
$LA_{Frost}$
$LA_{Heat}$
$TOL\_MGR_{Heat}$
$LA_{Mild}$
$LDI_{Frost}$
$(-)XMID_{Heat}$
$LDMC_{Heat}$
$MGR_{Mild}$
$MGR_{Heat}$
$RES(+)T2_{Mild}$
$TOL\_(-)XMID_{Heat}$

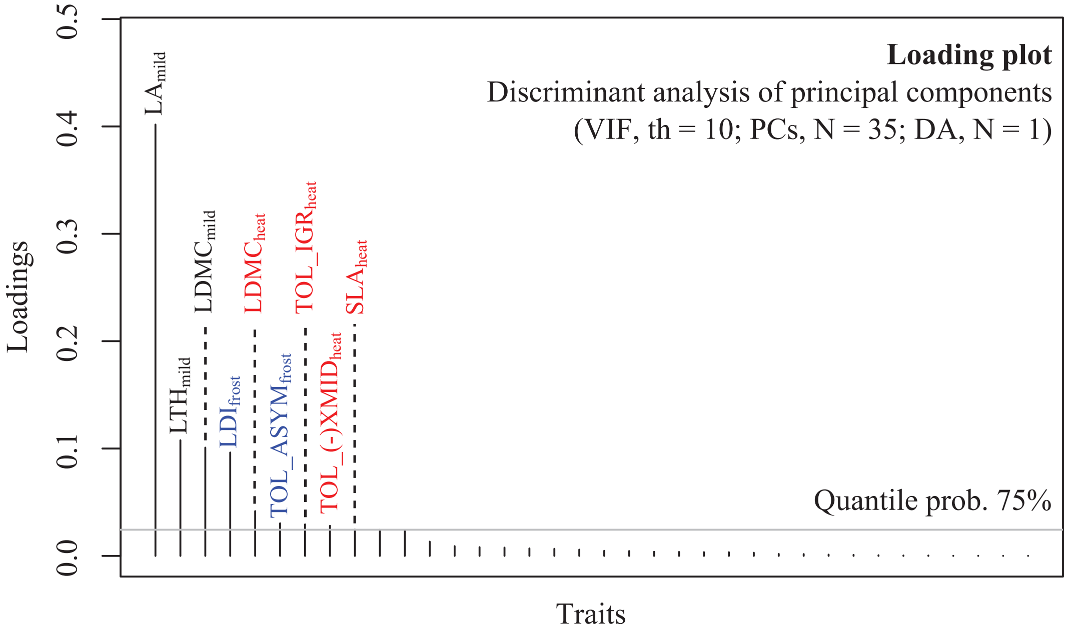
Loading plot
Discriminant analysis of principal components
(VIF, th = 10; PCs, N = 35; DA, N = 1)
Quantile prob. 75%
Loadings
0.0
0.1
0.2
0.3
0.4
0.5
Traits
$LA_{mild}$
$LTH_{mild}$
$LDMC_{mild}$
$LDI_{frost}$
$LDMC_{heat}$
$TOL\_ASYM_{frost}$
$TOL\_IGR_{heat}$
$TOL\_(-)XMID_{heat}$
$SLA_{heat}$

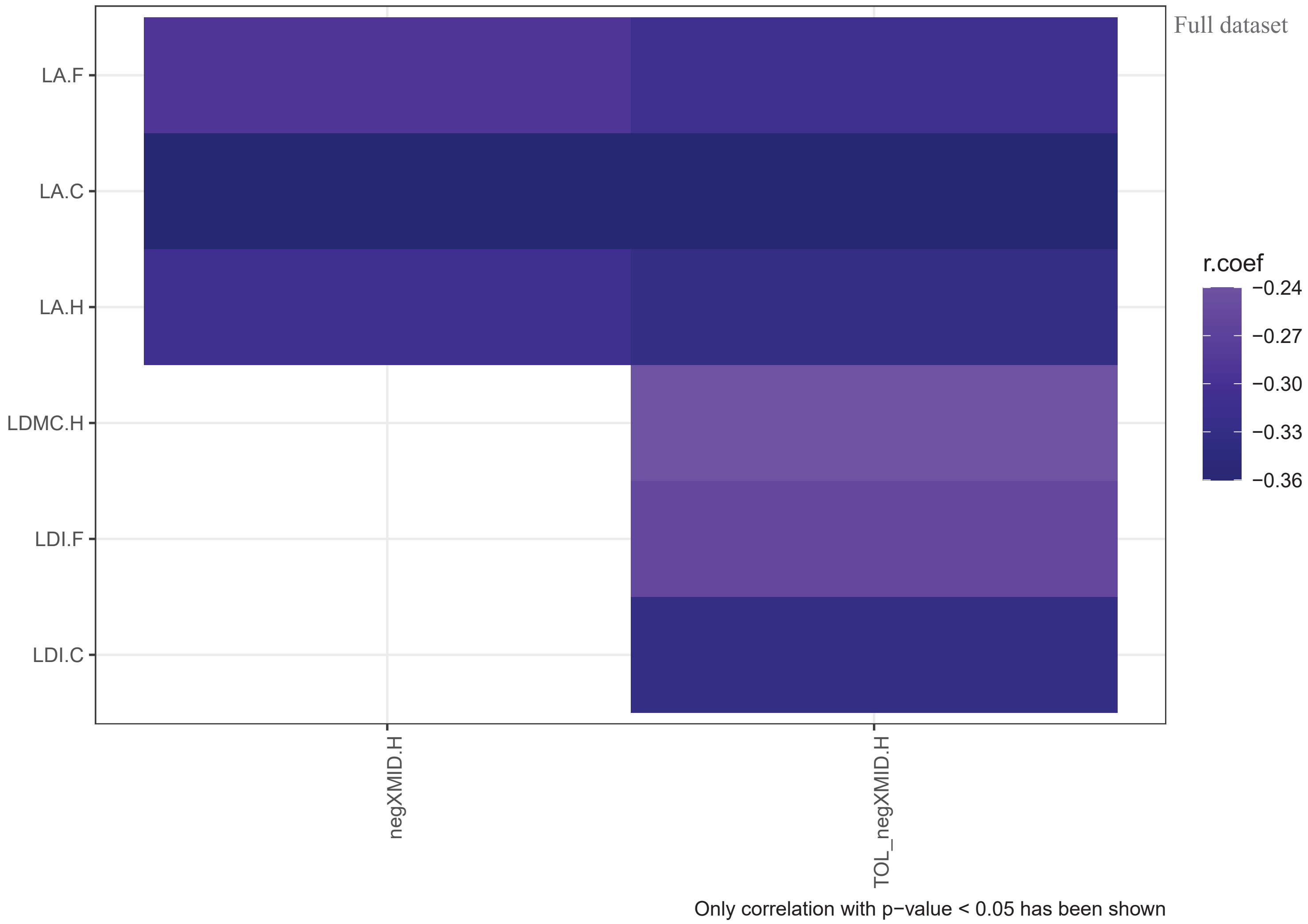
Full dataset
LA.F
LA.C
LA.H
LDMC.H
LDI.F
LDI.C
negXMID.H
TOL_negXMID.H
r.coef
−0.24
−0.27
−0.30
−0.33
−0.36
Only correlation with p−value < 0.05 has been shown

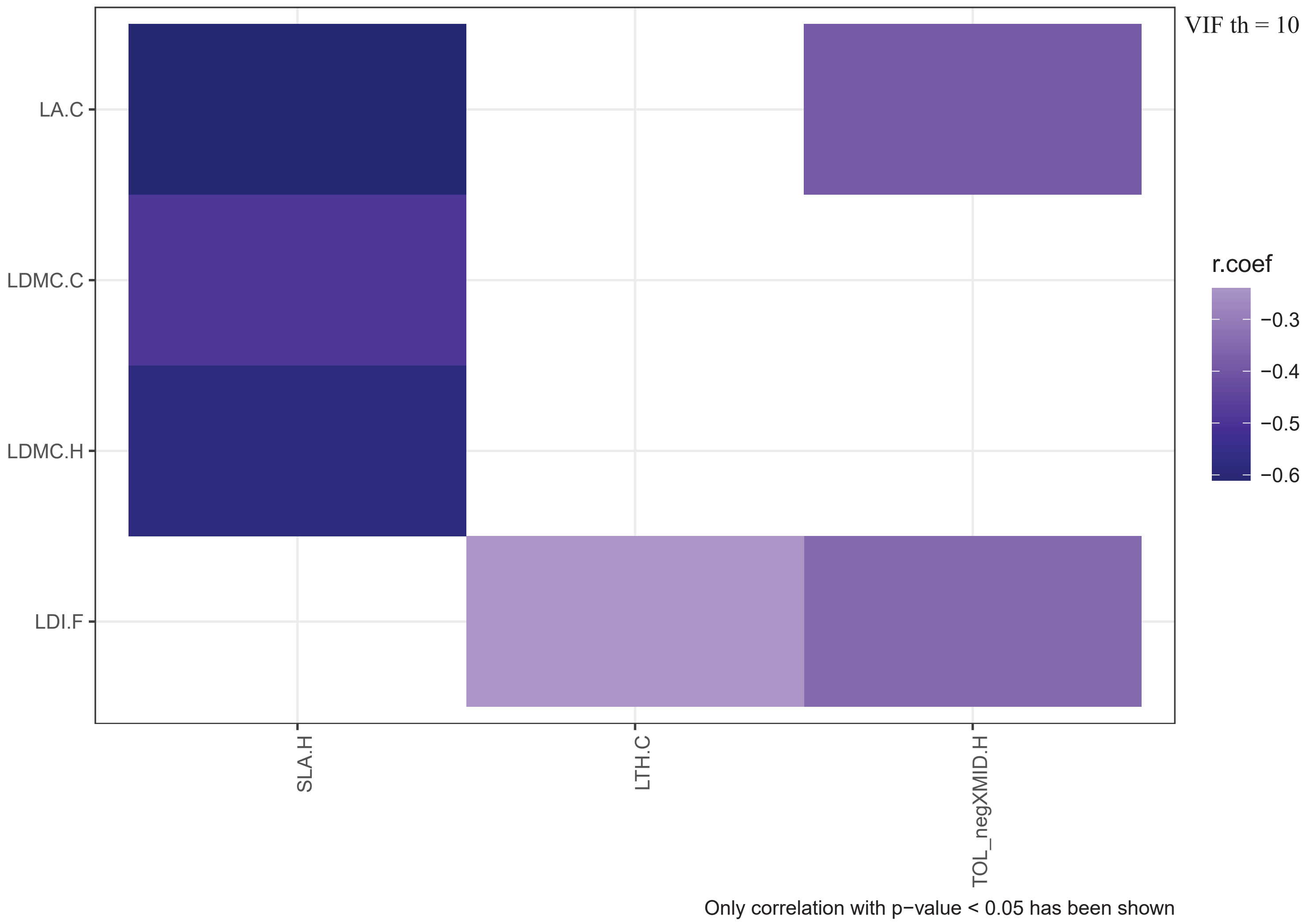
VIF th = 10
LA.C
LDMC.C
LDMC.H
LDI.F
SLA.H
LTH.C
TOL_negXMID.H
r.coef
−0.3
−0.4
−0.5
−0.6
Only correlation with p−value < 0.05 has been shown

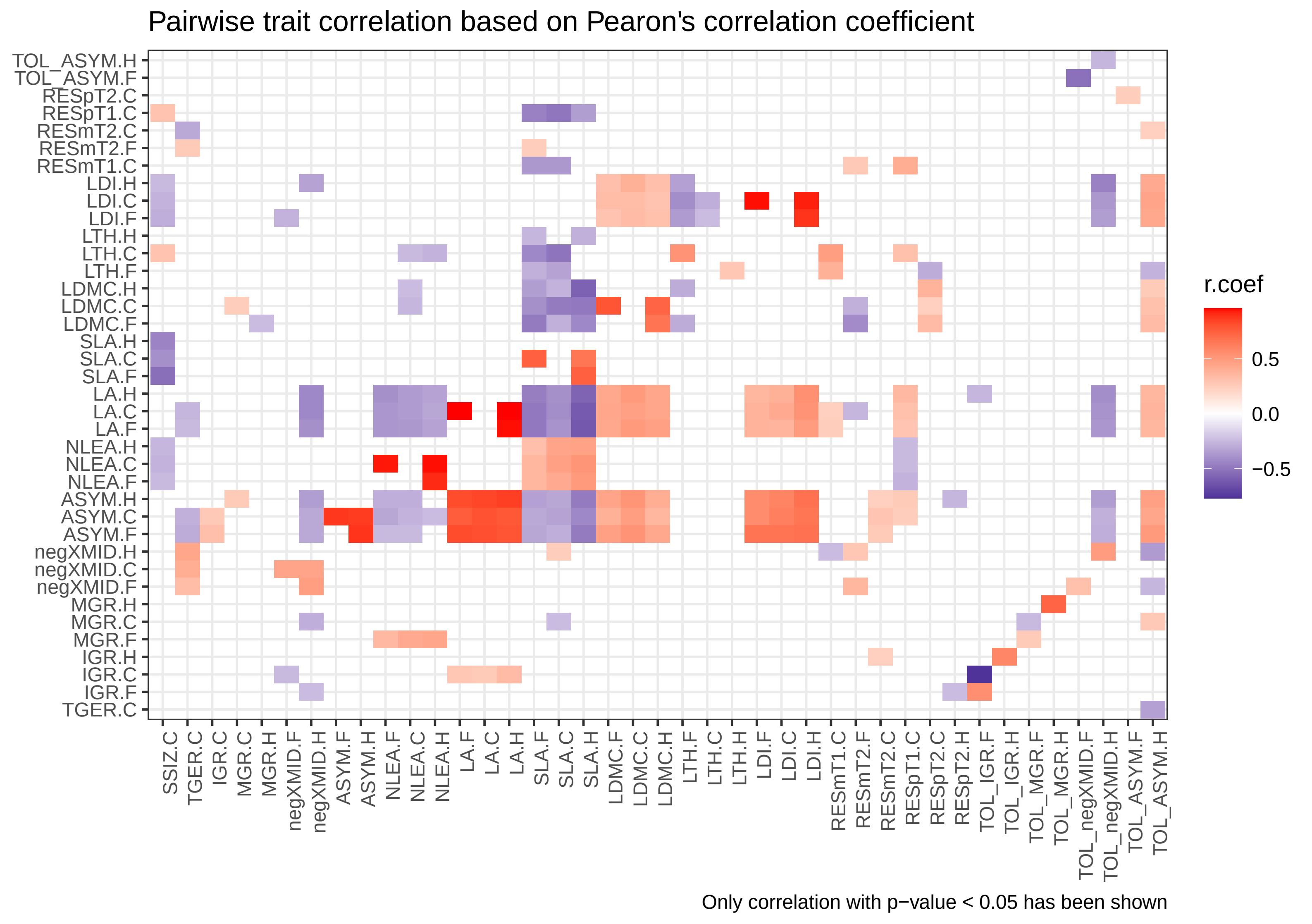

Pairwise trait correlation based on Pearon's correlation coefficient
TOL_ASYM.H
TOL_ASYM.F
RESpT2.C
RESpT1.C
RESmT2.C
RESmT2.F
RESmT1.C
LDI.H
LDI.C
LDI.F
LTH.H
LTH.C
LTH.F
LDMC.H
LDMC.C
LDMC.F
SLA.H
SLA.C
SLA.F
LA.H
LA.C
LA.F
NLEA.H
NLEA.C
NLEA.F
ASYM.H
ASYM.C
ASYM.F
negXMID.H
negXMID.C
negXMID.F
MGR.H
MGR.C
MGR.F
IGR.H
IGR.C
IGR.F
TGER.C
SSIZ.C
TGER.C
IGR.C
MGR.C
MGR.H
negXMID.F
negXMID.H
ASYM.F
ASYM.H
NLEA.F
NLEA.C
NLEA.H
LA.F
LA.C
LA.H
SLA.F
SLA.C
SLA.H
LDMC.F
LDMC.C
LDMC.H
LTH.F
LTH.C
LTH.H
LDI.F
LDI.C
LDI.H
RESmT1.C
RESmT2.F
RESmT2.C
RESpT1.C
RESpT2.C
RESpT2.H
TOL_IGR.F
TOL_IGR.H
TOL_MGR.F
TOL_MGR.H
TOL_negXMID.F
TOL_negXMID.H
TOL_ASYM.F
TOL_ASYM.H
r.coef
0.5
0.0
−0.5
Only correlation with p−value < 0.05 has been shown
